# The *N*-Terminal Carbamate is Key to High Cellular and Antiviral Potency for Boceprevir-Based SARS-CoV-2 Main Protease Inhibitors

**DOI:** 10.1101/2021.12.18.473330

**Authors:** Yugendar R. Alugubelli, Zhi Zachary Geng, Kai S. Yang, Namir Shaabani, Kaustav Khatua, Xinyu R. Ma, Erol C. Vatansever, Chia-Chuan Cho, Yuying Ma, Lauren Blankenship, Ge Yu, Banumathi Sankaran, Pingwei Li, Robert Allen, Henry Ji, Shiqing Xu, Wenshe Ray Liu

## Abstract

Boceprevir is an HCV NSP3 inhibitor that has been explored as a repurposed drug for COVID-19. It inhibits the SARS-CoV-2 main protease (M^Pro^) and contains an α-ketoamide warhead, a P1 β-cyclobutylalanyl moiety, a P2 dimethylcyclopropylproline, a P3 *tert*-butyl-glycine, and a P4 *N*-terminal *tert*-butylcarbamide. By introducing modifications at all four positions, we synthesized 20 boceprevir-based M^Pro^ inhibitors including PF-07321332 and characterized their M^Pro^ inhibition potency in test tubes (*in vitro*) and human host cells (*in cellulo*). Crystal structures of M^Pro^ bound with 10 inhibitors and antiviral potency of 4 inhibitors were characterized as well. Replacing the P1 site with a β-(S-2-oxopyrrolidin-3-yl)-alanyl (opal) residue and the warhead with an aldehyde leads to high *in vitro* potency. The original moieties at P2, P3 and the P4 *N*-terminal cap positions in boceprevir are better than other tested chemical moieties for high *in vitro* potency. In crystal structures, all inhibitors form a covalent adduct with the M^Pro^ active site cysteine. The P1 opal residue, P2 dimethylcyclopropylproline and P4 *N*-terminal *tert*-butylcarbamide make strong hydrophobic interactions with M^Pro^, explaining high *in vitro* potency of inhibitors that contain these moieties. A unique observation was made with an inhibitor that contains an P4 *N*-terminal isovaleramide. In its M^Pro^ complex structure, the P4 *N*-terminal isovaleramide is tucked deep in a small pocket of M^Pro^ that originally recognizes a P4 alanine side chain in a substrate. Although all inhibitors show high *in vitro* potency, they have drastically different *in cellulo* potency in inhibiting ectopically expressed M^Pro^ in human 293T cells. All inhibitors including PF-07321332 with a P4 *N*-terminal carbamide or amide have low *in cellulo* potency. This trend is reversed when the P4 *N*-terminal cap is changed to a carbamate. The installation of a P3 *O-tert*-butyl-threonine improves *in cellulo* potency. Three molecules that contain a P4 *N*-terminal carbamate were advanced to antiviral tests on three SARS-CoV-2 variants. They all have high potency with EC_50_ values around 1 μM. A control compound with a nitrile warhead and a P4 *N*-terminal amide has undetectable antiviral potency. Based on all observations, we conclude that a P4 *N*-terminal carbamate in a boceprevir derivative is key for high antiviral potency against SARS-CoV-2.

## ■ INTRODUCTION

COVID-19 is the currently ongoing pandemic that is caused by the coronavirus SARS-CoV-2 and has no effective medication for treatment. To address this emergency, a large variety of drug repurposing research has been conducted to identify approved medications that might be potentially used as COVID-19 treatments.^1–5^ Significant part of this research has been targeting the SARS-CoV-2 main protease (M^Pro^).^6–12^ M^Pro^ is a peptide fragment of two translation products pp1a and pp1ab of the SARS-COV-2 RNA genome after the virus infects human cells. Both pp1a and pp1ab are very large polypeptides that need to undergo proteolytic hydrolysis to form a number of nonstructural proteins (nsps). These nsps are essential for the virus to its genome replication in host cells, evasion from the host immune system, and packaging of new virions for infection of new host cells.^13^ Intervention of proteolytic hydrolysis of pp1a and pp1ab is considered as a viable approach to stop SARS-CoV-2 infection. There are two internal peptide fragments from pp1a and pp1b that function as cysteine proteases to hydrolyze all nsps. One is M^Pro^ and the other papain-like protease (PL^Pro^). As the main protease for this proteolytic process, M^Pro^ processes the majority of nsps. It is also more conserved than PL^Pro^. M^Pro^ genes in SARS-CoV and SARS-CoV-2 share 96% sequence identity.^2^ Targeting M^Pro^ for drug discovery will likely generate broad spectrum antivirals for coronaviruses. So far, a number of approved small molecule drugs have been confirmed as potent M^Pro^ inhibitors. One of which is boceprevir.^14–16^ Boceprevir is a peptidyl inhibitor of HCV NSP3.^17^ HCV NSP3 is a serine protease. Boceprevir contains an α-ketoamide warhead that forms a reversible covalent adduct with its active site serine for high inhibition potency.^18^ Besides the α-ketoamide warhead, boceprevir contains a P1 β-cyclobutylalanyl moiety, a P2 dimethylcyclopropylproline, a P3 *tert*-butylglycine, and a P4 *N*-terminal *tert*-butylcarbamide. Boceprevir has been shown as a potent inhibitor of M^Pro^ in multiple publications.^1, 14–16^ Its interactions with M^Pro^ have also been structurally characterized using X-ray protein crystallography.^14–16, 19, 20^ Although its discovery as a potent M^Pro^ inhibitor created excitement about its potential use as a COVID-19 treatment, boceprevir has moderate antiviral activity against SARS-CoV-2 and displays very low potency to inhibit M^Pro^ in a human cell host.^14, 15, 21^ In this work, we report a systematic structure-activity relationship (SARS) study of boceprevir-based molecules for improved cellular and antiviral potency against the SARS-CoV-2 M^Pro^.

## ■ RESULTS

### The Design and Synthesis of MPI29-47 and PF-07321332

In total, we designed 19 inhibitors as shown in Figure 1. The Pfizer compound PF-07321332 is included in our analysis due to its close similarity to boceprevir.^22^ M^Pro^ contains four binding pockets in its active site to interact with the P1, P2, P4, and P3’ amino acid residues in a substrate. The P1 residue in a substrate is strictly glutamine. Past efforts in developing M^Pro^ inhibitors have been either using a P1 β-(S-2oxopyrrolidin-3-yl)-alanyl (opal) residue in a peptidyl inhibitor or a pyridine moiety to engage the P1 binding pocket of M^Pro^ for strong binding.^6, 8 15, 23–30^ Boceprevir has a P1 β-cyclobutylalanyl moiety that displays loose binding to the M^Pro^ P1 binding pocket in its M^Pro^ complex structure,^14–16, 19, 20^ which explains its relatively moderate M^Pro^ inhibition potency. In our designed boceprevir derivatives, this site is replaced by opal. In M^Pro^-boceprevir complex structures, the P2 dimethylcyclopropylproline binds neatly to the M^Pro^ P2 binding pocket. Due to its nice fit to the M^Pro^ P2 binding pocket, this site is maintained as dimethylcyclopropylproline in most designed inhibitors. In a small number of inhibitors, other residues such as (*S*)-5-azaspiro[2,4]heptane-6-carboxylic acid in MPI37, neopentylglycine in MPI38, β-cyclopropylalanine in MPI39, and cyclohexylalanine in MPI45 at the P2 site are introduced. A previous study has indicated that neopentylglycine, β-cyclopropylalanine and cyclohexylalanine at this site lead to either high M^Pro^ inhibition potency in test tubes (*in vitro*) or in human host cells that express M^Pro^ (*in cellulo*)^31^ The P3 site is either maintained as *tert*-butylglycine or replaced by *O-tert*-butylthreonine. *tert*-Butylglycine at the P3 site in a peptidyl inhibitor is known to generate high *in vitro* potency and *O-tert*-butylthreonine is known to cause high *in cellulo* and antiviral potency.^21, 32^ Boceprevir has a P4 *N*-terminal *tert*-butylcarbamide cap. This site is either maintained as a carbamide or replaced by a carbamate or amide.

**Figure 1.**
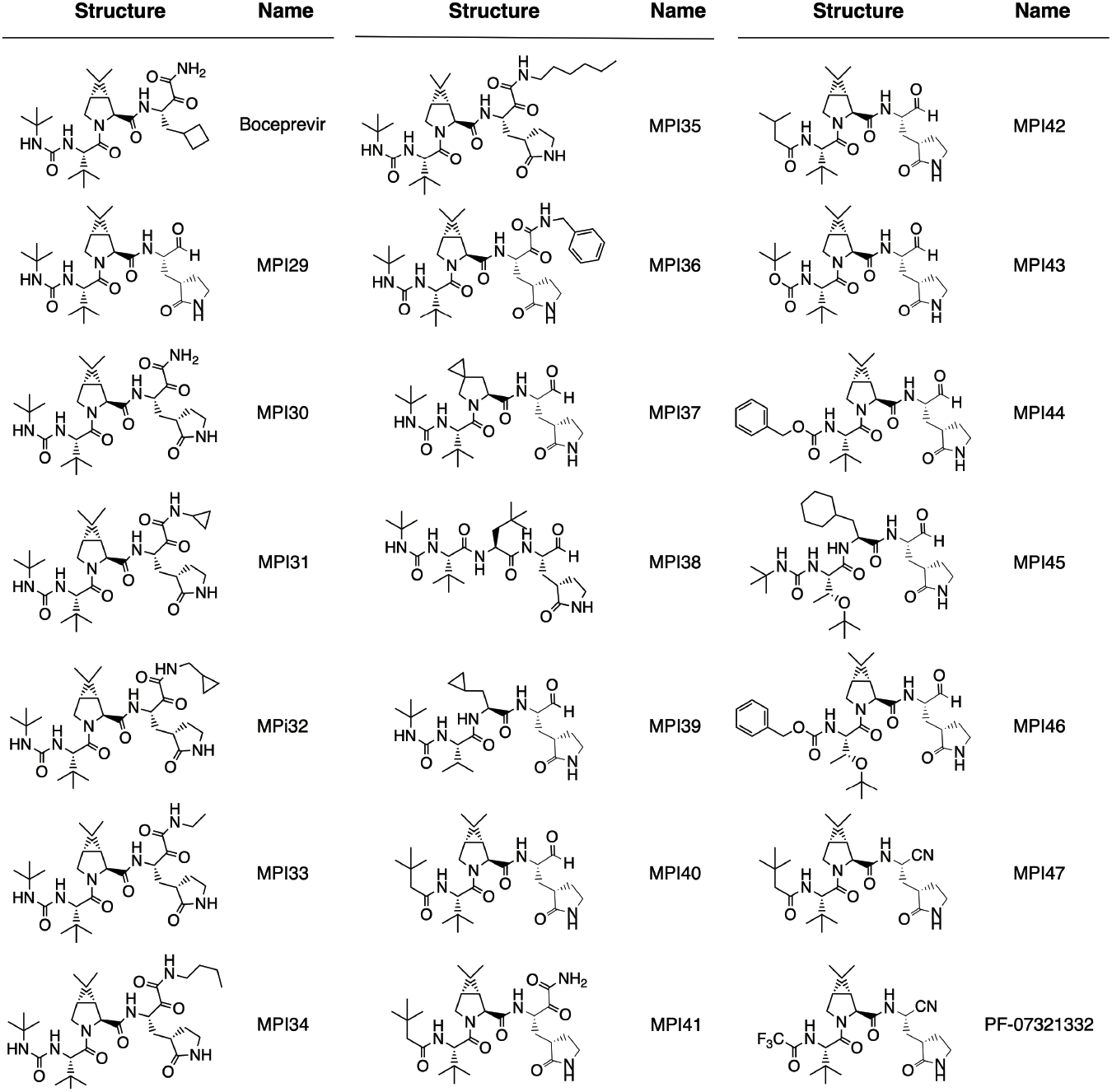
Structures of boceprevir, MPI29-47 and PF-07321332.

Boceprevir has an α-ketoamide warhead that covalently interacts with the M^Pro^ active site cysteine C145 to generate a hemithioacetal intermediate. Since the majority of currently known peptidyl inhibitors of M^Pro^ have an aldehyde warhead, we replaced the α-ketoamide warhead with an aldehyde in some designed inhibitors. We also tested nitrile at this position since PF-07321332 and some other developed M^Pro^ inhibitors have this warhead.^22, 33^ An advantage of the α-ketoamide warhead is its allowance of appending additional chemical moieties at its α-ketoamide nitrogen. All aldehyde- and nitrile-containing inhibitors will leave the M^Pro^ P3’ binding pocket empty when they bind to M^Pro^. A chemical appendage at the α-ketoamide nitrogen can potentially reach this pocket for additional interactions and therefore lead to high affinity to M^Pro^. MPI31-36 were designed for this purpose. The base molecule of MPI31-36 is MPI30 that is different from boceprevir only at the P1 residue. Please note that MPI30 is a previously developed M^Pro^ inhibitor with a name as ML1000, though the structural characterization of its complex with M^Pro^ has not been reported.^34^ We follow typical peptide coupling chemistry to synthesize all inhibitors. In general, the synthesis started from the P4 *N*-terminal cap to the *C*-terminal warhead. All compounds were strictly characterized to ensure their purity. Synthetic details and characterizations are provided in the supplementary information.

### Kinetic characterizations of MPI29-47 and PF-07321332 on their *in vitro* enzymatic inhibition potency

In a previous drug repurposing project, we established a protocol to characterize M^Pro^ inhibitors.^12^ In this protocol, a fluorogenic substrate Sub3 is used. We followed this protocol to characterize *in vitro* M^Pro^ inhibition potency for all synthesized inhibitors by determining their IC_50_ values. To conduct the assay, we preincubated 20 nM M^Pro^ with varied concentrations of an inhibitor for 30 min before 10 μM Sub3 was added and the fluorescent product formation was recorded in a fluorescence plate reader. All inhibitors exhibited well defined inhibition curves that started from 100% activity without an inhibitor and reached to almost total inhibition when 10 μM of an inhibitor was provided. Data are presented in Figure 2. We fit all collected data to a four-parameter variable slope inhibition equation in GraphPad 9.0 to obtain IC_50_ values for all inhibitors. Determined IC_50_ values are presented in Table 1. As shown in Table 1, MPI29 has the highest *in vitro* potency with an IC_50_ value as 9.3 nM. Since we used 20 nM M^Pro^ for the assay, this IC_50_ value has reached the detection limit. Real *in vitro* potency of MPI29 is likely higher than what the determined IC_50_ value indicates. MPI29 has an aldehyde warhead. MPI30 is different from MPI29 only at the warhead position. It has an α-ketoamide instead. MPI30 has a determined IC_50_ value as 40 nM. This value is similar to that reported by Westberg *et al*.^34^ This lower potency than MPI29 is expected since α-ketoamide is less chemically reactive than aldehyde toward a nucleophile. However, MPI30 is 100-fold more potent than boceprevir indicating the essential role of the P1 opal residue in improving interactions with M^Pro^. MPI31-36 all have much higher *in vitro* IC_50_ values than MPI30. Apparently adding a chemical appendage to the α-ketoamide nitrogen leads to less favored interactions with M^Pro^. Interestingly, MPI33 that has the smallest appendage has an *in vitro* IC_50_ value closest to MPI30. Although the benzyl appendage in MPI36 is bigger than that in MPI32-35, it is less detrimental to the *in vitro* potency indicating that the benzyl group possibly involves some favorable interactions with M^Pro^ in comparison to other *N*-substituents.

**Figure 2.**
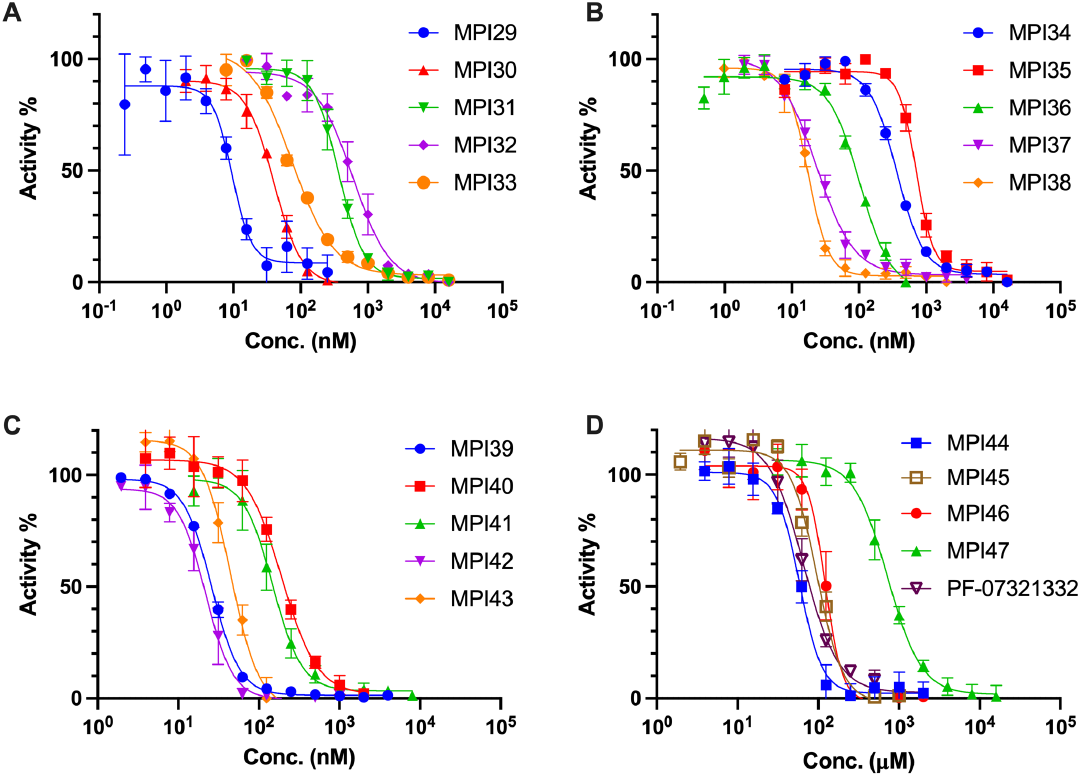
Inhibition curves of MPI29-47 and PF-07321332 on M^Pro^. Triplicate experiments were performed for each compound. For all experiments, 20 nM M^Pro^ was incubated with an inhibitor for 30 min before 10 *μ*M Sub3 was added. The M^Pro^-catalyzed Sub3 hydrolysis rate was determined by measuring linear increase of product fluorescence (Ex: 336 nm /Em: 455 nm) for 5 min.

**Table 1:**
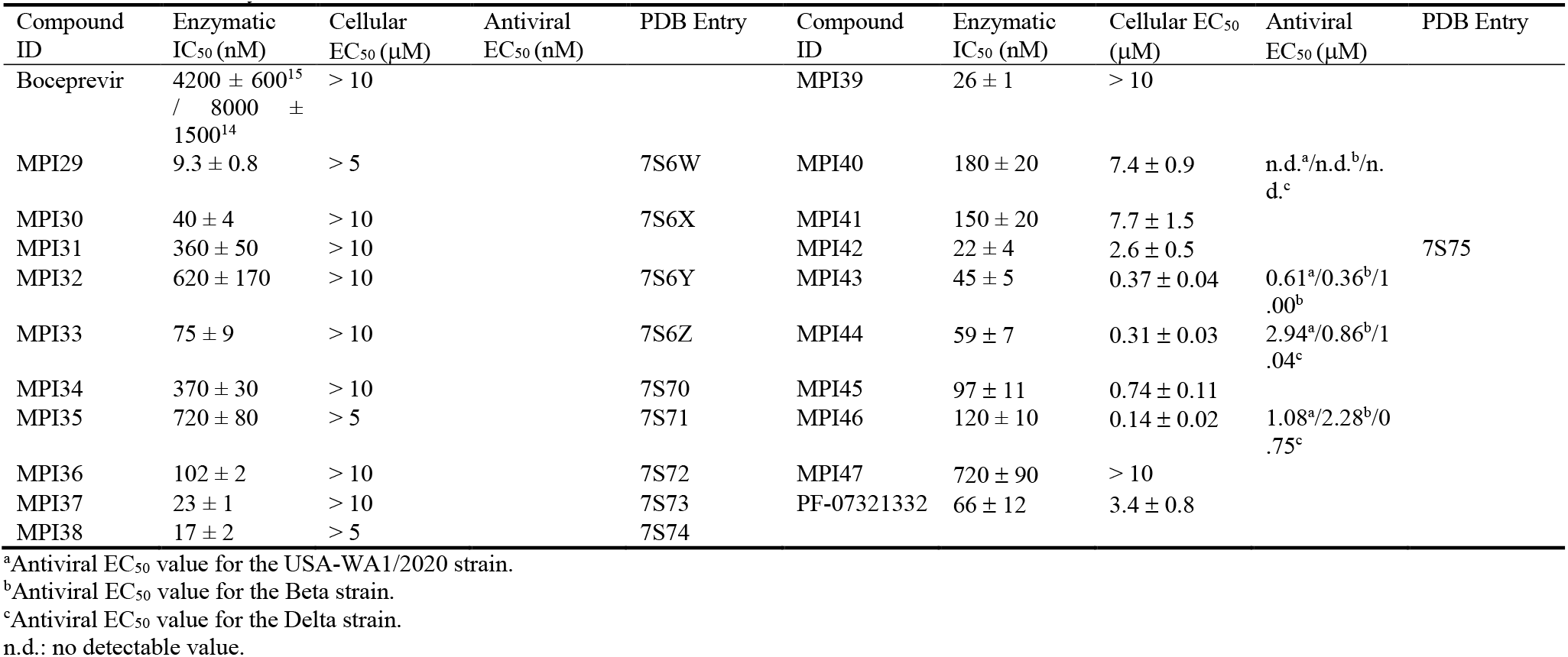
Determined enzymatic IC_50_, cellular EC_50_, and antiviral EC_50_ values of M^Pro^ inhibitors

| Compound ID | Enzymatic IC<sub>50</sub> (nM) | Cellular EC<sub>50</sub> (μM) | Antiviral EC<sub>50</sub> (nM) | PDB Entry | Compound ID | Enzymatic IC<sub>50</sub> (nM) | Cellular EC<sub>50</sub> (μM) | Antiviral EC<sub>50</sub> (μM) | PDB Entry |
| --- | --- | --- | --- | --- | --- | --- | --- | --- | --- |
| Boceprevir | 4200 ± 600<sup>15</sup><br>/ 8000 ± 1500<sup>14</sup> | > 10 |  |  | MPI39 | 26 ± 1 | > 10 |  |  |
| MPI29 | 9.3 ± 0.8 | > 5 |  | 7S6W | MPI40 | 180 ± 20 | 7.4 ± 0.9 | n.d.<sup>a</sup>/n.d.<sup>b</sup>/n.d.<sup>c</sup> |  |
| MPI30 | 40 ± 4 | > 10 |  | 7S6X | MPI41 | 150 ± 20 | 7.7 ± 1.5 |  |  |
| MPI31 | 360 ± 50 | > 10 |  |  | MPI42 | 22 ± 4 | 2.6 ± 0.5 |  | 7S75 |
| MPI32 | 620 ± 170 | > 10 |  | 7S6Y | MPI43 | 45 ± 5 | 0.37 ± 0.04 | 0.61<sup>a</sup>/0.36<sup>b</sup>/1.00<sup>b</sup> |  |
| MPI33 | 75 ± 9 | > 10 |  | 7S6Z | MPI44 | 59 ± 7 | 0.31 ± 0.03 | 2.94<sup>a</sup>/0.86<sup>b</sup>/1.04<sup>c</sup> |  |
| MPI34 | 370 ± 30 | > 10 |  | 7S70 | MPI45 | 97 ± 11 | 0.74 ± 0.11 |  |  |
| MPI35 | 720 ± 80 | > 5 |  | 7S71 | MPI46 | 120 ± 10 | 0.14 ± 0.02 | 1.08<sup>a</sup>/2.28<sup>b</sup>/0.75<sup>c</sup> |  |
| MPI36 | 102 ± 2 | > 10 |  | 7S72 | MPI47 | 720 ± 90 | > 10 |  |  |
| MPI37 | 23 ± 1 | > 10 |  | 7S73 | PF-07321332 | 66 ± 12 | 3.4 ± 0.8 |  |  |
| MPI38 | 17 ± 2 | > 5 |  | 7S74 |  |  |  |  |  |

<sup>a</sup>Antiviral EC<sub>50</sub> value for the USA-WA1/2020 strain.<sup>b</sup>Antiviral EC<sub>50</sub> value for the Beta strain.<sup>c</sup>Antiviral EC<sub>50</sub> value for the Delta strain.
n.d.: no detectable value.

MPI37 is different from MPI29 at the P2 residue. It has a determined *in vitro* IC_50_ value as 23 nM. Although replacing the P2 dimethylcyclopropylproline with (*S*)-5-azaspiro[2,4]heptane-6-carboxylic acid leads to less favorable interactions with M^Pro^, the effect is not dramatic. A similar observation was made with MPI38 that has a P2 neopentylglycine. MPI39 have both P2 and P3 sites different from MPI29. Both residues in MPI39 have been shown in a previous study favoring interactions with M^Pro^.^31^ Accordingly, we notice that MPI39 has relatively high *in vitro* potency. MPI40, MPI42 and MPI43 are different from MPI29 only in their P4 *N*-terminal cap. MPI40 has a P4 *N*-terminal amide that is different from MPI29 only at one atom position. However, this slight change leads to about 20-fold loss of *in vitro* potency. Intriguingly, removing one methyl group from the P4 *N*-terminal cap of MPI40, which leads to MPI42, improves *in vitro* potency for about 9 folds. Another interesting observation is on MPI43. MPI43 has a P4 *N*-terminal carbamate and is different from MPI29 and MPI40 only at the carbamate oxygen position. It has *in vitro* potency between MPI29 and MPI40. Therefore, switching a P4 *N*-terminal carbamide nitrogen to oxygen is less detrimental to *in vitro* potency than switching it to methylene. MPI41 is an α-ketoamide derivative of MPI40. It has *in vitro* potency surprisingly similar to MPI40 although α-ketoamide is expected to be less reactive than aldehyde. MPI44 is different from MPI43 at its P4 *N*-terminal *O*-alkyl group. It has a benzyl group instead of a *tert*-butyl group as in MPI43. Although two *O*-alkyl groups are structurally different, MPI43 and MPI44 have similar *in vitro* potency. In our previous work, we noticed that an *O*-benzyl group at this position loosely binds M^Pro^. Similar *in vitro* potency between MPI43 and MPI44 indicates that an *O-tert*-butyl group at this position might involve loose interactions with M^Pro^ as well. In our previous work, we discovered that MPI8, a peptidyl aldehyde has the highest *in cellulo* and antiviral potency among all inhibitors that we synthesized.^21, 31^ Due to the high *in vitro* potency of MPI29, we created chimera inhibitors MPI45 and MPI46 that integrate structural components from both MPI8 and MPI29. MPI45 is a chimera in which the P4 *N*-terminal cap of MPI29 is fused to the rest of MPI8 and MPI46 is a product made by switching the P2 residue in MPI8 to the one in MPI29. Both MPI45 and MPI46 have *in vitro* potency similar to MPI8 with an IC_50_ value about 100 nM.^32^ MPI47 is different from MPI40 only at its warhead. It has an *in vitro* IC_50_ value as 720 nM. Apparently switching the warhead to nitrile from either aldehyde or α-ketoamide leads to significant loss of *in vitro* potency. Although PF-07321332 and MPI47 are structurally similar, PF-07321332 has a more than 10-folder lower IC_50_ value than MPI47. A likely explanation is that the P4 N-terminal trifluoroacetamide cap in PF-07321332 involves unique interactions with M^Pro^ that do not exist for MPI47.

### X-Ray Crystallography analysis of M^Pro^ bound with 10 inhibitors

In order to structurally characterize our developed inhibitors in their complexes with M^Pro^. We crystalized M^Pro^ in its apo form and then soaked obtained crystals with our designed inhibitors for X-ray crystallography analysis. Using this approach, we successfully determined 10 M^Pro^-inhibitor complex structures to high resolutions. These inhibitors include MPI29, MPI30, MPI32-38, and MPI42. PDB entries for these M^Pro^-inhibitor complex structures are summarized in Table 1. In the active sites of all M^Pro^-inhibitor complexes, electron density is well shaped for unambiguous modeling of inhibitors except for MPI36 at the *N*-benzyl group of its α-ketoamide (Figure 3A). A covalent bond between C145 of M^Pro^ and the warhead of an inhibitor is clearly visible in all structures.

**Figure 3.**
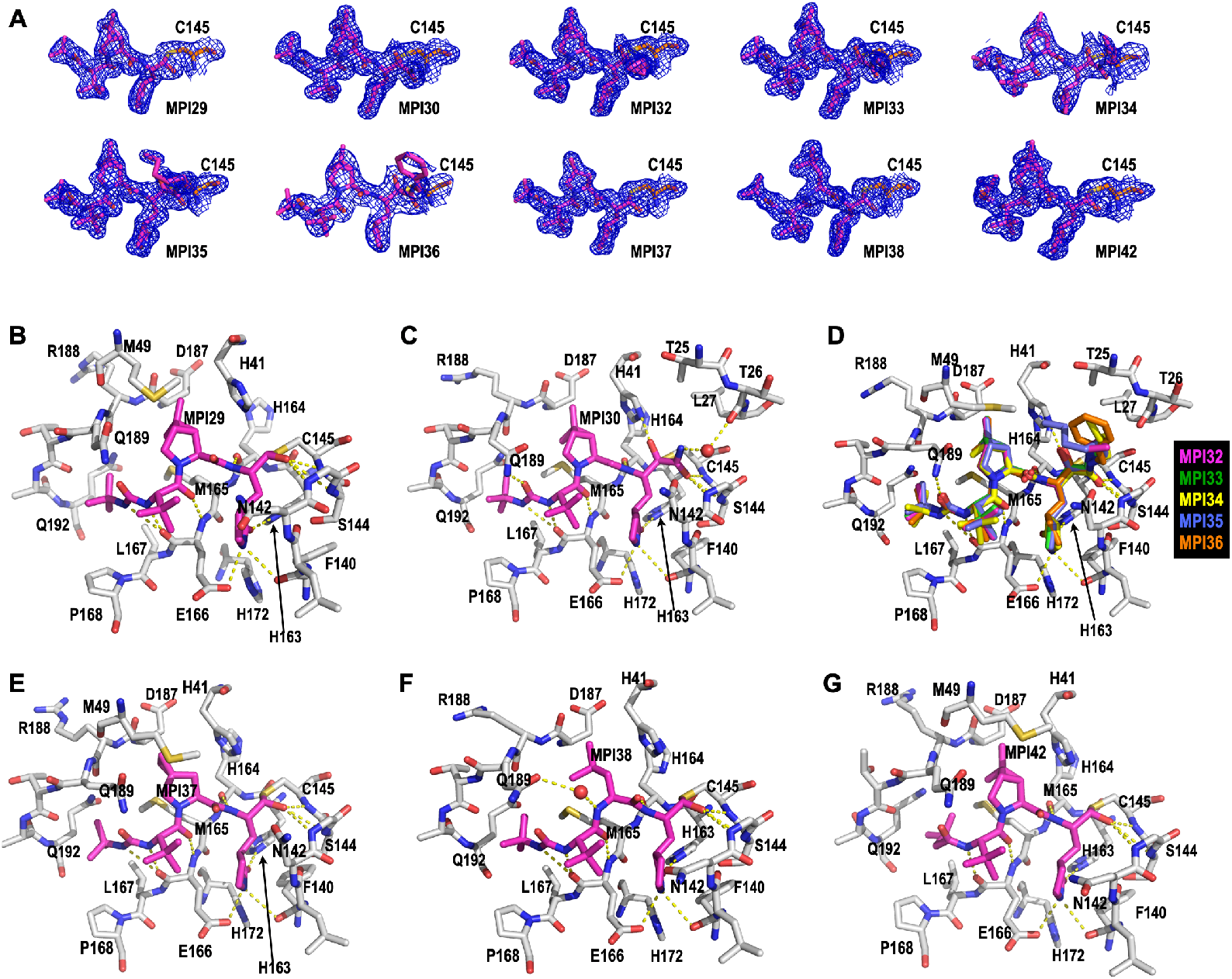
Crystal structures of M^Pro^ bound with 10 MPIs. (A) Contoured 2*F_o_-F_c_* maps at the 1*σ* level around 10 MPIs and C145 in the active site of M^Pro^. The active site structures for M^Pro^ bound with (B) MPI29, (C) MPI30, (D) MPI32-36, (E) MPI37, (F) MPI38, and (G) MPI42. Dashed yellow lines between inhibitors and M^Pro^ are potential hydrogen bonds.

In the M^Pro^-MPI29 complex as shown in Figure 3B, MPI29 forms a number of hydrogen bonds with M^Pro^. The amide of the P1 opal side chain lactam forms three hydrogen bonds with M^Pro^ residues including F140 at its backbone oxygen, H163 at one of its imidazole nitrogen atoms and E166 at its side chain carboxylate. There are two hydrogen bonds generated between two backbone amides of MPI29 and M^Pro^ residues including H164 at its backbone oxygen and E166 at its backbone nitrogen. The P4 *N*-terminal carbamide cap of MPI29 uses its two nitrogen atoms to form hydrogen bonds with the backbone oxygen of M^Pro^ E166. In previous published structures of M^Pro^ bound with peptidyl aldehyde inhibitors that have a P4 *N*-terminal carbamate, typically only one hydrogen bond is observed between the P4 *N*-terminal cap and M^Pro^ E166. One additional hydrogen bond that is formed between the P4 *N*-terminal cap and M^Pro^ E166 provides an explanation of very high *in vitro* potency of MPI29. In the M^Pro^-MPI29 complex, the hemithioacetal hydroxide is predominantly in an *S* configuration and poises to interact with three M^Pro^ backbone amide NH groups from G143, S144 and C145 that purportedly generates an oxyanion hole to stabilize the transition state of M^Pro^-catalyzed hydrolysis of a substrate. This favorable binding of the hemithioacetal hydroxide at the oxyanion hole likely contributes to strong *in vitro* potency for peptidyl aldehyde inhibitors. In M^Pro^-MPI29, the P2 dimethylcyclopropylproline is in van der Waals distances to most surrounding M^Pro^ residues in the P2 binding pocket, which indicates favorable hydrophobic interactions. The P3 *tert*-butylglycine doesn’t involve obvious interactions with M^Pro^ residues. However, the *tert*-butyl group in the P4 *N*-terminal cap fits nicely to the M^Pro^ binding pocket indicating favorable van der Waals interactions with surrounding M^Pro^ residues. All these favorable interactions likely contribute to the exceedingly high *in vitro* potency of MPI29.

MPI30 is different from MPI29 only at its warhead. It has an α-ketoamide. In its crystal structure as shown in Figure 2C, MPI30 forms interactions with M^Pro^ similar to MPI29 at the P1, P2 and P3 residues and the P4 *N*-terminal group. We observed one additional hydrogen bond that is formed between the P4 *N*-terminal carbamide oxygen and the side chain of M^Pro^ Q189. The keto group of the α-ketoamide of MPI30 covalently interacts with M^Pro^ C145 to generate a hemithioacetal. Unlike MPI29, the hemithioacetal hydroxide adopts a conformation pointing away from the oxyanion hole. It is in an unambiguously *S* configuration and interacts with H41 through two wa-ter-bridged hydrogen bonds. The amide part of the warhead points toward the oxyanion hole. The amide oxygen is assigned at the oxyanion hole due to its ability to form hydrogen bonds with three backbone NH groups from M^Pro^. The amide nitrogen interacts indirectly with the backbone oxygen of M^Pro^ T26 through two water-bridged hydrogen bonds. All these interactions likely contribute to the high *in vitro* potency of MPI30.

Since MPI32-36 have an MPI30 base structure and different *N*-appendages on the α-ketoamide, they are presented together in Figure 3C. Except at the appendage position, all other parts of MPI32-36 interact with M^Pro^ similar to MPI30. We designed MPI32-36 with a hope that their α-ketoamide *N*-appendages may reach the M^Pro^ P3’ binding pocket for favorable interactions. However, all *N*-appendages have a *Z* configuration that makes them point away from the M^Pro^ P3’ binding pocket. The appendage of MPI35 is long enough to fold back to the M^Pro^ P3’ pocket. Since the *N*-benzyl group of MPI36 has weak electron density around the phenyl ring, it is difficult to assess its interactions with M^Pro^. Its more favorable *in vitro* potency than MPI34 and MPI35 is likely due to higher structural rigidity of the *N*-benzyl group than two flexible *N*-alkyl groups in MPI34 and MPI35. The indirect interaction between the α-ketoamide nitrogen and M^Pro^ T26 that was observed in the M^Pro^-MPI30 structure is not observed in M^Pro^ complexes with MPI32-36. All *N*-appendages occupy the original water position and consequently remove this interaction. This might contribute to low *in vitro* potency of MPI32-36.

As shown in Figures 3E and 3F, MPI37 and MPI38 interact with M^Pro^ similar to MPI29. The P2 residue in MPI37 occupy slightly less space than that in MPI29. This may contribute to less *in vitro* potency of MPI37 than MPI29. In M^Pro^-MPI38, there exists a water molecule that bridges an indirect interaction between the P2 backbone nitrogen and the side chain of M^Pro^ Q189. MPI38 displays an M^Pro^ binding mode very similar to MPI18 that was previously developed and structurally resolved in its complex with M^Pro^ (PDB entry: 7RVR).^31^ The only difference is at the P4 *N*-terminal cap. MPI18 has a P4 *N*-terminal CBZ group that displays loose binding to M^Pro^ and cannot be unambiguously defined in its M^Pro^ complex structure. However, MPI38 has an *N-tert*-butylcarbamide cap whose structure can be clearly defined in the M^Pro^-MPI38 complex. MPI38 is slightly more potent than MPI18. This slightly higher potency might be attributed to more favorable interactions between the P4 *N*-terminal cap and the M^Pro^ P4 binding pocket.

MPI42 is structurally similar to MPI29 except that it has a P4 *N*-terminal isovaleramide. As shown in Figure 3G, MPI42 interacts with M^Pro^ similar to MPI29 except at the P4 *N*-terminal cap. The isovaleramide cap adopts a unique orientation that points deep into the M^Pro^ P4 binding pocket. This is different from most currently known peptidyl aldehyde inhibitors. M^Pro^ naturally prefers a P4 valine in its substrates. Isovaleramide is almost structurally identical to valine except that it doesn’t have an α-amide. It is apparent that isovaleramide engages favorable interactions with M^Pro^ by binding deep to the M^Pro^ P4 binding pocket. This explains its much higher *in vitro* potency than MPI40 that has an additional methyl group at the P4 *N*-terminal cap. The 3,3-dimethylbutamide cap in MPI40 is too bulky to fit into the relatively small M^Pro^ P4 binding pocket.

### Characterizations of *in cellulo* M^Pro^ inhibition potency of MPI29-47 and PF-07321332

When expressed in a human cell host, M^Pro^ leads to acute cytotoxicity and drives the host cell to undergo apoptosis. Using this unique feature, we previously developed a cell-based assay to characterize *in cellulo* potency of M^Pro^ inhibitors.^21^ In this assay, an inhibitor with *in cellulo* potency suppresses cytotoxicity from an M^Pro^-eGFP fusion protein that is ectopically expressed in 293T cells and consequently leads to host cell survival and enhanced overall expression of M^Pro^-eGFP that can be characterized by flow cytometry. We consider that this assay is more advantageous over a direct antiviral assay in the characterization of M^Pro^ inhibitors since an inhibitor may block functions of host proteases such as TMPRSS2, furin, and cathepsin L that are critical for SARS-CoV-2 infection and therefore provides false positive *in cellulo* potency results for M^Pro^ inhibitors.^35–37^ Using this assay, we characterized all synthesized inhibitors. Inhibitor-driven M^Pro^-eGFP expression data are presented in Figure 4 and the determined *in cellulo* EC_50_ values are summarized in Table 1. MPI29-39 that have a P4 *N*-terminal carbamide cap all showed low *in cellulo* potency. Since data for all these inhibitors do not reach a plateau at their highest tested concentrations, their EC_50_ values can only be estimated as higher than 5 or 10 μM. Although MPI29 has the most *in vitro* potency among all inhibitors that we synthesized, it has very weak potency in cells. MPI40-42 have a P4 *N*-terminal amide cap. In comparison to MPI29-39, they showed higher *in cellulo* potency although their determined EC_50_ values are in a single digit μM range and therefore are relatively high. MPI43 has a P4 *N*-terminal carbamate and is different from MPI29 and MPI40 at only one atom position. It has a determined *in cellulo* EC_50_ value as 0.37 μM. This *in cellulo* potency is 20-fold higher than MPI40 and 160-fold higher than MPI29. Considering that MPI29, MPI40 and MPI43 are different only at one atom position and MPI29 has much higher *in vitro* potency than MPI43, their drastically reversed potency in cells is intriguing. These three inhibitors must have very different plasma/cellular stability, cellular permeability, or both. MPI44 has a P4 *N*-terminal CBZ cap. It has a determined *in cellulo* EC_50_ value similar to that of MPI43. Since MPI43 and MPI44 have similar *in vitro* potency as well, the identity of a P4 *N*-terminal carbamate seems to have little effect on an inhibitor’s *in vitro* and *in cellulo* potency. MPI45 and MPI46 each have chemical moieties at two positions that are switched from that in MPI29 to that in MPI8. Both have high *in cellulo* potency. Their determined EC_50_ values are 0.74 and 0.14 μM respectively. It is evident that chemical moieties in MPI8 are optimal for high *in cellulo* potency. The *in cellulo* potency of MPI45 is much lower than that of MPI46 likely due to its P4 *N*-terminal carbamide cap. Both MPI47 and PF-07321332 have a P4 *N*-terminal amide, their determined *in cellulo* EC_50_ values are similar to that for MPI40-42 that also contain a P4 *N*-terminal amide. Much lower *in cellulo* potency of MPI47 than other four inhibitors is likely attributed to its low *in vitro* potency. Overall, inhibitors with a P4 *N*-terminal amide cap tend to have much lower *in cellulo* potency than that with a P4 *N*-terminal carbamate and for the two groups of inhibitors, their *in vitro* potency don’t correlate with their *in cellulo* potency. However, the three warheads aldehyde, α-ketoamide and nitrile don’t generate large discrepancy between *in vitro* and *in cellulo* potency of inhibitors.

**Figure 4.**
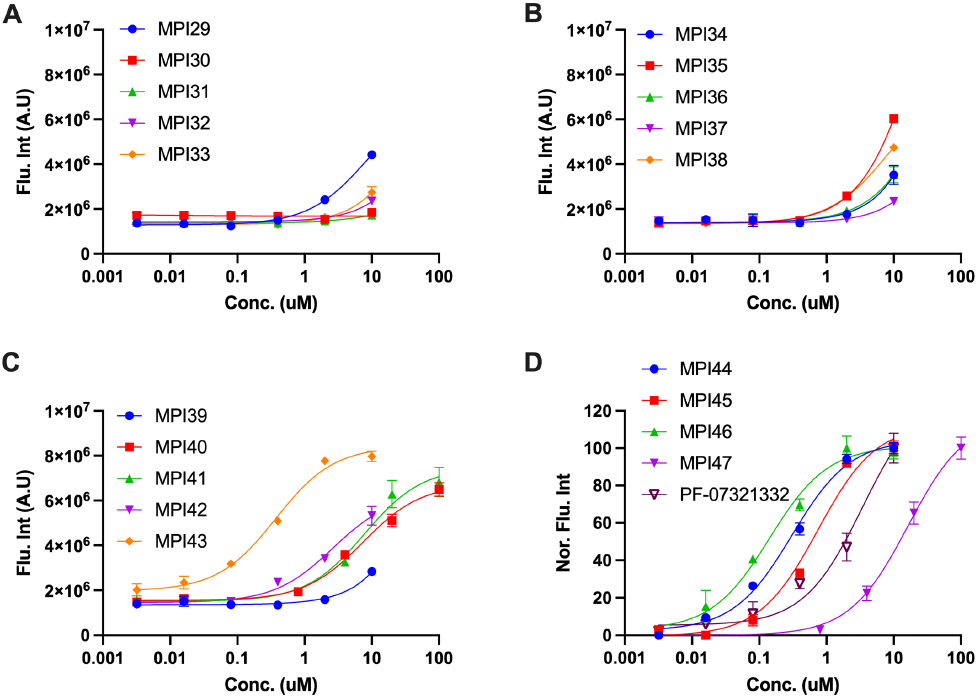
Cellular potency of MPI29-47 in their inhibition of M^Pro^ to drive host 293T cell survival and overall M^Pro^-eGFP expression. In D, fluorescent intensity is normalized due to that data for PF-07321332 was collected at a different time using different setups.

### Characterizations of antiviral potency of four selected inhibitors on three SARS-CoV-2 variants

Three most cellularly potent inhibitors MPI43, MPI44 and MPI45 were advanced to antiviral assays. One inhibitor MPI40 that has low cellular potency was characterized as well as a negative control. To quantify their antiviral EC_50_ values, we conducted plaque reduction neutralization tests of three SARS-CoV-2 variants including USA-WA1/2020, Beta and Delta in Vero E6 cells for all four inhibitors. We infected Vero E6 cells by virus in the presence of an inhibitor at various concentrations for three days and then quantified viral plaque reduction. Based on viral plaque reduction data, we determined antiviral EC_50_ values for all inhibitors. MPI40 showed no inhibition at all tested concentrations, which matches its detected low *in cellulo* potency. All other three inhibitors have high antiviral potency. Their determined antiviral EC_50_ values are similar and around 1 μM (Figure 5 and Table 1). MPI46 show the most well shaped antiviral data. It inhibits the Delta variants slightly better than the other two SARS-CoV-2 variants.

## ■ DISCUSSION

As a potential repurposed drug for COVID-19, boceprevir provided a high hope during the early phase of the COVID-19 pandemic. However, it is only moderate against SARS-CoV-2. Although it shows inhibition of M^Pro^, it has very weak potency to inhibit M^Pro^ in a human cell host.^21^ Lin *et al*. made ML1000 (MPI30 in our series) based on boceprevir. Although it showed high *in vitro* potency, its antiviral potency was minimal.^34^ In our study, both MPI29 and MPI30 are highly potent M^Pro^ inhibitors *in vitro*. The *in vitro* potency of MPI29 has even reached the detection limit of our kinetic assay. However, both inhibitors show very low *in cellulo* potency. Except MPI45 that has some MPI8 moieties, all other inhibitors that have a P4 *N*-terminal carbamide cap all display low *in cellulo* potency. Based on these data, we can conclude that M^Pro^ inhibitors that have a P4 *N*-terminal carbamide will have low cellular and antiviral potency. Although reasons for this low cellular and antiviral potency need to be further investigated, the emergency situation of COVID-19 demands to focus on other routes to develop M^Pro^ inhibitors as potential SARS-CoV-2 therapeutics. An alternative to the P4 *N*-terminal carbamide is amide. We synthesized several inhibitors that contain a P4 *N*-terminal amide. In comparison to inhibitors that have a P4 *N*-terminal carbamide, P4 amide-containing inhibitors are relatively more potent in cells but their cellular potency is moderate. However, all three inhibitors with a P4 *N*-terminal carbamate show high cellular potency and also high antiviral potency. Our collected data support a conclusion that a P4 *N*-terminal carbamate in a boceprevir-based M^Pro^ inhibitor is required for high cellular and antiviral potency. In other positions, a P1 opal residue, a P2 dimethylcyclopropylproline, and a P3 *tert*-butylglyine or *O-tert*-butylthreonine are necessary for high cellular and antiviral potency. In previous work, we discovered that a P3 *O-tert*-butylthreonine is critical for an M^Pro^ inhibitor to have high cellular and antiviral potency.^31^ This observation is supported by the high cellular and antiviral potency of MPI46. MPI8 was previously discovered as a highly potent SARS-CoV-2 inhibitor with an EC_50_ value as 30 nM.^21, 32, 38^ All boceprevir-based inhibitors that we developed in this work do not reach the potency of MPI8. To improve cellular and antiviral potency of inhibitors such as MPI43, MPI44 and MPI46, a possible route is to search for alternative *O*-alkyl groups in the P4 *N*-terminal carbamate cap for improved binding to the M^Pro^ P4 binding pocket. For the warhead, all current data indicate that aldehyde has the best activity. Changing it to either α-ketoamide or nitrile significantly decreases both *in vitro* and *in cellulo* potency of an inhibitor. Compounds with an aldehyde will lead to a concern of cytotoxicity. Cytotoxicity and potency of an aldehyde inhibitor for M^Pro^ needs to be carefully balanced when it is finally applied in animals or human patients.

**Figure 5.**
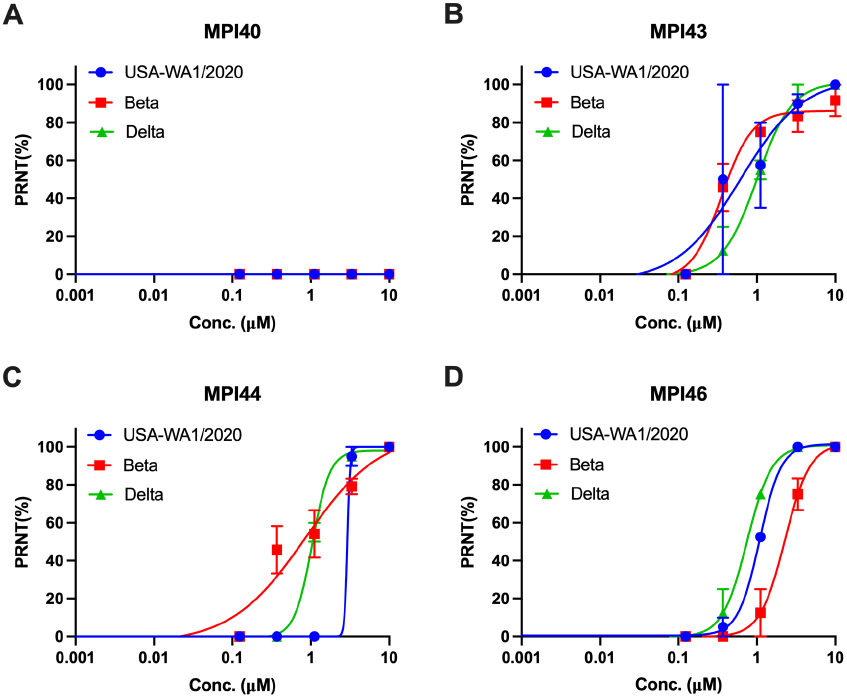
Plaque reduction neutralization tests (PRNTs) of MPI40, MPI43, MPI44, and MPI46 on their inhibition of three SARS-CoV-2 strains USA-WA1/2020, Beta and Delta in Vero E6 cells. Two repeats were conducted for each concentration.

We synthesized a number of α-ketoamide inhibitors with different alkyl substituents at the α-ketoamide nitrogen. The purpose was to make these *N*-appendages reach the M^Pro^ P3’ binding pocket. Unfortunately, all these inhibitors have low *in vitro* and *in cellulo* potency. Structures of their complexes with M^Pro^ all show a *Z* configuration at the α-ketoamide. It is evident that steric hinderance between the hemithioacetal hydroxide and an *N*-appendage prevents the *N*-appendage from adopting a favorable position to interact with the M^Pro^ P3’ binding pocket. Although further optimization in related M^Pro^ inhibitors may likely generate interactions with the M^Pro^ P3’ binding pocket, we suggest other routes to take advantage of the M^Pro^ P3’ binding pocket. In our characterized structures, the P4 *N*-terminal isovaleramide of MPI42 shows a unique conformation that binds deep into the M^Pro^ P4 binding pocket. This strong binding explains MPI42’s much higher *in vitro* potency than its close relative MPI40. We think MPI42 points to a unique way to develop potent M^Pro^ inhibitors by exploring P4 *N*-terminal chemical moieties with a similar size as isovaleramide.

## ■ CONCLUSIONS

Based on our systematic analysis of boceprevir-based M^Pro^ inhibitors, we conclude that a P4 *N*-terminal carbamate in an inhibitor is critical to yield high cellular and antiviral potency. However, chemical moieties at all sites including the P4 *N*-terminal carbamate and the warhead might be continuously optimized.

## ■ EXPERIMENTAL SECTION

## ■ ASSOCIATED CONTENT

### Supporting Information

The Supporting Information is available free of charge on the ACS Publications website.

Supplementary information for the synthesis of MPI29-47 and PF-07321332, NMR spectroscopy and HPLC chromatography figures, X-ray data collection and processing parameters, and all original flow cytometry graphs. All compounds are >95% pure by HPLC analysis. MPI40 and MPI46 contain two diastereomers.

## ACKNOWLEDGMENT

This work was supported by Welch Foundation (grant A-1715), DHHS-NIH-National Institute of Allergy and Infectious Diseases (R21AI164088), TAMU COS Strategic Transformative Research Program, and Texas A&M X Grants. The ALS-ENABLE beamlines are supported in part by the National Institutes of Health, National Institute of General Medical Sciences, grant P30 GM124169-01 and the Howard Hughes Medical Institute. The Advanced Light Source is a Department of Energy Office of Science User Facility under Contract No. DE-AC02-05CH11231.

---

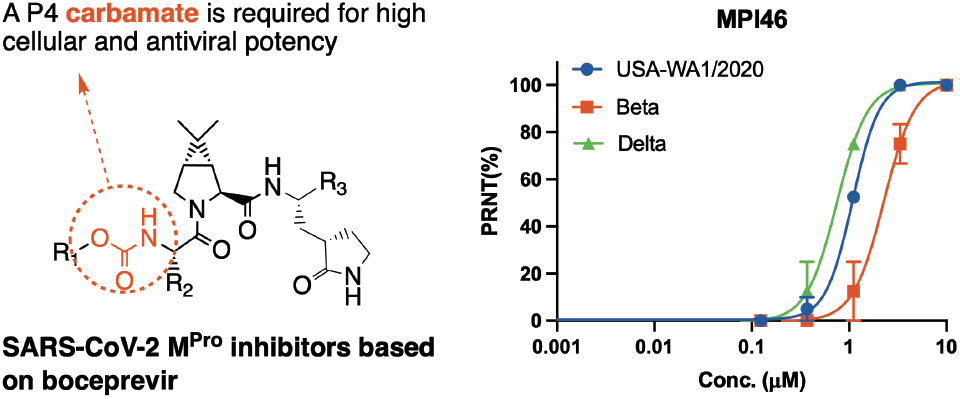

