## Supplementary Material for "The *N*-Terminal Carbamate is Key to High Cellular and Antiviral Potency for Boceprevir-Based SARS-CoV-2 Main Protease Inhibitors"

for

**Materials.** We purchased yeast extract from Thermo Fisher Scientific, tryptone from Gibco, Sub3 from Bachem, HEK 293T/17 cells from ATCC, DMEM with GlutaMax from Gibco, FBS from Gibco, polyethyleneimine from Polysciences, the trypsin-EDTA solution from Gibco. Chemicals used in this work were acquired from Sigma Aldrich, Chem Impex, Ambeed, A2B, etc.

**$M^{Pro}$ Expression and Purification.** The expression plasmid pET28a-His-SUMO-$M^{Pro}$ was constructed in a previous study. We used this construct to transform *E. coli* BL21(DE3) cells. A single colony grown on a LB plate containing 50 μg/mL kanamycin was picked and grown in 5 mL LB media supplemented with 50 μg/mL kanamycin overnight. We inoculated this overnight culture to 6 L 2YT media with 50 μg/mL kanamycin. Cells were grown to $OD_{600}$ as 0.8. At this point, we added 1 mM IPTG to induce the expression of His-SUMO-$M^{Pro}$. Induced cells were let grown for 3 h and then harvested by centrifugation at 12,000 rpm, 4 °C for 30 min. We resuspended cell pellets in 150 mL lysis buffer (20 mM Tris-HCl, 100 mM NaCl, 10 mM imidazole, pH 8.0) and lysed the cells by sonication on ice. We clarified the lysate by centrifugation at 16,000 rpm, 4 °C for 30 min. We decanted the supernatant and mixed with Ni-NTA resins (GenScript). We loaded the resins to a column, washed the resins with 10 volumes of lysis buffer, and eluted the bound protein using elution buffer (20 mM Tris-HCl, 100 mM NaCl, 250 mM imidazole, pH 8.0). We exchanged buffer of the elute to another buffer (20 mM Tris-HCl, 100 mM NaCl, 10 mM imidazole, 1 mM DTT, pH 8.0) using a HiPrep 26/10 desalting column (Cytiva) and digested the elute using 10 units SUMO protease overnight at 4 °C. The digested elute was subjected to Ni-NTA resins in a column to remove His-tagged SUMO protease, His-tagged SUMO tag, and undigested His-SUMO-$M^{Pro}$. We loaded the flow-through onto a Q-Sepharose column and purified $M^{Pro}$ using FPLC by running a linear gradient from 0 to 500 mM NaCl in a buffer (20 mM Tris-HCl, 1 mM DTT, pH 8.0). Fractions eluted from the Q-Sepharose column was concentrated and loaded onto a HiPrep 16/60 Sephacryl S-100 HR column and purified using a buffer containing 20 mM Tris-HCl, 100 mM NaCl, 1 mM DTT, and 1 mM EDTA at pH 7.8. The final purified was concentrated and stored in a -80 °C freezer.

***In Vitro* $M^{Pro}$ Inhibition Potency Characterizations of MPIs.** For most MPIs, we conducted the assay using 20 nM $M^{Pro}$ and 10 μM Sub3. For MPI13-14, 10 nM $M^{Pro}$ was used. We dissolved all inhibitors in DMSO as 10 mM stock solutions. Sub3 was dissolved in DMSO as a 1 mM stock solution and diluted 100 times in the final assay buffer containing 10 mM $Na_xH_yPO_4$, 10 mM NaCl, 0.5 mM EDTA, and 1.25% DMSO at pH 7.6. We incubated $M^{Pro}$ and an inhibitor in the final assay buffer for 30 min before adding the substrate to initiate the reaction catalyzed by $M^{Pro}$. The production format was monitored in a fluorescence plate reader with excitation at 336 nm and emission at 455 nm. More assay details can be found in a previous study.[1]

**X-Ray Crystallography Analysis of $M^{Pro}$-Inhibitor Complexes.** The production of crystals of $M^{pro}$-inhibitor complexes was following the previous protocols.[1] The data of $M^{pro}$ with MPI29, MPI32, MPI33, MPI35, MPI37 and MPI42 were collected on a Bruker Photon II detector. The data of $M^{pro}$ with MPI30, MPI34, MPI36 and MPI38 were collected at the Advanced Light Source (ALS) beamline 5.0.2 using a Pilatus3 6M detector. The diffraction data were indexed, integrated and scaled with iMosflm or PROTEUM3.[2] All the structures were determined by molecular replacement using the structure model of the free enzyme of the SARS-CoV-2 $M^{pro}$ [Protein Data Bank (PDB) ID code 7JPY] as the search model using Phaser in the Phenix package.[1,3] *JLigand* and *Sketcher* from the CCP4 suite were employed for the generation of PDB and geometric restraints for the inhibitors. The inhibitors were built into the Fo-Fc density by using *Coot*.[4] Refinement of all the structures was performed with Real-space Refinement in Phenix.[3] Details of data quality and structure refinement are summarized in Table S1. All structural figures were generated with PyMOL (https://www.pymol.org).

***In cellulo* $M^{Pro}$ Inhibition Potency Characterizations of MPIs.** We grew HEK 293T/17 cells in high-glucose DMEM with GlutaMAX supplement and 10% FBS in 10 cm culture plates under 37 °C and 5% $CO_2$ to 80-90% confluency and then transfected cells with the pLVX-$M^{Pro}$-eGFP-2 plasmid. 30 mg/mL polyethyleneimine and the total of 8 μg of the plasmid in 500 μL opti-MEM media were used for transfection. We incubated transfected cells overnight. On the second day, we collected cells using 0.05% trypsin-EDTA to detach them from plates, resuspended collected cells in the original growth media, adjusted the cell density to $5 \cdot 10^5$ cells/mL, added 500 μL adjusted cells to each well of a 48-well plate, and then added 100 μL of a drug solution in DMEM. We incubated treated cells under 37 °C and 5% $CO_2$ for 72 h. After 72 h incubation, cells were collected using trypsinization and centrifugation. We resuspended collected cells in 200 μL PBS and analyzed cells with fluorescence using a Cytoflex Research Flow Cytometer based on the size scatters (SSC-A and SSC-H) and forward scatter (FSC-A). We gated cells based on SSC-A and FSC-A then with SSC-A and SSC-H. Fluorescence was detected with excitation at 488 nm and emission at 525 nm. All collected data were converted to csv files and analyzed using a self-prepared MATLAB script for massive data processing. We sorted the FITC-A column from lowest to highest. A $10^6$ cutoff was set to separate the column to two groups with higher than $10^6$ as positive and lower than $10^6$ as negative. We integrated the positive group and divided the total integrated fluorescence intensity by the total cell positive cell counts as Flu. Int. shown in all graphs. The standard deviation of positive fluorescence was calculated as well. All processed data were plotted and fitted to a four-parameter Hill equation in GraphPad 9.0 to obtain determined $EC_{50}$ values.

**The Synthesis of MPIs.** All reagents and solvents for the synthesis were purchased from commercial sources and used without purification. All glassware was flame-dried prior to use. Thin-layer chromatography (TLC) was carried out on aluminum plates coated with 60 F254 silica gel. TLC plates were visualized under UV light (254 or 365 nm) or stained with 5% phosphomolybdic acid. Normal phase column chromatography was carried out using a Yamazen Small Flash AKROS system. Analytical reverse phase HPLC was carried out on a Shimazu LC20 HPLC system with an analytical C18 column. Semipreparative HPLC was carried out the same system with a semipreparative C18 column. The mobile phases were $H_2O$ with 0.1% formic acid (A) and acetone with 0.1% formic acid (B). NMR spectra were recorded on a Bruker AVANCE Neo 400 MHz or Varian INOVA 300 MHz spectrometer in specified deuterated solvents. High-resolution electrospray mass spectrometry was carried out on a Thermo Scientific Q Exactive Focus system. The purity of all compounds were confirmed by NMR and analytic HPLC-UV as ≥ 95%.

**HPLC analysis of MPIs.** All compounds were determined by using Thermo Scientific ultimate 3000 HPLC with binary pumps, using Acclaim 120 C18 column (2.1x150 mm, 5 μL). All compounds were analyzed using (MeOH/$H_2O$ 0.1% Formic acid)(v/v)(0.3mL/min) and calculated the peak areas at 254 or 214 nm.

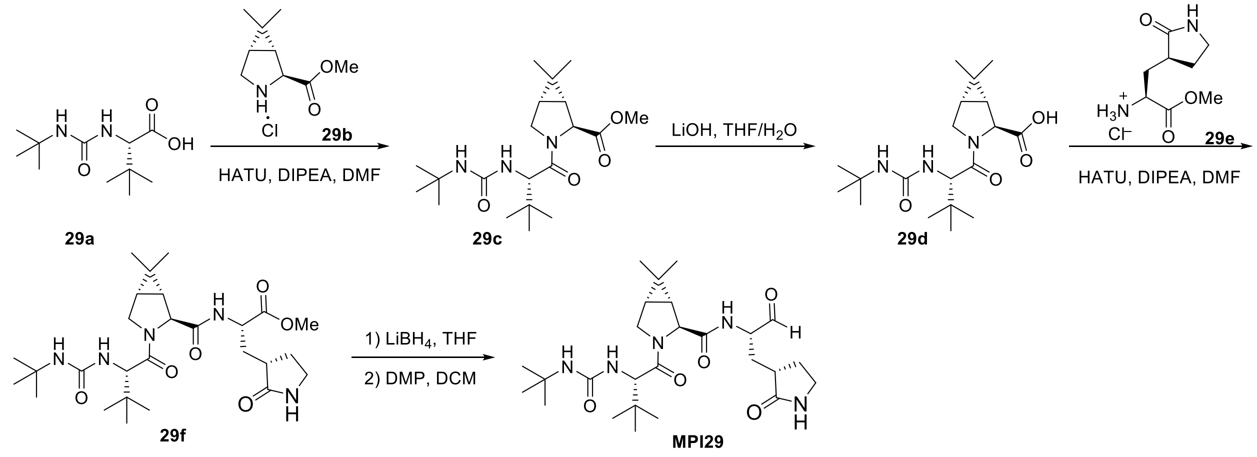


**Scheme 1**. The synthesis of compound **MPI29**

**Methyl (1R,2S,5S)-3-((S)-2-(3-(tert-butyl)ureido)-3,3-dimethylbutanoyl)-6,6-dimethyl-3-azabicyclo[3.1.0]hexane-2-carboxylate (29c).** To a solution of **29a** (4 mmol, 0.92 g) and **29b** (4 mmol, 0.82 mg) in anhydrous DMF (20 mL) was added DIPEA (10 mmol, 1.29 g) and was cooled to 0 °C. HATU (4.4 mmol, 1.67 g) was added to the solution under 0 °C and then stirred at room temperature overnight. The reaction mixture was then diluted with ethyl acetate (100 mL) and washed with saturated $NaHCO_3$ solution (2×50 mL), 1 M HCl solution (2×50 mL), and saturated brine solution (2×50 mL) sequentially. The organic layer was dried over anhydrous $Na_2SO_4$ and then concentrated *in vacuo*. The residue was then purified with flash chromatography (15-50% EtOAc in hexanes as the eluent) to afford **29c** as colorless oil (1.14 g, 75%).

**(1R,2S,5S)-3-((S)-2-(3-(tert-butyl)ureido)-3,3-dimethylbutanoyl)-6,6-dimethyl-3-azabicyclo[3.1.0]hexane-2-carboxylic acid (29d).** **29c** (3 mmol, 1.14 g) was dissolved in 10 mL of THF. A solution of LiOH·$H_2O$ (6 mmol, 250 mg) in 5 mL $H_2O$ was added to the solution. The mixture was stirred at room temperature for 3 h. Then THF was removed *in vacuo* and the aqueous layer was acidified with 1 M HCl and extracted with dichloromethane (3×20 mL). The organic layer was dried over anhydrous $Na_2SO_4$ and concentrated to yield **29d** as white solid (1.01 g, 92 %). $^1$H NMR (400 MHz, DMSO-$d_6$) δ 12.57 (s, 1H), 5.95 (s, 1H), 5.88 (d, $J$ = 10.0 Hz, 1H), 4.15 (d, $J$ = 10.0 Hz, 1H), 4.11 (s, 1H), 3.99 (d, $J$ = 10.4 Hz, 1H), 3.74 (dd, $J$ = 10.3, 5.3 Hz, 1H), 1.48 (dd,

*J* = 7.6, 5.1 Hz, 1H), 1.38 (d, *J* = 7.5 Hz, 1H), 1.17 (s, 9H), 1.00 (s, 3H), 0.91 (s, 9H), 0.82 (s, 3H).

**Methyl (S)-2-((1R,2S,5S)-3-((S)-2-(3-(tert-butyl)ureido)-3,3-dimethylbutanoyl)-6,6-dimethyl-3-azabicyclo[3.1.0]hexane-2-carboxamido)-3-((S)-2-oxopyrrolidin-3-yl)propanoate (29f).** To a solution of **29d** (1 mmol, 367 mg) and **29e** (1 mmol, 222 mg) in anhydrous DMF (5 mL) was added DIPEA (2 mmol, 258 mg) and was cooled to 0 °C. HATU (1.2 mmol, 456 mg) was added to the solution under 0 °C and then stirred at room temperature overnight. The reaction mixture was then diluted with ethyl acetate (50 mL) and washed with saturated $NaHCO_3$ solution (2×20 mL), 1 M HCl solution (2×20 mL), and saturated brine solution (2×20 mL) sequentially. The organic layer was dried over anhydrous $Na_2SO_4$ and then concentrated *in vacuo*. The residue was then purified with flash chromatography (1-10% methanol in dichloromethane as the eluent) to afford **29f** as white solid (321 mg, 60%). $^1$H NMR (400 MHz, Chloroform-*d*) δ 7.42 (d, *J* = 8.0 Hz, 1H), 6.01 (s, 1H), 5.17 – 4.89 (m, 1H), 4.64 (ddd, *J* = 11.6, 7.9, 4.0 Hz, 1H), 4.37 (s, 1H), 4.34 (s, 1H), 4.08 (d, *J* = 10.3 Hz, 1H), 3.88 (dt, *J* = 10.4, 2.7 Hz, 1H), 3.73 (s, 3H), 3.37 – 3.22 (m, 2H), 2.56 – 2.35 (m, 2H), 2.25 – 2.13 (m, 1H), 1.93 – 1.76 (m, 3H), 1.50 (d, *J* = 2.3 Hz, 2H), 1.26 (s, 9H), 1.02 (s, 3H), 0.97 (s, 9H), 0.89 (s, 3H).

**(1R,2S,5S)-3-((S)-2-(3-(tert-butyl)ureido)-3,3-dimethylbutanoyl)-6,6-dimethyl-N-((S)-1-oxo-3-((S)-2-oxopyrrolidin-3-yl)propan-2-yl)-3-azabicyclo[3.1.0]hexane-2-carboxamide (MPI29).** To a solution of **29f** (0.25 mmol, 133 mg) in anhydrous dichloromethane (5 mL) was added a solution of $LiBH_4$ in anhydrous THF (2 M, 0.25 mL, 0.5 mmol) at 0 °C. The resulting solution was stirred at the same temperature for 3 h. Then a saturated solution of $NH_4Cl$ (5 mL) was added dropwise to quench the reaction. The layers were separated, and the organic layer was washed with saturated brine solution (2×10 mL), dried over anhydrous $Na_2SO_4$. and evaporated to dryness. The residue was then dissolved in anhydrous dichloromethane (5 mL) and cooled to 0 °C. Dess-Martin periodinane (0.5 mmol, 212 mg) was added to the solution. The reaction mixture was then stirred at room temperature overnight. Then the reaction was quenched with a saturated $NaHCO_3$ solution containing 10 % $Na_2S_2O_3$. The layers were separated. The organic layer was then washed with saturated brine solution (2×10 mL), dried over anhydrous $Na_2SO_4$ and evaporated *in vacuo*. The residue was then purified with flash chromatography (1-10% methanol in dichloromethane as the eluent) to afford **MPI29** as white solid (64 mg, 51 %). $^1$H NMR (400

MHz, Chloroform-*d*) δ 9.54 (s, 1H), 6.34 (s, 1H), 5.20 (d, *J* = 10.0 Hz, 1H), 4.64 (s, 1H), 4.54 – 4.44 (m, 1H), 4.36 (t, *J* = 5.0 Hz, 2H), 4.10 (d, *J* = 10.4 Hz, 1H), 3.91 (dd, *J* = 10.4, 5.2 Hz, 1H), 3.31 (dq, *J* = 17.5, 9.3, 8.0 Hz, 2H), 2.53 (q, *J* = 8.1 Hz, 1H), 2.40 (ddd, *J* = 17.9, 9.3, 4.8 Hz, 1H), 1.98 (ddt, *J* = 21.6, 13.8, 5.3 Hz, 2H), 1.89 – 1.76 (m, 1H), 1.58 – 1.43 (m, 2H), 1.25 (s, 9H), 1.03 (s, 3H), 0.95 (s, 9H), 0.90 (s, 3H). $^{13}C$ NMR (101 MHz, $CDCl_3$) δ 199.8, 180.1, 172.8, 172.2, 157.3, 60.8, 57.8, 57.3, 50.2, 48.4, 40.5, 37.7, 34.7, 30.6, 30.2, 29.4, 28.5, 28.0, 26.6, 26.3, 19.3, 12.7.

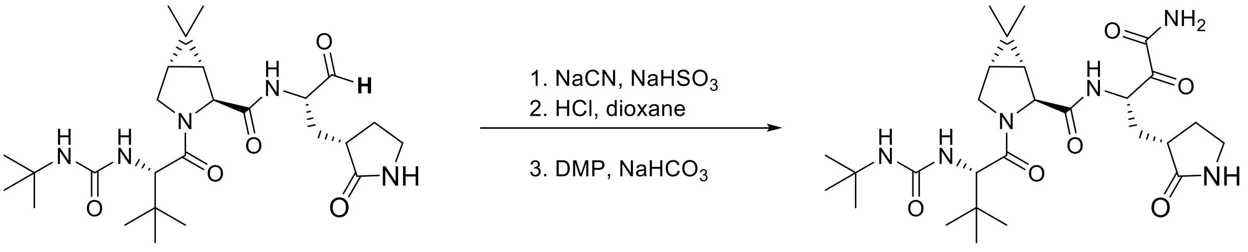


**Scheme 2**: The synthesis of compound **MPI30**

**Synthesis of (1R,2S,5S)-N-((S)-4-amino-3,4-dioxo-1-((S)-2-oxopyrrolidin-3-yl) butan-2-yl)-3-((S)-2-(3-(tert-butyl)ureido)-3,3-dimethylbutanoyl)-6,6-dimethyl-3-azabicyclo[3.1.0]hexane-2-carboxamide (MPI30).** To a solution of **MPI29** (500 mg, 0.9 mmol) in dichloromethane (25 mL) was added $NaHSO_3$ (480 mg, 4.6 mmol) slowly. The reaction was allowed to stir at RT for 30 min. Then NaCN (230 mg, 4.6 mmol), dissolved in 5 mL water was added to the reaction mixture slowly. The reaction mixture was stirred at RT for overnight. The mixture was washed with water, sat. NaCl, dried over $Na_2SO_4$ and concentrated. ESI-MS was used to confirm the formation of cyanohydrin intermediate, which was carried forward to the next step without further purification. To a solution of the cyanohydrin intermediate (350 mg, 0.65 mmol) in 1,4-dioxane (10 mL) was added dropwise a HCl solution in 1,4-dioxane (4 M, 10 mL). The resulting solution was stirred at room temperature for 3 h. Then residue was then concentrated *on vacuo* to afford the hydroxyamide intermediate. ESI-MS was used to confirm the formation of cyanohydrin intermediate, which was carried forward to the next step without further purification. In the final step, to a solution of the hydroxyamide intermediate (260 mg, 0.47 mmol, 1.0 equiv.,) in anhydrous DCM (10 mL) was added Dess-Martin reagent (616 g, 1.42 mmol, 5.0 equiv.,) slowly at 0 °C. Then the reaction mixture was stirred at RT for 2 h. The formation of the desired product was confirmed by ESI-MS study. A solution of $NaHCO_3$ and $Na_2S_2O_3$ was added to quench the reaction. After 10 min, the mixture was washed with water, sat. NaCl, dried over $Na_2SO_4$ and concentrated. The residue was purified by column chromatography (MeOH: DCM = 1:10 v/v) to yield **MPI30** as a white solid (156 mg, yield 60 %). $^1$H NMR (400 MHz, DMSO-$d_6$) δ 7.49 – 7.31 (m, 2H), 5.90 – 5.72 (m, 2H), 4.19 – 3.96 (m, 3H), 3.92 – 3.79 (m, 1H), 3.76 – 3.62 (m, 1H), 3.14 – 2.82 (m, 3H), 2.29 (q, $J$ = 10.2 Hz, 1H), 2.12 (dd, $J$ = 12.4, 7.4 Hz, 1H), 1.88 – 1.73 (m, 1H), 1.63 – 1.46 (m, 1H), 1.44 – 1.31 (m, 2H), 1.31 – 1.24 (m, 1H), 1.22 – 1.15 (m, 1H), 1.14 – 1.03 (m, 9H), 0.96 – 0.89 (m, 3H), 0.86 – 0.73 (m, 11H). $^{13}$C NMR (100 MHz, DMSO-$d_6$) δ 197.87, 178.66, 172.32, 171.66, 171.31, 171.23, 163.53, 157.88, 60.01, 57.28, 49.41, 34.57, 34.52, 29.58, 27.77, 26.88, 19.11, 13.05.

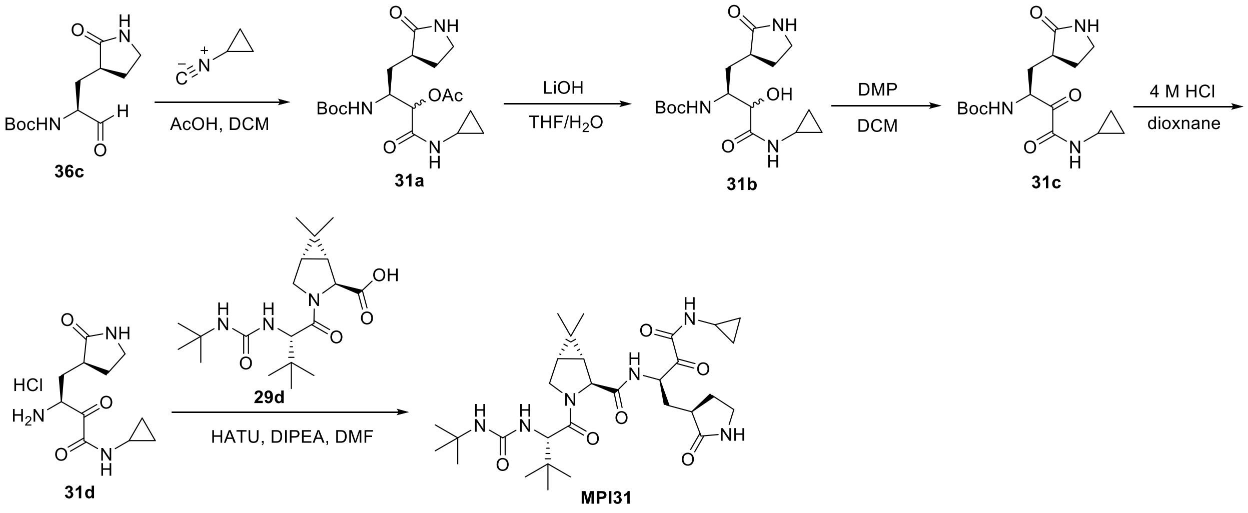


**Scheme 3**. The synthesis of compound **MPI31**

**(3S)-3-((tert-butoxycarbonyl)amino)-1-(cyclopropylamino)-1-oxo-4-((S)-2-oxopyrrolidin-3-yl)butan-2-yl acetate (31a).** To a solution of **36c** (100 mg, 0.39 mmol) in DCM was added acetic acid (47 mg, 0.78 mmol) and cyclopropyl isocyanide (52 mg, 0.78 mmol). The resulting solution was stirred at room temperature overnight and concentrated *in vacuo*. The crude product was then used without purification for the next step.

**tert-butyl ((2S)-4-(cyclopropylamino)-3-hydroxy-4-oxo-1-((S)-2-oxopyrrolidin-3-yl)butan-2-yl)carbamate (31b).** To a solution of crude **31a** from last step in THF (5 mL) was added an aqueous solution of LiOH (65 mg, 1.56 mmol). The reaction mixture was stirred at room temperature for 3 h and then diluted with water (10 mL), extracted with dichloromethane (2×10 mL). The organic layers were combined and dried with anhydrous $Na_2SO_4$ and concentrated *in vacuo*. The crude product was used without further purification for the next step.

**tert-butyl((S)-4-(cyclopropylamino)-3,4-dioxo-1-((S)-2-oxopyrrolidin-3-yl)butan-2-yl)carbamate (31c).** To a solution of crude **31b** from last step in dichloromethane (5 mL) was added Dess-Martin periodinane (330 mg, 0.78 mmol) at 0 °C. The reaction mixture was added stirred at room temperature for 3 h. Then the reaction was quenched with a saturated $NaHCO_3$ solution containing 10 % $Na_2S_2O_3$. The layers were separated. The organic layer was then washed with saturated brine solution (2×10 mL), dried over anhydrous $Na_2SO_4$, and evaporated *in vacuo*. The residue was purified by flash chromatography (0-10% methanol in dichloromethane as eluent) to yield **31c** as white solid (51 mg, 38 %). $^1$H NMR (400 MHz, Chloroform-*d*) δ 7.12 – 6.95 (m, 1H), 6.34 (d, *J* = 21.4 Hz, 1H), 5.77 (d, *J* = 7.8 Hz, 1H), 5.08 (ddd, *J* = 11.0, 7.7, 3.3 Hz, 1H), 3.44 – 3.23 (m, 2H), 2.75 (tp, *J* = 7.6, 3.8 Hz, 1H), 2.60 – 2.41 (m, 2H), 2.08 – 1.80 (m, 3H), 1.40 (d, *J* = 4.8 Hz, 9H), 0.82 (ddd, *J* = 7.3, 3.1, 1.4 Hz, 2H), 0.59 (heptd, *J* = 4.8, 3.8, 1.6 Hz, 2H).

**(1R,2S,5S)-3-((S)-2-(3-(tert-butyl)ureido)-3,3-dimethylbutanoyl)-N-((R)-4-(cyclopropylamino)-3,4-dioxo-1-((R)-2-oxopyrrolidin-3-yl)butan-2-yl)-6,6-dimethyl-3-azabicyclo[3.1.0]hexane-2-carboxamide (MPI31).** To a stirred solution of **31c** (34 mg, 0.1 mmol) in 1,4-dioxane (1 mL) was added a 4 M HCl solution in dioxane (4 mL). The reaction mixture was stirred at room temperature for 1 h and then concentrated *in vacuo*. The residue **31d** was then dissolved in DMF (2 mL). To the solution was then added **29d** (37 mg, 0.1 mmol) and DIPEA (39 mg, 0.3 mmol). HATU (46 mg, 0.12 mmol) was added to the solution at 0 °C. The reaction mixture was then stirred at room temperature overnight before being diluted with 20 mL EtOAc and washed with saturated $NaHCO_3$ solution (2×20 mL), 1 M HCl solution (2×20 mL), and saturated brine solution (2×20 mL) sequentially. The organic layer was then dried with anhydrous $Na_2SO_4$ and concentrated *in vacuo*. The residue was purified with flash chromatography to yield **MPI31** as white solid (8 mg, 14 %). $^1$H NMR (400 MHz, Chloroform-*d*) δ 7.78 (s, 1H), 6.97 (d, *J* = 3.8 Hz, 1H), 5.70 (s, 1H), 5.43 – 5.30 (m, 1H), 4.87 (s, 1H), 4.36 (s, 2H), 4.08 (d, *J* = 10.3 Hz, 1H), 3.86 (ddd, *J* = 17.3, 10.2, 5.0 Hz, 1H), 3.34 (td, *J* = 18.6, 16.9, 8.0 Hz, 2H), 2.76 (tq,

*J* = 7.6, 3.8 Hz, 1H), 2.65 – 2.53 (m, 1H), 2.55 – 2.43 (m, 1H), 2.16 – 1.88 (m, 3H), 1.49 (d, *J* = 4.5 Hz, 2H), 1.26 (s, 9H), 1.05 (d, *J* = 3.2 Hz, 1H), 1.03 (s, 3H), 0.98 (d, *J* = 4.8 Hz, 9H), 0.88 (s, 3H), 0.86 – 0.81 (m, 2H), 0.65 – 0.54 (m, 2H).

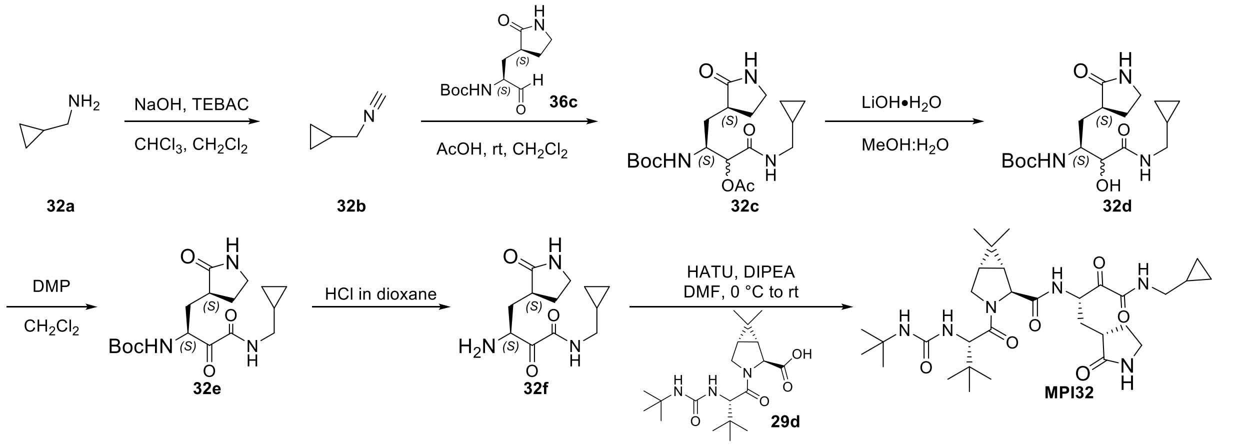


**Scheme 4**. The synthesis of compound **MPI32**

**tert-butyl ((S)-4-((cyclopropylmethyl)amino)-3,4-dioxo-1-((S)-2-oxopyrrolidin-3-yl)butan-2-yl)carbamate (32e).** Amine **32a** (9.9mmol, 1 equiv.,.) in dichloromethane (5.0 mL) was added to a solution of sodium hydroxide (2.36 g, 59.1 mmol, 6 equiv.,.) in water (2.5mL). Then N-benzyl-N,N,N-triethylammonium chloride (45 mg, 0.20 mmol, 0.02 equiv.,.) and chloroform (4.8 mL, 59.1 mmol, 6 equiv.) were added respectively. The mixture was stirred at room temperature (RT.) for 12 h. The organic layer was extracted with a syringe and directly purified by flash chromatography using dichloromethane to obtain the desired isocyanide. Eluent in a test tube (~10.0 mL) with most pungent odour was used directly in the next step. Aldehyde **36c** (0.46 mmol, 1 equiv.,.) was dissolved in the solution of isocyanide in dichloromethane, followed by addition of acetic acid (57 µL, 0.92 mmol, 2 equiv.). The mixture was stirred overnight at RT. All the volatiles were removed under vacuum and the residue was dissolved in a mixture of methanol and water (10.0 mL, v/v = 4/1). Lithium hydroxide monohydrate (42 mg, 0.37 mmol, 3 equiv.,.) was added in one portion and the resulting mixture was stirred at RT. for 1-4 h. Then it was neutralized by 0.1 M hydrochloric acid and concentrated under reduced pressure. The residue was extracted with acetic ester (20.0 mL × 3) and the combined organic layers were dried over anhydrous magnesium sulfate, filtered, and concentrated under vacuum. The residue was dissolved in anhydrous dichloromethane (10.0 mL) and added Dess-Martin reagent (275 mg, 0.63 mmol, 1.5 equiv.,.)

slowly at 0 °C. Then the reaction mixture was stirred at RT. for 1-2 h. A solution of sodium bicarbonate and sodium thiosulfate was added to quench the reaction. After 10 min, dichloromethane (10.0 mL × 2) was added to extract the mixture. The organic phase was washed with brine (10.0 mL × 2), dried over anhydrous sodium sulfate, filtered, and concentrated. The residue was purified by column chromatography (dichloromethane: methanol, 35:1 v/v) to afford the pure product as a white solid **32e** (70%).$^{1}$H NMR (400 MHz, Chloroform-*d*) δ 7.03 (s, 1H), 5.70 (d, *J* = 26.9 Hz, 1H), 5.17 (s, 1H), 3.38 (s, 2H), 3.19 (q, *J* = 6.8 Hz, 2H), 2.70 – 2.48 (m, 1H), 2.17 – 1.79 (m, 3H), 1.46 (s, 9H), 1.12 – 0.90 (m, 1H), 0.57 (t, *J* = 7.3 Hz, 2H), 0.26 (t, *J* = 5.2 Hz, 2H).

**(S)-3-amino-N-(cyclopropylmethyl)-2-oxo-4-((S)-2-oxopyrrolidin-3-yl)butanamide hydrochloride (32f):**

To a solution of **32e** (100 mg, 0.28 mmol)in 1,4-dioxane (10 mL) was added dropwise a HCl solution in 1,4-dioxane (4 M, 0.7 mL). The resulting solution was stirred at room temperature for 2 h. Then residue was then concentrated *on vacuo* to afford **32f** as light-yellow hygroscopic solid (65 mg, 80%). $^{1}$H NMR (400 MHz, Deuterium Oxide) δ 3.57 – 3.42 (m, 1H), 3.17 (dt, *J* = 9.7, 4.8 Hz, 2H), 3.01 – 2.77 (m, 2H), 2.59 – 2.44 (m, 1H), 2.27 – 2.10 (m, 1H), 1.77 – 1.44 (m, 3H), 0.92 – 0.72 (m, 1H), 0.39 – 0.21 (m, 2H), 0.09 – -0.13 (m, 2H).

**(1R,2S,5S)-3-((S)-2-(3-(tert-butyl)ureido)-3,3-dimethylbutanoyl)-N-((S)-4-((cyclopropylmethyl)amino)-3,4-dioxo-1-((S)-2-oxopyrrolidin-3-yl)butan-2-yl)-6,6-dimethyl-3-azabicyclo[3.1.0]hexane-2-carboxamide (MPI32).** To a solution of **29d** (0.19 mmol, 70 mg) and **32f** (0.19 mmol, 65 mg) in anhydrous DMF (5 mL) was added DIPEA (0.76 mmol, 0.52 g) and was cooled to 0 °C. HATU (0.24 mmol, 94 mg) was added at 0 °C and then stirred at room temperature overnight. The reaction mixture was then diluted with ethyl acetate (20 mL) and washed with saturated $NaHCO_3$ solution (2×10 mL), 1 M HCl solution (2×10 mL), and saturated brine solution (2×10 mL) sequentially. The organic layer was dried over anhydrous $Na_2SO_4$ and then concentrated *on vacuo*. The residue was then purified with flash chromatography (0-10% MeOH in Dichloromethane as the eluent) to afford **MPI32** as white solid (60 mg, 55%).$^{1}$H NMR (400 MHz, Chloroform-*d*) δ 7.60 (s, 1H), 6.88 (t, *J* = 5.8 Hz, 1H), 6.23 (s, 1H), 5.23 – 5.12 (m, 1H), 5.04 (d, *J* = 10.1 Hz, 1H), 4.59 – 4.46 (m, 1H), 4.19 – 4.03 (m, 1H), 3.95 – 3.77 (m, 1H), 3.20

– 2.97 (m, 2H), 2.97 – 2.89 (m, 2H), 2.45 – 2.01 (m, 3H), 1.82 – 1.71 (m, 2H), 1.33 – 1.20 (m, 3H), 1.03 (s, 9H), 0.80 (s, 3H), 0.74 (s, 9H), 0.66 (s, 3H), 0.40 – 0.22 (m, 2H), 0.12 – -0.08 (m, 2H). $^{13}$C NMR (101 MHz, $CDCl_3$) δ 195.55, 179.99, 172.74, 171.50, 159.14, 157.35, 77.37, 60.57, 57.79, 55.30, 53.30, 50.19, 50.15, 44.27, 40.49, 38.51, 34.72, 30.36, 29.35, 27.86, 27.67, 26.54, 26.36, 26.30, 19.20, 12.64, 10.33, 3.62, 3.57.

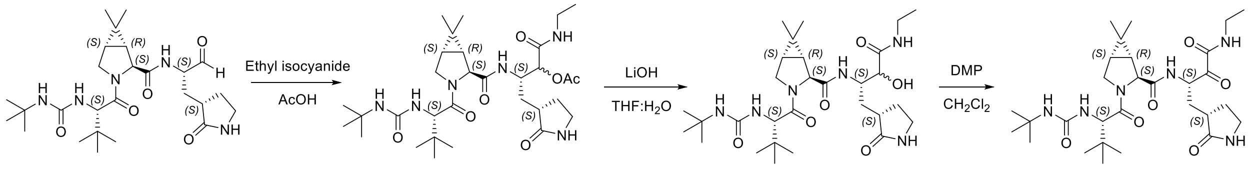


**Scheme 5**. The synthesis of compound **MPI33**

**(3S)-3-((1R,2S,5S)-3-((S)-2-(3-(tert-butyl)ureido)-3,3-dimethylbutanoyl)-6,6-dimethyl-3-azabicyclo[3.1.0]hexane-2-carboxamido)-1-(ethylamino)-1-oxo-4-((S)-2-oxopyrrolidin-3-yl)butan-2-yl acetate (33a).** To a solution of **MPI29** (120 mg, 0.24 mmol, 1.0 equiv.,) in anhydrous DCM (10 mL) at 0 °C. Then added ethyl isocyanide (21 μL, 0.28 mmol, 1.2 equiv.,) and acetic acid (30 μL, 0.48 mmol, 2.0 equiv.,). Then the reaction mixture was stirred at RT overnight. After the reaction was completed, remove the solvent in vacuum and purified by column chromatography (MeOH: DCM = 1:15 v/v) to yield **33a** as a white solid (140 mg, yield 96%). $^{1}$H NMR (400 MHz, Chloroform-*d*) δ 7.0 (d, $J$ = 9.6 Hz, 1H), 6.9 (s, 1H), 6.8 (s, 1H), 6.6 (t, $J$ = 5.7 Hz, 1H), 5.4 – 5.3 (m, 1H), 5.1 (dd, $J$ = 46.6, 4.6 Hz, 1H), 4.9 (d, $J$ = 10.2 Hz, 1H), 4.5 – 4.3 (m, 1H), 4.3 (d, $J$ = 9.9 Hz, 1H), 4.1 (d, $J$ = 7.5 Hz, 1H), 4.0 (d, $J$ = 10.3 Hz, 1H), 3.9 – 3.7 (m, 1H), 3.2 (qd, $J$ = 11.8, 9.9, 5.4 Hz, 4H), 2.5 – 2.3 (m, 2H), 2.1 (d, $J$ = 21.9 Hz, 3H), 1.7 (dq, $J$ = 11.9, 9.0 Hz, 1H), 1.5 – 1.3 (m, 2H), 1.3 (dd, $J$ = 7.7, 2.1 Hz, 1H), 1.2 (d, $J$ = 1.5 Hz, 9H), 1.1 – 1.0 (m, 3H), 0.9 (d, $J$ = 1.5 Hz, 3H), 0.9 (d, $J$ = 4.4 Hz, 9H), 0.8 (s, 3H). $^{13}$C NMR (100 MHz, Chloroform-*d*) δ 180.5, 172.6, 171.6, 169.7, 167.6, 157.4, 74.8, 61.0, 57.6, 50.0, 48.3, 40.4, 37.8, 34.7, 34.3, 32.8, 31.9, 30.6, 29.4, 28.2, 27.9, 26.5, 20.9, 19.1, 14.7, 12.6.

**(1R,2S,5S)-3-((S)-2-(3-(tert-butyl)ureido)-3,3-dimethylbutanoyl)-N-((2S)-4-(ethylamino)-3-hydroxy-4-oxo-1-((S)-2-oxopyrrolidin-3-yl)butan-2-yl)-6,6-dimethyl-3-azabicyclo[3.1.0]hexane-2-carboxamide(33b).** To a solution of **33a** (140 mg, 0.23 mmol, 1.0 equiv.,) in 3:1 MeOH/$H_2O$ (8 mL) was added LiOH.$H_2O$ (20 mg, 0.46 mmol, 2.0 equiv.,) at 0 °C.

The reaction was stirred at RT for 1 h. After completion, the reaction mixture was neutralized with 0.5 M HCl solution and remove the MeOH in vacuum. then extracted with DCM. The organic layer was washed with sat. NaCl, dried over $Na_2SO_4$ and concentrated. The residue was purified by column chromatography (MeOH: DCM = 1:10 v/v) to yield **33b** as a white solid (90 mg, yield 70%). $^1$H NMR (400 MHz, Chloroform-*d*) δ 7.1 (s, 2H), 6.9 (s, 1H), 5.6 (s, 1H), 5.4 (s, 1H), 5.0 (s, 1H), 4.3 (t, *J* = 11.4 Hz, 2H), 4.1 (d, *J* = 21.6 Hz, 2H), 4.0 (d, *J* = 10.4 Hz, 1H), 3.8 (dd, *J* = 10.4, 5.4 Hz, 1H), 3.2 (dt, *J* = 18.8, 8.6 Hz, 4H), 2.5 – 2.4 (m, 1H), 2.3 (s, 1H), 2.1 (d, *J* = 13.8 Hz, 1H), 1.7 (q, *J* = 10.8, 10.3 Hz, 1H), 1.5 (t, *J* = 11.6 Hz, 1H), 1.4 – 1.4 (m, 1H), 1.3 (d, *J* = 7.6 Hz, 1H), 1.2 (s, 9H), 1.1 (t, *J* = 7.2 Hz, 3H), 0.9 (s, 3H), 0.9 (s, 9H), 0.8 (s, 3H). $^{13}$C NMR (100 MHz, Chloroform-*d*) δ 181.3, 172.7, 172.1, 171.5, 157.5, 77.3, 73.1, 67.1, 61.0, 57.6, 50.0, 49.8, 48.4, 40.6, 38.1, 34.8, 34.1, 29.4, 28.1, 27.9, 26.6, 19.2, 14.9, 12.6.

**(1R,2S,5S)-3-((S)-2-(3-(tert-butyl)ureido)-3,3-dimethylbutanoyl)-N-((S)-4-(ethylamino)-3,4-dioxo-1-((S)-2-oxopyrrolidin-3-yl)butan-2-yl)-6,6-dimethyl-3-azabicyclo[3.1.0]hexane-2-carboxamide (MPI33).** To a solution of **33b** (90 mg, 0.16 mmol, 1.0 equiv.,) in anhydrous DCM (10 mL) was added Dess-Martin reagent (130 mg, 0.32 mmol, 2.0 equiv.,) slowly at 0 $^o$C. Then the reaction mixture was stirred at RT for 2 h. A solution of $NaHCO_3$ and $Na_2S_2O_3$ was added to quench the reaction. After 10 min, the mixture was washed with water, sat. NaCl, dried over $Na_2SO_4$ and concentrated. The residue was purified by column chromatography (MeOH: DCM = 1:10 v/v) to yield **MPI33** as a white solid (50 mg, yield 54%). $^1$H NMR (400 MHz, Methanol-$d_4$) δ 4.4 – 4.2 (m, 3H), 4.1 – 3.9 (m, 2H), 3.3 – 3.2 (m, 4H), 2.7 – 2.5 (m, 1H), 2.5 – 2.3 (m, 1H), 2.1 (qd, *J* = 13.8, 3.3 Hz, 1H), 2.0 – 1.6 (m, 2H), 1.6 (ddd, *J* = 18.5, 7.7, 5.2 Hz, 1H), 1.5 – 1.3 (m, 1H), 1.3 (d, *J* = 2.0 Hz, 9H), 1.2 (td, *J* = 7.2, 4.2 Hz, 3H), 1.1 (d, *J* = 10.3 Hz, 3H), 1.0 – 1.0 (m, 9H), 1.0 (d, *J* = 7.8 Hz, 3H). $^{13}$C NMR (100 MHz, Methanol-$d_4$) δ 195.4, 181.5, 172.3, 171.9, 170.5, 158.3, 60.6, 57.4, 52.1, 49.3, 48.5, 40.0, 37.6, 34.4, 34.1, 31.0, 29.3, 28.3, 27.8, 27.8, 25.7, 19.0, 13.6, 11.8.

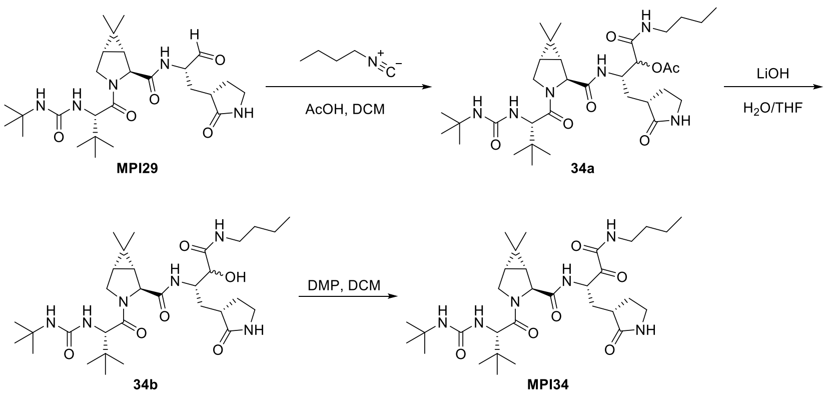


**Scheme 6**. The synthesis of compound **MPI34**

**(3S)-3-((1R,2S,5S)-3-((S)-2-(3-(tert-butyl)ureido)-3,3-dimethylbutanoyl)-6,6-dimethyl-3-azabicyclo[3.1.0]hexane-2-carboxamido)-1-(butylamino)-1-oxo-4-((S)-2-oxopyrrolidin-3-yl)butan-2-yl acetate (34a).** To a stirred solution of **MPI29** (90 mg, 0.18 mmol) in dichloromethane was added acetic acid (22 mg, 21 μL, 0.26 mmol) and n-butyl isocyanide (30 mg, 41 μL, 0.36 mmol). The reaction mixture was then stirred at room temperature overnight and concentrated *in vacuo*. The crude product was used without purification for the next step.

**(1R,2S,5S)-3-((S)-2-(3-(tert-butyl)ureido)-3,3-dimethylbutanoyl)-N-((2S)-4-(butylamino)-3-hydroxy-4-oxo-1-((S)-2-oxopyrrolidin-3-yl)butan-2-yl)-6,6-dimethyl-3-azabicyclo[3.1.0]hexane-2-carboxamide (34b).** To a solution of crude **34a** from last step in THF (5 mL) was added an aqueous solution of LiOH (36 mg, 0.72 mmol). The reaction mixture was stirred at room temperature for 3 h and then diluted with water (10 mL), extracted with dichloromethane (2×10 mL). The organic layers were combined and dried with anhydrous $Na_2SO_4$ and concentrated *in vacuo*. The crude product was used without further purification for the next step.

**(1R,2S,5S)-3-((S)-2-(3-(tert-butyl)ureido)-3,3-dimethylbutanoyl)-N-((S)-4-(butylamino)-3,4-dioxo-1-((S)-2-oxopyrrolidin-3-yl)butan-2-yl)-6,6-dimethyl-3-azabicyclo[3.1.0]hexane-2-carboxamide (MPI34).** To a solution of crude **34b** from last step in dichloromethane (5 mL) was added Dess-Martin periodinane (152 mg, 0.36 mmol) at 0 °C. The reaction mixture was added stirred at room temperature for 3 h. Then the reaction was quenched with a saturated $NaHCO_3$ solution containing 10 % $Na_2S_2O_3$. The layers were separated. The organic layer was then washed with saturated brine solution (2×10 mL), dried over anhydrous $Na_2SO_4$ and evaporated *in vacuo*. The residue was purified by flash chromatography (0-10% methanol in dichloromethane as eluent) to yield **MPI34** as white solid (55 mg, 51 %). $^1$H NMR (400 MHz, Chloroform-*d*) δ 7.77 (d, *J* = 7.0 Hz, 1H), 6.99 (t, *J* = 6.1 Hz, 1H), 6.27 (s, 1H), 5.37 (ddd, *J* = 10.5, 6.9, 3.4 Hz, 1H), 5.13 (s, 1H), 4.35 (s, 1H), 4.31 (s, 1H), 4.06 (d, *J* = 10.4 Hz, 1H), 3.87 (dt, *J* = 10.4, 2.7 Hz, 1H), 3.46 (s, 1H), 3.38 – 3.17 (m, 4H), 2.65 – 2.54 (m, 1H), 2.45 (tdd, *J* = 10.8, 7.5, 2.1 Hz, 1H), 2.09 – 1.90 (m, 3H), 1.58 – 1.43 (m, 4H), 1.39 – 1.29 (m, 2H), 1.25 (s, 9H), 1.01 (s, 3H), 0.98 – 0.82 (m, 15H). $^{13}$C NMR (101 MHz, $CDCl_3$) δ 195.6, 179.9, 172.7, 171.5, 159.3, 157.3, 60.6, 57.8, 53.3, 50.7, 50.2, 48.3, 40.4, 39.1, 38.4, 34.7, 32.7, 31.2, 30.3, 29.4, 28.2, 27.8, 26.5, 26.3, 20.0, 19.2, 13.7, 12.6.

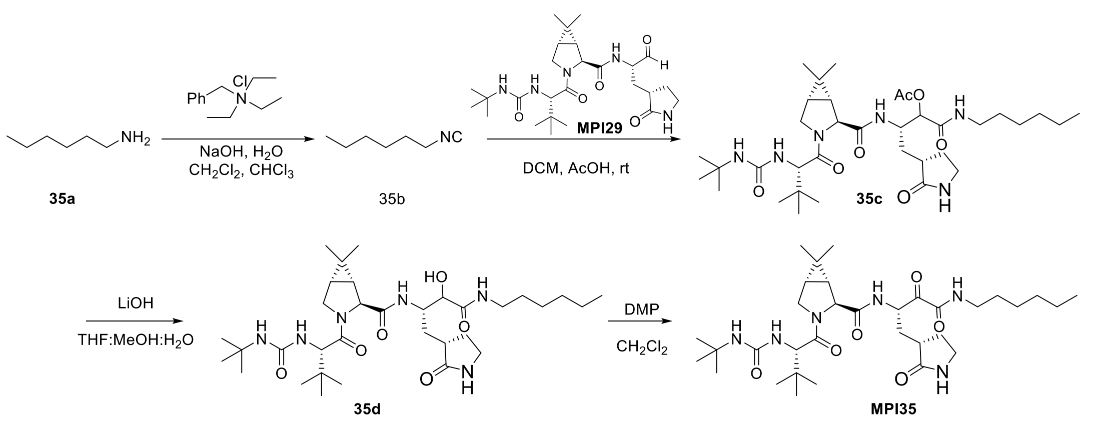


**Scheme 7**. The synthesis of compound **MPI35**

**(1R,2S,5S)-3-((S)-2-(3-(tert-butyl)ureido)-3,3-dimethylbutanoyl)-N-((S)-4-(hexylamino)-3,4-dioxo-1-((S)-2-oxopyrrolidin-3-yl)butan-2-yl)-6,6-dimethyl-3-azabicyclo[3.1.0]hexane-2-carboxamide (MPI35).** Amine **35a** (6.9 mmol, 1 equiv.,.) in dichloromethane (5.0 mL) was added to a solution of sodium hydroxide (1.66 g, 41 mmol, 6 equiv.,.) in water (2.5mL). Then N-benzyl-N,N,N-triethylammonium chloride (32 mg, 0.13 mmol, 0.02 equiv.,.) and chloroform (3.3 mL, 41 mmol, 6 equiv.,.) were added respectively. The mixture was stirred at room temperature (RT.) for 12 h. The organic layer was extracted with a syringe and directly purified by flash chromatography using dichloromethane to obtain the desired isocyanide. Eluent in a test tube (~10.0 mL) with most pungent odour was used directly in the next step. Aldehyde **MPI29** (0.2 mmol, 1 equiv.,.) was dissolved in the solution of isocyanide in dichloromethane, followed by addition of acetic acid (25 μL, 0.4 mmol, 2 equiv.,.). The mixture was stirred overnight at RT. All the volatiles were removed under vacuum and the residue was dissolved in a mixture of methanol and water (10.0 mL, v/v = 4/1). Lithium hydroxide monohydrate (19 mg, 0.16 mmol, 3 equiv.,.) was added in one portion and the resulting mixture was stirred at RT. for 1-4 h. Then it was neutralized by 0.1 M hydrochloric acid and concentrated under reduced pressure. The residue was extracted with Ethyl acetate (20.0 mL × 3) and the combined organic layers were dried over anhydrous magnesium sulfate, filtered, and concentrated under vacuum. The residue was dissolved in anhydrous dichloromethane (10.0 mL) and added Dess-Martin reagent (102.6 mg, 0.24 mmol, 1.5 equiv.,.) slowly at 0 °C. Then the reaction mixture was stirred at RT. for 1-2 h. A solution of sodium bicarbonate and sodium thiosulfate was added to quench the reaction. After 10 min, dichloromethane (10.0 mL × 2) was added to extract the mixture. The organic phase was washed with brine (10.0 mL × 2), dried over anhydrous sodium sulfate, filtered, and concentrated. The residue was purified by column chromatography (dichloromethane: methanol, 35:1 v/v) to afford the pure product as a white solid **MPI35** (70%). $^1$H NMR (400 MHz, Chloroform-*d*) δ 7.75 (d, *J* = 6.9 Hz, 1H), 6.95 (t, *J* = 6.2 Hz, 1H), 6.21 (s, 1H), 5.44 – 5.31 (m, 1H), 4.39 – 4.30 (m, 2H), 4.06 (d, *J* = 9.9 Hz, 1H), 3.86 (dd, *J* = 10.4, 5.0 Hz, 1H), 3.28 (dq, *J* = 13.5, 7.2, 6.7 Hz, 5H), 2.69 – 2.54 (m, 1H), 2.54 – 2.45 (m, 1H), 2.12 – 1.86 (m, 3H), 1.60 – 1.41 (m, 4H), 1.36 – 1.15 (m, 17H), 1.02 (s, 3H), 0.96 (s, 9H), 0.90 – 0.87 (m, 6H). $^{13}$C NMR (101 MHz, $CDCl_3$) δ 195.54, 179.77, 172.68, 171.41, 159.28, 157.23, 60.50, 57.82, 53.33, 50.78, 50.25, 48.30, 40.40, 39.45, 38.39, 34.73, 32.67, 31.37, 30.23, 29.36, 29.14, 28.28, 27.82, 26.53, 26.32, 22.50, 19.17, 14.00, 12.64.

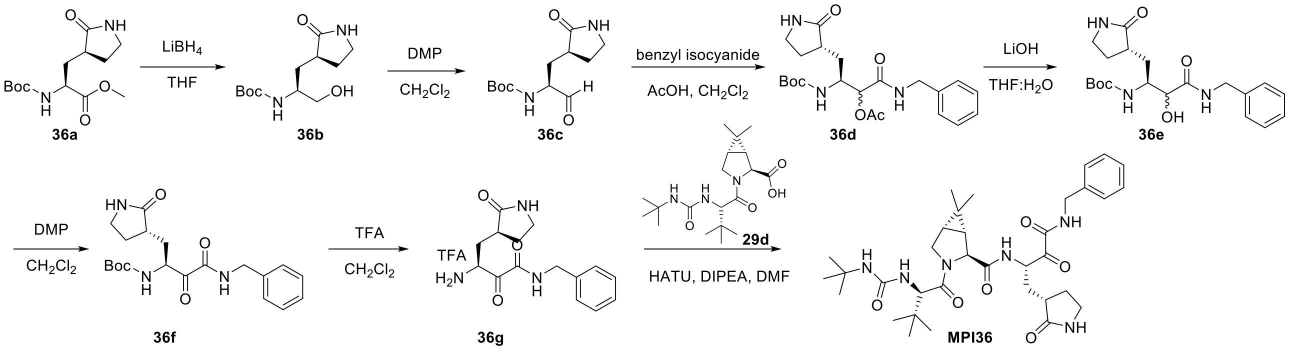


**Scheme 8**. The synthesis of compound **MPI36**.

**Tert-butyl ((S)-1-hydroxy-3-((S)-2-oxopyrrolidin-3-yl)propan-2-yl)carbamate (36b).**

To a solution of methyl (S)-2-((tert-butoxycarbonyl)amino)-3-((S)-2-oxopyrrolidin-3-yl)propanoate **36a** (400 mg, 1.4 mmol, 1.0 equiv.,.,) in anhydrous THF (10 mL) at 0 °C was added $LiBH_4$ (2.0 M in THF, 2.0 mL, 4.2 mmol, 3.0 equiv.,.,). The mixture was stirred at RT for 2 h. After the reaction was completed, excess reactants were consumed by slow addition of $H_2O$. The mixture was diluted with $H_2O$ and extracted with EtOAc, washed with sat. NaCl, dried over $Na_2SO_4$ and concentrated. The residue was purified by column chromatography (MeOH: EA = 1:10 v/v) to afford the pure product **36b** as a white solid (320 mg, yield 88%). $^1$H NMR (400 MHz, Methanol-*d*$_4$) δ 3.7 – 3.6 (m, 1H), 3.6 – 3.4 (m, 2H), 3.4 (s, 2H), 2.5 – 2.3 (m, 2H), 1.9 (q, *J* = 12.3, 10.4 Hz, 2H), 1.5 (s, 10H). $^{13}$C NMR (100 MHz, Methanol-*d*$_4$) δ 182.8, 158.3, 80.0, 65.8, 51.8, 41.5, 39.7, 34.0, 28.8, 28.7.

**Tert-butyl ((S)-1-oxo-3-((S)-2-oxopyrrolidin-3-yl)propan-2-yl)carbamate (36c).**

To a solution of **36b** (320 mg, 1.2 mmol, 1.0 equiv.) in anhydrous DCM (10 mL) was added Dess-Martin reagent (1.0 g, 2.4 mmol, 2.0 equiv.) slowly at 0 °C. Then the reaction mixture was stirred at RT for 2 h. A solution of $NaHCO_3$ and $Na_2S_2O_3$ was added to quench the reaction. After 10 min, the mixture was washed with water, sat. NaCl, dried over $Na_2SO_4$ and concentrated. The residue was purified by column chromatography (MeOH: DCM = 1:15 v/v) to yield **36c** as a white solid (230 mg, yield 72%). $^1$H NMR (400 MHz, Chloroform-*d*) δ 9.6 (s, 1H), 6.4 (s, 1H), 6.1 (d, *J* = 6.7 Hz, 1H), 4.2 (p, *J* = 6.2, 5.7 Hz, 1H), 3.4 – 3.3 (m, 2H), 2.5 (dd, *J* = 17.6, 11.1 Hz, 2H), 2.0 – 1.9 (m, 2H), 1.4 (d, *J* = 6.7 Hz, 10H). $^{13}$C NMR (100 MHz, Chloroform-*d*) δ 200.4, 180.2, 156.3, 80.2, 68.1, 58.8, 40.7, 38.0, 30.5, 28.4.

**(3S)-1-(benzylamino)-3-((tert-butoxycarbonyl)amino)-1-oxo-4-((S)-2-oxopyrrolidin-3-yl)butan-2-yl acetate (36d).** To a solution of **36c** (230 mg, 0.9 mmol, 1.0 equiv.,) in anhydrous DCM (10 mL) at 0 °C. Then added benzyl isocyanide (129 mg, 1.1 mmol, 1.2 equiv.,) and acetic acid (103 μL, 1.8 mmol, 2.0 equiv.,). Then the reaction mixture was stirred at RT overnight. After the reaction was completed, remove the solvent in vacuum and purified by column chromatography (MeOH: DCM = 1:15 v/v) to yield **36d** as a white solid (200 mg, yield 51%). $^1$H NMR (400 MHz, Chloroform-*d*) δ 7.3 – 7.1 (m, 5H), 6.8 (t, *J* = 5.8 Hz, 1H), 6.6 (s, 1H), 5.3 (d, *J* = 9.8 Hz, 1H), 5.2 (d, *J* = 3.6 Hz, 1H), 4.4 – 4.3 (m, 2H), 4.2 (ddt, *J* = 13.8, 7.2, 3.5 Hz, 1H), 3.3 – 3.1 (m, 2H), 2.4 – 2.2 (m, 2H), 2.1 (s, 3H), 1.9 (ddd, *J* = 17.3, 9.5, 3.5 Hz, 1H), 1.8 – 1.6 (m,

1H), 1.3 (s, 10H). $^{13}C$ NMR (100 MHz, Chloroform-*d*) δ 180.2, 169.6, 167.9, 155.7, 137.9, 128.7, 127.7, 127.5, 79.7, 74.9, 53.5, 50.0, 43.3, 40.4, 38.0, 33.4, 28.3, 20.8.

**Tert-butyl ((2S)-4-(benzylamino)-3-hydroxy-4-oxo-1-((S)-2-oxopyrrolidin-3-yl) butan-2-yl) carbamate (36e).** To a solution of **36d** (200 mg, 0.5 mmol, 1.0 equiv.,) in 3:1 MeOH/$H_2O$ (8 mL) was added LiOH.$H_2O$ (42 mg, 1.0 mmol, 2.0 equiv.,) at 0 °C. The reaction was stirred at RT for 1 h. After completion, the reaction mixture was neutralized with 0.5 M HCl solution and remove the MeOH in vacuum. then extracted with DCM. The organic layer was washed with sat. NaCl, dried over $Na_2SO_4$ and concentrated. The residue was purified by column chromatography (MeOH: DCM = 1:15 v/v) to yield **36e** as a white solid (160 mg, yield 89%). $^1H$ NMR (400 MHz, Chloroform-*d*) δ 7.4 (t, *J* = 6.1 Hz, 1H), 7.2 (dq, *J* = 15.5, 8.2, 7.4 Hz, 5H), 6.6 (s, 1H), 5.6 (d, *J* = 5.2 Hz, 1H), 5.5 (d, *J* = 9.3 Hz, 1H), 4.4 (dd, *J* = 14.9, 6.3 Hz, 1H), 4.3 (dd, *J* = 14.9, 5.5 Hz, 1H), 4.2 – 4.0 (m, 2H), 3.2 (dq, *J* = 17.2, 9.1 Hz, 2H), 2.4 – 2.2 (m, 2H), 2.1 – 1.9 (m, 1H), 1.7 (dq, *J* = 17.3, 9.1 Hz, 1H), 1.6 (p, *J* = 5.8 Hz, 1H), 1.3 (s, 9H). $^{13}C$ NMR (100 MHz, Chloroform-*d*) δ 181.0, 172.4, 156.1, 138.1, 128.6, 127.7, 127.4, 79.6, 73.3, 53.5, 51.1, 43.1, 40.6, 38.0, 32.9, 28.3.

**Tert-butyl ((S)-4-(benzylamino)-3,4-dioxo-1-((S)-2-oxopyrrolidin-3-yl)butan-2-yl)carbamate (36f).** To a solution of **36e** (160 mg, 0.4 mmol, 1.0 equiv.,) in anhydrous DCM (10 mL) was added Dess-Martin reagent (340 mg, 0.8 mmol, 2.0 equiv.,) slowly at 0 °C. Then the reaction mixture was stirred at RT for 2 h. A solution of $NaHCO_3$ and $Na_2S_2O_3$ was added to quench the reaction. After 10 min, the mixture was washed with water, sat. NaCl, dried over $Na_2SO_4$ and concentrated. The residue was purified by column chromatography (MeOH: DCM = 1:15 v/v) to yield **36f** as a white solid (150 mg, yield 94%). $^1H$ NMR (400 MHz, Chloroform-*d*) δ 7.4 (t, *J* = 6.4 Hz, 1H), 7.3 – 7.2 (m, 5H), 6.7 (s, 1H), 5.9 (d, *J* = 7.7 Hz, 1H), 5.0 (ddd, *J* = 11.4, 7.7, 3.4 Hz, 1H), 4.4 (d, *J* = 6.2 Hz, 2H), 3.3 – 3.2 (m, 2H), 2.5 (qd, *J* = 8.8, 5.4 Hz, 1H), 2.4 (d, *J* = 8.5 Hz, 1H), 2.3 – 2.2 (m, 1H), 1.9 – 1.8 (m, 2H), 1.3 (d, *J* = 3.8 Hz, 9H). $^{13}C$ NMR (100 MHz, Chloroform-*d*) δ 196.3, 180.1, 159.4, 155.8, 136.9, 128.8, 127.9, 127.9, 79.9, 53.5, 50.6, 43.4, 40.5, 38.5, 33.1, 28.3.

**(1R,2S,5S)-N-((S)-4-(benzylamino)-3,4-dioxo-1-((S)-2-oxopyrrolidin-3-yl)butan-2-yl)-3-((R)-2-(3-(tert-butyl)ureido)-3,3-dimethylbutanoyl)-6,6-dimethyl-3-azabicyclo[3.1.0]hexane-2-carboxamide (MPI36).** To a solution of **36f** (70 mg, 0.18 mmol, 1.0

equiv.,) in anhydrous DCM (5 mL) at 0 °C, and then TFA (140 μL, 1.8 mmol, 10 equiv.,) was added. The mixture was stirred for 2 h. After the reaction was completed, remove the solvent in vacuum. The residue was dissolved in anhydrous DMF at 0 °C, and then (1R,2S,5S)-3-((R)-2-(3-(tert-butyl)ureido)-3,3-dimethylbutanoyl)-6,6-dimethyl-3-azabicyclo[3.1.0]hexane-2-carboxylic acid **29d** (81 mg, 0.22 mmol, 1.2 equiv.,), HATU (103 mg, 0.27 mmol, 1.5 equiv.,), DIPEA (160 μL, 0.9 mmol, 5.0 equiv.,) was added sequentially. The mixture was stirred at RT overnight. The mixture was diluted with EtOAc and washed with water, 1M HCl, sat. NaCl, dried over $Na_2SO_4$, and concentrated. The residue was purified by column chromatography (MeOH: DCM = 1:15 v/v) to afford the pure product **MPI36** as a white solid (75 mg, yield 65%). $^1$H NMR (400 MHz, Methanol-$d_4$) δ 7.4 – 7.3 (m, 4H), 7.3 – 7.2 (m, 1H), 6.0 (d, $J$ = 8.7 Hz, 1H), 5.8 (d, $J$ = 10.0 Hz, 1H), 4.4 (s, 1H), 4.3 (dt, $J$ = 10.4, 2.6 Hz, 1H), 4.2 (s, 1H), 4.0 (dd, $J$ = 10.2, 4.2 Hz, 1H), 3.9 (dt, $J$ = 10.1, 4.9 Hz, 1H), 3.3 (s, 1H), 3.2 (dt, $J$ = 13.7, 5.3 Hz, 1H), 2.6 (p, $J$ = 11.3 Hz, 1H), 2.4 – 2.3 (m, 1H), 2.3 – 2.0 (m, 2H), 1.7 – 1.6 (m, 1H), 1.5 (dtd, $J$ = 15.1, 7.7, 3.6 Hz, 1H), 1.3 (s, 10H), 1.1 – 0.9 (m, 15H). $^{13}$C NMR (100 MHz, Chloroform-$d$) δ 195.5, 180.0, 172.8, 171.5, 159.3, 157.4, 136.8, 128.9, 128.7, 127.9, 60.6, 57.8, 53.3, 50.1, 48.3, 44.1, 43.4, 40.5, 38.4, 34.7, 32.7, 29.4, 28.2, 26.6, 26.3, 19.2, 12.7.

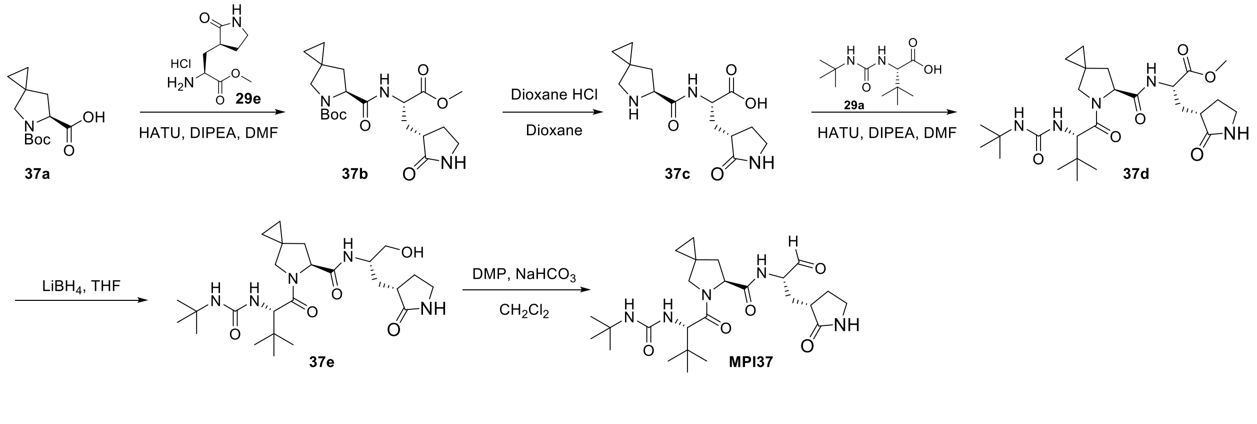


**Scheme 9**: The synthesis of compound **MPI37**

**Synthesis of tert-butyl (S)-6-(((S)-1-methoxy-1-oxo-3-((S)-2-oxopyrrolidin-3-yl)propan-2-yl)carbamoyl)-5-azaspiro[2.4]heptane-5-carboxylate (37b).** To a solution of **37a** (2.24 mmol, 0.54 g) and **29e** (2.47 mmol, 0.55 g) in anhydrous DMF (10 mL) was added DIPEA (8.96 mmol, 1.26 g) and was cooled to 0 °C. HATU (3 mmol, 1.1 g) was added to the solution under 0 °C and then stirred at room temperature overnight. The reaction mixture was then diluted with ethyl acetate (50 mL) and washed with saturated $NaHCO_3$ solution (2×20 mL), 1 M HCl solution (2×20 mL), and saturated brine solution (2×20 mL) sequentially. The organic layer was dried over anhydrous $Na_2SO_4$ and then concentrated *on vacuo*. The residue was then purified with flash chromatography (50-100% EtOAc in hexanes as the eluent) to afford **37b** as white solid (440 mg, 48%). $^1$H NMR (400 MHz, Methanol-$d_4$) δ 4.65 – 4.44 (m, 1H), 4.39 (d, $J$ = 8.7 Hz, 1H), 3.72 (d, $J$ = 2.0 Hz, 3H), 3.32 (t, $J$ = 1.6 Hz, 2H), 2.51 – 2.29 (m, 2H), 2.27 – 2.04 (m, 1H), 1.95 – 1.69 (m, 3H), 1.45 (s, 9H), 0.65 – 0.45 (m, 4H). $^{13}$C NMR (100 MHz, Methanol-$d_4$) δ 180.09, 174.46, 172.32, 172.11, 150.77, 128.61, 120.78, 80.27, 79.86, 78.23, 77.90, 77.58, 60.94, 53.82, 53.47, 51.56, 51.05, 50.49, 47.56, 47.35, 47.14, 40.17, 39.13, 38.67, 38.18, 32.51, 27.47, 27.38, 20.46, 19.69, 12.27, 11.23.

**Synthesis of methyl (S)-3-((S)-2-oxopyrrolidin-3-yl)-2-((S)-5-azaspiro[2.4]heptane-6-carboxamido)propanoate (37c).** To a solution of **37b** (0.44 g, 0.93 mmol) in 1,4-dioxane (10 mL) was added dropwise a HCl solution in 1,4-dioxane (4 M, 10 mL). The resulting solution was stirred at room temperature for 1 h. Then residue was then concentrated *on vacuo* to afford **37c** as light-yellow hygroscopic solid (0.3 g, 90%). $^1$H NMR (400 MHz, Methanol-$d_4$) δ 4.75 – 4.37 (m, 1H), 3.77 (d, $J$ = 3.8 Hz, 5H), 3.49 – 3.19 (m, 6H), 2.70 – 2.31 (m, 4H), 2.31 – 2.05 (m, 2H), 2.03 – 1.73 (m, 2H), 0.86 – 0.70 (m, 4H). $^{13}$C NMR (100 MHz, Methanol-$d_4$) δ 180.20, 171.88, 150.73, 72.18, 71.06, 66.77, 60.82, 52.42, 51.71, 48.34, 48.14, 48.12, 47.93, 47.91, 47.72, 47.70, 47.50, 47.48, 47.29, 47.27, 47.06, 38.39, 37.54, 27.23, 20.22, 10.05, 8.67.

**Synthesis of methyl (S)-2-((S)-5-((S)-2-(3-(tert-butyl)ureido)-3,3-dimethylbutanoyl)-5-azaspiro[2.4]heptane-6-carboxamido)-3-((S)-2-oxopyrrolidin-3-yl)propanoate (37d)**. To a solution of **37c** (1.18 mmol, 0.5 g) and **29a** (1.3 mmol, 0.3 g) in anhydrous DMF (15 mL) was added DIPEA (4.7 mmol, 0.82 mL) and was cooled to 0 °C. HATU (1.53 mmol, 0.58 g) was added at 0 °C and then stirred at room temperature overnight. The reaction mixture was then diluted with ethyl acetate (50 mL) and washed with saturated $NaHCO_3$ solution (2×20 mL), 1 M HCl solution (2×20 mL), and saturated brine solution (2×20 mL) sequentially. The organic layer was dried over anhydrous $Na_2SO_4$ and then concentrated *on vacuo*. The residue was then purified with flash chromatography (MeOH: DCM = 1:10 v/v) to afford **37d** as white solid (400 mg, 53%). $^1$H NMR (400 MHz, Chloroform-$d$) δ 7.49 (d, $J$ = 8.3 Hz, 1H), 6.51 (s, 1H), 5.70 (s, 1H), 4.70 – 4.55 (m,

1H), 4.48 (dd, $J$ = 8.1, 6.3 Hz, 1H), 4.36 (s, 1H), 3.75 – 3.66 (m, 1H), 3.65 (s, 3H), 3.59 (t, $J$ = 9.0 Hz, 1H), 3.23 (s, 0H), 3.11 (dd, $J$ = 7.5, 4.4 Hz, 1H), 2.68 – 2.44 (m, 1H), 2.44 – 2.27 (m, 1H), 2.06 (dd, $J$ = 12.6, 6.3 Hz, 1H), 1.89 (dd, $J$ = 12.6, 8.1 Hz, 1H), 1.77 (dd, $J$ = 10.3, 2.1 Hz, 1H), 1.19 (s, 10H), 0.91 (s, 11H), 0.60 – 0.39 (m, 4H).

**Synthesis of (S)-5-((S)-2-(3-(tert-butyl)ureido)-3,3-dimethylbutanoyl)-N-((S)-1-hydroxy-3-((S)-2-oxopyrrolidin-3-yl)propan-2-yl)-5-azaspiro[2.4]heptane-6-carboxamide (37e).** To a solution of **37d** (180 mg, 0.3 mmol, 1.0 equiv.,) in anhydrous THF (10 mL) at 0 °C was added $LiBH_4$ (2.0 M in THF, 0.7 mL, 1.4 mmol, 5.0 equiv.,). The mixture was stirred at RT for 2 h. After the reaction was completed, excess reactants were consumed by slow addition of $H_2O$. The mixture was diluted with $H_2O$ and extracted with EtOAc, washed with sat. NaCl, dried over $Na_2SO_4$ and concentrated. The residue was purified by column chromatography (MeOH: DCM = 1:10 v/v) to afford the pure product **37e** as a white solid (85 mg, yield 49%). $^1$H NMR (400 MHz, Chloroform-*d*) δ 4.70 (d, $J$ = 7.6 Hz, 1H), 4.49 (t, $J$ = 7.2 Hz, 2H), 4.35 (s, 2H), 3.83 – 3.64 (m, 3H), 3.59 (s, 1H), 3.34 – 3.15 (m, 6H), 2.50 (s, 2H), 2.36 (d, $J$ = 6.2 Hz, 4H), 2.09 – 1.95 (m, 3H), 1.89 (dd, $J$ = 12.5, 8.0 Hz, 2H), 1.85 – 1.62 (m, 3H), 1.56 – 1.37 (m, 1H), 1.22 (d, $J$ = 5.5 Hz, 26H), 0.92 (s, 27H), 0.57 (d, $J$ = 3.4 Hz, 6H).

**Synthesis of (S)-5-((S)-2-(3-(tert-butyl)ureido)-3,3-dimethylbutanoyl)-N-((S)-1-oxo-3-((S)-2-oxopyrrolidin-3-yl)propan-2-yl)-5-azaspiro[2.4]heptane-6-carboxamide (MPI37)**. To a solution of **37e** (85 mg, 0.14 mmol, 1.0 equiv.,) in anhydrous DCM (10 mL) was added Dess-Martin reagent (182 mg, 0.42 mmol, 3.0 equiv.,) slowly at 0 °C. Then the reaction mixture was stirred at RT for 2 h. A solution of $NaHCO_3$ and $Na_2S_2O_3$ was added to quench the reaction. After 10 min, the mixture was washed with water, sat. NaCl, dried over $Na_2SO_4$ and concentrated. The residue was purified by column chromatography (MeOH: DCM = 1:10 v/v) to yield **MPI37** as a white solid (67 mg, yield 79 %). $^1$H NMR (400 MHz, Chloroform-*d*) δ 9.48 (s, 1H), 7.86 (d, $J$ = 7.7 Hz, 1H), 6.53 (s, 1H), 5.46 (d, $J$ = 9.6 Hz, 1H), 4.89 (s, 1H), 4.57 – 4.42 (m, 1H), 4.34 (d, $J$ = 9.7 Hz, 1H), 3.77 – 3.52 (m, 2H), 3.38 – 3.10 (m, 2H), 2.70 – 2.46 (m, 8H), 2.13 – 1.71 (m, 1H), 1.21 (s, 12H), 0.91 (d, $J$ = 5.2 Hz, 15H), 0.62 – 0.38 (m, 6H). $^{13}$C NMR (100 MHz, Chloroform-*d*) δ 199.99, 180.49, 172.67, 172.56, 157.18, 60.89, 57.21, 57.01, 56.12, 50.17, 40.50, 37.69, 37.41, 35.52, 30.45, 29.49, 28.19, 26.45, 21.57.

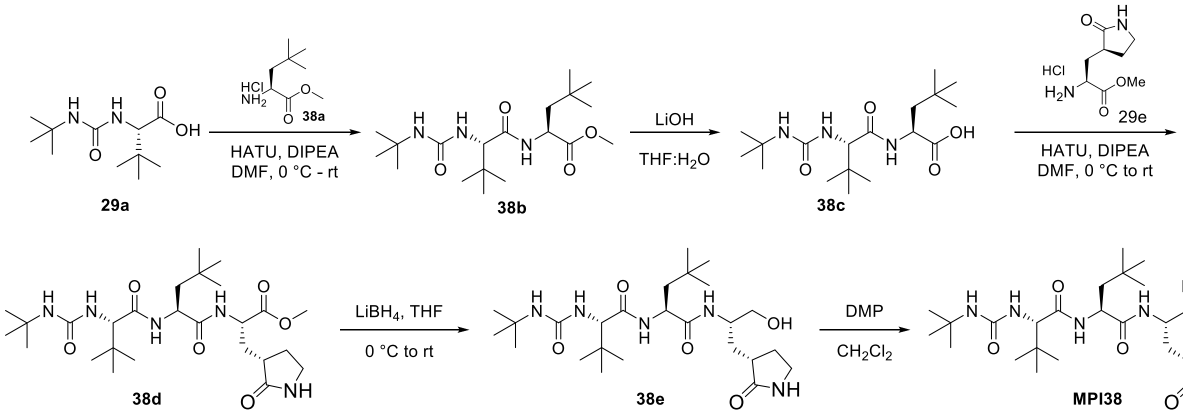

**Scheme 10**: The synthesis of compound **MPI38**.

**(S)-methyl 2-((S)-2-(3-(tert-butyl)ureido)-3,3-dimethylbutanamido)-4,4-dimethylpentanoate (38b):** The amino acid methyl ester hydrochloride **38a** (0.4 g, 1.74 mmol) and **29a** (406 g, 2.08 mmol) were dissolved in dry DMF (20 mL) and the reaction was cooled to 0 °C. HATU (0.86 g, 2.26 mmol) and DIPEA (1.24 mL, 6.95 mmol) were added, and the reaction mixture was allowed warm up to room temperature and stirred for 12 h. The mixture was then poured into water (50 mL) and extracted with ethyl acetate (4×20 mL). The organic layer was washed with aqueous hydrochloric acid 10% v/v (2×20 mL), saturated aqueous $NaHCO_3$ (2×20 mL), brine (2×20 mL) and dried over $Na_2SO_4$. The organic phase was evaporated to dryness and the crude material purified by silica gel column chromatography (15-50% EtOAc in hexanes as the eluent) to afford **38b** white solid (460 mg, 71%). $^1$H NMR (400 MHz, Chloroform-*d*) δ 6.48 (d, *J* = 7.9 Hz, 1H), 5.36 (d, *J* = 9.3 Hz, 1H), 4.70 (s, 1H), 4.55 (td, *J* = 8.0, 4.3 Hz, 1H), 4.09 (d, *J* = 9.4 Hz, 1H), 3.68 (s, 3H), 1.77 (dd, *J* = 14.3, 4.3 Hz, 1H), 1.49 (dd, *J* = 14.3, 8.1 Hz, 1H), 1.28 (s, 9H), 0.98 (s, 9H), 0.91 (s, 9H). $^{13}$C NMR (101 MHz, $CDCl_3$): δ 173.51, 171.85, 157.26, 61.30, 52.12, 50.19, 45.93, 34.63, 30.63, 29.55, 29.48, 26.68.

**(S)-2-((S)-2-(3-(tert-butyl)ureido)-3,3-dimethylbutanamido)-4,4-dimethylpentanoic acid (38c).** The peptide **38b** (450 mg, 1.2 mmol) was dissolved in THF/$H_2O$ (1:1, 20 mL). LiOH (120 mg, 3.02 mmol) was added at 0 °C. The mixture was stirred at room temperature overnight. Then THF was removed *on vacuum* and the aqueous layer was acidified with 1 M HCl and extracted with dichloromethane (3 x 20 mL). The organic layer was dried over anhydrous $Na_2SO_4$ and concentrated to yield **38c** as white solid (380 mg, 85%). $^{1}H$ NMR (400 MHz, DMSO-$d_6$) δ 12.37 (s, 1H), 8.16 (d, *J* = 8.1 Hz, 1H), 5.98 (s, 1H), 5.84 (d, *J* = 9.8 Hz, 1H), 4.30 (td, *J* = 8.5, 3.2 Hz, 1H), 4.01 (d, *J* = 9.8 Hz, 1H), 1.75 – 1.48 (m, 2H), 1.18 (s, 9H), 0.88 (s, 9H), 0.87 (s, 9H).

**(6S,9S,12S)-methyl-6-(tert-butyl)-2,2-dimethyl-9-neopentyl-4,7,10-trioxo-12-(((S)-2-oxopyrrolidin-3-yl) methyl)-3,5,8,11-tetraazatridecan-13-oate (38d).** The amino acid methyl ester hydrochloride **29e** (136 mg, 0.616 mmol) and **38c** (200 mg, 0.56 mmol) were dissolved in dry DMF (10 mL) and the reaction was cooled to 0 °C. HATU (252 mg, 0.672 mmol) and DIPEA (0.4 mL, 2.24 mmol) were added, and the reaction mixture was allowed warm up to room temperature and stirred for 12 h. The mixture was then poured into water (50 mL) and extracted with ethyl acetate (4×20 mL). The organic layer was washed with aqueous hydrochloric acid 10% v/v (2×20 mL), saturated aqueous $NaHCO_3$ (2×20 mL), brine (2×20 mL) and dried over $Na_2SO_4$. The organic phase was evaporated to dryness and the crude material purified by silica gel column chromatography (0-10% MeOH in $CH_2Cl_2$ as the eluent) to afford **38d** white gummy solid (220 mg, 74%). $^{1}H$ NMR (400 MHz, Chloroform-*d*) δ 8.07 (s, 1H), 7.76 (s, 1H), 7.56 (d, *J* = 8.9 Hz, 1H), 4.83 (s, 1H), 4.68 (ddd, *J* = 12.1, 8.8, 3.3 Hz, 1H), 4.59 (dd, *J* = 9.3, 3.3 Hz, 1H), 4.07 (d, *J* = 9.7 Hz, 1H), 3.68 (s, 3H), 3.45 – 3.30 (m, 2H), 2.50 – 2.33 (m, 2H), 2.30 – 2.19 (m, 1H), 2.19 – 2.06 (m, 1H), 1.94 – 1.73 (m, 3H), 1.27 (s, 9H), 0.96 (s, 9H), 0.90 (s, 9H). $^{13}C$ NMR (101 MHz, $CDCl_3$): δ 179.98, 173.02, 172.15, 171.65, 157.32, 61.01, 55.43, 53.44, 51.21, 50.01, 46.76, 40.56, 38.27, 34.48, 34.29, 30.58, 29.66, 29.48, 29.43, 27.73, 26.55.

**(S)-2-((S)-2-(3-(tert-butyl)ureido)-3,3-dimethylbutanamido)-N-((S)-1-hydroxy-3-((S)-2-oxopyrrolidin-3-yl)propan-2-yl)-4,4-dimethylpentanamide (38e).** To a stirred solution of compound **38d** (200 mg, 0.381 mmol) in THF (8 mL) was added $LiBH_4$ (2.0 M in THF, 1.2 mL, 1.14 mmol) in several portions at 0 °C under a nitrogen atmosphere. The reaction mixture was stirred at 0 °C for 1 h, then allowed to warm up to room temperature, and stirred for an additional 2 h. The reaction was quenched by the drop wise addition of 1.0 M HCl (aq) (1.2 mL) with cooling in an ice bath. The solution was diluted with ethyl acetate and $H_2O$. The phases were separated, and the aqueous layer was extracted with ethyl acetate (3×15 mL). The organic phases were combined, dried over $MgSO_4$, filtered, and concentrated on a rotavapor to give a yellow oily residue. Column chromatographic purification of the residue (6% MeOH in $CH_2Cl_2$ as the eluent) afforded **38e** as a white solid (150 mg, 79%). [1]H NMR (400 MHz, Chloroform-*d*) δ 5.82 (d, *J* = 9.2 Hz, 1H), 4.33 (dd, *J* = 8.0, 4.5 Hz, 2H), 3.94 – 3.85 (m, 2H), 3.55 – 3.37 (m, 2H), 3.24 – 3.14 (m, 2H), 2.47 – 2.35 (m, 1H), 2.34 – 2.24 (m, 1H), 1.95 – 1.87 (m, 1H), 1.78 – 1.64 (m, 2H), 1.60 – 1.42 (m, 2H), 1.23 (s, 9H), 0.90 (s, 18H).

**(S)-2-((S)-2-(3-(tert-butyl)ureido)-3,3-dimethylbutanamido)-4,4-dimethyl-N-((S)-1-oxo-3-((S)-2-oxopyrrolidin-3-yl)propan-2-yl)pentanamide (MPI38).** To a solution of **38e** (120 mg, 0.241 mmol) in $CH_2Cl_2$ (6 mL) was added $NaHCO_3$ (83 mg, 4 equiv.,) and the Dess-Martin reagent (314 mg, 0.724 mmol, 3 equiv.). The resulting mixture was stirred at rt for 12 h. Then the reaction was quenched with a saturated $NaHCO_3$ solution containing 10 % $Na_2S_2O_3$. The layers were separated. The organic layer was then washed with saturated brine solution, dried over anhydrous $Na_2SO_4$, and concentrated *on vacuum*. The residue was then purified with flash chromatography afford **MPI38** as white solid (80 mg, 67%). [1]H NMR (400 MHz, Chloroform-*d*) δ 9.45 (s, 1H), 4.63 – 4.24 (m, 2H), 3.92 (d, *J* = 25.5 Hz, 1H), 3.36 – 3.25 (m, 2H), 2.55 – 2.33 (m, 2H), 1.97 – 1.72 (m, 3H), 1.54 – 1.41 (m, 2H), 1.23 (s, 9H), 0.91 (s, 9H), 0.87 (s, 9H). [13]C NMR (101 MHz, DMSO): δ 201.04, 178.62, 173.44, 172.54, 171.37, 157.54, 157.45, 60.02, 56.54, 50.49, 49.39, 45.72, 37.48, 34.78, 30.68, 29.98, 29.85, 29.73, 27.69, 27.04.

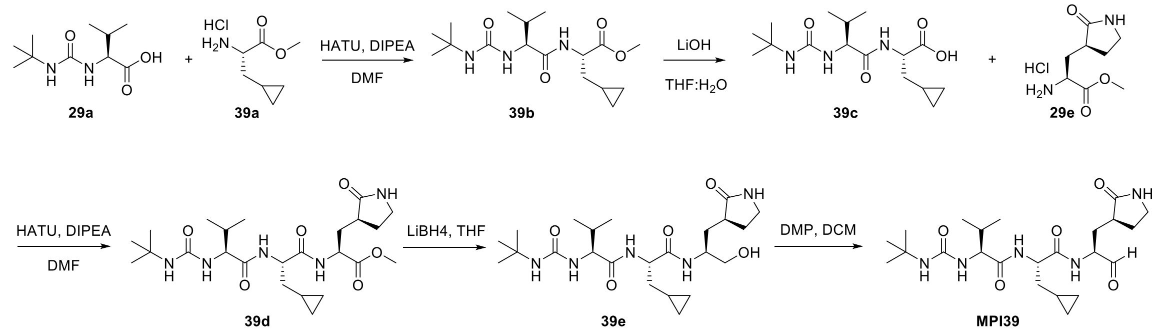


**Scheme S11**: The synthesis of compound **MPI39**.

**Methyl (S)-2-((S)-2-(3-(tert-butyl)ureido)-3-methylbutanamido)-3-cyclopropylpropanoate (39b)**. **39b** was prepared as a white solid following a similar procedure to **29c**. (yield 62%). [1]H NMR (400 MHz, Methanol-$d_4$) δ 4.5 (dd, *J* = 7.8, 5.7 Hz, 1H), 4.1 – 4.0 (m, 1H), 3.7 (s, 3H), 2.1 – 2.0 (m, 1H), 1.8 – 1.7 (m, 1H), 1.6 (ddd, *J* = 13.8, 7.5, 5.8 Hz, 1H), 1.3 (s, 9H), 1.0 (dd, *J* = 22.1,

6.8 Hz, 6H), 0.9 – 0.7 (m, 1H), 0.6 – 0.4 (m, 2H), 0.2 – 0.0 (m, 2H). $^{13}$C NMR (100 MHz, Methanol-$d_4$) δ 175.0, 174.0, 159.8, 59.6, 54.2, 52.5, 50.8, 37.6, 32.5, 29.7, 19.8, 18.2, 8.4, 5.1, 4.7.

**(S)-2-((S)-2-(3-(tert-butyl)ureido)-3-methylbutanamido)-3-cyclopropylpropanoic acid (39c). 39c** was prepared as a white solid following a similar procedure to **29d** (yield 83%), the residue was used in the next step without further purification.

**Methyl (6S,9S,12S)-9-(cyclopropylmethyl)-6-isopropyl-2,2-dimethyl-4,7,10-trioxo-12-(((S)-2-oxopyrrolidin-3-yl)methyl)-3,5,8,11-tetraazatridecan-13-oate (39d). 39d** was prepared as a white solid following a similar procedure to **29f** (yield 55%). $^1$H NMR (400 MHz, Methanol-$d_4$) δ 4.5 (dd, *J* = 11.8, 4.2 Hz, 1H), 4.4 (d, *J* = 7.2 Hz, 1H), 4.0 (d, *J* = 6.1 Hz, 1H), 3.7 (d, *J* = 5.3 Hz, 3H), 3.3 (s, 2H), 2.6 – 2.5 (m, 1H), 2.3 (dt, *J* = 14.4, 5.7 Hz, 1H), 2.2 (td, *J* = 13.1, 11.8, 4.1 Hz, 1H), 2.1 (dq, *J* = 13.7, 6.9 Hz, 1H), 1.8 (q, *J* = 10.0 Hz, 2H), 1.6 (ddt, *J* = 21.3, 14.4, 7.3 Hz, 2H), 1.3 (d, *J* = 6.3 Hz, 9H), 0.9 (dd, *J* = 22.8, 6.9 Hz, 7H), 0.5 (ddd, *J* = 18.9, 9.6, 5.1 Hz, 2H), 0.2 – -0.0 (m, 2H). $^{13}$C NMR (100 MHz, Methanol-$d_4$) δ 181.7, 174.8, 174.4, 173.5, 159.8, 59.7, 55.2, 52.8, 51.7, 50.8, 41.4, 39.4, 38.1, 33.9, 32.4, 29.7, 28.7, 19.9, 18.1, 8.3, 5.1, 5.0.

**(S)-2-(3-(tert-butyl)ureido)-N-((S)-3-cyclopropyl-1-(((S)-1-hydroxy-3-((S)-2-oxopyrrolidin-3-yl)propan-2-yl)amino)-1-oxopropan-2-yl)-3-methylbutanamide (39e). 39e** was prepared as a white solid following a similar procedure to **37e** (yield 86%). $^1$H NMR (400 MHz, Methanol-$d_4$) δ 4.4 (q, *J* = 7.2 Hz, 1H), 4.0 (d, *J* = 5.9 Hz, 2H), 3.6 – 3.4 (m, 2H), 3.3 – 3.2 (m, 2H), 2.6 – 2.4 (m, 1H), 2.4 (dddd, *J* = 12.5, 8.5, 7.0, 2.7 Hz, 1H), 2.2 – 1.9 (m, 2H), 1.8 – 1.7 (m, 2H), 1.6 – 1.5 (m, 2H), 1.3 (s, 9H), 0.9 (dd, *J* = 23.1, 6.8 Hz, 6H), 0.8 (qdd, *J* = 8.0, 6.5, 3.9 Hz, 1H), 0.6 – 0.4 (m, 2H), 0.2 – 0.0 (m, 2H). $^{13}$C NMR (100 MHz, Methanol-$d_4$) δ 182.6, 175.0, 174.4, 159.9, 65.5, 60.1, 55.8, 50.8, 50.6, 41.5, 39.5, 38.1, 33.7, 32.2, 29.8, 29.0, 19.9, 18.1, 8.5, 5.1, 5.0.

**(S)-2-(3-(tert-butyl)ureido)-N-((S)-3-cyclopropyl-1-oxo-1-(((S)-1-oxo-3-((S)-2-oxopyrrolidin-3-yl)propan-2-yl)amino)propan-2-yl)-3-methylbutanamide (MPI39). MPI39** was prepared as a white solid following a similar procedure to **MPI29** (yield 78%). $^1$H NMR (400 MHz, Chloroform-*d*) δ 9.5 (s, 1H), 7.8 – 7.7 (m, 1H), 6.6 (d, *J* = 23.0 Hz, 1H), 5.5 (s, 1H), 5.2 – 5.1 (m,

1H), 4.7 – 4.4 (m, 2H), 4.2 – 4.0 (m, 1H), 3.3 (dd, $J$ = 10.5, 5.4 Hz, 3H), 2.4 (s, 2H), 2.0 (td, $J$ = 12.0, 11.1, 6.2 Hz, 2H), 1.7 (dd, $J$ = 46.1, 5.3 Hz, 4H), 1.3 (s, 9H), 1.0 – 0.8 (m, 6H), 0.7 (s, 1H), 0.4 (t, $J$ = 7.6 Hz, 2H), 0.2 – 0.0 (m, 2H).

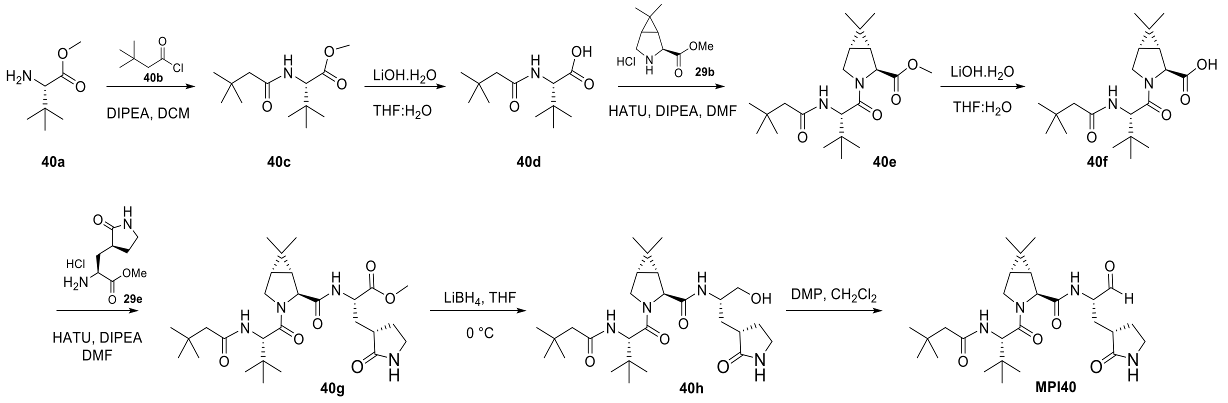


**Scheme 12**: The synthesis of compound **MPI40**

**(S)-methyl 2-(3,3-dimethylbutanamido)-3,3-dimethylbutanoate (40c).** To a stirred solution of **40a** (1.0 g, 5.52 mmol) and DIPEA (2.33 mL, 16.57 mmol) in dry $CH_2Cl_2$ (20 mL) was added **40b** (0.76 mL, 6.07 mmol) at 0 °C. After 10 h at rt, the reaction mixture was evaporated in vacuo. Purification by silica gel chromatography (Hexanes/EtOAc = 7:3). 150 mg of compound **40c** isolated. Yield 74%. $^{1}$H NMR (400 MHz, Chloroform-*d*) δ 5.87 (d, $J$ = 9.3 Hz, 1H), 4.46 (d, $J$ = 9.2 Hz, 1H), 3.71 (s, 3H), 2.11 (d, $J$ = 1.9 Hz, 2H), 1.04 (s, 9H), 0.97 (s, 9H). $^{13}$C NMR (101 MHz, $CDCl_3$) δ: 172.41, 171.41, 59.83, 51.75, 50.72, 34.53, 31.00, 29.84, 26.63.

**(S)-2-(3,3-dimethylbutanamido)-3,3-dimethylbutanoic acid (40d).** The compound **40c** (1.0 g, 4.11 mmol) was dissolved in THF/$H_2O$ (1:1, 30 mL). LiOH (432 mg, 10.25 mmol) was added at 0 °C. The mixture was stirred at room temperature overnight. Then THF was removed *on vacuum* and the aqueous layer was acidified with 1 M HCl and extracted with dichloromethane (3 x 10 mL). The organic layer was dried over anhydrous $Na_2SO_4$ and concentrated. Crude product **40d** directly used next step without further purification.

**(1R,2S,5S)-methyl 3-((S)-2-(3,3-dimethylbutanamido)-3,3-dimethylbutanoyl)-6,6-dimethyl-3-azabicyclo[3.1.0]hexane-2-carboxylate (40e).** The amino acid methyl ester hydrochloride **29b**

(0.716 g, 3.49 mmol) and the amino acid **40d** (0.8 g, 3.49 mmol) were dissolved in dry DMF (30 mL) and the reaction was cooled to 0 °C. HATU (1.72 g, 4.54 mmol) and DIPEA (2.43 mL, 13.96 mmol) were added, and the reaction mixture was allowed warm up to room temperature and stirred for 12 h. The mixture was then poured into water (50 mL) and extracted with ethyl acetate (4×20 mL). The organic layer was washed with aqueous hydrochloric acid 10% v/v (2×20 mL), saturated aqueous $NaHCO_3$ (2×20 mL), brine (2×20 mL) and dried over $Na_2SO_4$. The organic phase was evaporated to dryness and the crude material purified by silica gel column chromatography (15-50% EtOAc in n-hexane as the eluent) to afford **40e** white solid (0.75 g, 56%).$^1$H NMR (400 MHz, Chloroform-*d*) δ 4.67 (d, *J* = 9.8 Hz, 1H), 4.26 (s, 1H), 4.15 – 4.11 (m, 1H), 3.70 (s, 3H), 3.65 (d, *J* = 10.2 Hz, 1H), 2.09 (d, *J* = 1.2 Hz, 2H), 1.52 (dd, *J* = 7.6, 5.5 Hz, 1H), 1.40 (d, *J* = 7.6 Hz, 1H), 1.07 – 1.01 (m, 14H), 1.01 – 0.96 (m, 14H).

**(1R,2S,5S)-3-((S)-2-(3,3-dimethylbutanamido)-3,3-dimethylbutanoyl)-6,6-dimethyl-3-azabicyclo[3.1.0]hexane-2-carboxylic acid (40f).** The peptide **40e** (250 mg, 0.657 mmol) was dissolved in THF/$H_2O$ (1:1, 20 mL). LiOH (69 mg, 1.64 mmol) was added at 0 °C. The mixture was stirred at room temperature overnight. Then THF was removed *on vacuum* and the aqueous layer was acidified with 1 M HCl and extracted with dichloromethane (3 x 10 mL). The organic layer was dried over anhydrous $Na_2SO_4$ and concentrated to yield **40f** as white solid (200 mg).$^1$H NMR (400 MHz, Chloroform-*d*) δ 6.18 (d, *J* = 9.4 Hz, 1H), 4.66 (d, *J* = 9.3 Hz, 1H), 4.31 (s, 1H), 4.16 – 4.08 (m, 1H), 3.67 (d, *J* = 10.4 Hz, 1H), 2.09 (dd, *J* = 6.3, 4.0 Hz, 2H), 1.64 – 1.50 (m, 2H), 1.07 (s, 3H), 1.07 – 0.96 (m, 21H).

**(S)-methyl 2-((1R,2S,5S)-3-((S)-2-(3,3-dimethylbutanamido)-3,3-dimethylbutanoyl)-6,6-dimethyl-3-azabicyclo[3.1.0]hexane-2-carboxamido)-3-((S)-2-oxopyrrolidin-3-yl)propanoate (40g).** The amino acid methyl ester hydrochloride 2**9e** (0.123 g, 0.592 mmol) and the acid **40f** (0.18 g, 0.492 mmol) were dissolved in dry DMF (10 mL) and the reaction was cooled to 0 °C. HATU (229 mg, 0.639 mmol) and DIPEA (0.32 mL, 1.96 mmol) were added, and the reaction mixture was allowed warm up to room temperature and stirred for 12 h. The mixture was then poured into water (50 mL) and extracted with ethyl acetate (4×20 mL). The organic layer was washed with aqueous hydrochloric acid 10% v/v (2×20 mL), saturated aqueous $NaHCO_3$ (2×20 mL), brine (2×20 mL) and dried over $Na_2SO_4$. The organic phase was evaporated to dryness and the crude material purified by silica gel column chromatography (0-10% MeOH in $CH_2Cl_2$ as the eluent) to afford **40g** white gummy solid (200 mg, 76%).$^{1}$H NMR (400 MHz, Chloroform-*d*) δ 6.21 (d, *J* = 9.0 Hz, 1H), 6.13 (s, 1H), 4.64 – 4.54 (m, 2H), 4.26 (s, 1H), 4.17 (dd, *J* = 10.5, 5.1 Hz, 1H), 3.75 (d, *J* = 4.3 Hz, 1H), 3.72 (s, 3H), 3.67 (d, *J* = 10.4 Hz, 1H), 3.42 – 3.28 (m, 2H), 2.50 – 2.40 (m, 2H), 2.24 – 2.14 (m, 1H), 2.10 (d, *J* = 5.6 Hz, 2H), 1.96 – 1.81 (m, 2H), 1.57 – 1.49 (m, 2H), 1.06 (s, 3H), 1.04 (s, 9H), 1.02 (s, 9H), 0.98 (s, 3H).

**(1R,2S,5S)-3-((S)-2-(3,3-dimethylbutanamido)-3,3-dimethylbutanoyl)-N-((S)-1-hydroxy-3-((S)-2-oxopyrrolidin-3-yl)propan-2-yl)-6,6-dimethyl-3-azabicyclo[3.1.0]hexane-2-carboxamide (40h).** To a stirred solution of compound **40g** (200 mg, 0.376 mmol) in THF (8 mL) was added $LiBH_4$ (2.0 M in THF, 0.56 mL, 1.13 mmol) in several portions at 0 °C under a nitrogen atmosphere. The reaction mixture was stirred at 0 °C for 1 h, then allowed to warm up to room temperature, and stirred for an additional 2 h. The reaction was quenched by the drop wise addition of 1.0 M HCl (aq) (1.2 mL) with cooling in an ice bath. The solution was diluted with ethyl acetate and $H_2O$. The phases were separated, and the aqueous layer was extracted with ethyl acetate (3×15 mL). The organic phases were combined, dried over $MgSO_4$, filtered, and concentrated on a rotavapor to give a oily residue. Column chromatographic purification of the residue (6% MeOH in $CH_2Cl_2$ as the eluent) afforded **40h** as white solid (120 mg, 63%). $^{1}$H NMR (400 MHz, Chloroform-*d*) δ 6.75 (d, *J* = 8.6 Hz, 1H), 6.18 (d, *J* = 6.5 Hz, 1H), 6.02 (s, 1H), 4.35 (dd, *J* = 10.5, 5.3 Hz, 1H), 4.23 (s, 1H), 4.15 (d, *J* = 6.6 Hz, 1H), 4.06 (d, *J* = 12.4 Hz, 1H), 3.68 – 3.58 (m, 2H), 3.34 – 3.22 (m, 2H), 2.44 (dddd, *J* = 27.3, 13.1, 10.4, 6.6 Hz, 2H), 2.19 – 2.04 (m, 3H), 1.83 (dq, *J* = 11.7, 8.7 Hz, 1H), 1.56 – 1.42 (m, 3H), 1.07 – 0.99 (m, 21H), 0.96 (s, 3H). $^{13}$C NMR (101 MHz, $CDCl_3$) δ 180.44, 173.62, 171.77, 171.06, 64.70, 62.82, 58.97, 53.44, 50.18, 50.03, 48.51, 40.38, 38.21, 33.98, 32.60, 32.25, 31.07, 29.87, 28.37, 27.33, 26.79, 26.03, 19.82, 13.23.

**(1R,2S,5S)-3-((S)-2-(3,3-dimethylbutanamido)-3,3-dimethylbutanoyl)-6,6-dimethyl-N-((S)-1-oxo-3-((S)-2-oxopyrrolidin-3-yl)propan-2-yl)-3-azabicyclo[3.1.0]hexane-2-carboxamide (MPI40).** To a solution of **40h** (100 mg, 0.197 mmol) in $CH_2Cl_2$ (10 mL) was added $NaHCO_3$ (68 mg, 4 equiv.,) and the Dess-Martin reagent (257 mg, 0.592 mmol, 3 equiv.,). The resulting mixture was stirred at rt for 2 h. Then the reaction was quenched with a saturated $NaHCO_3$ solution containing 10 % $Na_2S_2O_3$. The layers were separated. The organic layer was then washed with saturated brine solution, dried over anhydrous $Na_2SO_4$, and concentrated *on vacuum*. The residue was then purified with flash chromatography afford **MPI40** as white solid (70 mg, 70%). $^1$H NMR (400 MHz, Chloroform-*d*) δ 9.40 (s, 1H), 7.67 (d, *J* = 7.6 Hz, 1H), 6.37 – 6.11 (m, 2H), 4.46 – 4.39 (m, 1H), 4.33 (d, *J* = 7.6 Hz, 2H), 4.26 (ddd, *J* = 10.4, 4.0, 1.4 Hz, 1H), 3.66 (d, *J* = 10.4 Hz, 1H), 3.37 – 3.29 (m, 2H), 2.50 – 2.36 (m, 2H), 2.08 – 2.01 (m, 3H), 1.96 – 1.81 (m, 2H), 1.54 – 1.47 (m, 2H), 1.05 (s, 3H), 1.02 (s, 9H), 1.00 (s, 9H), 0.98 (s, 3H). $^{13}$C NMR (101 MHz, $CDCl_3$) δ 200.18, 179.91, 172.85, 172.18, 170.79, 61.96, 58.18, 56.99, 53.44, 50.24, 48.40, 40.43, 37.96, 34.41, 32.03, 31.00, 29.97, 29.87, 28.31, 27.31, 26.73, 26.14, 19.74, 13.19.

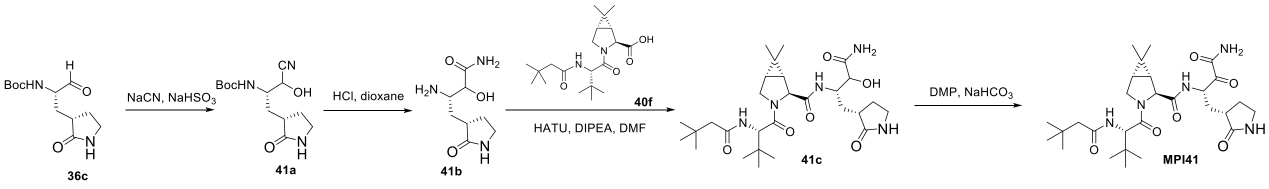


**Scheme 13**: The synthesis of compound **MPI41**

**Synthesis of tert-butyl ((2S)-1-cyano-1-hydroxy-3-((S)-2-oxopyrrolidin-3-yl)propan-2-yl)carbamate (41a).** To a solution of **36c** (300 mg, 1.17 mmol) in dichloromethane (25 mL) was added $NaHSO_3$ (610 mg, 5.8 mmol) slowly. The reaction was allowed to stir at RT for 30 min. Then NaCN (300 mg, 5.8 mmol), dissolved in 5 mL water was added to the reaction mixture slowly. The reaction mixture was stirred at RT for overnight. The mixture was washed with water, sat. NaCl, dried over $Na_2SO_4$ and concentrated. ESI-MS was used to confirm the formation of cyanohydrin intermediate **41a**, which was carried forward to the next step without further purification.

**Synthesis of (3S)-3-amino-2-hydroxy-4-((S)-2-oxopyrrolidin-3-yl)butanamide (41b).** To a solution of the cyanohydrin intermediate **41a** (250 mg, 0.88 mmol) in 1,4-dioxane (10 mL) was added dropwise a HCl solution in 1,4-dioxane (4 M, 10 mL). The resulting solution was stirred at room temperature for 3 h. Then residue was then concentrated *on vacuo* to afford the Boc-deprotected hydroxyamide intermediate. ESI-MS was used to confirm the formation of cyanohydrin intermediate **41b**, which was carried forward to the next step without further purification.

**Synthesis of (1R,2S,5S)-N-((2S)-4-amino-3-hydroxy-4-oxo-1-((S)-2-oxopyrrolidin-3-yl)butan-2-yl)-3-((S)-2-(3,3-dimethylbutanamido)-3,3-dimethylbutanoyl)-6,6-dimethyl-3-azabicyclo[3.1.0]hexane-2-carboxamide (41c).** To a solution of **40f** (0.29 mmol, 110 mg) and **41b** (0.32 mmol, 76 mg) in anhydrous DMF (6 mL) was added DIPEA (1.16 mmol, 150 mg, 0.21 mL) and was cooled to 0 °C. HATU (0.38 mmol, 145 mg) was added to the solution under 0 °C and then stirred at room temperature overnight. The reaction mixture was then diluted with ethyl acetate (50 mL) and washed with saturated $NaHCO_3$ solution (2×20 mL), 1 M HCl solution (2×20 mL), and saturated brine solution (2×20 mL) sequentially. The organic layer was dried over anhydrous $Na_2SO4$ and then concentrated *on vacuo*. The residue was purified by column chromatography (MeOH: DCM = 1:10 v/v) to afford the pure product **41c** (65 mg, 40.8%).

**Synthesis of (1R,2S,5S)-N-((S)-4-amino-3,4-dioxo-1-((S)-2-oxopyrrolidin-3-yl)butan-2-yl)-3-((S)-2-(3,3-dimethylbutanamido)-3,3-dimethylbutanoyl)-6,6-dimethyl-3-azabicyclo[3.1.0]hexane-2-carboxamide (MPI41).** To a solution of **41c** (65 mg, 0.12 mmol, 1.0 equiv.,) in anhydrous DCM (10 mL) was added Dess-Martin reagent (155 mg, 0.35 mmol, 3.0 equiv.,) slowly at 0 $^{o}$C. Then the reaction mixture was stirred at RT for 2 h. A solution of $NaHCO_3$ and $Na_2S_2O_3$ was added to quench the reaction. After 10 min, the mixture was washed with water, sat. NaCl, dried over $Na_2SO_4$ and concentrated. The residue was purified by column chromatography (MeOH: DCM = 1:10 v/v) to yield **MPI41** as a white solid (37 mg, yield 57 %). $^{1}$H NMR (400 MHz, Chloroform-*d*) δ 8.23 – 7.92 (m, 1H), 6.67 (s, 1H), 6.26 (s, 1H), 5.82 (s, 1H), 5.41 (s, 1H), 5.19 – 4.97 (m, 1H), 4.67 – 3.96 (m, 4H), 3.71 – 3.47 (m, 1H), 3.39 – 3.02 (m, 2H), 2.61 – 2.47 (m, 1H), 2.47 – 2.30 (m, 1H), 2.13 – 1.79 (m, 4H), 1.52 – 1.31 (m, 2H), 1.31 – 1.11 (m, 1H), 1.07 – 0.85 (m, 11H), 0.85 – 0.68 (m, 1H).

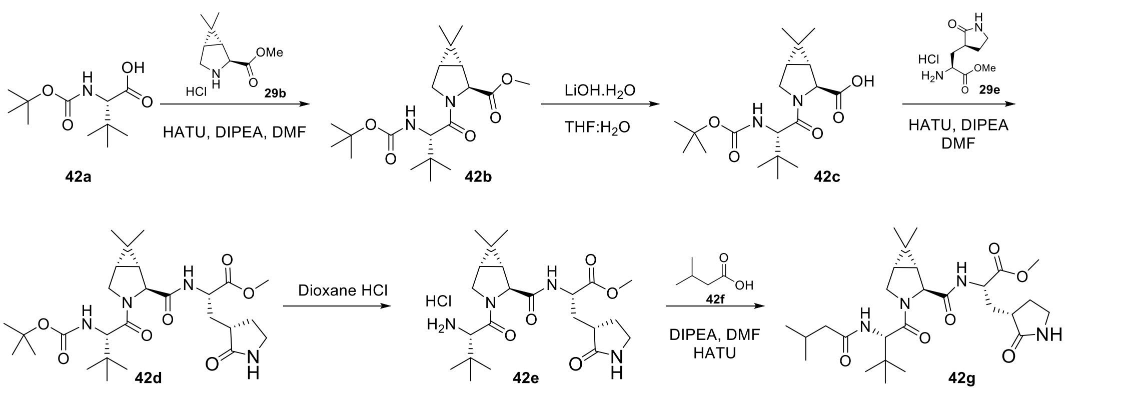


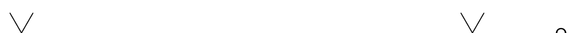

**Scheme 14**: The synthesis of compound **MPI42**

**(1R,2S,5S)-methyl-3-((S)-2-((tert-butoxycarbonyl)amino)-3,3-dimethylbutanoyl)-6,6-dimethyl-3-azabicyclo[3.1.0]hexane-2-carboxylate (42b).** To a solution of **42a** (3.9 mmol, 0.8 g) and **29b** (4.29 mmol, 0.992 g) in anhydrous DMF (20 mL) was added DIPEA (15.6 mmol, 2.72 mL) and was cooled to 0 °C. HATU (5.07 mmol, 1.78 g) was added to the solution under 0 °C and then stirred at room temperature overnight. The reaction mixture was then diluted with ethyl acetate (50 mL) and washed with saturated $NaHCO_3$ solution (2×20 mL), 1 M HCl solution (2×20 mL), and saturated brine solution (2×20 mL) sequentially. The organic layer was dried over anhydrous $Na_2SO_4$ and then concentrated *on vacuum*. The residue was then purified with flash chromatography (30-70% EtOAc in hexanes as the eluent) to afford **42b** as white solid (1.3 g, 87%). $^1$H NMR (400 MHz, Chloroform-*d*) δ 5.10 (d, *J* = 10.2 Hz, 1H), 4.46 (s, 1H), 4.20 (d, *J* = 10.3 Hz, 1H), 3.99 (d, *J* = 10.2 Hz, 1H), 3.89 – 3.83 (m, 1H), 3.74 (s, 3H), 1.48 – 1.42 (m, 2H), 1.39 (s, 9H), 1.02 (d, *J* = 6.1 Hz, 12H), 0.90 (s, 3H).$^{13}$C NMR (101 MHz, $CDCl_3$): δ 171.99, 171.07, 155.97, 79.58, 59.16, 58.56, 52.25, 47.69, 34.96, 30.35, 28.37, 28.23, 27.38, 26.26, 19.44, 12.39.

**(1R,2S,5S)-3-((S)-2-((tert-butoxycarbonyl)amino)-3,3-dimethylbutanoyl)-6,6-dimethyl-3-azabicyclo[3.1.0]hexane-2-carboxylic acid (42c).** The peptide **42b** (1.3 g, 3.4 mmol) was dissolved in THF/$H_2O$ (1:1, 30 mL). LiOH (331 mg, 8.5 mmol) was added at 0 °C. The mixture was stirred at room temperature overnight. Then THF was removed *on vacuum* and the aqueous layer was acidified with 1 M HCl and extracted with dichloromethane (3 x 30 mL). The organic layer was dried over anhydrous $Na_2SO_4$ and concentrated to yield **42c** as white solid (1.3 g, crude).

[1]H NMR (400 MHz, Chloroform-*d*) δ 5.19 (dd, *J* = 10.7, 4.9 Hz, 1H), 4.46 (s, 1H), 4.23 (d, *J* = 10.2 Hz, 1H), 4.04 (d, *J* = 10.4 Hz, 1H), 3.84 (dd, *J* = 10.4, 5.4 Hz, 1H), 1.70 – 1.63 (m, 1H), 1.53 – 1.44 (m, 1H), 1.39 (s, 9H), 1.05 (s, 3H), 1.00 (s, 9H), 0.89 (s, 3H).

**(S)-methyl 2-((1R,2S,5S)-3-((S)-2-((tert-butoxycarbonyl)amino)-3,3-dimethylbutanoyl)-6,6-dimethyl-3-azabicyclo[3.1.0]hexane-2-carboxamido)-3-((S)-2-oxopyrrolidin-3-yl)propanoate (42d).** To a solution of **42c** (3.53 mmol, 1.3 g) and **29e** (3.88 mmol, 840 mg) in anhydrous DMF (20 mL) was added DIPEA (14.1 mmol, 2.4 mL) and was cooled to 0 °C. HATU (4.23 mmol, 1.57 g) was added to the solution under 0 °C and then stirred at room temperature overnight. The reaction mixture was then diluted with ethyl acetate (50 mL) and washed with saturated $NaHCO_3$ solution (2×20 mL), 1 M HCl solution (2×20 mL), and saturated brine solution (2×20 mL) sequentially. The organic layer was dried over anhydrous $Na_2SO_4$ and then concentrated *on vacuum*. The residue was then purified with flash chromatography (0-10% MeOH in Dichloromethane as the eluent) to afford **42d** as white solid (1.51 g, 79%). [1]H NMR (400 MHz, Chloroform-*d*): δ 7.41 (t, *J* = 7.2 Hz, 1H), 6.21 (d, *J* = 53.6 Hz, 1H), 5.12 (d, *J* = 10.2 Hz, 1H), 4.59 (ddd, *J* = 11.1, 7.5, 3.8 Hz, 1H), 4.31 (d, *J* = 1.7 Hz, 1H), 4.18 (dd, *J* = 10.3, 2.8 Hz, 1H), 4.01 – 3.85 (m, 2H), 3.72 (s, 3H), 3.41 – 3.20 (m, 2H), 2.50 (m, 1H), 2.44 – 2.37 (m, 1H), 2.23 – 2.11 (m, 1H), 1.90 – 1.76 (m, 2H), 1.55 – 1.48 (m, 2H), 1.37 (s, 9H), 1.02 (s, 3H), 0.97 (s, 9H), 0.87 (s, 3H).[13]C NMR (101 MHz, $CDCl_3$): δ 179.72, 172.25, 171.41, 155.94, 79.60, 60.73, 52.45, 51.07, 48.29, 40.38, 38.62, 37.99, 34.92, 33.47, 30.36, 28.22, 27.73, 26.35, 26.24, 19.20, 12.53.

**(S)-methyl 2-((1R,2S,5S)-3-((S)-2-amino-3,3-dimethylbutanoyl)-6,6-dimethyl-3-azabicyclo[3.1.0]hexane-2-carboxamido)-3-((S)-2-oxopyrrolidin-3-yl)propanoate (42e).** To a solution of **42d** (1.0 g, 1.86 mmol) in 1,4-dioxane (10 mL) was added drop wise a HCl solution in 1,4-dioxane (4 M, 0.6 mL). The resulting solution was stirred at room temperature for 2 h. Then residue was then concentrated *on vacuum* to afford **42e** as light-yellow hygroscopic solid (700 mg).[1]H NMR (400 MHz, Deuterium Oxide) δ 4.46 (dd, *J* = 11.4, 4.0 Hz, 1H), 4.35 (s, 1H), 3.97 (s, 1H), 3.93 (dd, *J* = 12.0, 7.0 Hz, 1H), 3.73 – 3.64 (m, 5H), 3.37 – 3.21 (m, 2H), 2.63 – 2.50 (m, 1H), 2.28 – 2.18 (m, 1H), 2.14 – 2.04 (m, 1H), 1.94 – 1.76 (m, 2H), 1.62 (t, *J* = 6.6 Hz, 1H), 1.46 (d, *J* = 7.6 Hz, 1H), 1.32 – 1.24 (m, 2H), 1.02 (s, 9H), 0.99 (s, 3H), 0.88 (s, 3H).

**(S)-methyl 2-((1R,2S,5S)-3-((S)-3,3-dimethyl-2-(3-methyl butanamido) butanoyl)-6,6-dimethyl-3-azabicyclo[3.1.0]hexane-2-carboxamido)-3-((S)-2-oxopyrrolidin-3-yl)propanoate (42g).** To a solution of **42e** (0.48 mmol, 0.230 g) and **42f** (0.58 mmol, 0.06 g) in anhydrous DMF (10 mL) was added DIPEA (1.94 mmol, 0.35 mL) and was cooled to 0 °C. HATU (0.63 mmol, 0.24 g) was added to the solution under 0 °C and then stirred at room temperature overnight. The reaction mixture was then diluted with ethyl acetate (50 mL) and washed with saturated $NaHCO_3$ solution (2×20 mL), 1 M HCl solution (2×20 mL), and saturated brine solution (2×20 mL) sequentially. The organic layer was dried over anhydrous $Na_2SO_4$ and then concentrated *on vacuo*. The residue was then purified with flash chromatography (0-10% MeOH in $CH_2Cl_2$ as the eluent) to afford **42g** as white solid (200 mg, 79%).$^1$H NMR (400 MHz, Chloroform-*d*) δ 7.40 (d, *J* = 7.7 Hz, 1H), 6.06 (s, 1H), 5.71 (s, 1H), 4.62 (dd, *J* = 8.6, 6.2 Hz, 2H), 4.33 (s, 1H), 3.98 – 3.95 (m, 2H), 3.76 (s, 3H), 3.75 – 3.71 (m, 1H), 3.42 – 3.30 (m, 2H), 2.62 – 2.40 (m, 2H), 2.18 (ddd, *J* = 14.2, 11.0, 5.1 Hz, 1H), 2.06 (d, *J* = 3.2 Hz, 2H), 1.96 – 1.81 (m, 2H), 1.59 – 1.53 (m, 2H), 1.05 (s, 3H), 1.02 (s, 9H), 0.93 (dd, *J* = 8.7, 5.0 Hz, 6H), 0.88 (s, 3H).

**(1R,2S,5S)-3-((S)-3,3-dimethyl-2-(3-methylbutanamido)butanoyl)-N-((S)-1-hydroxy-3-((S)-2-oxopyrrolidin-3-yl)propan-2-yl)-6,6-dimethyl-3-azabicyclo[3.1.0]hexane-2-carboxamide (42h).** To a stirred solution of compound **42g** (200 mg, 0.384 mmol) in THF (8 mL) was added $LiBH_4$ (2.0 M in THF, 1.0 mL, 1.92 mmol) in several portions at 0 °C under a nitrogen atmosphere. The reaction mixture was stirred at 0 °C for 1 h, then allowed to warm up to room temperature, and stirred for an additional 2 h. The reaction was quenched by the drop wise addition of 1.0 M HCl (aq) (1.2 mL) with cooling in an ice bath. The solution was diluted with ethyl acetate and $H_2O$. The phases were separated, and the aqueous layer was extracted with ethyl acetate (3×15 mL). The organic phases were combined, dried over $MgSO_4$, filtered, and concentrated on a rotavapor to give a yellow oily residue. Column chromatographic purification of the residue (6% MeOH in $CH_2Cl_2$ as the eluent) afforded **42h** as white solid (110 mg, 58%).$^1$H NMR (400 MHz, Chloroform-*d*) δ 7.31 (d, *J* = 8.1 Hz, 1H), 6.50 (s, 1H), 6.33 (d, *J* = 9.6 Hz, 1H), 4.58 (d, *J* = 9.6 Hz, 1H), 4.24 (s, 1H), 4.07 – 3.98 (m, 1H), 3.93 (d, *J* = 3.1 Hz, 2H), 3.62 – 3.57 (m, 2H), 3.31 – 3.23 (m, 2H), 2.50 (qd, *J* = 8.6, 4.9 Hz, 1H), 2.38 (dt, *J* = 9.0, 2.6 Hz, 1H), 2.07 – 1.98 (m, 4H), 1.82 – 1.73 (m, 1H), 1.59 – 1.53 (m, 1H), 1.51 – 1.46 (m, 1H), 1.43 (d, *J* = 7.7 Hz, 1H), 1.00 (s, 4H), 0.97 (s, 9H), 0.91 – 0.85 (m, 6H), 0.83 (s, 3H). $^{13}$C NMR (101 MHz, $CDCl_3$) δ 181.03, 172.35,

171.76, 170.94, 65.34, 61.19, 56.92, 49.95, 48.47, 45.70, 40.49, 38.06, 35.20, 32.43, 30.85, 28.49, 27.77, 26.52, 26.21, 22.39, 22.34, 19.17, 12.62.

**(1R,2S,5S)-3-((S)-3,3-dimethyl-2-(3-methylbutanamido)butanoyl)-6,6-dimethyl-N-((S)-1-oxo-3-((S)-2-oxopyrrolidin-3-yl)propan-2-yl)-3-azabicyclo[3.1.0]hexane-2-carboxamide (MPI42).** To a solution of **42h** (100 mg, 0.203 mmol) in $CH_2Cl_2$ (6 mL) was added $NaHCO_3$ (65 mg, 4 equiv.,) and the Dess-Martin reagent (264 mg, 0.609 mmol, 3 equiv.,). The resulting mixture was stirred at rt for 12 h. Then the reaction was quenched with a saturated $NaHCO_3$ solution containing 10 % $Na_2S_2O_3$. The layers were separated. The organic layer was then washed with saturated brine solution, dried over anhydrous $Na_2SO_4$, and concentrated *on vacuum*. The residue was then purified with flash chromatography afford **MPI42** as white solid (75 mg, 75%). $^1$H NMR (400 MHz, Chloroform-*d*) δ 9.47 (s, 1H), 7.83 (d, *J* = 6.8 Hz, 1H), 6.07 (td, *J* = 17.2, 16.6, 5.9 Hz, 2H), 4.55 (dd, *J* = 9.7, 1.4 Hz, 1H), 4.40 (ddd, *J* = 8.9, 6.8, 5.2 Hz, 1H), 4.29 (s, 1H), 3.90 (d, *J* = 2.8 Hz, 2H), 3.35 – 3.21 (m, 2H), 2.57 – 2.42 (m, 1H), 2.39 – 2.30 (m, 1H), 2.01 – 1.96 (m, 3H), 1.94 – 1.89 (m, 2H), 1.77 (ddt, *J* = 17.2, 8.3, 4.9 Hz, 1H), 1.51 – 1.42 (m, 2H), 0.98 (s, 3H), 0.93 (s, 9H), 0.87 – 0.82 (m, 6H), 0.81 (s, 3H). $^{13}$C NMR (101 MHz, $CDCl_3$) δ 199.47, 179.95, 172.23, 172.07, 170.84, 60.82, 57.46, 56.85, 48.37, 45.86, 40.45, 37.64, 35.28, 30.73, 30.02, 28.60, 27.80, 26.47, 26.23, 22.40, 22.34, 19.27, 12.59.

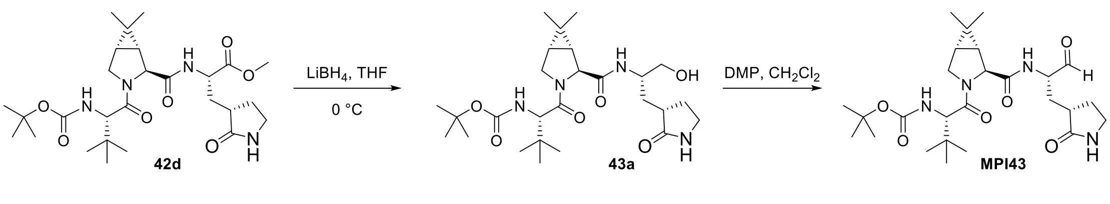


**Scheme 15**: The synthesis of compound **MPI43**

**tert-butyl ((S)-1-((1R,2S,5S)-2-(((S)-1-hydroxy-3-((S)-2-oxopyrrolidin-3-yl)propan-2-yl)carbamoyl)-6,6-dimethyl-3-azabicyclo[3.1.0]hexan-3-yl)-3,3-dimethyl-1-oxobutan-2-yl)carbamate (43a).** To a stirred solution of compound 4**2d** (1.0 g, 1.86 mmol) in THF (30 mL) was added $LiBH_4$ (2.0 M in THF, 4.5 mL, 9.32 mmol) in several portions at 0 °C under a nitrogen atmosphere. The reaction mixture was stirred at 0 °C for 1 h, then allowed to warm up to room temperature, and stirred for an additional 2 h. The reaction was quenched by the drop wise addition of 1.0 M HCl (aq) (1.2 mL) with cooling in an ice bath. The solution was diluted with ethyl acetate and $H_2O$. The phases were separated, and the aqueous layer was extracted with ethyl acetate (3×50 mL). The organic phases were combined, dried over $MgSO_4$, filtered, and concentrated on a rotavapor to give a yellow oily residue. Column chromatographic purification of the residue (6% MeOH in $CH_2Cl_2$ as the eluent) afforded **43a** a white solid (810 mg, 85%). $^1$H NMR (400 MHz, Chloroform-*d*): δ 7.34 (d, *J* = 8.0 Hz, 1H), 6.38 (s, 1H), 5.18 (d, *J* = 10.2 Hz, 1H), 4.27 (s, 1H), 4.19 (d, *J* = 10.2 Hz, 1H), 4.07 – 3.88 (m, 3H), 3.79 (t, *J* = 6.2 Hz, 1H), 3.66 – 3.56 (m, 2H), 3.44 (d, *J* = 3.3 Hz, 1H), 3.31 – 3.25 (m, 2H), 2.58 – 2.47 (m, 2H), 2.38 (dddd, *J* = 12.1, 8.6, 5.7, 3.2 Hz, 1H), 2.06 – 1.96 (m, 1H), 1.83 – 1.72 (m, 1H), 1.61 – 1.53 (m, 1H), 1.50 – 1.44 (m, 2H), 1.37 (s, 9H), 1.01 (s, 3H), 0.97 (s, 9H), 0.86 (s, 3H). $^{13}$C NMR (101 MHz, $CDCl_3$) δ 181.05, 171.92, 171.43, 155.94, 79.63, 65.42, 61.15, 58.70, 50.63, 50.17, 48.35, 40.48, 37.97, 34.93, 32.35, 30.83, 28.64, 28.23, 27.75, 26.38, 26.24, 19.20, 12.55.

**tert-butyl ((S)-1-((1R,2S,5S)-6,6-dimethyl-2-(((S)-1-oxo-3-((S)-2-oxopyrrolidin-3-yl)propan-2-yl)carbamoyl)-3-azabicyclo[3.1.0]hexan-3-yl)-3,3-dimethyl-1-oxobutan-2-yl)carbamate (MPI43).** To a solution of **43a** (110 mg, 0.216 mmol) in $CH_2Cl_2$ (6 mL) was added $NaHCO_3$ (73 mg, 4 eq) and the Dess-Martin reagent (275 mg, 0.649 mmol, 3 eq). The resulting mixture was stirred at rt for 12 h. Then the reaction was quenched with a saturated $NaHCO_3$ solution containing 10 % $Na_2S_2O_3$. The layers were separated. The organic layer was then washed with saturated brine solution, dried over anhydrous $Na_2SO_4$ and concentrated *on vacuum*. The residue was then purified with flash chromatography afford **MPI43** as white solid (90 mg, 82%). $^1$H NMR (400 MHz, Chloroform-*d*) δ 9.47 (s, 1H), 7.84 (t, *J* = 7.1 Hz, 1H), 6.03 (s, 1H), 5.26 – 4.99 (m, 1H), 4.39 (ddd, *J* = 9.0, 6.8, 5.0 Hz, 1H), 4.32 (s, 1H), 4.15 (d, *J* = 10.3 Hz, 1H), 3.97 – 3.81 (m, 2H), 3.28 (dtd, *J* = 11.2, 9.3, 8.9, 2.4 Hz, 2H), 2.58 – 2.42 (m, 1H), 2.38 – 2.28 (m, 1H), 1.91 (ddd, *J* = 15.5, 8.8, 4.2 Hz, 2H), 1.84-1.71 (m, 1H), 1.46 (d, *J* = 2.2 Hz, 2H), 1.33 (s, 9H), 0.98 (s, 3H), 0.92 (s, 9H), 0.83 (s, 3H).$^{13}$C NMR (101 MHz, $CDCl_3$) δ 199.53, 180.05, 172.16, 171.37, 155.96, 79.65, 60.78, 60.40, 58.66, 57.43, 48.26, 40.48, 37.55, 34.95, 30.68, 29.99, 28.57, 28.44, 28.24, 27.81, 26.35, 26.26, 19.27, 12.54.

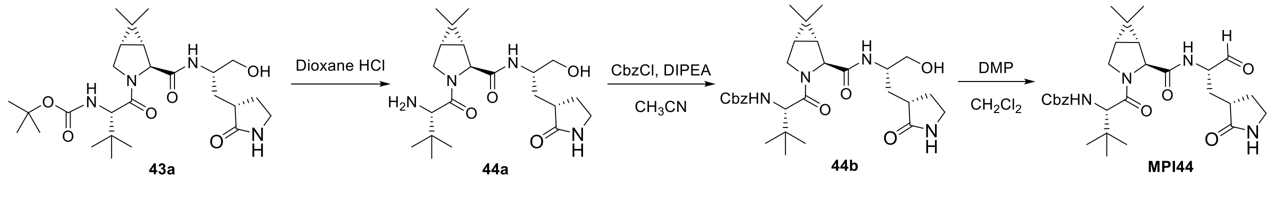

**Scheme 16**: The synthesis of compound **MPI44**

**(1R,2S,5S)-3-((S)-2-amino-3,3-dimethylbutanoyl)-N-((S)-1-hydroxy-3-((S)-2-oxopyrrolidin-3-yl)propan-2-yl)-6,6-dimethyl-3-azabicyclo[3.1.0]hexane-2-carboxamide hydrochloride(44a).** To a solution of **43a** (250 mg, 0.492 mmol) in 1,4-dioxane (10 mL) was added drop wise a HCl solution in 1,4-dioxane (4 M, 1.2 mL). The resulting solution was stirred at room temperature for 2 h. Then residue was then concentrated *on vacuum* to afford **44a** as light-yellow hygroscopic solid (150 mg). $^1$H NMR (400 MHz, Deuterium Oxide) δ 4.28 (s, 1H), 3.96 (s, 1H), 3.95 – 3.88 (m, 1H), 3.70 – 3.63 (m, 1H), 3.53 (ddq, *J* = 17.8, 11.5, 6.2 Hz, 2H), 3.30 (td, *J* = 9.9, 2.7 Hz, 1H), 3.22 (t, *J* = 8.6 Hz, 1H), 2.48 (p, *J* = 8.5 Hz, 1H), 2.27 – 2.14 (m, 1H), 1.91 – 1.72 (m, 2H), 1.61 (t, *J* = 6.7 Hz, 1H), 1.53 – 1.38 (m, 2H), 1.01 (s, 9H), 0.97 (s, 3H), 0.86 (s, 3H).$^{13}$C NMR (101 MHz, $D_2O$): δ 182.63, 173.18, 167.07, 66.55, 63.80, 61.69, 58.98, 49.27, 48.72, 40.61, 38.25, 34.41, 31.68, 30.70, 27.77, 26.78, 25.37, 25.00, 19.11, 11.98.

**benzyl ((S)-1-((1R,2S,5S)-2-(((S)-1-hydroxy-3-((S)-2-oxopyrrolidin-3-yl)propan-2-yl)carbamoyl)-6,6-dimethyl-3-azabicyclo[3.1.0]hexan-3-yl)-3,3-dimethyl-1-oxobutan-2-yl)carbamate (44b).** To a solution of **44a** (100 mg, 0.225 mmol) was taken up in $CH_3CN$ (10 mL). The solution was cooled to 0 °C with an ice bath and benzyl chloroformate (46 mg, 0.27 mmol) was added followed by a slow addition of diisopropylethylamine (0.118 mL, 0.675 mmol). The reaction was allowed to warm to rt and stirred. The reaction was monitored and determined to be complete at 1h by TLC. Saturated aqueous $NaHCO_3$ was added. The layers were separated, and the aqueous layer was washed with $CH_2Cl_2$ (20 mL). The organics were combined and dried over $Na_2SO_4$. The solids were filtered, and solvent removed under reduced pressure. Compound **44b** was isolated by silica gel chromatography as white solid, yield 50 mg, 41 %.$^1$H NMR (400 MHz, Chloroform-*d*) δ 7.39 (d, *J* = 8.0 Hz, 1H), 7.34-7.24 (m, 5H), 6.30 (s, 1H), 5.64 (d, *J* = 9.9 Hz, 1H), 5.09 – 4.96 (m, 2H), 4.22 (d, *J* = 9.6 Hz, 2H), 4.03 – 3.89 (m, 2H), 3.83 (d, *J* = 10.2 Hz, 1H), 3.64 – 3.50 (m, 2H), 3.19 (dd, *J* = 9.2, 4.5 Hz, 2H), 2.49 (qd, *J* = 8.4, 4.9 Hz, 1H), 2.32 (ddt, *J* = 12.8, 8.8, 4.5 Hz, 1H), 2.01 (td, *J* = 10.4, 5.4 Hz, 1H), 1.69 (td, *J* = 9.8, 2.9 Hz, 1H), 1.49 (ddd, *J* = 18.9, 7.9, 4.6 Hz, 2H), 1.41 (d, *J* = 7.7 Hz, 1H), 0.98 (s, 3H), 0.93 (s, 9H), 0.81 (s, 3H).$^{13}$C NMR (101 MHz, $CDCl_3$) δ 181.11, 171.88, 170.76, 156.42, 136.42, 128.49, 128.11, 127.99, 66.86, 65.50,

61.21, 59.36, 50.06, 48.40, 40.45, 37.93, 35.31, 32.30, 30.99, 28.57, 27.84, 26.60, 26.42, 26.25, 19.21, 12.70.

**benzyl ((S)-1-((1R,2S,5S)-6,6-dimethyl-2-(((S)-1-oxo-3-((S)-2-oxopyrrolidin-3-yl)propan-2-yl)carbamoyl)-3-azabicyclo[3.1.0]hexan-3-yl)-3,3-dimethyl-1-oxobutan-2-yl)carbamate (MPI44).** To a solution of **44b** (50 mg, 0.092 mmol) in $CH_2Cl_2$ (6 mL) was added $NaHCO_3$ (32 mg, 4 equiv.,) and the Dess-Martin reagent (120 mg, 0.27 mmol, 3 equiv.,). The resulting mixture was stirred at rt for 3 h. Then the reaction was quenched with a saturated $NaHCO_3$ solution containing 10 % $Na_2S_2O_3$. The layers were separated. The organic layer was then washed with saturated brine solution, dried over anhydrous $Na_2SO_4$, and concentrated *on vacuum*. The residue was then purified with flash chromatography afford **MPI44** as white solid (40 mg, 80%). $^1$H NMR (400 MHz, Chloroform-*d*) δ 9.52 (s, 1H), 7.99 (d, *J* = 7.4 Hz, 1H), 7.46 – 7.17 (m, 5H), 6.30 (s, 1H), 5.66 (s, 1H), 5.15 – 5.00 (m, 2H), 4.44 (ddd, *J* = 9.7, 7.1, 4.7 Hz, 1H), 4.35 (s, 1H), 4.26 (d, *J* = 9.9 Hz, 1H), 4.03 – 3.84 (m, 2H), 3.35 – 3.21 (m, 2H), 2.62 – 2.51 (m, 1H), 2.40 – 2.28 (m, 1H), 2.07 – 1.98 (m, 1H), 1.96 – 1.86 (m, 1H), 1.84 – 1.71 (m, 1H), 1.59 – 1.47 (m, 2H), 1.03 (s, 3H), 0.96 (s, 9H), 0.86 (s, 3H). $^{13}$C NMR (101 MHz, $CDCl_3$) δ 199.66, 180.22, 172.18, 170.71, 156.42, 136.37, 128.52, 128.47, 128.14, 128.00, 66.91, 60.91, 59.29, 57.27, 53.45, 48.33, 40.47, 37.52, 35.32, 30.90, 27.90, 26.36, 26.24, 19.30, 12.67.

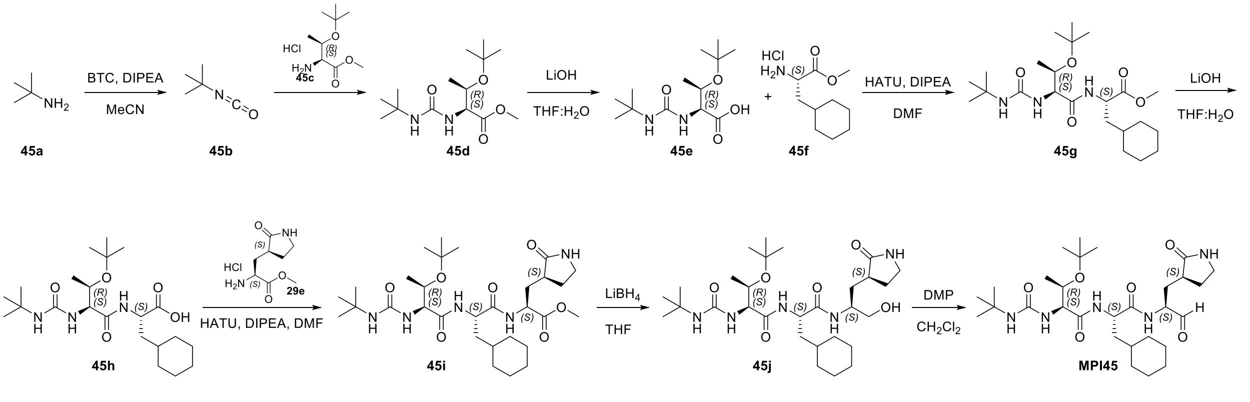


**Scheme 17**: The synthesis of compound **MPI45**

**Methyl O-(tert-butyl)-N-(tert-butylcarbamoyl)-L-threoninate (45d)**. To a solution of tert-butylamine (710µL, 6.74 mmol, 1.0 eq) in MeCN was added DIPEA (3.6 mL, 20.2 mmol, 3.0 eq) at 0 °C. The reaction mixture was stirred for 20 minutes. BTC (1.0 g, 3.37 mmol 0.5 eq) was added dropwise, then warmed to RT for 0.5 h. H-Thr(tBu)-OMe hydrochloride **45c** (2.0g, 8.76mmol 1.3

eq) was added. The reaction mixture was then allowed to stir at RT overnight. The solvent was removed in vacuo and the resulted residue was purified by column chromatography (hexane: EA= 1:1 v/v) to yield 4**5d** (1.0 g, yield 51%). $^1$H NMR (400 MHz, Methanol-$d_4$) δ 4.2 – 4.0 (m, 2H), 3.6 (s, 3H), 1.2 (s, 9H), 1.1 (d, $J$ = 6.1 Hz, 3H), 1.0 (s, 9H). $^{13}$C NMR (100 MHz, Methanol-$d_4$) δ 174.2, 160.3, 74.8, 68.9, 59.8, 52.4, 50.8, 29.8, 28.8, 21.3.

**O-(tert-butyl)-N-(tert-butylcarbamoyl)-L-threonine (45e). 45e** was prepared as a white solid following a similar procedure to **29d** (yield 85%), the residue was used in the next step without further purification.

**Methyl (S)-2-((2S,3R)-3-(tert-butoxy)-2-(3-(tert-butyl)ureido)butanamido)-3-cyclohexylpropanoate (45g). 45g** was prepared as a light yellow oil following a similar procedure to **29c**. (yield 72%).

**(S)-2-((2S,3R)-3-(tert-butoxy)-2-(3-(tert-butyl)ureido)butanamido)-3-cyclohexylpropanoic acid (45h). 45h** was prepared as a white solid following a similar procedure to **29d** (yield 75%), the residue was used in the next step without further purification.

**Methyl (6S,9S,12S)-6-((R)-1-(tert-butoxy)ethyl)-9-(cyclohexylmethyl)-2,2-dimethyl-4,7,10-trioxo-12-(((S)-2-oxopyrrolidin-3-yl)methyl)-3,5,8,11-tetraazatridecan-13-oate (45i). 45i** was prepared as a white solid following a similar procedure to **29f** (yield 40%). $^1$H NMR (400 MHz, Methanol-$d_4$) δ 4.6 – 4.4 (m, 2H), 4.2 – 4.1 (m, 2H), 3.8 (s, 3H), 3.4 – 3.3 (m, 2H), 2.5 (qd, $J$ = 10.3, 4.0 Hz, 1H), 2.4 – 2.3 (m, 1H), 2.2 (ddd, $J$ = 14.0, 11.5, 4.0 Hz, 1H), 1.9 – 1.8 (m, 4H), 1.8 – 1.6 (m, 4H), 1.6 (ddd, $J$ = 14.0, 8.3, 5.6 Hz, 1H), 1.5 (td, $J$ = 9.7, 9.0, 5.5 Hz, 1H), 1.3 (s, 11H), 1.2 (s, 10H), 1.1 (d, $J$ = 6.1 Hz, 3H), 1.1 – 0.9 (m, 2H). $^{13}$C NMR (100 MHz, Methanol-$d_4$) δ 181.5, 174.6, 173.5, 173.4, 159.7, 75.3, 68.9, 59.9, 52.8, 52.5, 51.7, 50.8, 41.4, 41.3, 39.4, 35.1, 34.6, 33.9, 33.8, 29.7, 28.9, 28.6, 27.5, 27.3, 27.2, 19.9.

**(2S,3R)-3-(tert-butoxy)-2-(3-(tert-butyl)ureido)-N-((S)-3-cyclohexyl-1-(((S)-1-hydroxy-3-((S)-2-oxopyrrolidin-3-yl)propan-2-yl)amino)-1-oxopropan-2-yl)butanamide (45j). 45j** was prepared as a white solid following a similar procedure to **37e** (yield 76%). $^1$H NMR (400 MHz,

Chloroform-*d*) δ 8.1 (d, *J* = 9.3 Hz, 1H), 7.8 (d, *J* = 7.2 Hz, 1H), 6.3 – 6.2 (m, 1H), 6.0 (s, 1H), 5.4 (s, 1H), 4.3 – 4.0 (m, 4H), 3.8 – 3.5 (m, 2H), 3.3 – 3.1 (m, 2H), 2.6 – 2.3 (m, 3H), 1.8 (dq, *J* = 15.2, 7.7 Hz, 1H), 1.7 – 1.5 (m, 6H), 1.4 – 1.3 (m, 2H), 1.3 (d, *J* = 5.0 Hz, 23H), 1.0 (d, *J* = 6.4 Hz, 3H), 0.9 – 0.8 (m, 2H). $^{13}$C NMR (100 MHz, Chloroform-*d*) δ 181.8, 172.4, 171.9, 157.5, 75.3, 67.3, 66.2, 57.4, 52.0, 49.7, 48.7, 40.4, 38.6, 38.3, 34.6, 34.1, 33.0, 32.8, 29.5, 28.3, 27.7, 26.3, 26.1, 26.1, 16.3.

**(2S,3R)-3-(tert-butoxy)-2-(3-(tert-butyl)ureido)-N-((S)-3-cyclohexyl-1-oxo-1-(((S)-1-oxo-3-((S)-2-oxopyrrolidin-3-yl)propan-2-yl)amino)propan-2-yl)butanamide (MPI45). MPI45** was prepared as a white solid following a similar procedure to **MPI29** (yield 67%). $^{1}$H NMR (400 MHz, Chloroform-*d*) δ 9.5 (s, 1H), 8.1 (d, *J* = 6.4 Hz, 1H), 7.6 (d, *J* = 8.0 Hz, 1H), 6.9 (s, 1H), 5.7 (d, *J* = 6.0 Hz, 1H), 4.5 (td, *J* = 8.5, 5.4 Hz, 1H), 4.3 (ddd, *J* = 10.8, 7.0, 4.4 Hz, 1H), 4.2 (dd, *J* = 6.0, 3.7 Hz, 1H), 4.1 – 4.0 (m, 1H), 3.4 – 3.1 (m, 2H), 2.5 – 2.3 (m, 2H), 2.1 – 1.9 (m, 1H), 1.8 (ddd, *J* = 13.2, 8.1, 4.2 Hz, 1H), 1.8 – 1.5 (m, 8H), 1.5 (ddd, *J* = 14.1, 8.9, 5.6 Hz, 1H), 1.3 – 1.1 (m, 21H), 1.1 (d, *J* = 12.0 Hz, 1H), 1.0 (d, *J* = 6.3 Hz, 3H), 1.0 – 0.8 (m, 2H). $^{13}$C NMR (100 MHz, Chloroform-*d*) δ 199.9, 180.1, 173.0, 171.6, 157.3, 75.1, 67.6, 58.0, 57.2, 51.3, 50.1, 40.5, 39.8, 37.9, 34.1, 33.6, 32.6, 30.0, 29.5, 28.3, 28.2, 26.3, 26.2, 26.0, 17.6.

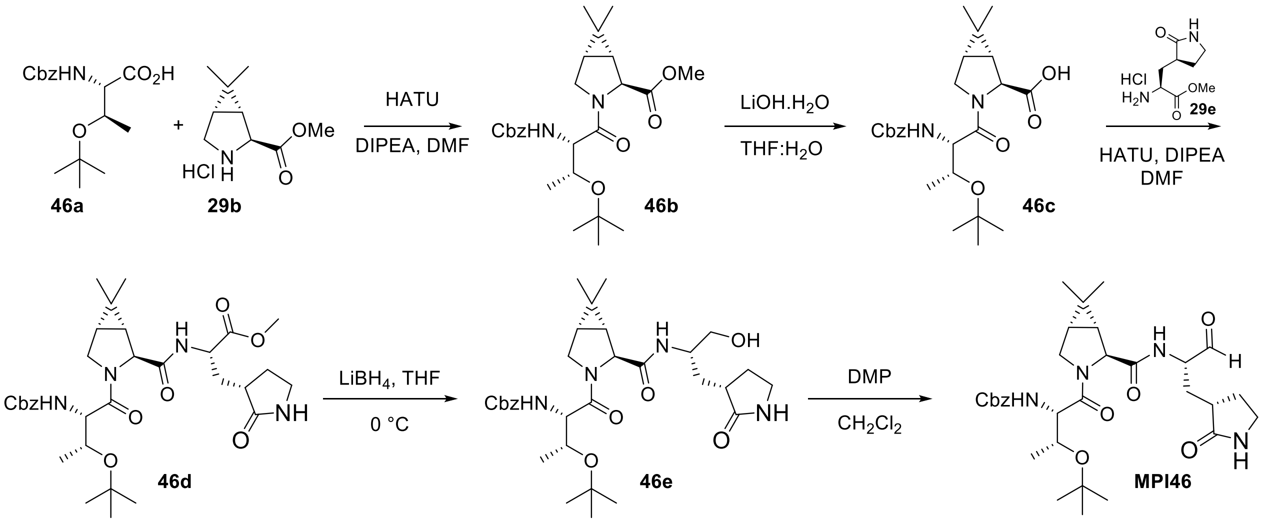


**Scheme S18**: The synthesis of compound **MPI46**

**(1R,2S,5S)-methyl 3-((2S,3R)-2-(((benzyloxy)carbonyl)amino)-3-(tert-butoxy)butanoyl)-6,6-dimethyl-3-azabicyclo[3.1.0]hexane-2-carboxylate (46b).** To a solution of **46a** (1.61 mmol, 500 mg) and **29b** (1.61 mmol, 365 mg) in anhydrous DMF (10 mL) was added DIPEA (6.46 mmol, 1.15 mL) and was cooled to 0 °C. HATU (1.93 mmol, 798 mg) was added to the solution under 0 °C and then stirred at room temperature overnight. The reaction mixture was then diluted with ethyl acetate (20 mL) and washed with saturated $NaHCO_3$ solution (2×10 mL), 1 M HCl solution (2×10 mL), and saturated brine solution (2×10 mL) sequentially. The organic layer was dried over anhydrous $Na_2SO_4$ and then concentrated *on vacuo*. The residue was then purified with flash chromatography (20-70% EtOAc in Hexanes as the eluent) to afford **46b** as white solid (500 mg, 67%). $^1$H NMR (400 MHz, Chloroform-*d*) δ 7.48 – 7.32 (m, 5H), 5.15 – 5.04 (m, 2H), 4.47 (s, 1H), 4.33 (dd, *J* = 8.3, 6.1 Hz, 1H), 4.10 – 4.06 (m, 1H), 3.90 – 3.81 (m, 2H), 3.76 (s, 3H), 1.54 – 1.44 (m, 2H), 1.21 (s, 9H), 1.07 (s, 6H), 0.94 (s, 3H).

**(1R,2S,5S)-3-((2S,3R)-2-(((benzyloxy)carbonyl)amino)-3-(tert-butoxy)butanoyl)-6,6-dimethyl-3-azabicyclo[3.1.0]hexane-2-carboxylic acid (45c).** The peptide **46b** (500 mg, 1.08 mmol) was dissolved in THF/$H_2O$ (1:1, 10 mL). LiOH (114 mg, 2.71 mmol) was added at 0 °C. The mixture was stirred at room temperature overnight. Then THF was removed *on vacuum* and the aqueous layer was acidified with 1 M HCl and extracted with dichloromethane (3 x 10 mL). The organic layer was dried over anhydrous $Na_2SO_4$ and concentrated to yield **46c** as white solid (315 mg, 65%). The crude product directly used for next step.

**(S)-methyl 2-((1R,2S,5S)-3-((2S,3R)-2-(((benzyloxy)carbonyl)amino)-3-(tert-butoxy)butanoyl)-6,6-dimethyl-3-azabicyclo[3.1.0]hexane-2-carboxamido)-3-((S)-2-oxopyrrolidin-3-yl)propanoate (46d).** To a solution of **46c** (1.12 mmol, 0.5 g) and **29e** (1.34 mmol, 0.3 g) in anhydrous DMF (10 mL) was added DIPEA (4.48 mmol, 0.8 mL) and was cooled to 0 °C. HATU (1.45 mmol, 0.554 g) was added to the solution under 0 °C and then stirred at room temperature overnight. The reaction mixture was then diluted with ethyl acetate (50 mL) and washed with saturated $NaHCO_3$ solution (2×20 mL), 1 M HCl solution (2×20 mL), and saturated brine solution (2×20 mL) sequentially. The organic layer was dried over anhydrous $Na_2SO_4$ and then concentrated *on vacuo*. The residue was then purified with flash chromatography (0-10% MeOH in Dichloromethane as the eluent) to afford 4**6d** as white gummy solid (520 mg, 75%).$^1$H NMR (400 MHz, Chloroform-*d*) δ 7.46 – 7.28 (m, 5H), 5.13 – 4.98 (m, 2H), 4.60 (ddd, *J* = 10.6, 7.8, 4.4 Hz, 1H), 4.39 (d, *J* = 14.0 Hz, 2H), 4.08 (dd, *J* = 10.7, 5.3 Hz, 1H), 3.95 – 3.81 (m, 1H),

3.81 – 3.70 (m, 4H), 3.31 (dd, *J* = 8.9, 6.3 Hz, 2H), 2.53 – 2.38 (m, 2H), 2.25 – 2.04 (m, 1H), 1.96 – 1.78 (m, 2H), 1.65 – 1.55 (m, 1H), 1.52 – 1.47 (m, 1H), 1.21 (s, 9H), 1.12 (d, *J* = 6.3 Hz, 3H), 1.03 (s, 3H), 0.88 (s, 3H).

**Benzyl ((2S,3R)-3-(tert-butoxy)-1-((1R,2S,5S)-2-(((S)-1-hydroxy-3-((S)-2-oxopyrrolidin-3-yl)propan-2-yl)carbamoyl)-6,6-dimethyl-3-azabicyclo[3.1.0]hexan-3-yl)-1-oxobutan-2-yl)carbamate (46e).** To a stirred solution of compound **46d** (500 mg, 0.81 mmol) in THF (10 mL) was added $LiBH_4$ (2.0 M in THF, 2 mL, 4.06 mmol) in several portions at 0 °C under a nitrogen atmosphere. The reaction mixture was stirred at 0 °C for 1 h, then allowed to warm up to room temperature, and stirred for an additional 2 h. The reaction was quenched by the drop wise addition of 1.0 M HCl (aq) (1.2 mL) with cooling in an ice bath. The solution was diluted with ethyl acetate and $H_2O$. The phases were separated, and the aqueous layer was extracted with ethyl acetate (3×15 mL). The organic phases were combined, dried over $MgSO_4$, filtered, and concentrated on a rotavapor to give a yellow oily residue. Column chromatographic purification of the residue (6% MeOH in $CH_2Cl_2$ as the eluent) afforded **46e** as a white solid (250 mg, 52%). $^1$H NMR (400 MHz, Chloroform-*d*) δ 7.41 – 7.27 (m, 5H), 5.07 (d, *J* = 3.1 Hz, 2H), 4.45 (dd, *J* = 8.1, 5.4 Hz, 0.7H), 4.39 (s, 0.7H), 4.26 (dd, *J* = 7.9, 3.4 Hz, 0.7H), 4.10 (dd, *J* = 10.5, 5.3 Hz, 1.3H), 3.99 (q, *J* = 6.2 Hz, 0.7H), 3.81 – 3.71 (m, 1.3H), 3.70 – 3.60 (m, 1.3H), 3.56 (dd, *J* = 11.6, 5.3 Hz, 0.7H), 3.42 – 3.37 (m, 0.3H), 3.35 – 3.23 (m, 2H), 2.48 – 2.34 (m, 2H), 2.06 – 1.95 (m, 1H), 1.88 – 1.75 (m, 1H), 1.63 – 1.40 (m, 3H), 1.26 (s, 9H), 1.13 (d, *J* = 6.4 Hz, 3H), 1.05 – 0.99 (m, 4H), 0.88 (s, 2H).

**Benzyl ((2S,3R)-3-(tert-butoxy)-1-((1R,2S,5S)-6,6-dimethyl-2-(((S)-1-oxo-3-((S)-2-oxopyrrolidin-3-yl)propan-2-yl)carbamoyl)-3-azabicyclo[3.1.0]hexan-3-yl)-1-oxobutan-2-yl)carbamate (MPI46).** To a solution of **46e** (250 mg, 0.42 mmol) in $CH_2Cl_2$ (6 mL) was added $NaHCO_3$ (146 mg, 4 equiv.,) and the Dess-Martin reagent (550 mg, 1.27 mmol, 3 equiv.,). The resulting mixture was stirred at rt for 3 h. Then the reaction was quenched with a saturated $NaHCO_3$ solution containing 10 % $Na_2S_2O_3$. The layers were separated. The organic layer was then washed with saturated brine solution, dried over anhydrous $Na_2SO_4$ and concentrated *on vacuum*. The residue was then purified with flash chromatography afford **MPI6** as white solid (150 mg, 60%). $^1$H NMR (400 MHz, Chloroform-*d*) δ 9.47 (s, 0.7H), 9.35 (s, 0.3H), 7.37 – 7.19 (m, 5H), 5.10 – 4.95 (m, 2H), 4.47 – 4.14 (m, 3H), 4.10 – 3.96 (m, 1H), 3.92 – 3.83 (m, 1H), 3.77 – 3.57 (m, 2H),

3.22 (pd, $J$ = 9.0, 5.4 Hz, 2H), 2.49 – 2.35 (m, 1H), 2.35 – 2.23 (m, 1H), 1.97 – 1.82 (m, 1H), 1.80 – 1.66 (m, 2H), 1.58 – 1.38 (m, 2H), 1.14 (s, 9H), 1.04 (d, $J$ = 6.2 Hz, 3H), 0.96 (dd, $J$ = 10.8, 5.3 Hz, 4H), 0.84 (s, 2H). $^{13}$C NMR (101 MHz, $CDCl_3$) δ 200.62, 199.51, 180.39, 179.80, 173.04, 172.09, 169.20, 168.94, 155.83, 155.37, 136.52, 136.24, 128.54, 128.49, 128.20, 128.14, 128.07, 75.09, 75.07, 68.54, 66.93, 62.00, 61.00, 58.73, 57.39, 57.26, 57.14, 48.41, 40.76, 40.39, 37.61, 31.07, 30.37, 29.67, 28.98, 28.41, 28.21, 27.94, 27.31, 26.40, 26.18, 25.50, 20.04, 19.36, 18.64, 18.08, 13.70, 12.77.

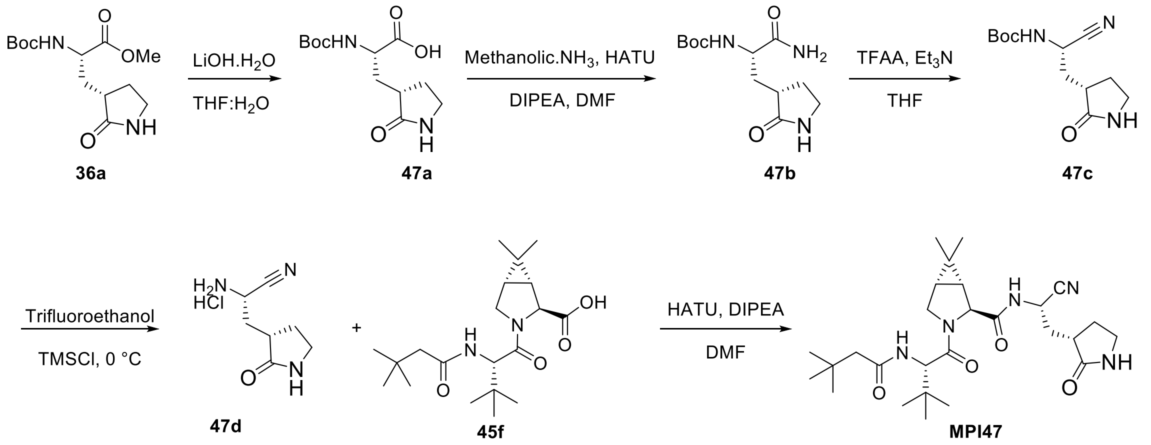


**Scheme S19**: The synthesis of compound **MPI47**

**(S)-2-((tert-butoxycarbonyl)amino)-3-((S)-2-oxopyrrolidin-3-yl)propanoic acid (47a).** The compound **36a** (2 g, 6.99 mmol) was dissolved in THF/$H_2O$ (1:1, 30 mL). LiOH (734 mg, 17.48 mmol) was added at 0 °C. The mixture was stirred at room temperature overnight. Then THF was removed *on vacuum* and the aqueous layer was acidified with 1 M HCl and extracted with dichloromethane (3 x 50 mL). The organic layer was dried over anhydrous $Na_2SO_4$ and concentrated to yield **47a** as white solid (1.1 g). $^1$H NMR (400 MHz, Chloroform-*d*) δ 6.90 (s, 1H), 5.68 (s, 1H), 4.38 (s, 1H), 3.41 (ddt, $J$ = 17.3, 9.8, 6.6 Hz, 2H), 2.69 – 2.55 (m, 1H), 2.51 – 2.37 (m, 1H), 2.27 – 2.14 (m, 1H), 1.99 – 1.80 (m, 2H), 1.44 (s, 9H).

**tert-butyl ((S)-1-amino-1-oxo-3-((S)-2-oxopyrrolidin-3-yl)propan-2-yl)carbamate (47b).** The methanolic ammonia (0.787 mL, 5.51 mmol) and the acid **47a** (0.5 g, 1.84 mmol) were dissolved in dry DMF (10 mL) and the reaction was cooled to 0 °C. HATU (0.908 g, 2.39 mmol) and DIPEA (1.28 mL, 7.35 mmol) were added, and the reaction mixture was allowed warm up to room temperature and stirred for 12 h. The mixture was then poured into water (50 mL) and extracted with ethyl acetate (4×20 mL). The organic layer was washed with aqueous hydrochloric acid 10% v/v (2×20 mL), saturated aqueous $NaHCO_3$ (2×20 mL), brine (2×20 mL) and dried over $Na_2SO_4$. The organic phase was evaporated to dryness and the crude material purified by silica gel column chromatography (0-10% MeOH in $CH_2Cl_2$ as the eluent) to afford **47b** white gummy solid (300 mg, 60%). $^1$H NMR (400 MHz, Chloroform-*d*) δ 7.15 (s, 1H), 6.47 (s, 1H), 6.02 (s, 1H), 5.89 (s, 1H), 4.37 (d, *J* = 10.1 Hz, 1H), 3.49 – 3.30 (m, 2H), 2.53 (qd, *J* = 8.5, 6.5 Hz, 1H), 2.45 – 2.32 (m, 1H), 2.06 (h, *J* = 8.5 Hz, 1H), 1.92 – 1.80 (m, 2H), 1.43 (s, 9H).

**tert-butyl ((S)-1-cyano-2-((S)-2-oxopyrrolidin-3-yl)ethyl)carbamate (47c).** $Et_3N$ (0.47 mL, 3.32 mmol) is added to **47b** (300 mg, 1.11 mmol) dissolved in anhydrous THF (10 mL). The stirred solution is cooled in an ice bath and trifluoroacetic anhydride (0.23 mL, 1.66 mmol) is added drop wise. The mixture is allowed to reach room temperature, and after 1 h the reaction is quenched with water (10 mL). The residue obtained after evaporation of the organic phase is extracted with ether (3 x 30 mL); The organic layer is dried ($Na_2SO_4$) and concentrated. The organic phase was evaporated to dryness and the crude material purified by silica gel column chromatography (0-10% MeOH in $CH_2Cl_2$ as the eluent) to afford **47c** white solid (270 mg, 90%). $^1$H NMR (400 MHz, Chloroform-*d*) δ 6.54 (s, 1H), 5.98 (d, *J* = 7.8 Hz, 1H), 4.66 (t, *J* = 8.1 Hz, 1H), 3.42 – 3.29 (m, 2H), 2.45 (dddt, *J* = 23.8, 11.9, 5.9, 3.4 Hz, 2H), 2.28 (ddd, *J* = 14.1, 9.6, 6.3 Hz, 1H), 1.99 – 1.77 (m, 2H), 1.45 (s, 9H). $^{13}$C NMR (101 MHz, $CDCl_3$): δ 178.99, 154.82, 118.95, 45.63, 40.47, 37.82, 34.44, 28.26, 28.23, 8.51.

**(S)-2-amino-3-((S)-2-oxopyrrolidin-3-yl)propanenitrile hydrochloride (47d). 47c** (270 mg, 1.11 mmol) dissolved in anhydrous trifluoroethanol (10 mL). The stirred solution is cooled in an ice bath and TMSCl (0.2 mL) is added drop wise. The mixture is same temperature until the reaction completion. After completion remove solvent by rotavapor, crude **47d** directly used for next step. $^1$H NMR (400 MHz, Deuterium Oxide) δ 4.74 (d, *J* = 2.6 Hz, 1H), 3.45 – 3.24 (m, 2H), 2.76 (dtd, *J* = 10.1, 8.4, 6.1 Hz, 1H), 2.45 – 2.30 (m, 1H), 2.27 – 2.02 (m, 2H), 1.85 (tdd, *J* = 15.6, 11.6, 6.8 Hz, 1H).

**(1R,2S,5S)-N-((S)-1-cyano-2-((S)-2-oxopyrrolidin-3-yl)ethyl)-3-((S)-2-(3,3-dimethylbutanamido)-3,3-dimethylbutanoyl)-6,6-dimethyl-3-azabicyclo[3.1.0]hexane-2-carboxamide (MPI47).** The compound **47d** (57 mg, 0.3 mmol) and the acid **45f** (0.1 g, 0.273 mmol) were dissolved in dry DMF (10 mL) and the reaction was cooled to 0 °C. HATU (125 mg, 0.355 mmol) and DIPEA (0.19 mL, 1.09 mmol) were added, and the reaction mixture was allowed warm up to room temperature and stirred for 12 h. The mixture was then poured into water (20 mL) and extracted with ethyl acetate (4×20 mL). The organic layer was washed with aqueous hydrochloric acid 10% v/v (2×20 mL), saturated aqueous $NaHCO_3$ (2×20 mL), brine (2×20 mL) and dried over $Na_2SO_4$. The organic phase was evaporated to dryness and the crude material purified by silica gel column chromatography (0-10% MeOH in $CH_2Cl_2$ as the eluent) to afford **MPI47** white solid (55 mg, 40%). $^1$H NMR (400 MHz, Chloroform-*d*) δ 7.75 (d, *J* = 8.5 Hz, 1H), 5.06 (q, *J* = 8.1 Hz, 1H), 4.67 – 4.50 (m, 1H), 4.39 – 4.10 (m, 3H), 3.77 – 3.60 (m, 2H), 3.34 (ddt, *J* = 15.5, 8.6, 4.5 Hz, 2H), 2.56 – 2.37 (m, 2H), 2.35 – 2.23 (m, 1H), 2.14 – 2.04 (m, 3H), 2.00 – 1.83 (m, 2H), 1.48 (t, *J* = 3.6 Hz, 2H), 1.05 – 0.98 (m, 24H).

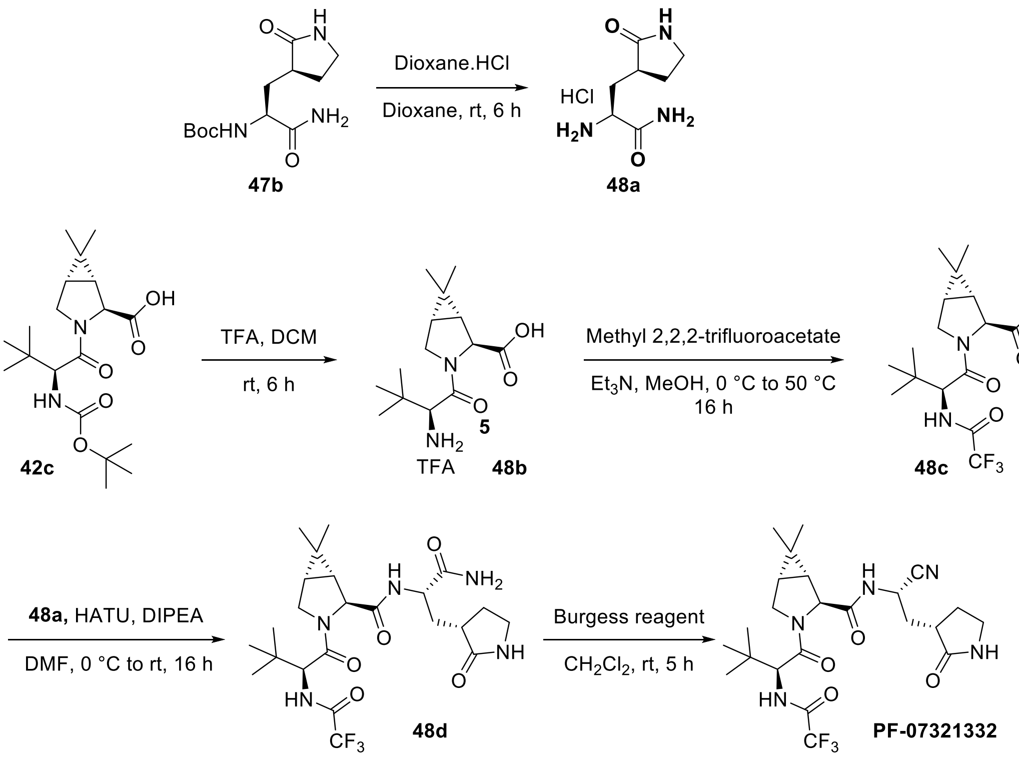

**Scheme 20:** The synthesis of compound **PF-07321332**

**3-[(3S)-2-Oxopyrrolidin-3-yl]-L-alaninamide (48a), HCl salt**. To a solution of **47b** (400 mg, 1.47 mmol) in 1,4-dioxane (10 mL) was added dropwise a HCl solution in 1,4-dioxane (4 M, 3.7 mL). The resulting solution was stirred at room temperature for 6 h. Then residue was then concentrated *on vacuo* to afford **48a** as light-yellow hygroscopic solid (365 mg). Crude **48a** compound directly used next step.

**(1R,2S,5S)-6,6-Dimethyl-3-(3-methyl-L-valyl)-3-azabicyclo[3.1.0]hexane-2-carboxylic acid, TFA salt (48b).** A solution of TFA (1 ml, 13.3 mol) was added to a solution of **42c** (500 mg, 1.33 mmol) in dichloromethane (10 ml), and the reaction mixture was stirred at 25 °C for 6 h. Removal of solvents afforded TFA salt of **48b** as a white solid (450 mg). This material was used directly in the following step.

**(1R,2S,5S)-6,6-Dimethyl-3-[3-methyl-N-(trifluoroacetyl)-L-valyl]-3-azabicyclo[3.1.0]hexane-2-carboxylic acid (48c).** To a 0 °C solution of the TFA salt of **48b** (500 mg, 1.31 mmol) in methanol (3 ml) was added triethylamine (0.64 ml, 4.58 mmol), followed by Methyl trifluoroacetate (0.2 ml, 1.96 mmol), whereupon the reaction mixture was allowed to warm to 50 °C, and was stirred for 16 h. It was then concentrated in vacuo at 50 °C, and the residue was diluted with water (10 ml) and adjusted to a pH of 3 to 4 by addition of 1 M HCl. After extraction of the aqueous layer with ethyl acetate (3 x 20 ml), the combined organic layers were washed with saturated aqueous sodium chloride solution (20 ml), dried over sodium sulfate, filtered, and concentrated to afford **48c** as a white solid (400 mg). $^{1}$H NMR (400 MHz, DMSO-d6) δ 9.44 (d, J = 8.5 Hz, 1H), 4.44 (d, J = 8.5 Hz, 1H), 4.16 (s, 1H), 3.85 (dd, J = 10.5, 5.3 Hz, 1H), 3.73 (d, J = 10.5 Hz, 1H), 1.54 (dd, J = 7.6, 5.1 Hz, 1H), 1.44 (d, J = 7.5 Hz, 1H), 1.01 (d, J = 3.5 Hz, 12H), 0.83 (s, 3H).

**(1R,2S,5S)-N-{(2S)-1-Amino-1-oxo-3-[(3S)-2-oxopyrrolidin-3-yl]propan-2-yl}-6,6-dimethyl-3-[3-methyl-N-(trifluoroacetyl)-L-valyl]-3-azabicyclo[3.1.0]hexane-2-carboxamide (48d).** To a 0 °C solution of TFA salt **48a** (184 mg, 0.906 mmol) and **48c** (300 mg, 0.824 mmol) in a mixture of N,N-dimethylformamide (10 ml) was added HATU (375 mg, 0.989 mmol), followed by drop-wise addition of NMM (0.27 ml, 3.29 mmol). The reaction mixture was then allowed to warm to 25 °C and was stirred for 16 h, The reaction mixture was then diluted with ethyl acetate (20 mL) and washed with saturated $NaHCO_3$ solution (2×10 mL), 1 M HCl solution (2×10 mL), and saturated brine solution (2×10 mL) sequentially. The organic layer was dried over anhydrous $Na_2SO_4$ and then concentrated on vacuo. The residue was then purified with flash chromatography (0-10% MeOH in DCM as the eluent) to afford **48d** as white solid (100 mg, 21%). $^1$H NMR (400 MHz, DMSO-d6) δ 8.42 (d, J = 6.7 Hz, 1H), 8.09 (d, J = 9.3 Hz, 1H), 7.06 (d, J = 18.7 Hz, 2H), 6.16 (s, 1H), 4.49 (d, J = 9.2 Hz, 1H), 4.36 – 4.27 (m, 1H), 4.19 (d, J = 1.6 Hz, 1H), 4.11 – 4.02 (m, 1H), 3.63 (d, J = 10.3 Hz, 1H), 3.21 (dd, J = 9.4, 6.7 Hz, 2H), 2.41 (q, J = 8.1 Hz, 1H), 2.34 – 2.23 (m, 1H), 2.01 – 1.79 (m, 2H), 1.74 (td, J = 10.0, 2.5 Hz, 1H), 1.47 – 1.36 (m, 2H), 0.98 (d, J = 1.6 Hz, 12H), 0.80 (d, J = 1.6 Hz, 3H).

**(1R,2S,5S)-N-{(1S)-1-Cyano-2-[(3S)-2-oxopyrrolidin-3-yl]ethyl}-6,6-dimethyl-3-[3-methylN-(trifluoroacetyl)-L-valyl]-3-azabicyclo[3.1.0]hexane-2-carboxamide (PF-07321332)**. Methyl N-(triethylammoniosulfonyl)carbamate (Burgess reagent; 115 mg, 0.483 mmol) was added to a solution of **48d** (100 mg, 0.193 mmol) in dichloromethane (3 ml). After the reaction mixture had been stirred at rt for 5 h. The reaction mixture was quenched by a mixture of saturated aqueous sodium bicarbonate solution (20 ml) and saturated aqueous sodium chloride solution (10 ml). The separated organic phase was concentrated. The residue was then purified with flash chromatography (0-10% MeOH in DCM as the eluent) to afford **PF-07321332** as white solid (60 mg, 62%). $^1$H NMR (400 MHz, DMSO-$d_6$) δ 9.42 (d, $J$ = 8.4 Hz, 1H), 9.03 (d, $J$ = 8.5 Hz, 1H), 7.67 (s, 1H), 5.03 – 4.94 (m, 1H), 4.42 (d, $J$ = 8.5 Hz, 1H), 4.16 (s, 1H), 3.92 (dd, $J$ = 10.3, 5.5 Hz, 1H), 3.70 (d, $J$ = 10.3 Hz, 1H), 3.15 (t, $J$ = 9.0 Hz, 1H), 3.05 (q, $J$ = 9.2, 8.7 Hz, 1H), 2.40 (td, $J$ = 10.3, 9.6, 4.1 Hz, 1H), 2.13 (ddt, $J$ = 27.7, 14.2, 6.2 Hz, 2H), 1.78 – 1.67 (m, 2H), 1.58 (dd, $J$ = 7.6, 5.3 Hz, 1H), 1.33 (d, $J$ = 7.6 Hz, 1H), 1.04 (s, 3H), 0.99 (s, 9H), 0.86 (s, 3H).

**Table S1. Statistics of crystallographic analysis of $M^{Pro}$ in complexed with different inhibitors.**

| Protein/Ligand (PDI entry) | **MPI29** (7S6W) | **MPI30** (7S6X) | **MPI32** (7S6Y) | **MPI33** (7S6Z) | **MPI34** (7S70) |
|---|---|---|---|---|---|
| Data Collection | | | | | |
| Space group | P1 | C121 | C121 | C121 | P1 |
| cell dimensions | | | | | |
| *a*, *b*, *c* (Å) | 56.01 61.16 63.89 | 94.91 80.64 54.32 | 95.26 81.29 54.31 | 95.30 81.31 54.26 | 55.12 62.07 62.29 |
| α, β, γ (°) | 80.26 68.29 70.33 | 90.00 116.90 90.00 | 90.00 116.97 90.00 | 90.00 116.95 90.00 | 80.41 69.53 69.61 |
| Resolution (Å) | 24.83-2.29 (2.37-2.29) | 58.38-1.80 (1.84-1.80) | 24.21-1.85 (1.89-1.85) | 24.20-1.85 (1.89-1.85) | 58.19-2.60 (2.72-2.60) |
| $R_{merge}$ | 0.219 (2.633) | 0.070 (1.242) | 0.100 (1.217) | 0.074 (0.752) | 0.173 (0.896) |
| $I/\sigma I$ | 9.8 (1.0) | 12.4 (1.1) | 12.3 (1.4) | 19.4 (2.8) | 4.6 (0.7) |
| Completeness (%) | 98.7 (88.8) | 95.5 (89.7) | 99.9 (100.0) | 99.9 (100.0) | 99.9 (99.7) |
| Redundancy | 10.8 (9.1) | 6.2 (5.5) | 12.3 (9.3) | 11.8 (8.3) | 4.8 (4.4) |
| Refinement | | | | | |
| Resolution (Å) | 24.827 - 2.287 | 48.442 - 1.800 | 24.206 - 1.850 | 24.205 - 1.850 | 49.026 - 2.600 |
| No. Reflections | 32743 (2845) | 32247 (3054) | 28445 (2874) | 30207 (2961) | 11357 (1120) |
| $R_{work}/R_{free}$ | 0.1986 /0.2418 | 0.1878/0.2069 | 0.2290/0.2580 | 0.2018/0.2215 | 0.2489/0.3224 |
| No. atoms | | | | | |
| Protein | 4852 | 2356 | 2406 | 2404 | 2360 |
| Water | 229 | 148 | 168 | 230 | 20 |
| *B* factors | | | | | |
| Protein | 23.652 | 39.306 | 32.434 | 29.981 | 55.277 |
| Water | 36.727 | 41.465 | 33.846 | 33.947 | 49.108 |
| R.m.s deviations | | | | | |
| Bond lengths (Å) | 0.009 | 0.009 | 0.009 | 0.009 | 0.010 |
| Bond angles (°) | 1.37 | 1.26 | 1.29 | 1.24 | 1.34 |

| Protein/Ligand (PDI entry) | **MPI35** (7S71) | **MPI36** (7S72) | **MPI37** (7S73) | **MPI38** (7S74) | **MPI42** (7S75) |
|---|---|---|---|---|---|
| Data Collection | | | | | |
| Space group | C121 | P1 | C121 | C121 | C121 |
| cell dimensions | | | | | |
| *a*, *b*, *c* (Å) | 95.36 81.35 54.46 | 55.25 61.85 62.10 | 97.11 80.93 54.61 | 96.56 80.86 54.22 | 98.84 80.37 51.94 |
| α, β, γ (°) | 90.00 117.17 90.00 | 80.35 69.50 69.65 | 90.00 116.77 90.00 | 90.00 117.23 90.00 | 90.00 114.76 90.00 |
| Resolution (Å) | 24.23-1.85 (1.89-1.85) | 58.00-2.50 (2.60-2.50) | 24.38-1.85 (1.89-1.85) | 58.86-1.70 (1.73-1.70) | 47.16-1.80 (1.84-1.80) |
| $R_{merge}$ | 0.092 (2.322) | 0.487 (3.127) | 0.073 (0.462) | 0.286 (2.276) | 0.085 (1.087) |
| $I/\sigma I$ | 16.3 (1.4) | 2.7 (0.7) | 18.4 (3.4) | 3.5 (1.2) | 12.0 (1.2) |
| Completeness (%) | 100.0 (100.0) | 99.4 (98.9) | 99.6 (99.2) | 96.3 (95.1) | 99.0 (95.5) |
| Redundancy | 10.1 (7.7) | 6.4 (6.7) | 9.4 (6.6) | 3.6 (3.8) | 7.8 (3.5) |
| Refinement | | | | | |
| Resolution (Å) | 24.235 - 1.850 | 49.150 - 2.500 | 23.878 - 1.850 | 48.211 - 1.700 | 42.629 - 1.800 |
| No. Reflections | 31202 (2961) | 12641 (1252) | 32073 (3183) | 76581 (7457) | 33631 (3255) |
| $R_{work}/R_{free}$ | 0.1945/0.2186 | 0.2411/0.2958 | 0.2058/0.2381 | 0.2507/0.2722 | 0.2696/0.3086 |
| No. atoms | | | | | |
| Protein | 2408 | 2363 | 2398 | 4704 | 2398 |
| Water | 210 | 36 | 250 | 366 | 161 |
| *B* factors | | | | | |
| Protein | 32.036 | 52.233 | 27.454 | 31.917 | 32.667 |
| Water | 36.452 | 46.333 | 32.194 | 36.152 | 34.743 |
| R.m.s deviations | | | | | |
| Bond lengths (Å) | 0.009 | 0.010 | 0.012 | 0.010 | 0.008 |
| Bond angles (°) | 1.33 | 1.27 | 2.03 | 1.49 | 1.59 |

### NMR Spectroscopies of Synthesized Compounds

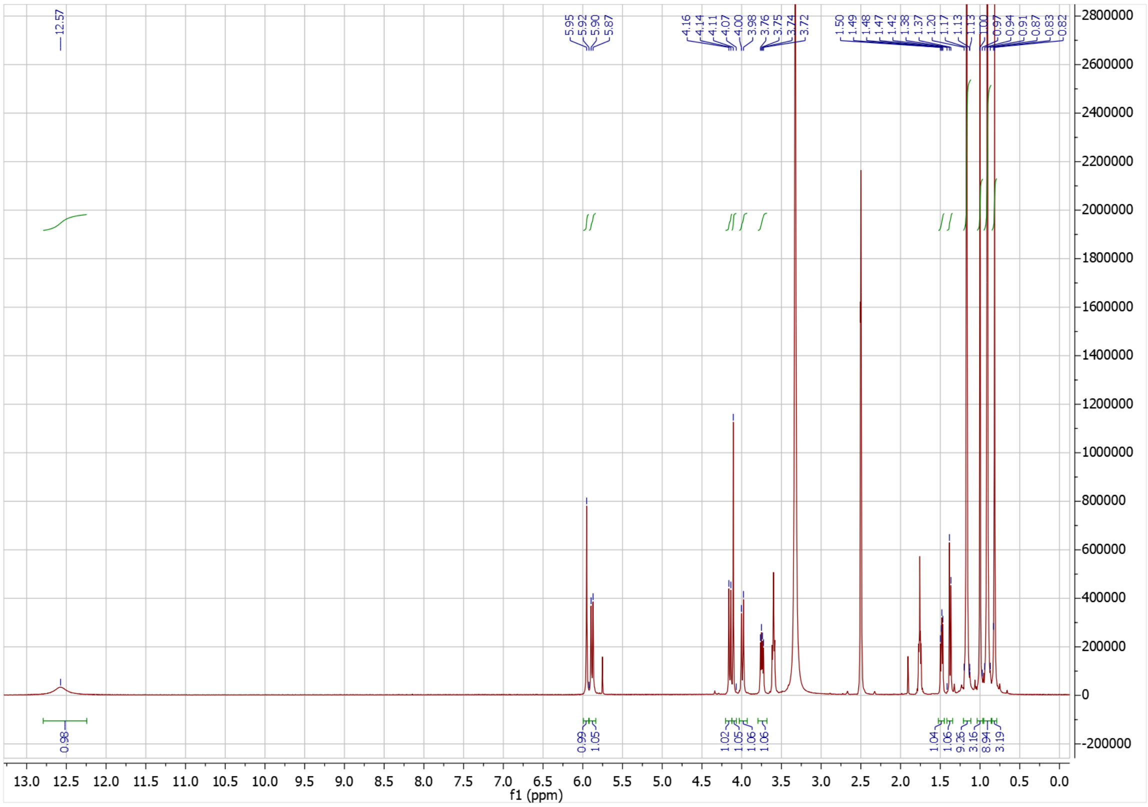

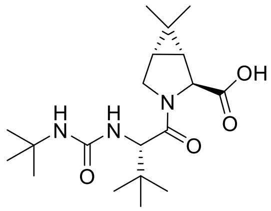

**29d**

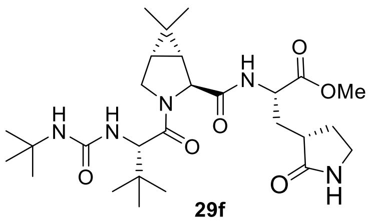

**29f**

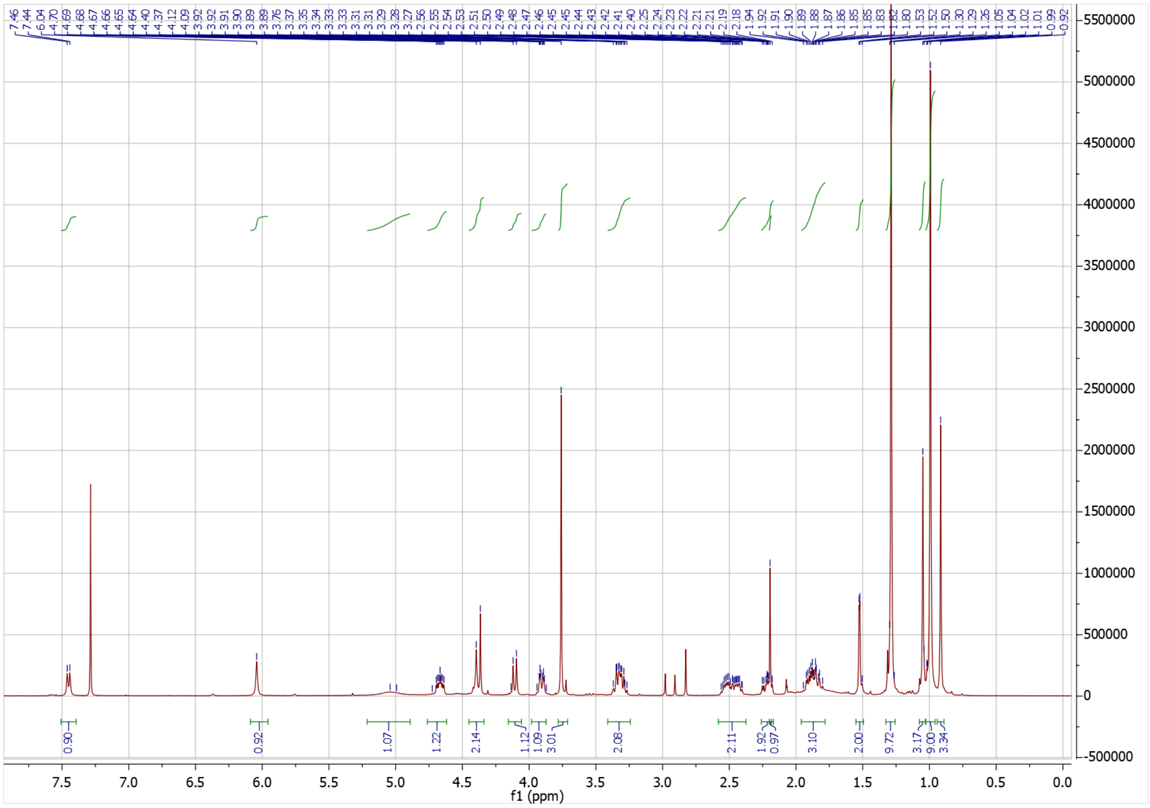

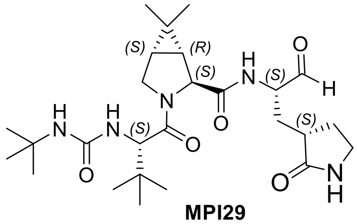

**MPI29**

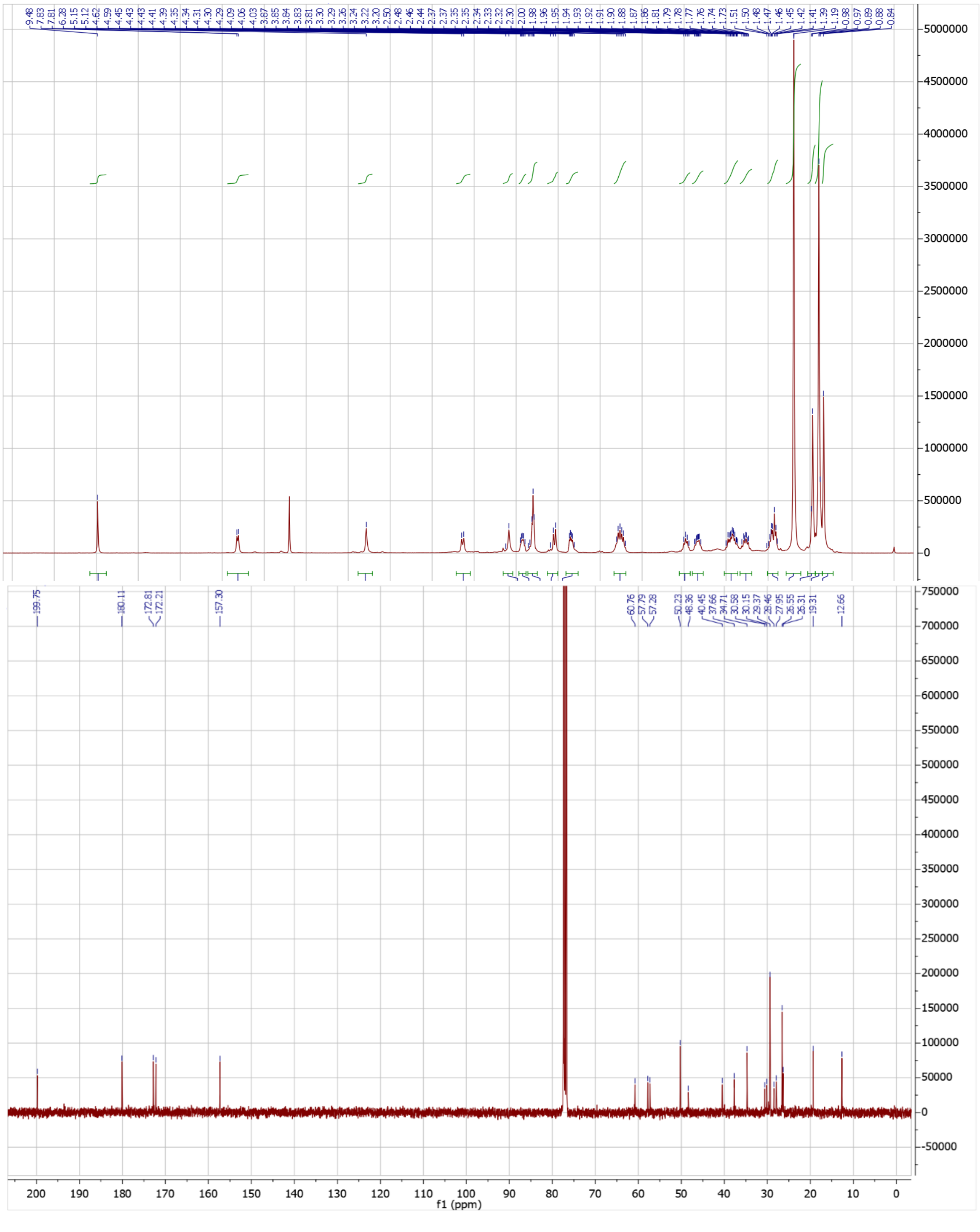

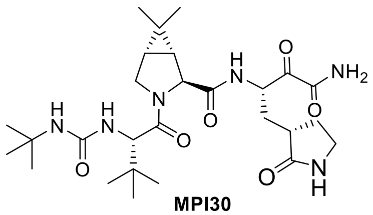

**MPI30**

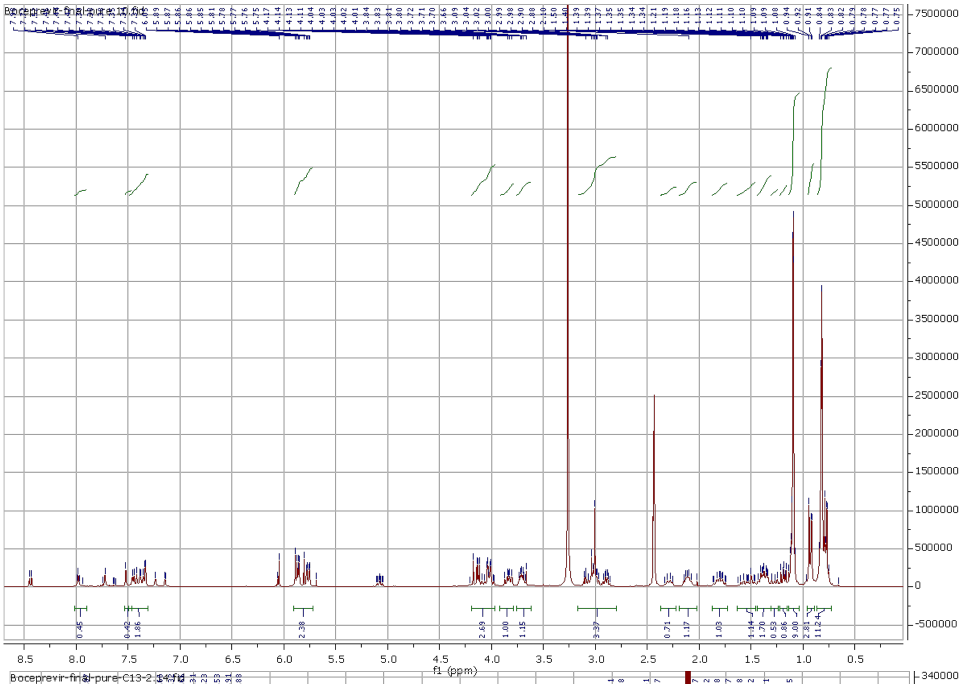

f1 (ppm)


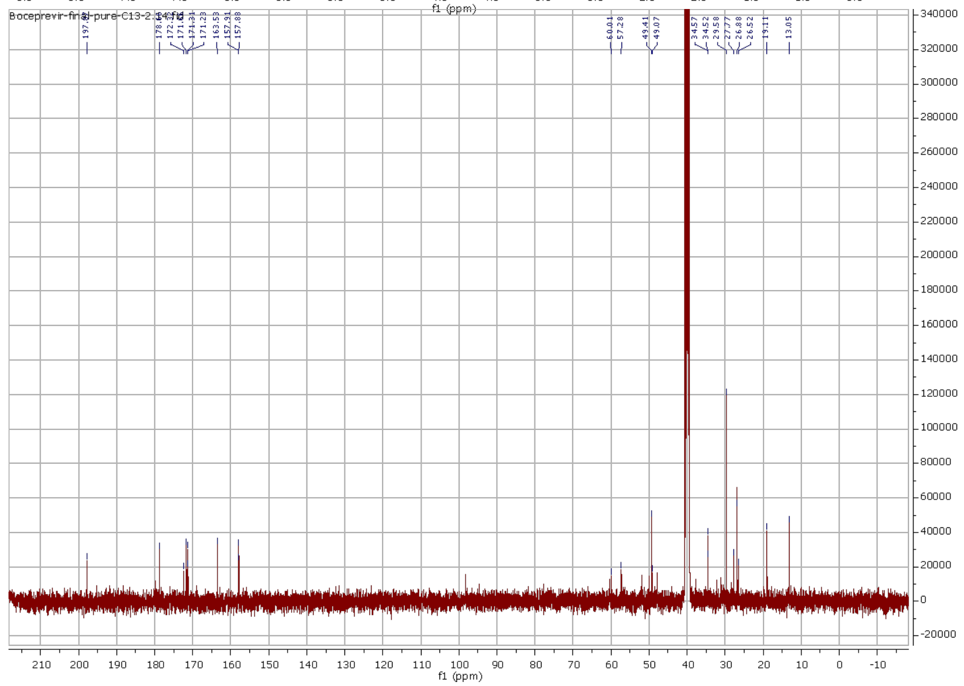

Boceprevir-final-pure-C13-2.1.fid
197.79
178.44
171.31
171.23
163.53
157.31
157.88
60.01
57.28
49.41
49.07
34.57
34.52
29.58
27.77
26.88
26.52
19.11
13.05
f1 (ppm)

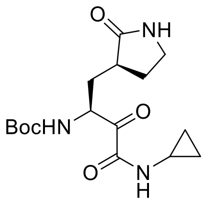

**31c**

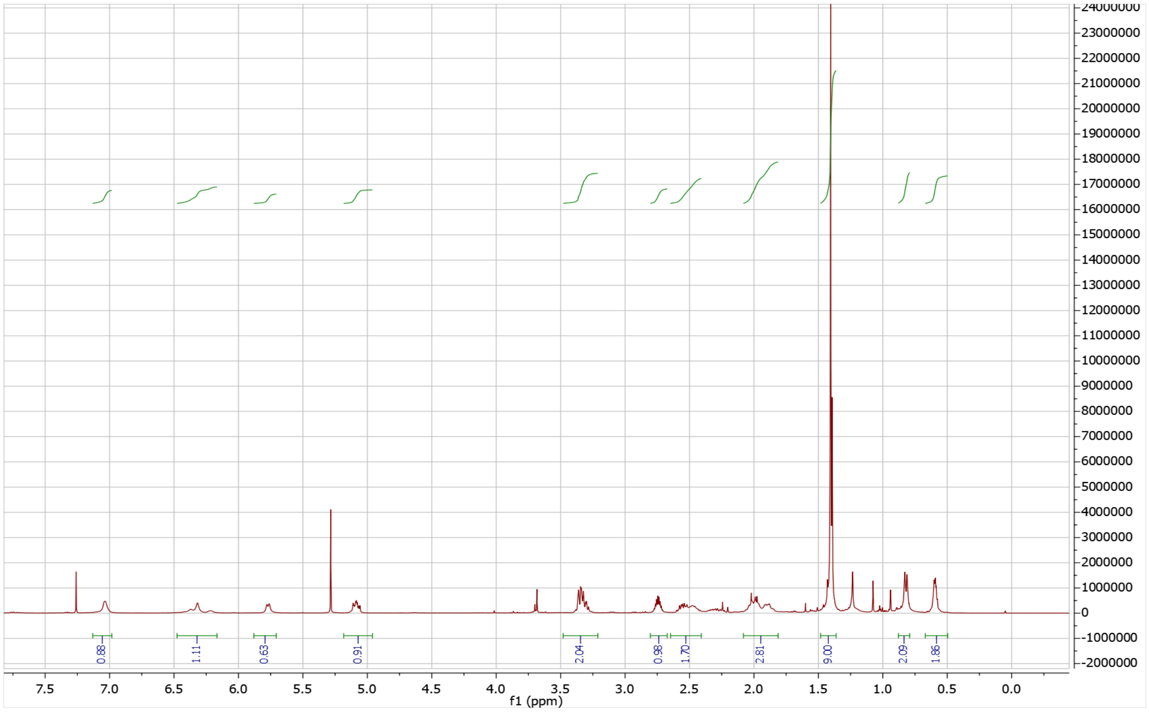

7.5
7.0
6.5
6.0
5.5
5.0
4.5
4.0
3.5
3.0
2.5
2.0
1.5
1.0
0.5
0.0
f1 (ppm)
0.88
1.11
0.63
0.91
2.04
0.98
1.70
2.81
9.00
2.09
1.86
24000000
23000000
22000000
21000000
20000000
19000000
18000000
17000000
16000000
15000000
14000000
13000000
12000000
11000000
10000000
9000000
8000000
7000000
6000000
5000000
4000000
3000000
2000000
1000000
0
-1000000
-2000000

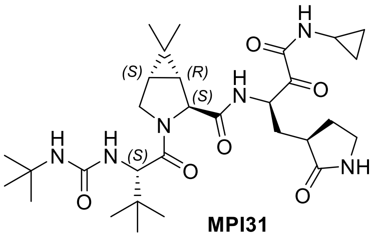

**MPI31**

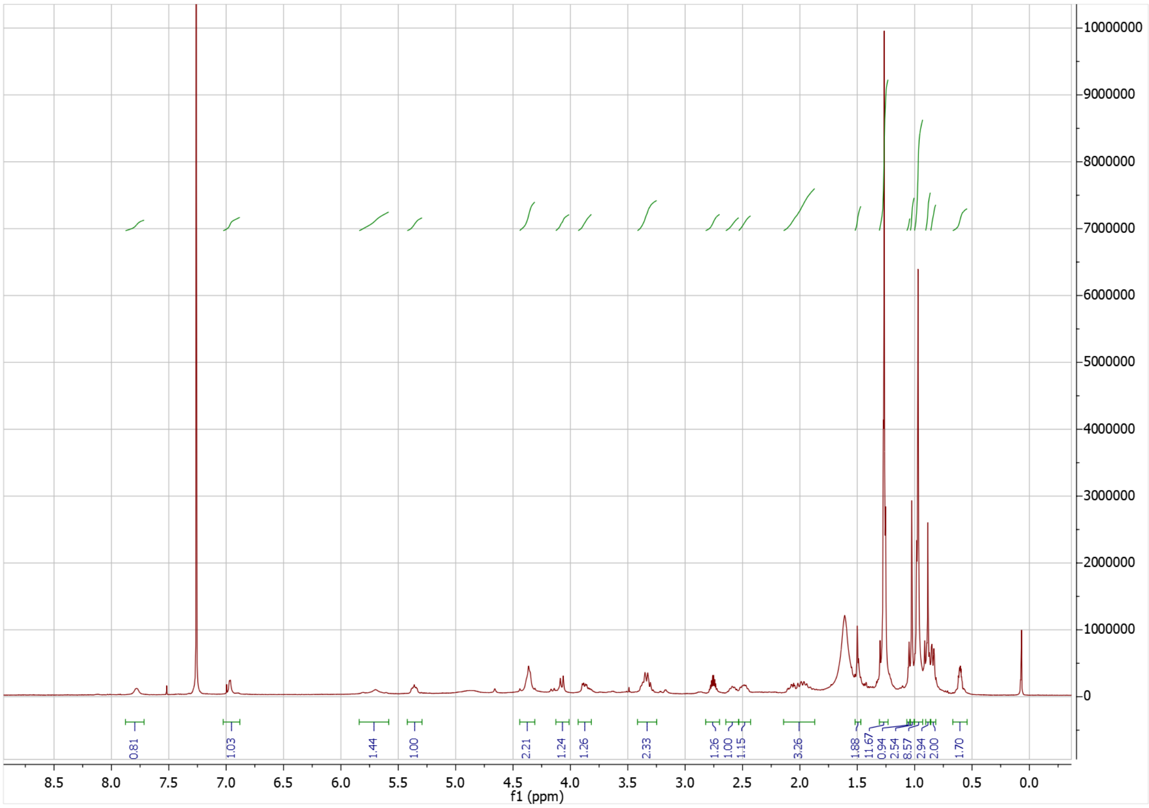

0.81
1.03
1.44
1.00
2.21
1.24
1.26
2.33
1.26
1.00
1.15
3.26
1.88
11.67
0.94
2.54
8.57
2.94
2.00
1.70
f1 (ppm)

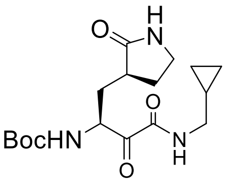
O
H
N
O
BocHN
N
H
O

**32e**

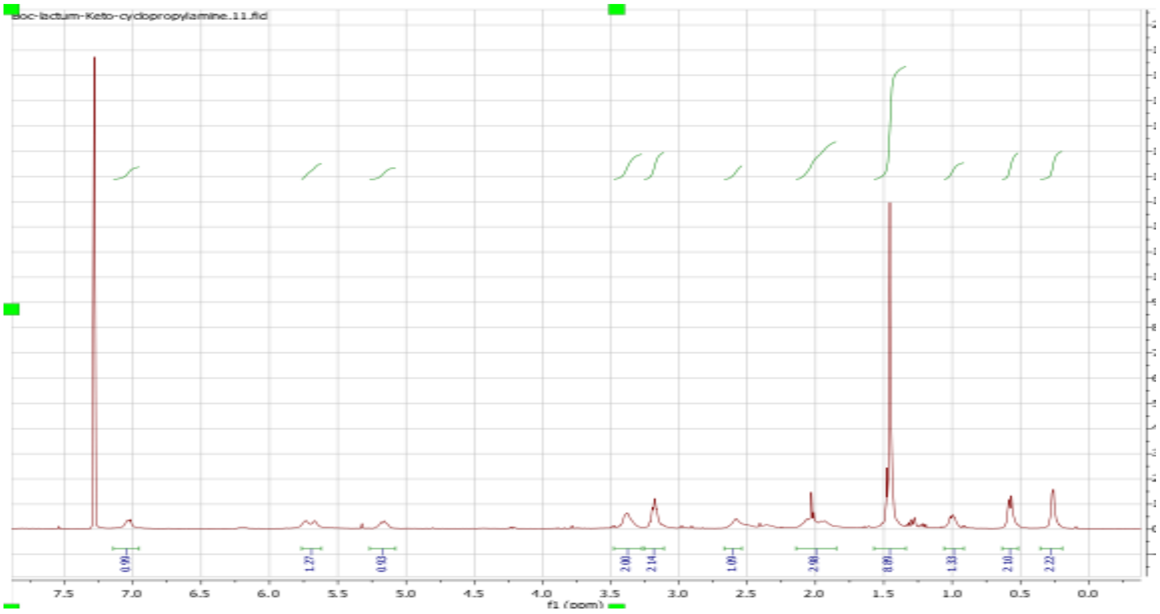
Boc-lactum-Keto-cyclopropylamine.11.fid
7.5
7.0
6.5
6.0
5.5
5.0
4.5
4.0
3.5
3.0
2.5
2.0
1.5
1.0
0.5
0.0
f1 (ppm)

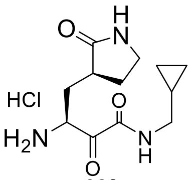
O
H
N
HCl
O
$H_2N$
N
H
O

**32f**

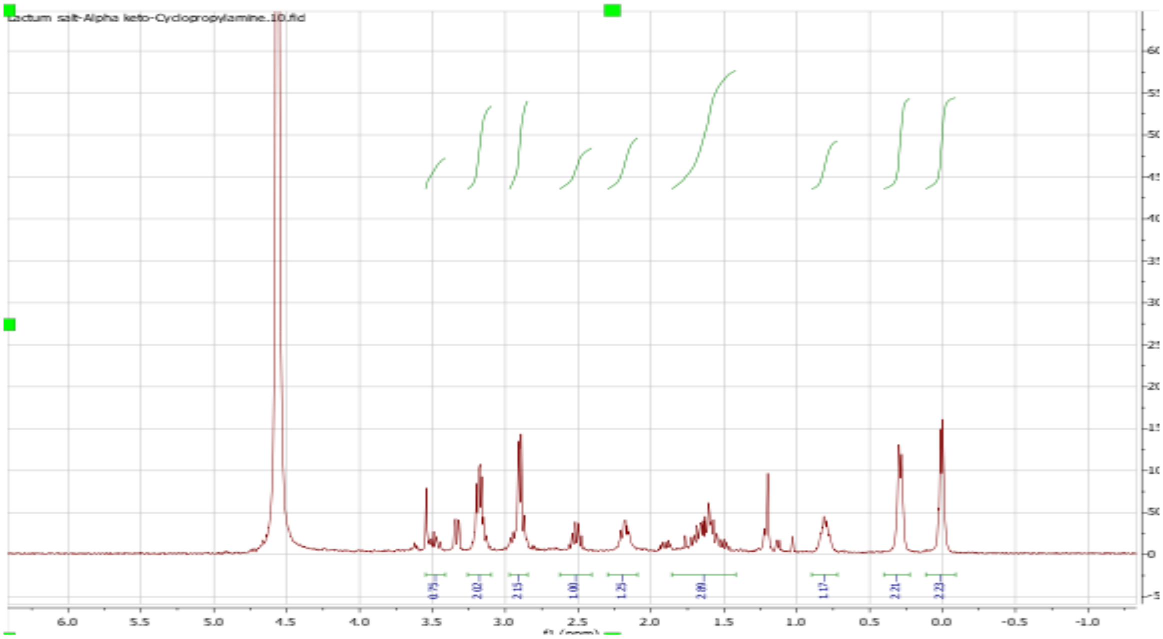
Lactum salt-Alpha keto-Cyclopropylamine.10.fid
6.0
5.5
5.0
4.5
4.0
3.5
3.0
2.5
2.0
1.5
1.0
0.5
0.0
-0.5
-1.0
f1 (ppm)

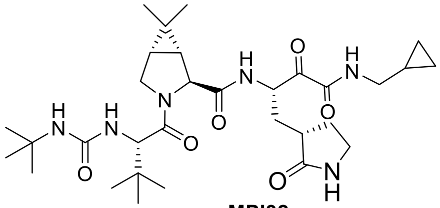

**MPI32**

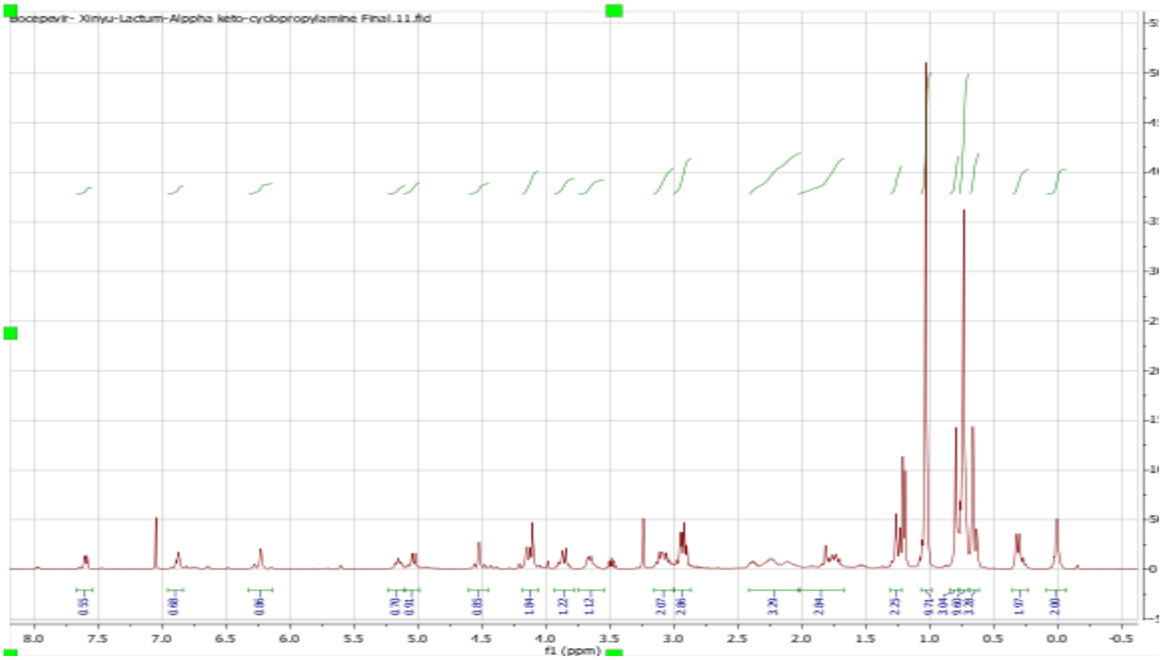
Bocepevir- Xinyu-Lactum-Alppha keto-cyclopropylamine Final.11.fid
f1 (ppm)

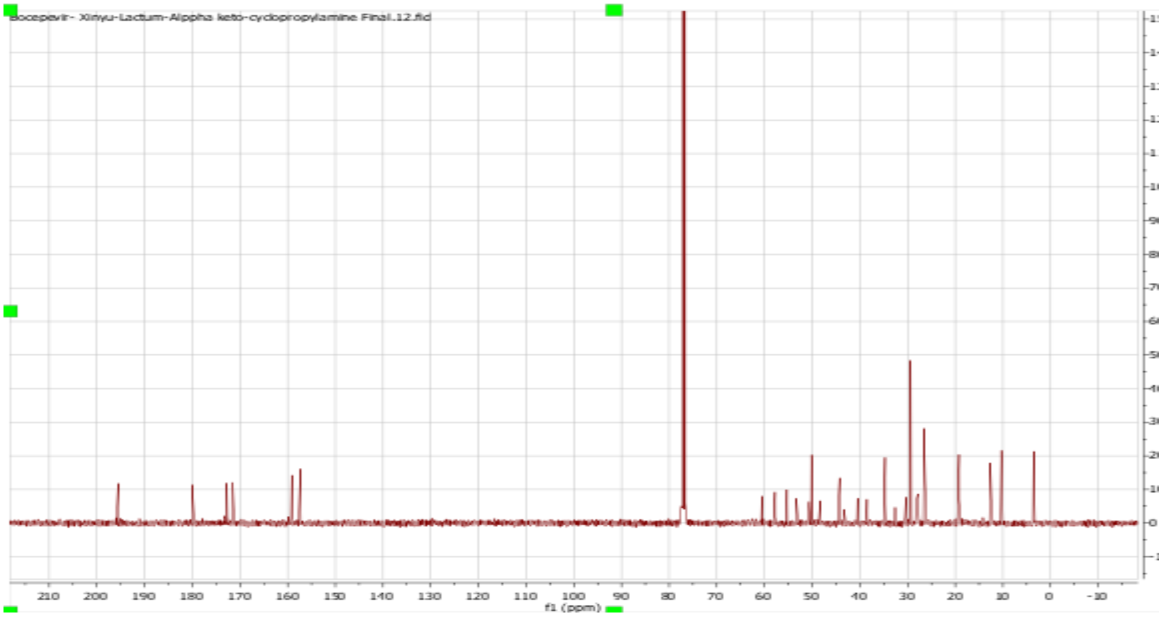
Bocepevir- Xinyu-Lactum-Alppha keto-cyclopropylamine Final.12.fid
f1 (ppm)

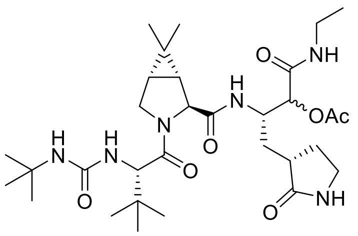

**33a**

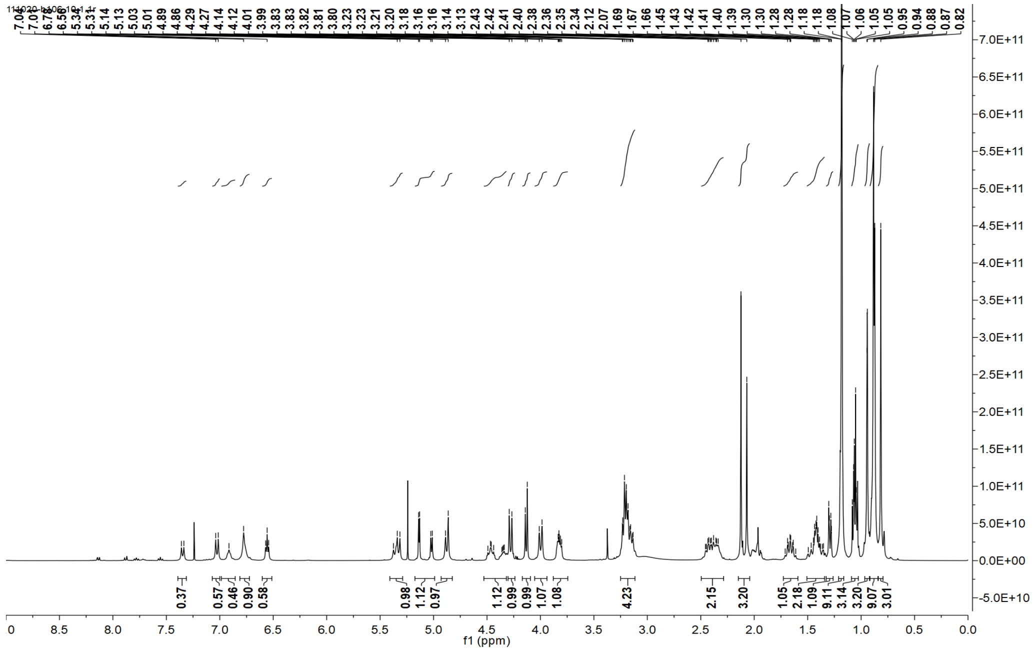

0.37
0.57
0.46
0.90
0.58
0.98
1.12
0.97
1.12
0.99
0.99
1.07
1.08
4.23
2.15
3.20
1.05
2.18
1.09
9.11
3.14
3.20
9.07
3.01
f1 (ppm)


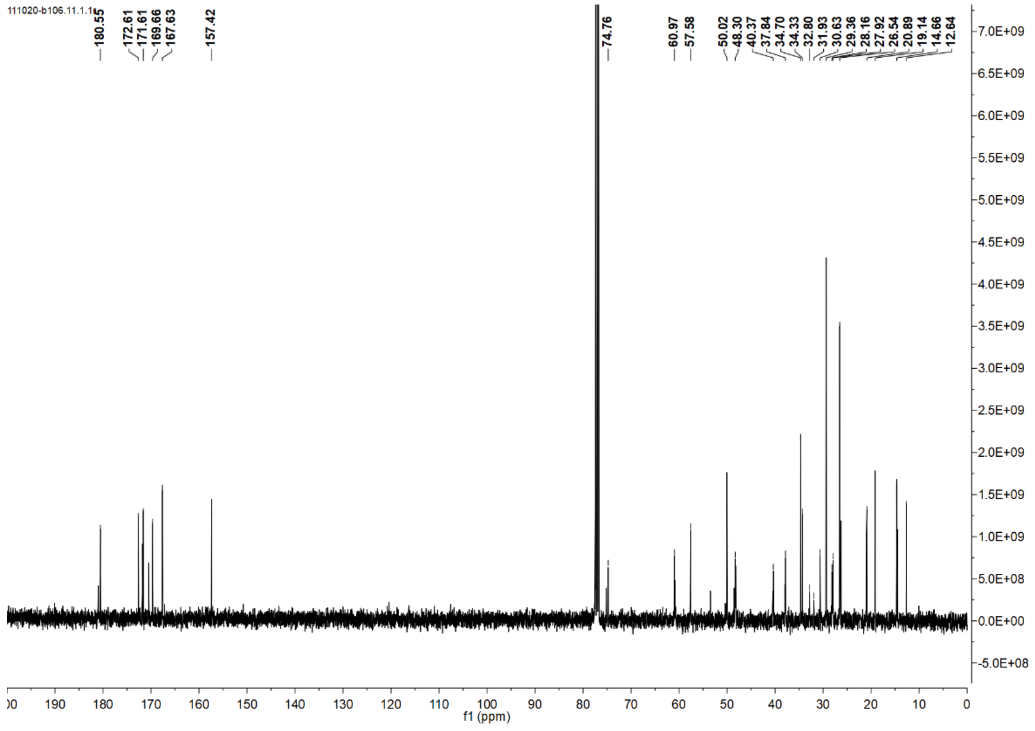

111020-b106.11.1.1r
180.55
172.61
171.61
169.66
167.63
157.42
74.76
60.97
57.58
50.02
48.30
40.37
37.84
34.70
34.33
32.80
31.93
30.63
29.36
28.16
27.92
26.54
20.89
19.14
14.66
12.64
f1 (ppm)

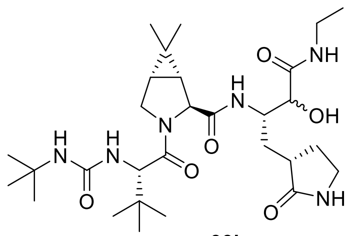
O
NH
H
N
OH
N
O
H
N
H
N
N
O
O
O
NH

**33b**

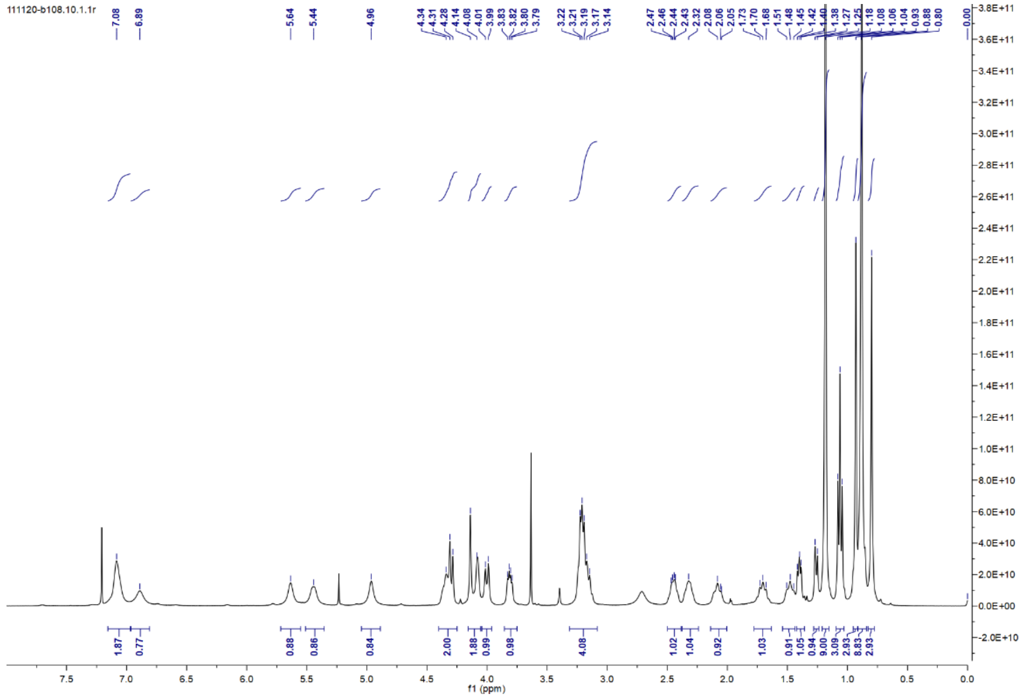
111120-b108.10.1.1r
7.08
6.89
5.64
5.44
4.96
1.87
0.77
0.88
0.86
0.84
2.00
1.88
0.99
0.98
4.08
1.02
1.04
0.92
1.03
f1 (ppm)

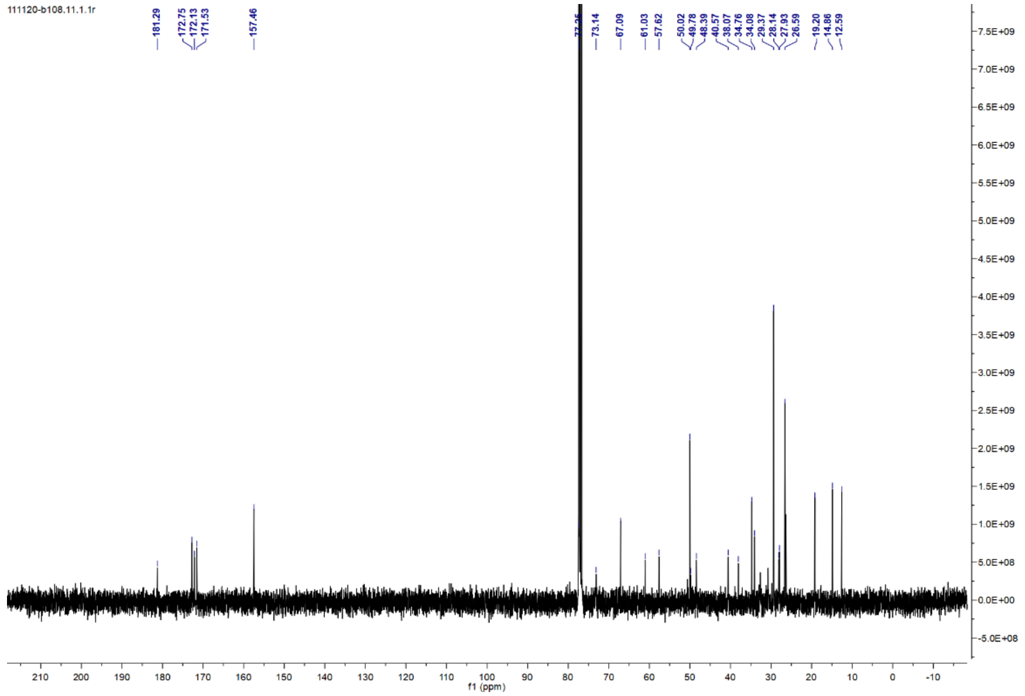
111120-b108.11.1.1r
181.29
172.75
172.13
171.53
157.46
73.14
67.09
61.03
57.62
50.02
49.78
48.39
40.57
38.07
34.76
34.08
29.37
28.14
27.93
26.59
19.20
14.86
12.59
f1 (ppm)

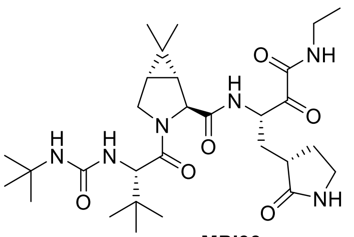

NH
H
N
O
O
O
O
N
H
N
H
N
O
O
NH


**MPI33**

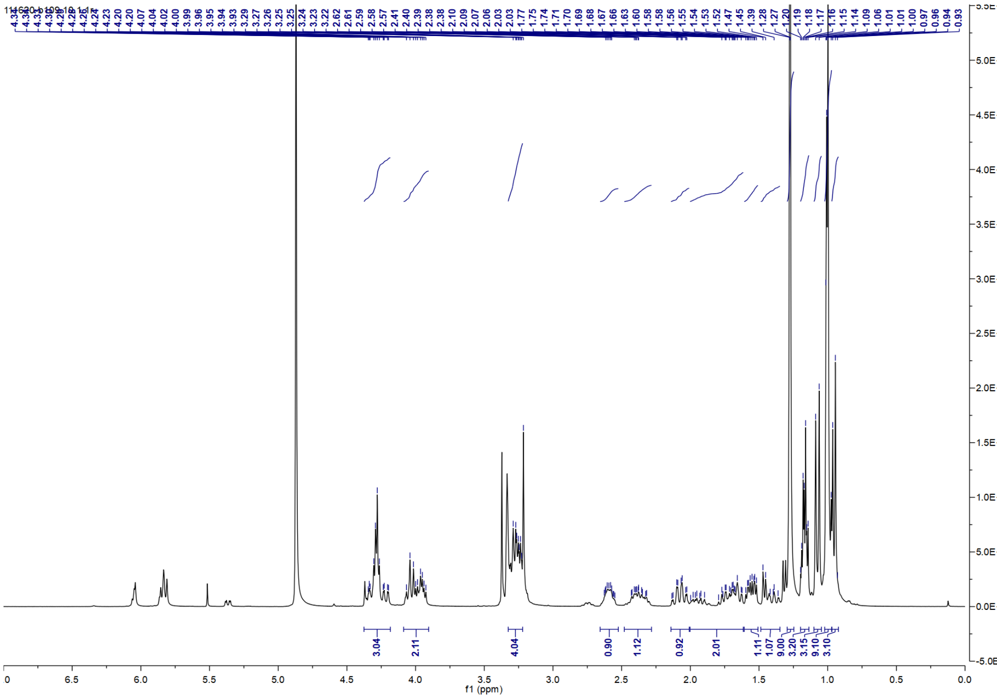

3.04
2.11
4.04
0.90
1.12
0.92
2.01
1.11
1.07
9.00
3.20
3.15
9.10
3.10
f1 (ppm)


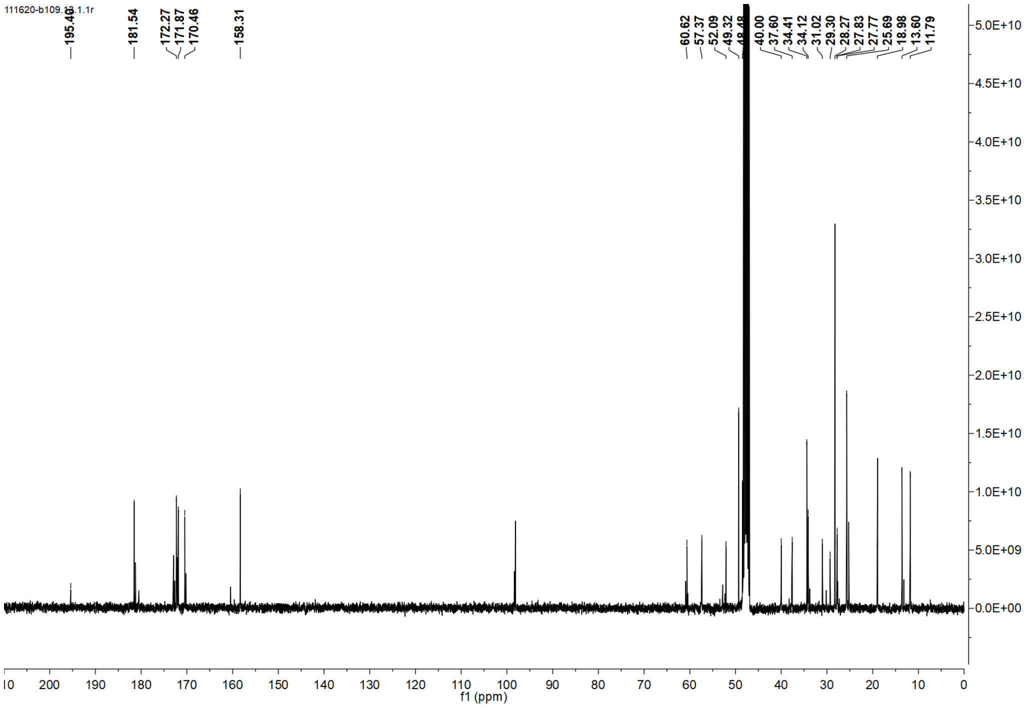

111620-b109.1.1r
181.54
172.27
171.87
170.46
158.31
60.62
57.37
52.09
49.32
40.00
37.60
34.41
34.12
31.02
29.30
28.27
27.83
27.77
25.69
18.98
13.60
11.79
f1 (ppm)

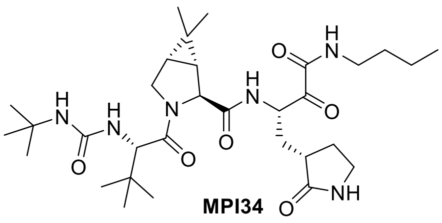

**MPI34**

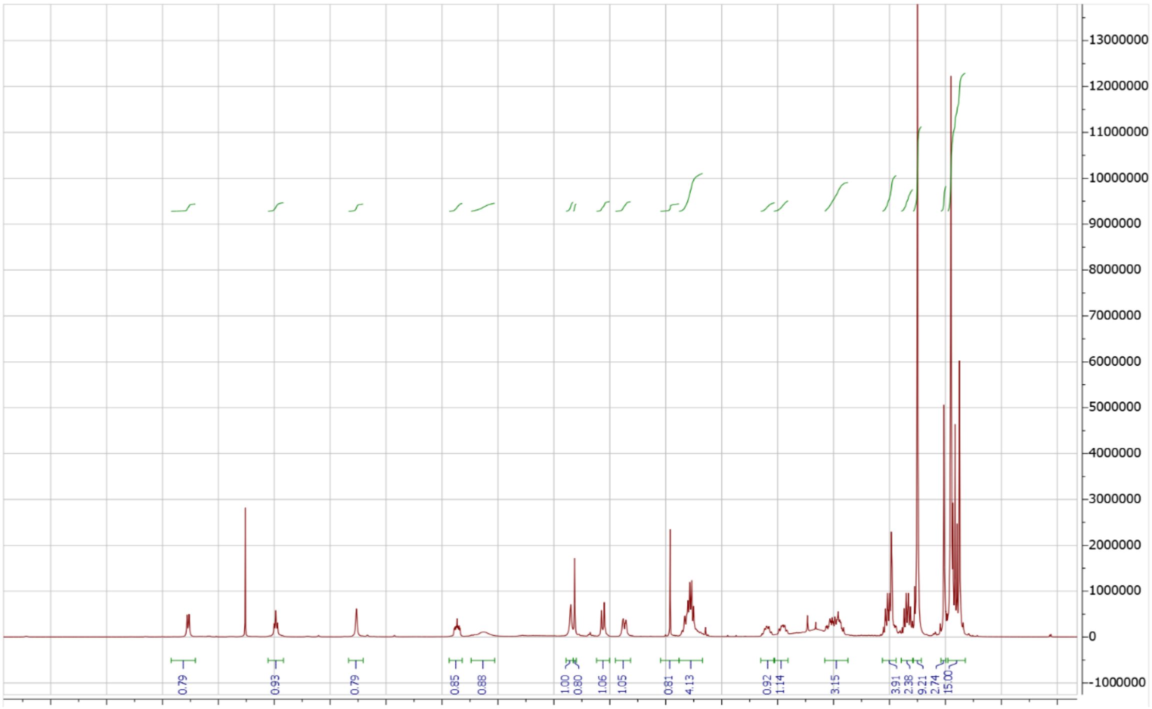

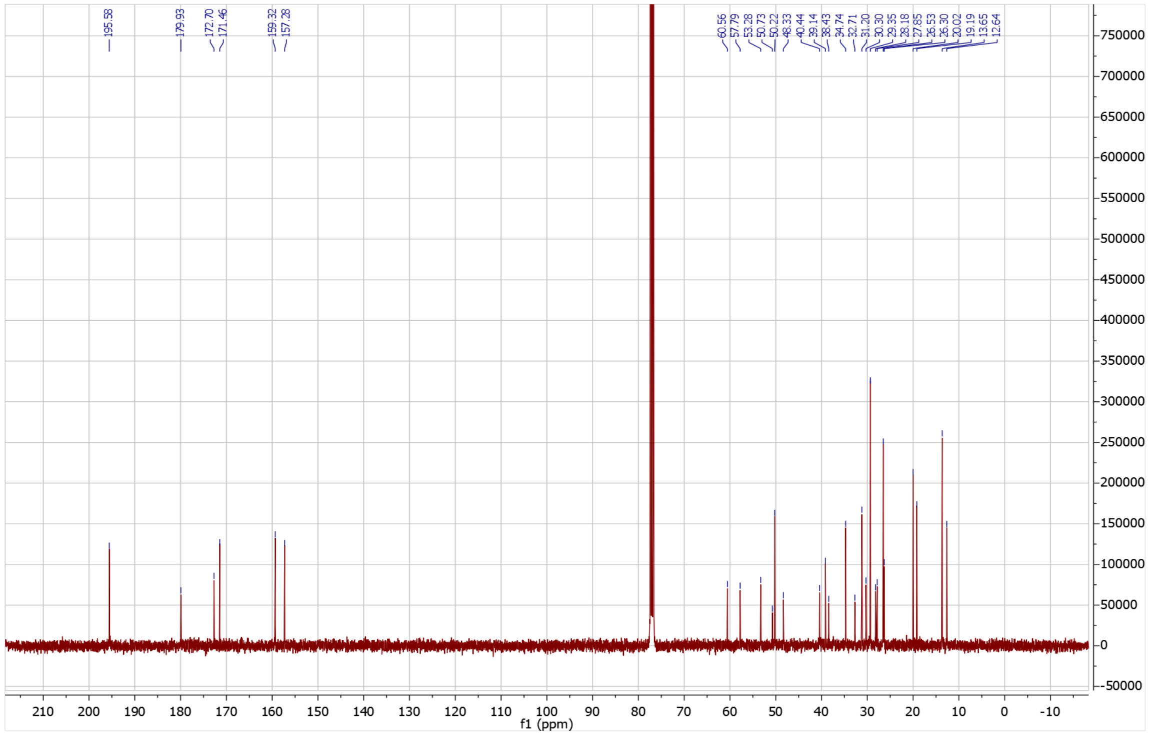

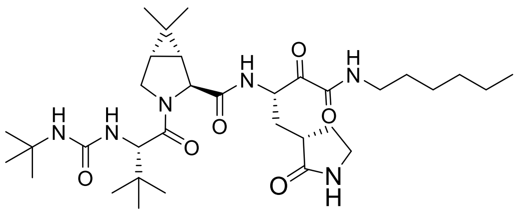

**MPI35**

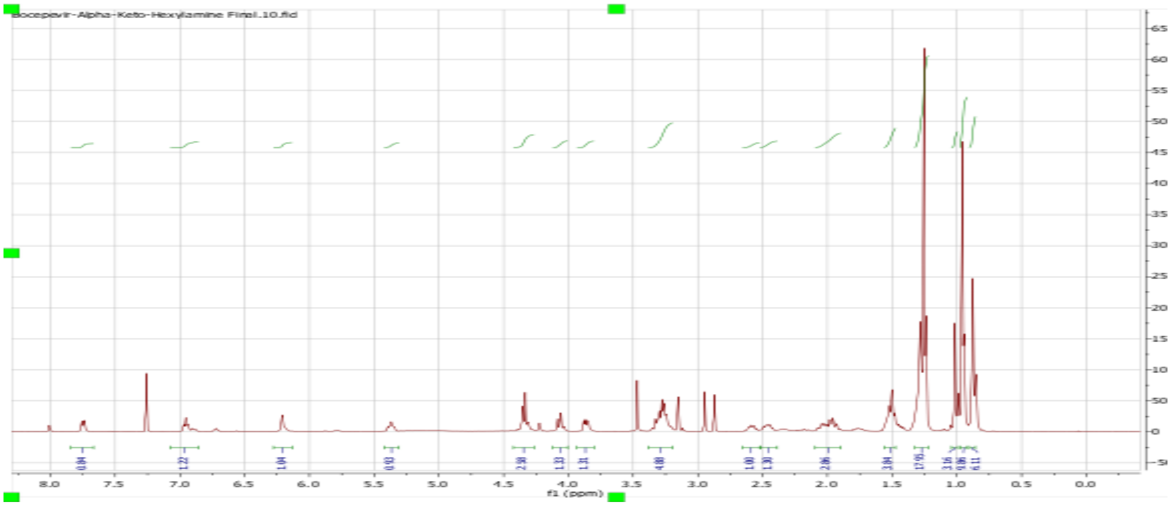
Bocepevir-Alpha-Keto-Hexylamine Final.10.fid
f1 (ppm)

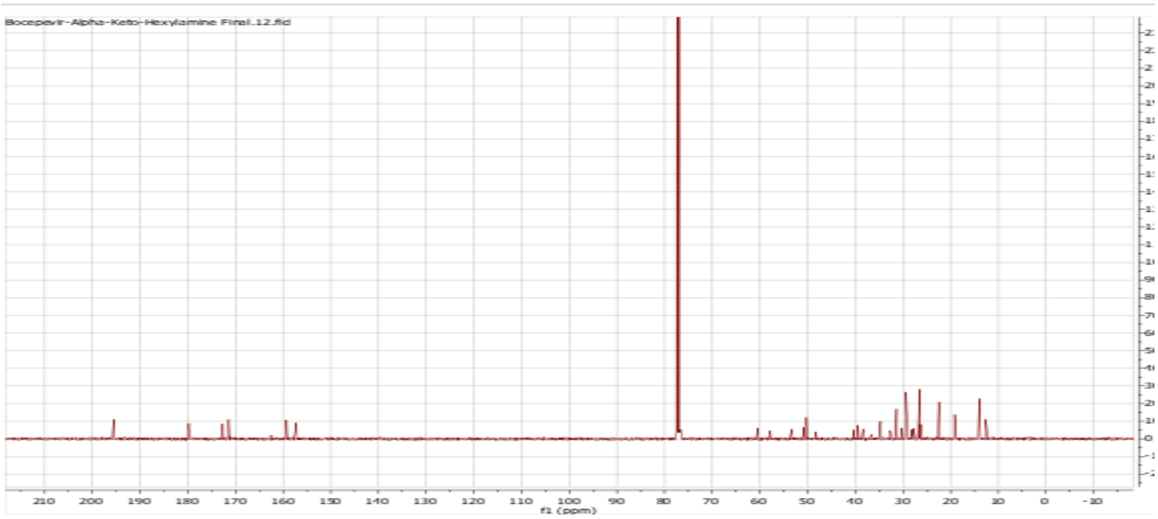
Bocepevir-Alpha-Keto-Hexylamine Final.12.fid
f1 (ppm)

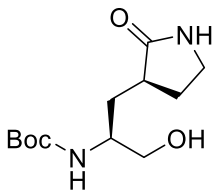

**36b**

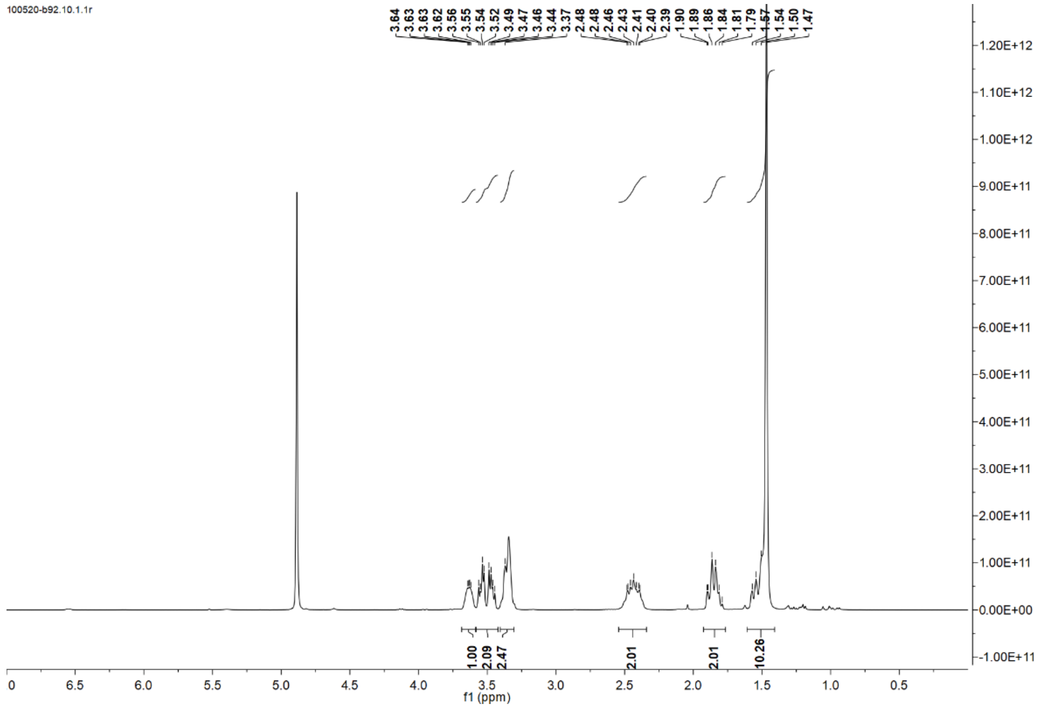
100520-b92.10.1.1r
3.64
3.63
3.63
3.62
3.56
3.55
3.54
3.52
3.49
3.47
3.46
3.44
3.37
2.48
2.48
2.46
2.43
2.41
2.40
2.39
1.90
1.89
1.86
1.84
1.81
1.79
1.57
1.54
1.50
1.47
1.00
2.09
2.47
2.01
2.01
10.26
f1 (ppm)

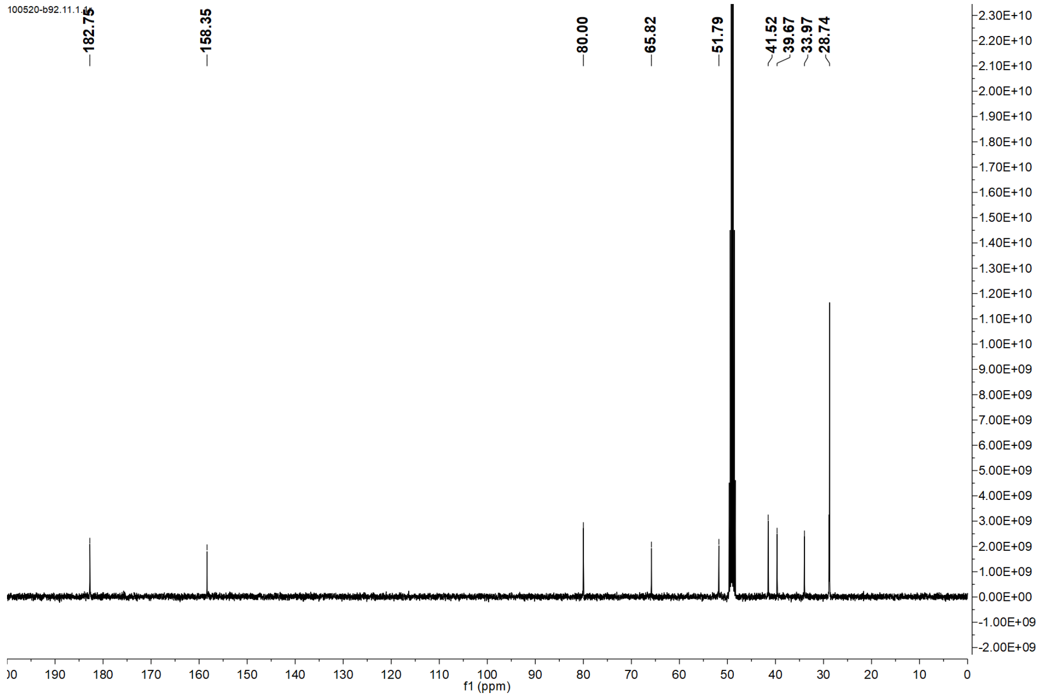
100520-b92.11.1.1r
182.75
158.35
80.00
65.82
51.79
41.52
39.67
33.97
28.74
f1 (ppm)

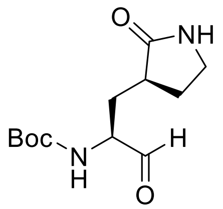
O
NH
Boc
N
H
H
O

**36c**

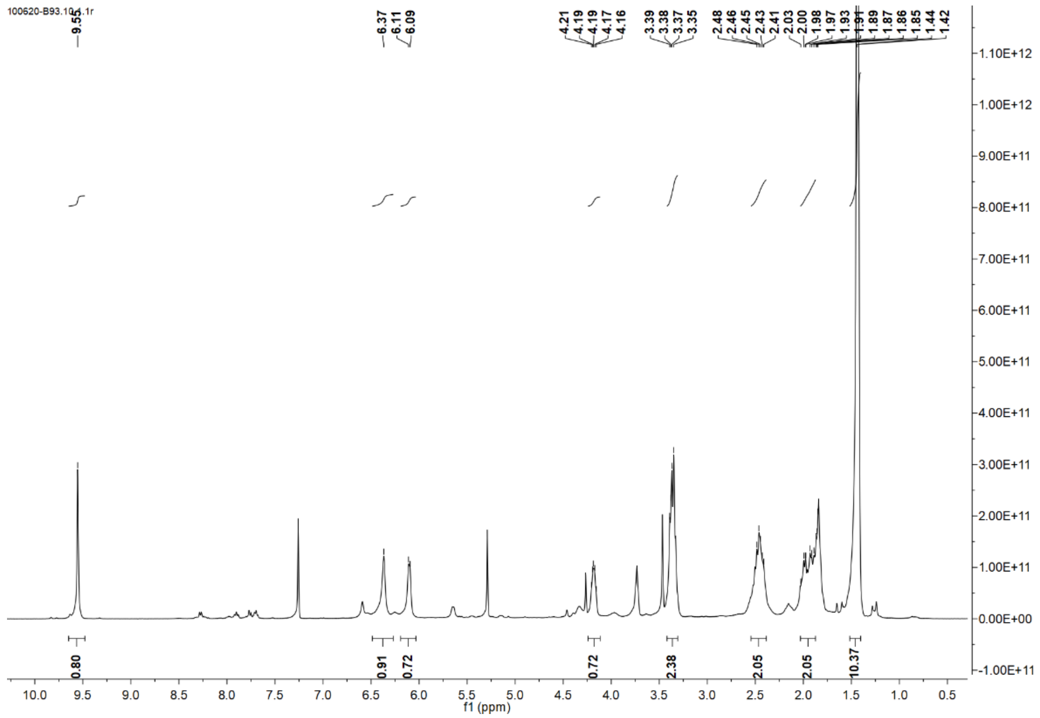
100620-B93.10.1.1r
9.55
6.37
6.11
6.09
4.21
4.19
4.19
4.17
4.16
3.39
3.38
3.37
3.35
2.48
2.46
2.45
2.43
2.41
2.03
2.00
1.98
1.97
1.93
1.91
1.89
1.87
1.86
1.85
1.44
1.42
0.80
0.91
0.72
0.72
2.38
2.05
2.05
10.37
f1 (ppm)

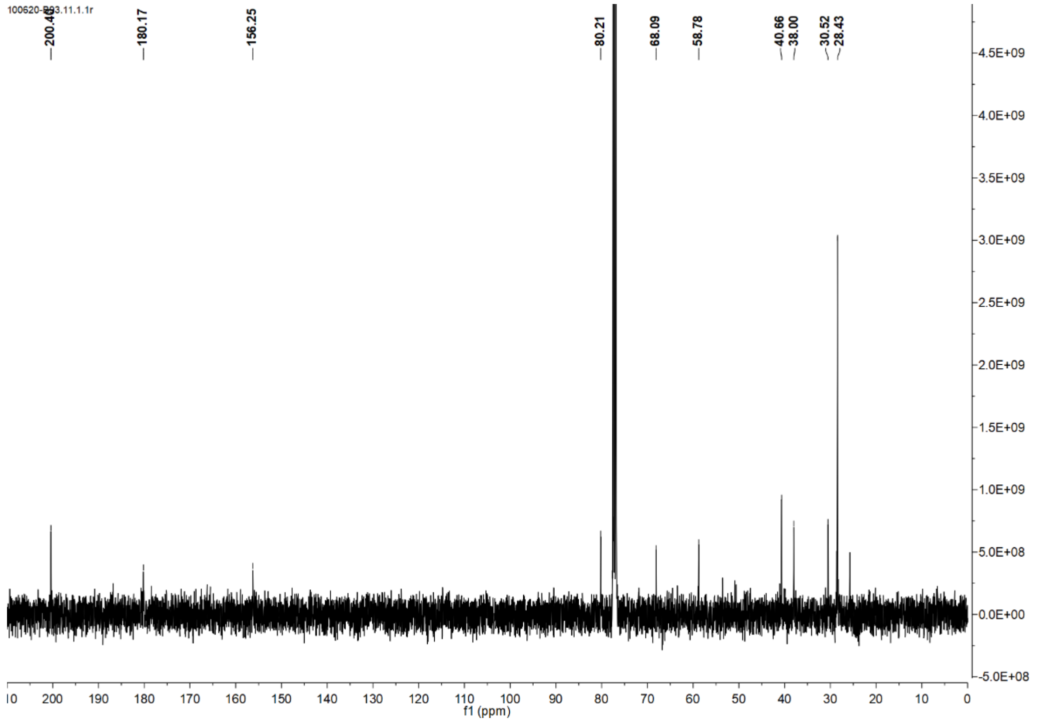
100620-B93.11.1.1r
200.40
180.17
156.25
80.21
68.09
58.78
40.66
38.00
30.52
28.43
f1 (ppm)

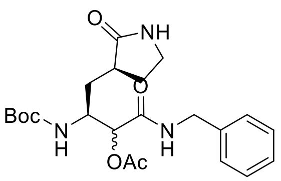
O
NH
O
Boc
N
H
OAc
N
H

**36d**

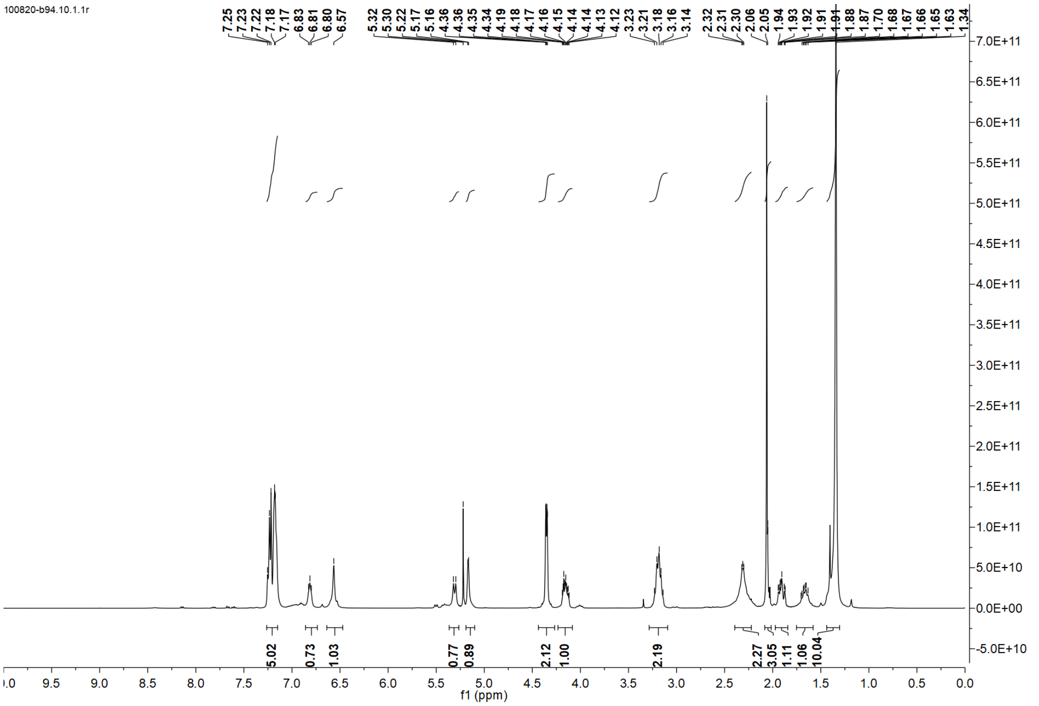
100820-b94.10.1.1r
7.25 7.23 7.22 7.18 7.17 6.83 6.81 6.80 6.57
5.32 5.30 5.22 5.17 5.16 4.36 4.36 4.35 4.34 4.19 4.18 4.17 4.16 4.15 4.14 4.14 4.13 4.12 3.23 3.21 3.18 3.16 3.14
2.32 2.31 2.30 2.06 2.05 1.94 1.93 1.92 1.91 1.91 1.88 1.87 1.70 1.68 1.67 1.66 1.65 1.63 1.34
5.02 0.73 1.03 0.77 0.89 2.12 1.00 2.19 2.27 3.05 1.11 1.06 10.04
f1 (ppm)

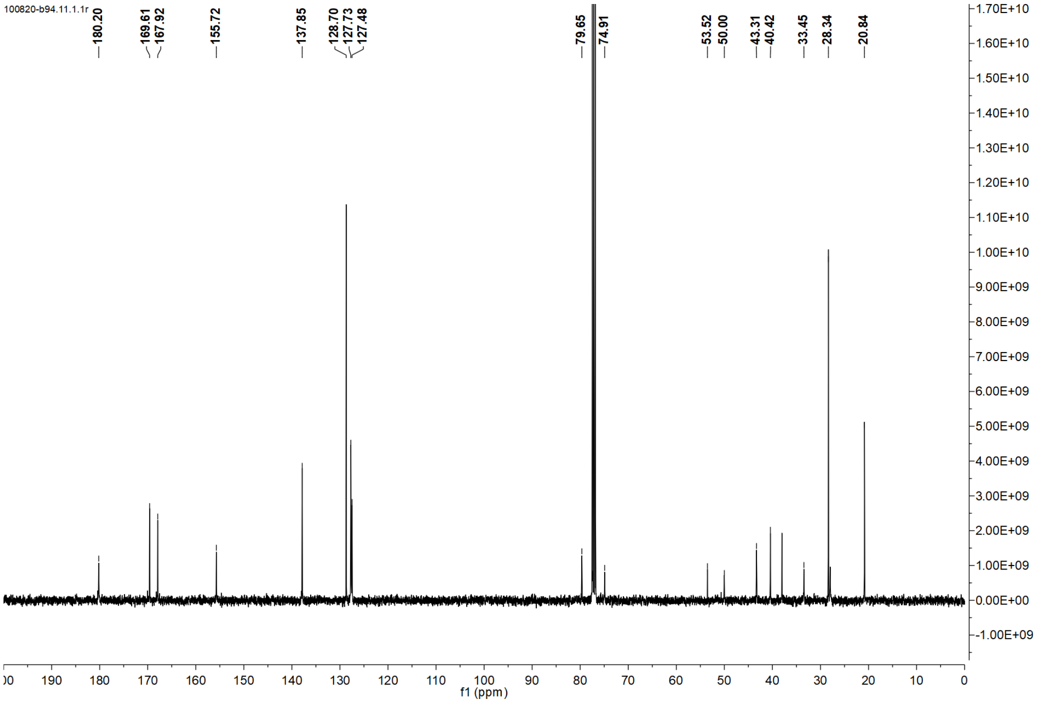
100820-b94.11.1.1r
180.20
169.61 167.92
155.72
137.85
128.70 127.73 127.48
79.65 74.91
53.52 50.00
43.31 40.42
33.45
28.34
20.84
f1 (ppm)

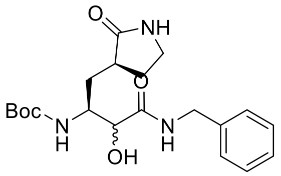

O
NH
O
Boc
N
H
OH
N
H


**36e**

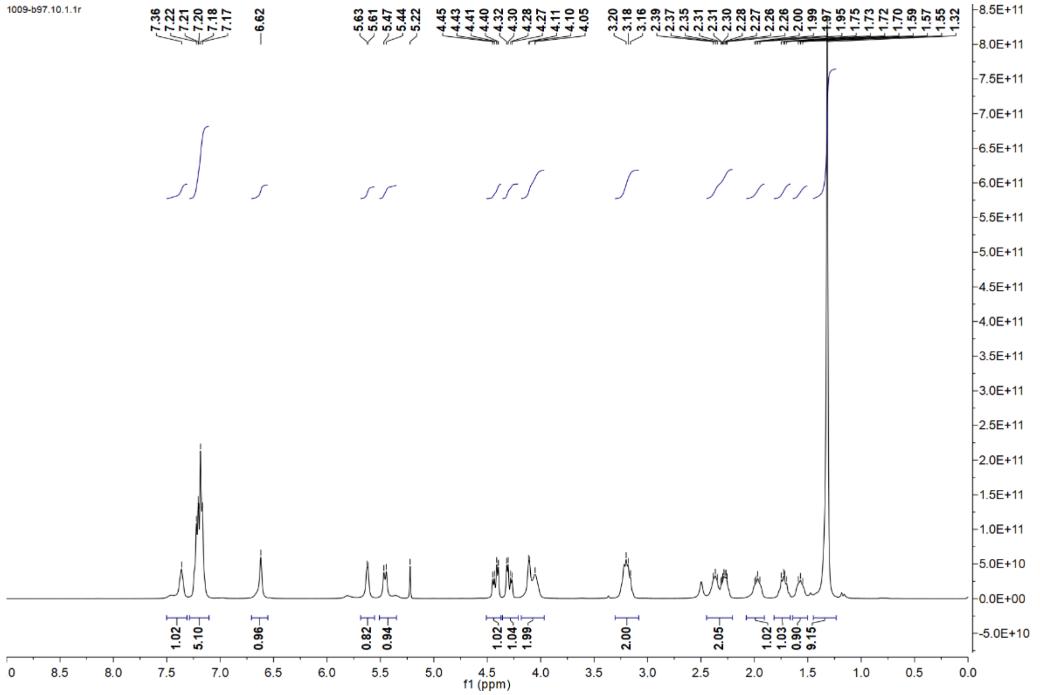

1009-b97.10.1.1r
7.36
7.22
7.21
7.20
7.18
7.17
6.62
5.63
5.61
5.47
5.44
5.22
4.45
4.43
4.41
4.40
4.32
4.30
4.28
4.27
4.11
4.10
4.05
3.20
3.18
3.16
2.39
2.37
2.35
2.31
2.31
2.30
2.28
2.27
2.26
2.26
2.00
1.99
1.97
1.95
1.75
1.73
1.72
1.70
1.59
1.57
1.55
1.32
1.02
5.10
0.96
0.82
0.94
1.02
1.04
1.99
2.00
2.05
1.02
1.03
0.90
9.15
f1 (ppm)


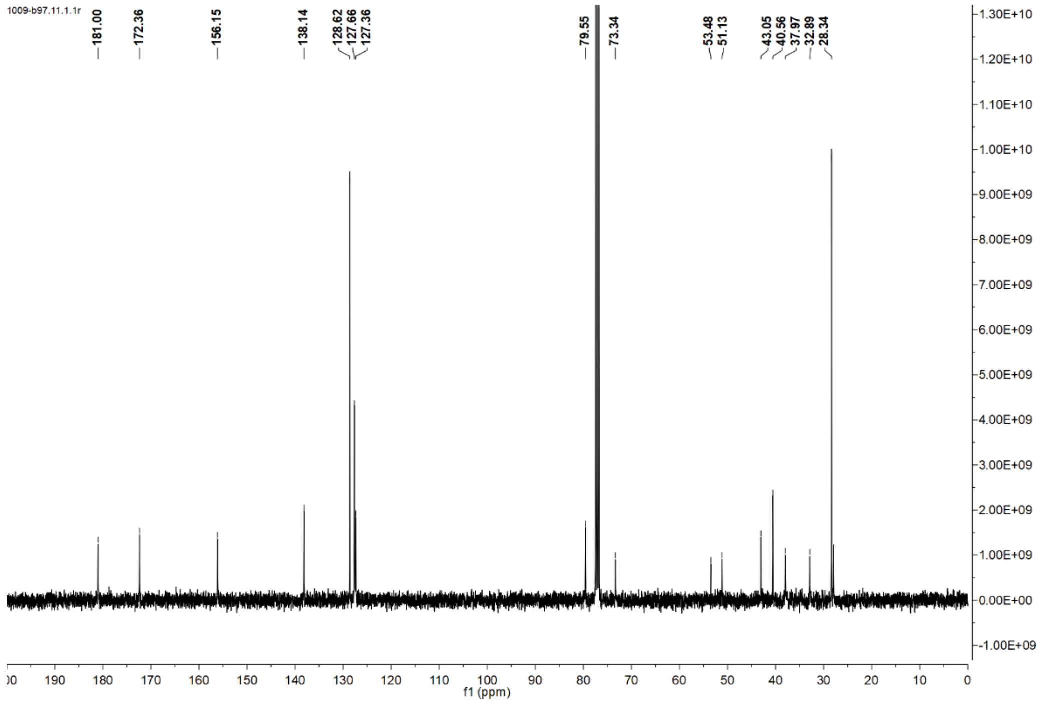

1009-b97.11.1.1r
181.00
172.36
156.15
138.14
128.62
127.66
127.36
79.55
73.34
53.48
51.13
43.05
40.56
37.97
32.89
28.34
f1 (ppm)

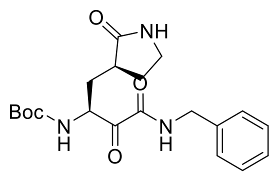

**36f**

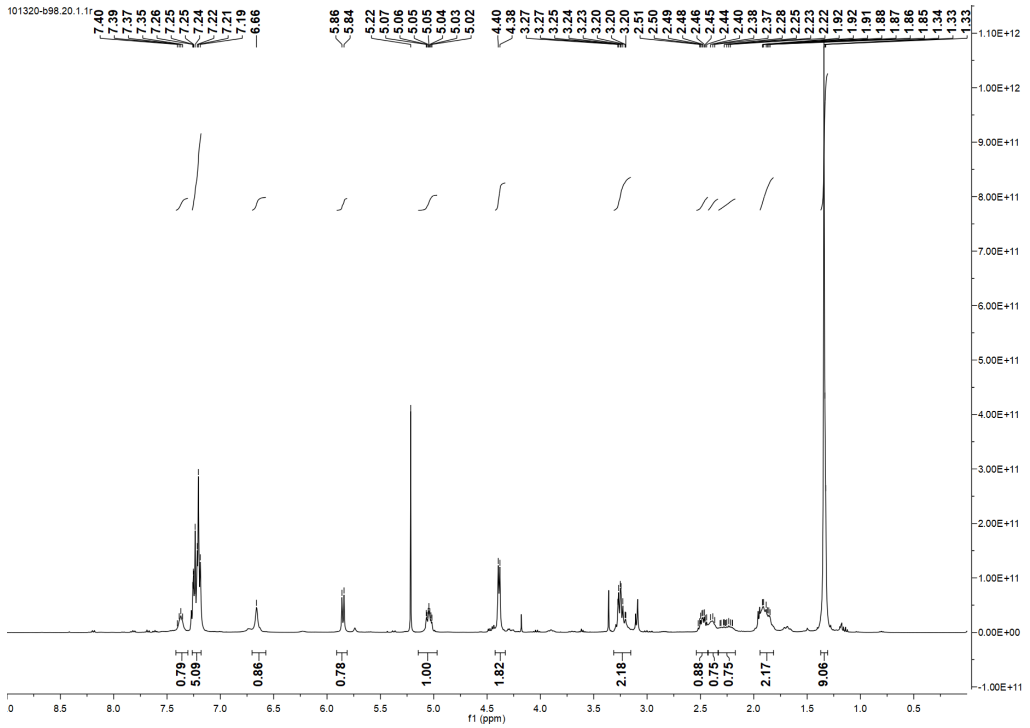
101320-b98.20.1.1r
7.40 7.39 7.37 7.35 7.26 7.25 7.25 7.24 7.22 7.21 7.19 6.66 5.86 5.84 5.22 5.07 5.06 5.05 5.05 5.04 5.03 5.02 4.40 4.38 3.27 3.25 3.24 3.23 3.20 3.20 3.20 2.51 2.50 2.49 2.48 2.46 2.45 2.44 2.40 2.38 2.37 2.28 2.25 2.23 2.22 1.92 1.92 1.91 1.88 1.87 1.86 1.85 1.34 1.33 1.33
0.79 5.09 0.86 0.78 1.00 1.82 2.18 0.88 0.75 0.75 2.17 9.06
f1 (ppm)

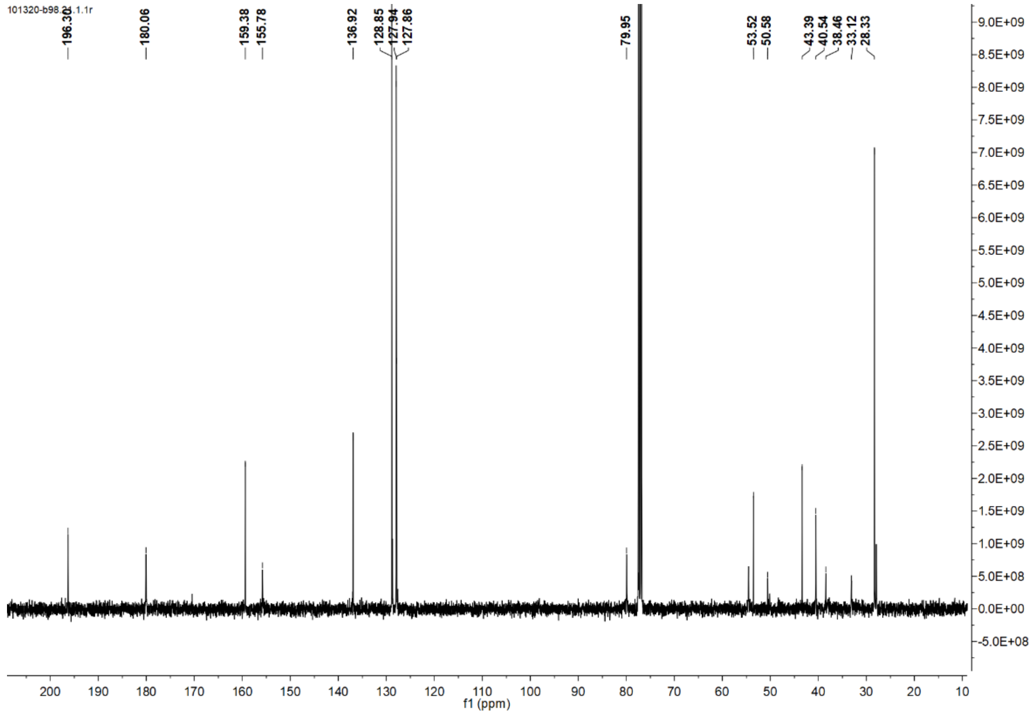
196.30 180.06 159.38 155.78 136.92 128.85 127.94 127.86 79.95 53.52 50.58 43.39 40.54 38.46 33.12 28.33
f1 (ppm)

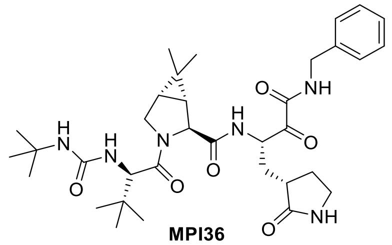

**MPI36**

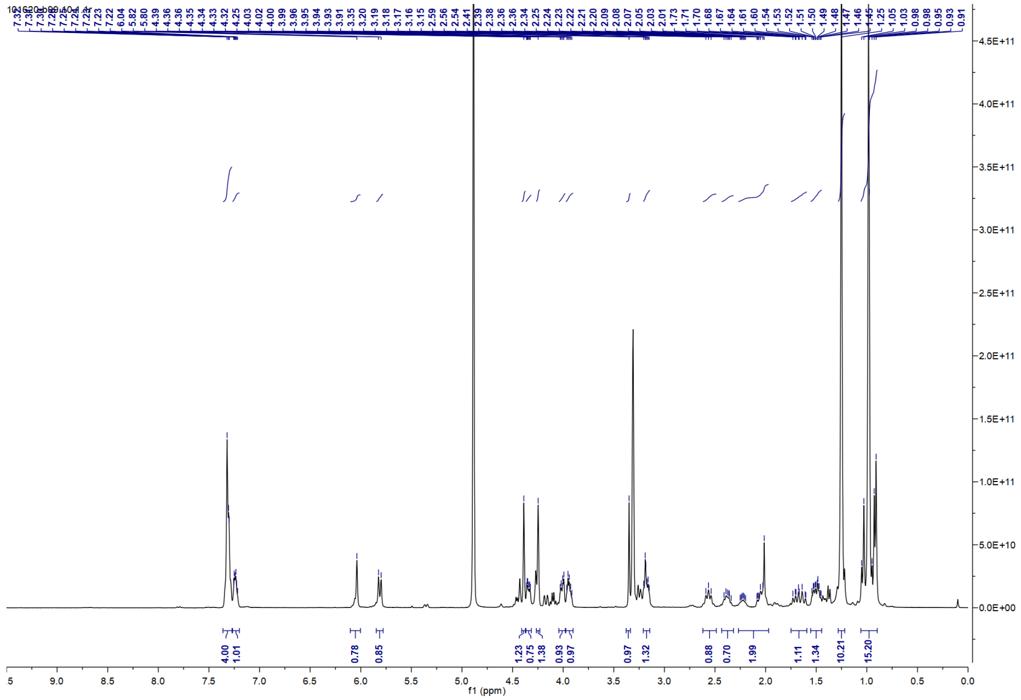

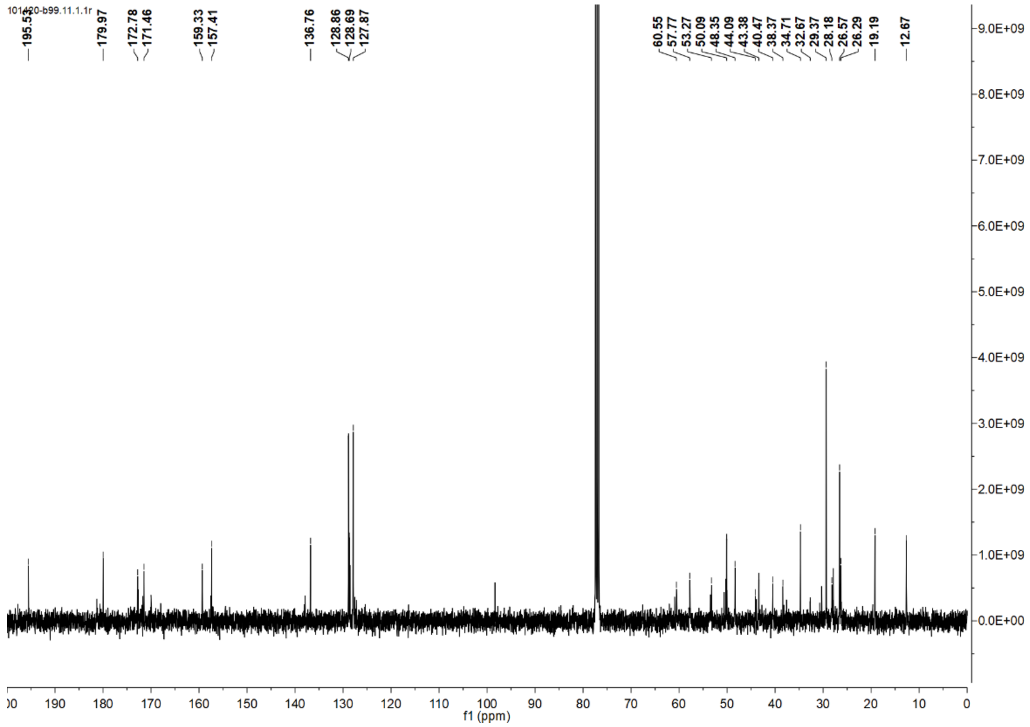

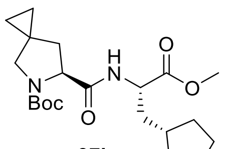

**37b**

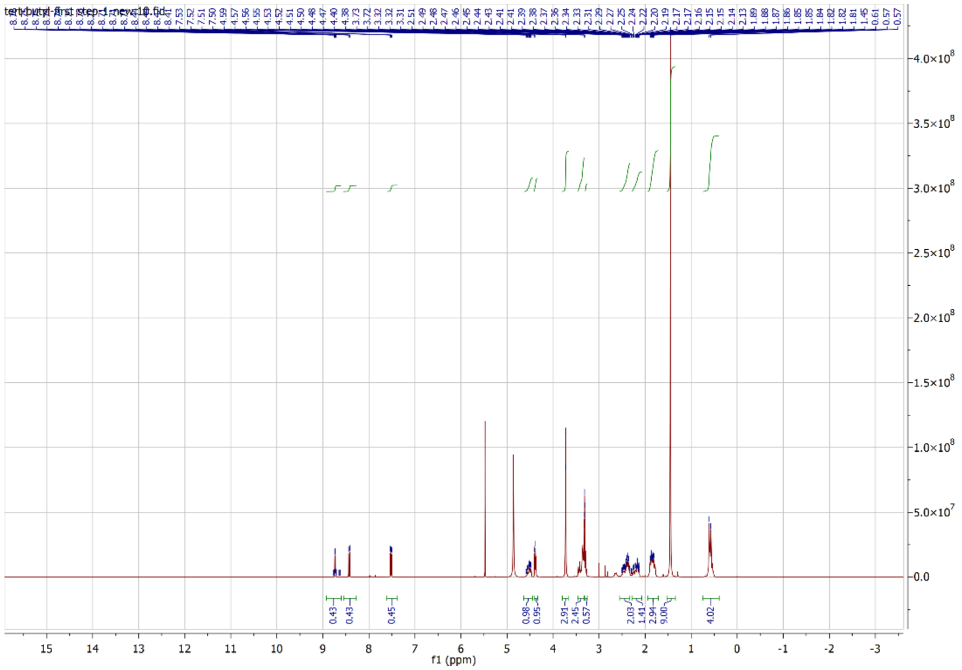

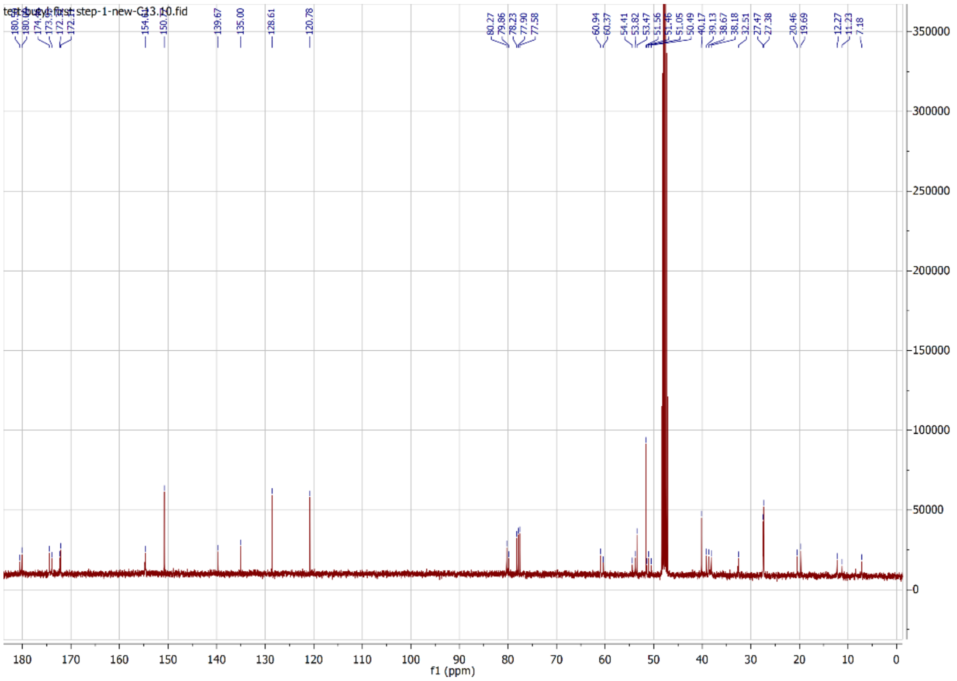

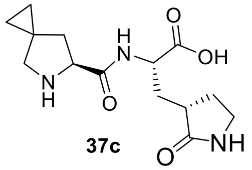

**37c**

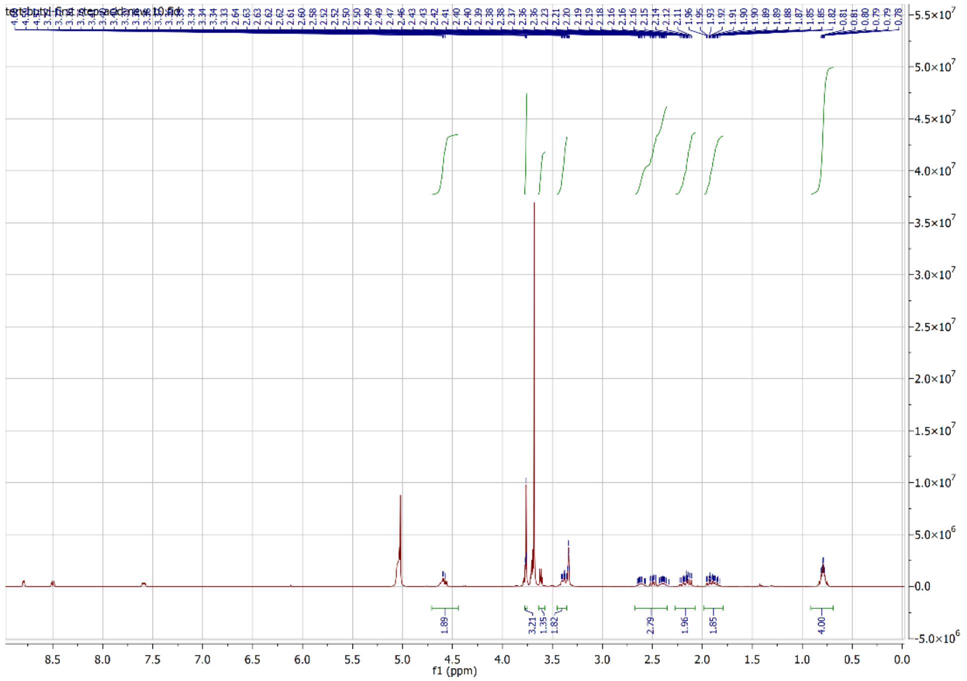


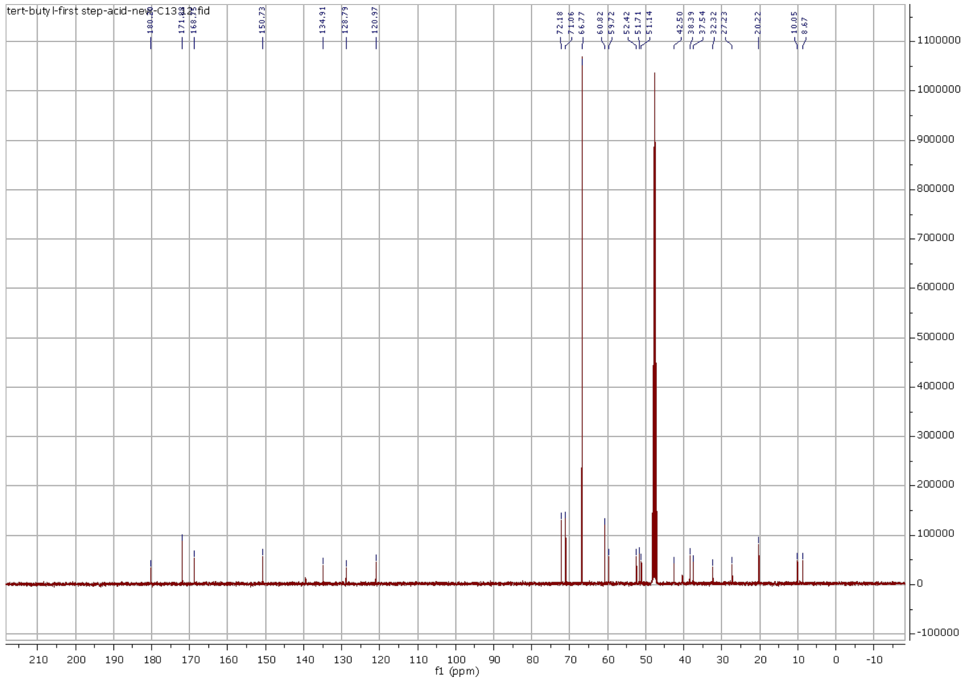

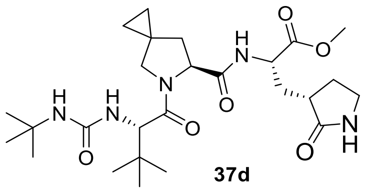
H
N
H
N
O
N
O
H
N
O
O
O
N
H
O

**37d**

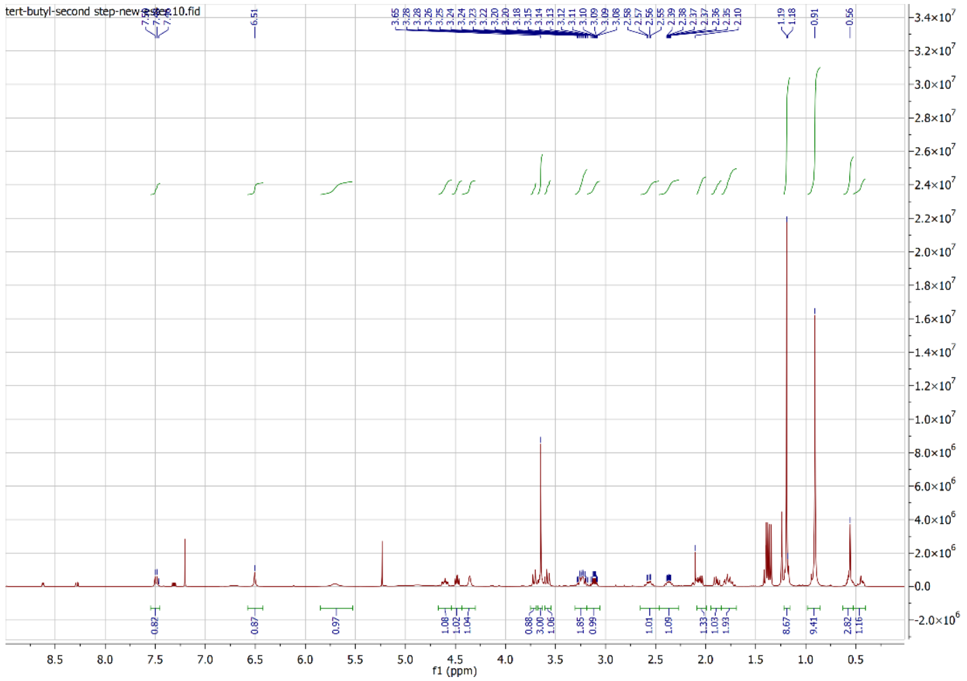
tert-butyl-second step-new ester10.fid
f1 (ppm)

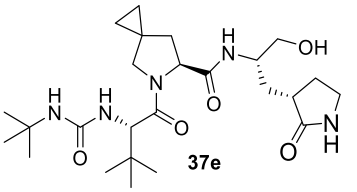

OH
N
H
O
37e


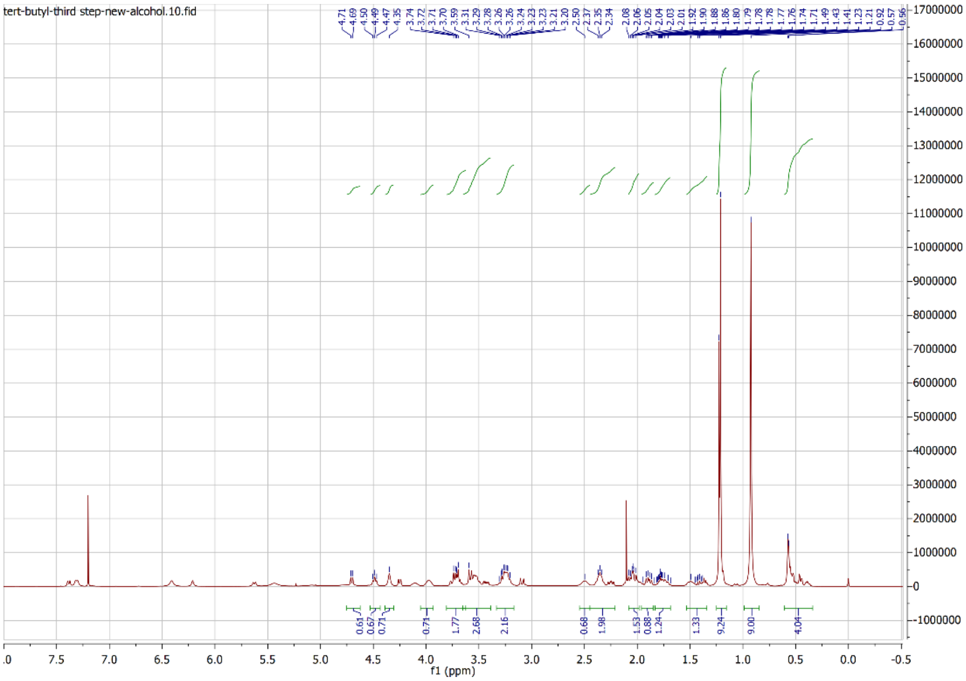

tert-butyl-third step-new-alcohol.10.fid
f1 (ppm)

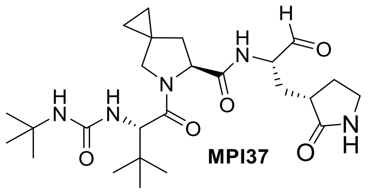
MPI37

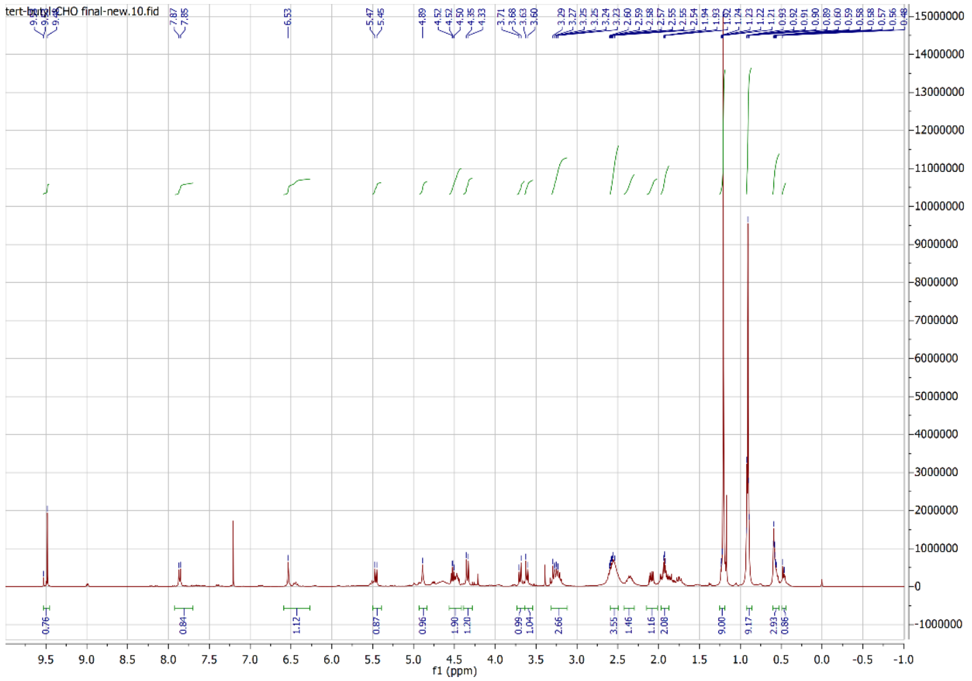
tert-butyl-CHO final-new.10.fid
f1 (ppm)

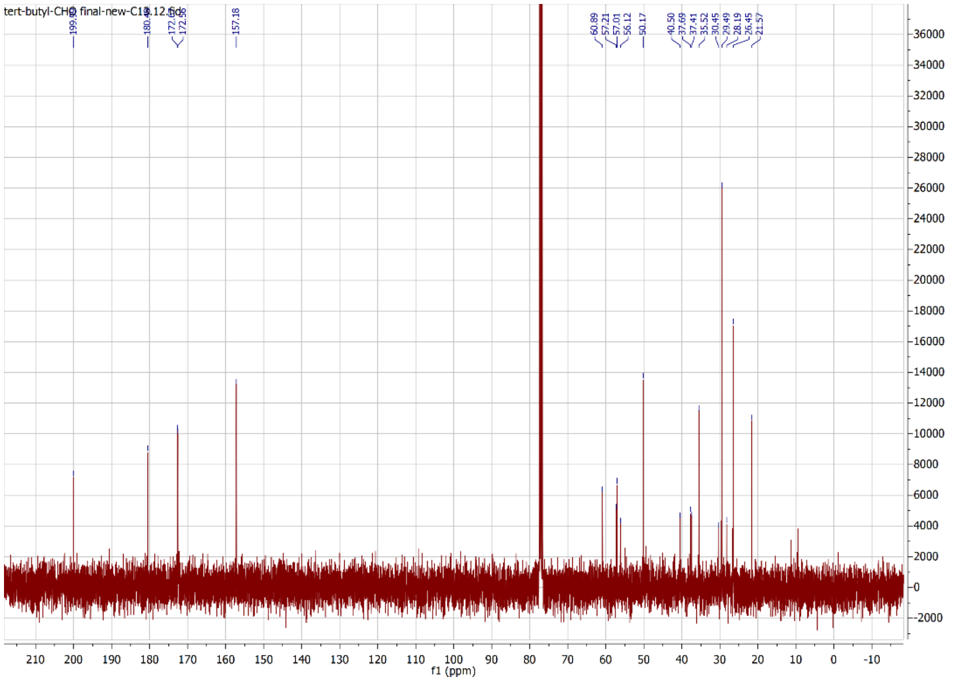
tert-butyl-CHO final-new-C13.12.fid
199.92
180.18
172.69
172.55
157.18
60.89
57.21
57.01
56.12
50.17
40.50
37.69
37.41
35.52
30.45
29.49
28.19
26.45
21.57
f1 (ppm)

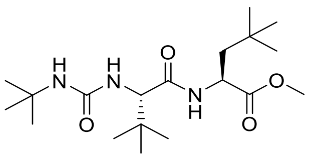

38b

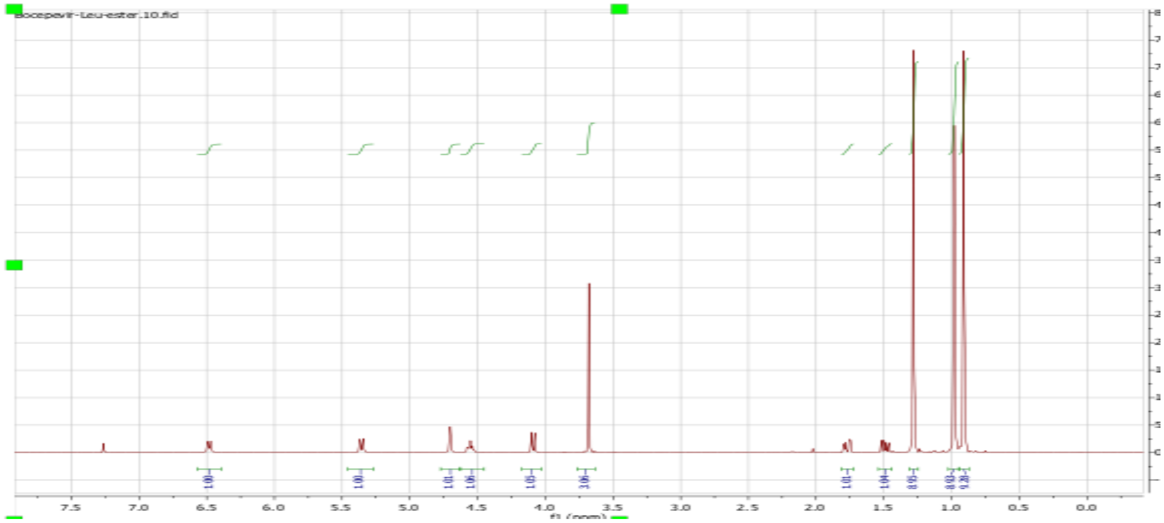
Bocepevir-Leu-ester.10.fid
f1 (ppm)

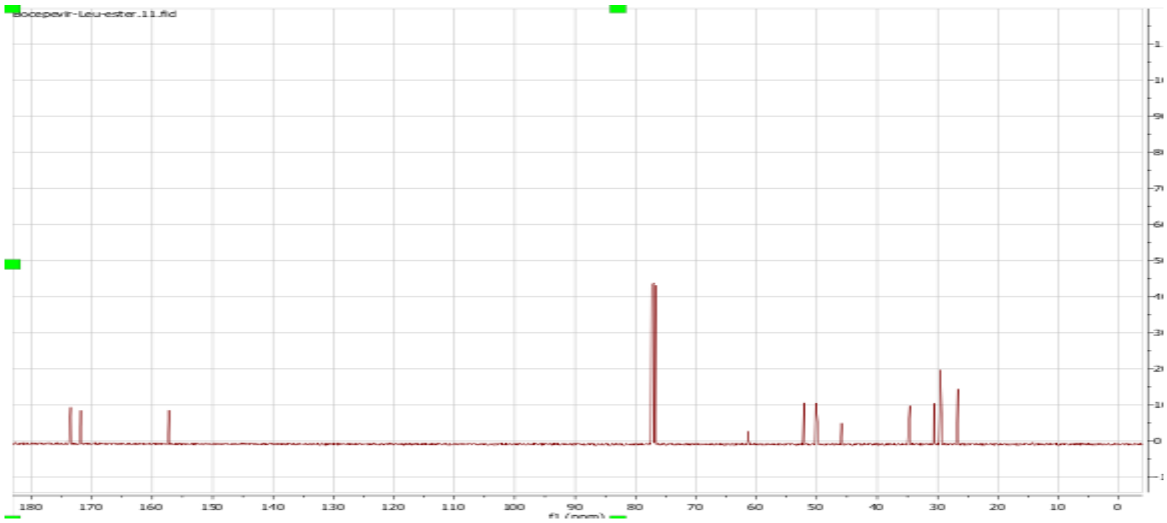
Bocepevir-Leu-ester.11.fid
f1 (ppm)

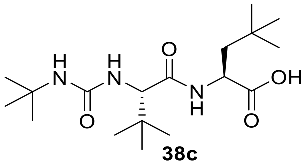

**38c**

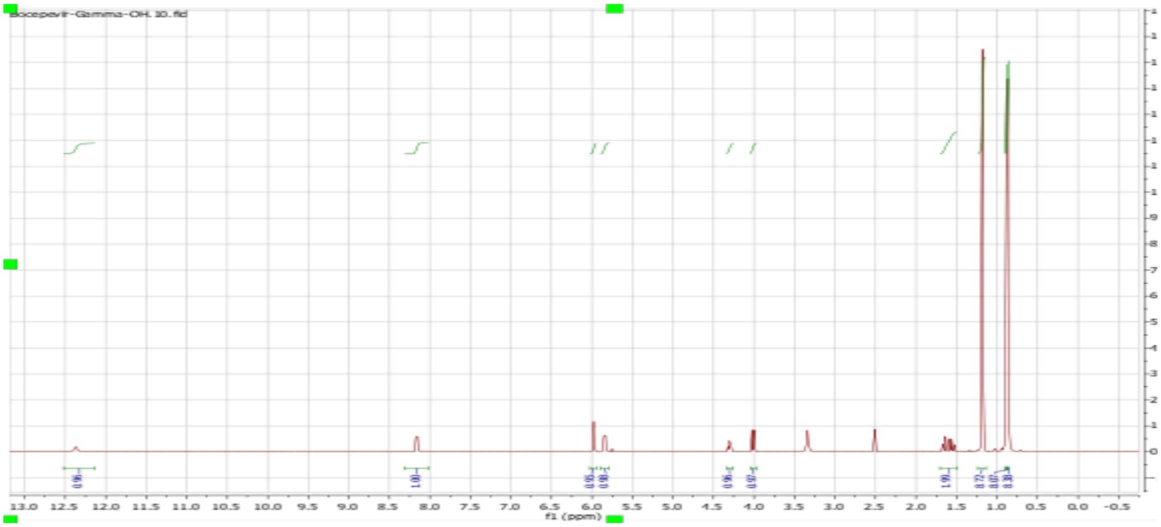
Boceprevir-Gamma-OH. 10.fid
13.0 12.5 12.0 11.5 11.0 10.5 10.0 9.5 9.0 8.5 8.0 7.5 7.0 6.5 6.0 5.5 5.0 4.5 4.0 3.5 3.0 2.5 2.0 1.5 1.0 0.5 0.0 -0.5
f1 (ppm)

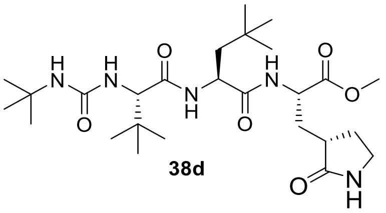

**38d**

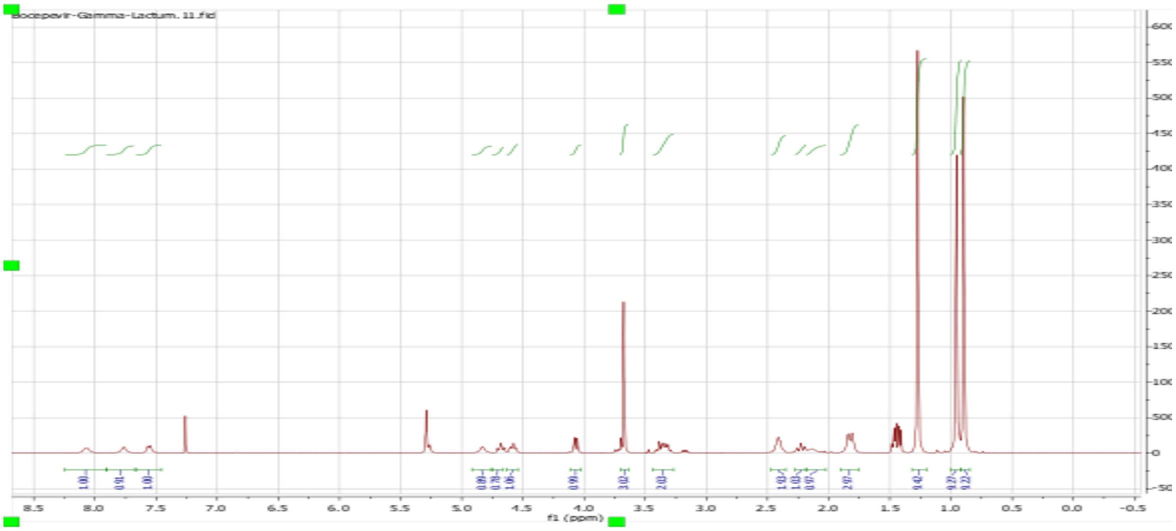

Bocepevir-Gamma-Lactum.11.fid
f1 (ppm)


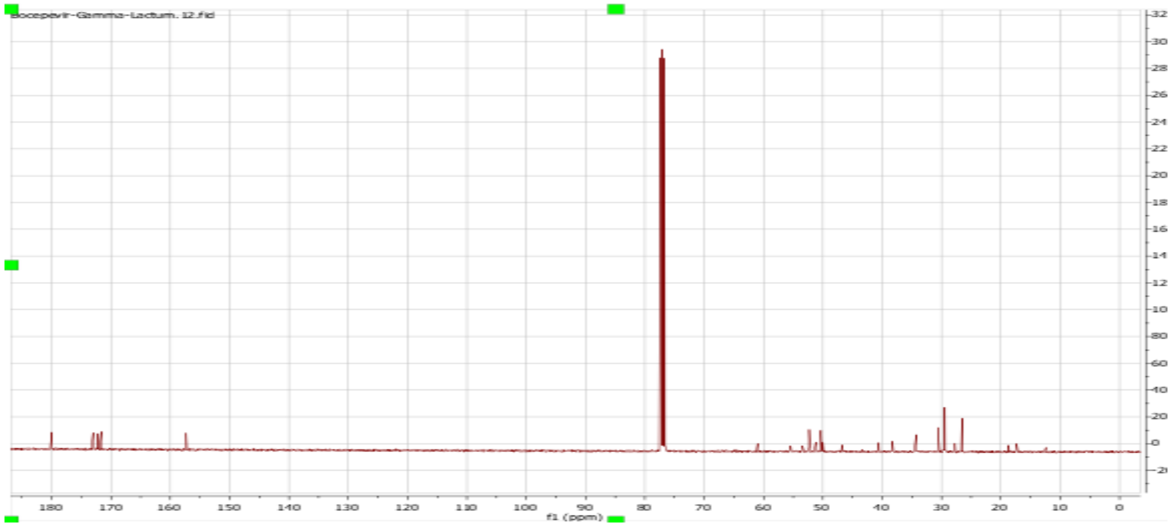

Bocepevir-Gamma-Lactum.12.fid
f1 (ppm)

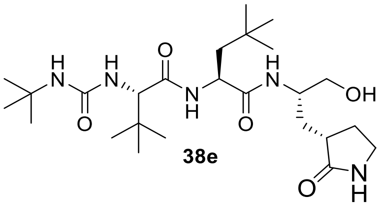

H
N
O
OH
NH


**38e**

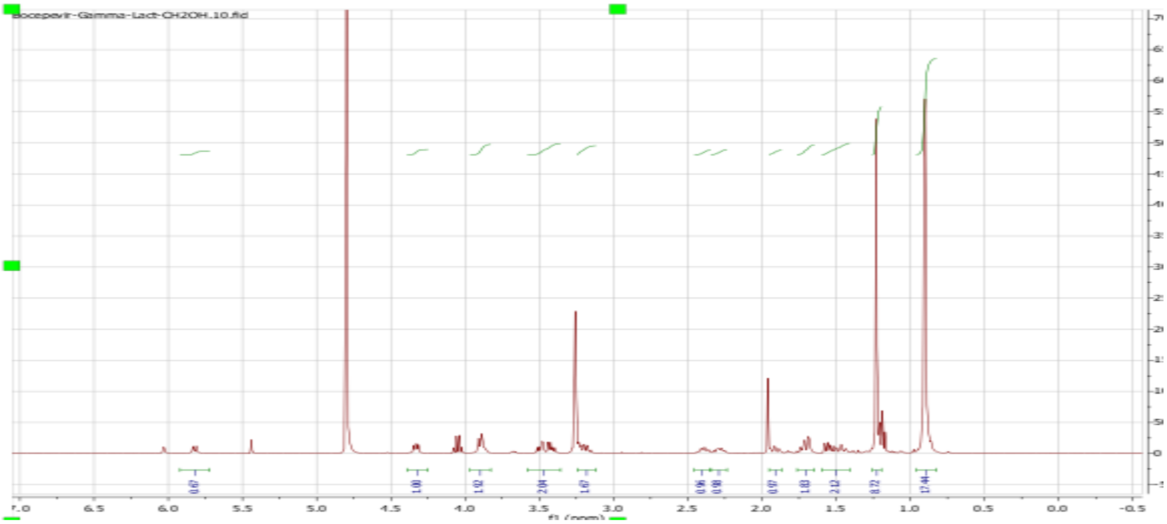

Bocepevir-Gamma-Lact-CH2OH.10.fid
0.67
1.00
1.92
2.04
1.67
0.96
0.98
0.97
1.03
2.12
8.72
17.44
6.5
6.0
5.5
5.0
4.5
4.0
3.5
3.0
2.5
2.0
1.5
1.0
0.5
0.0
-0.5

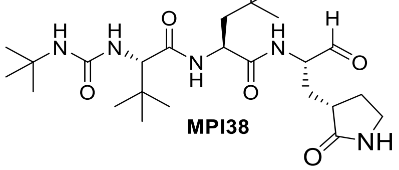
H
N
O
NH
O
MPI38

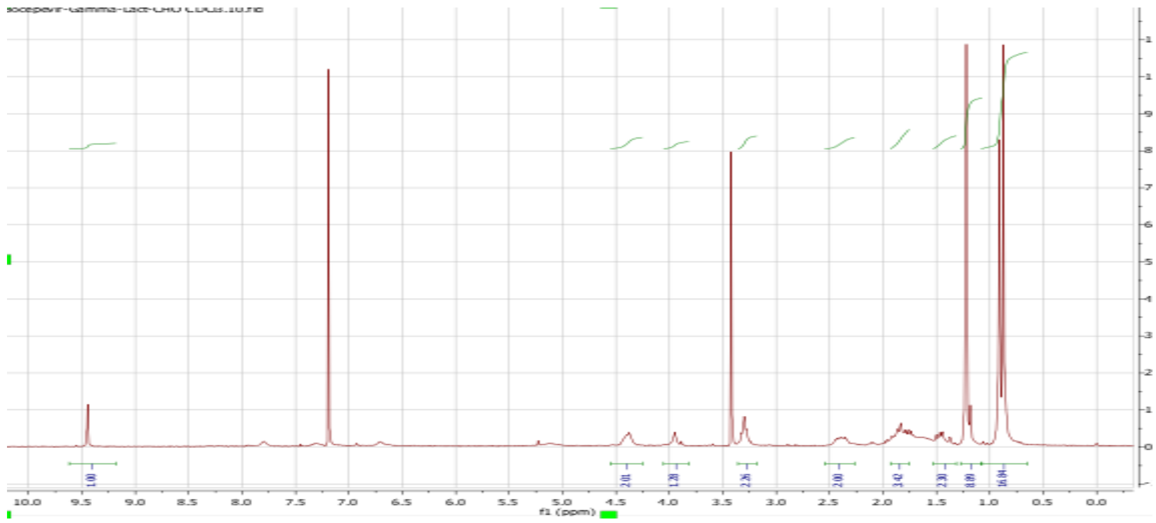
f1 (ppm)

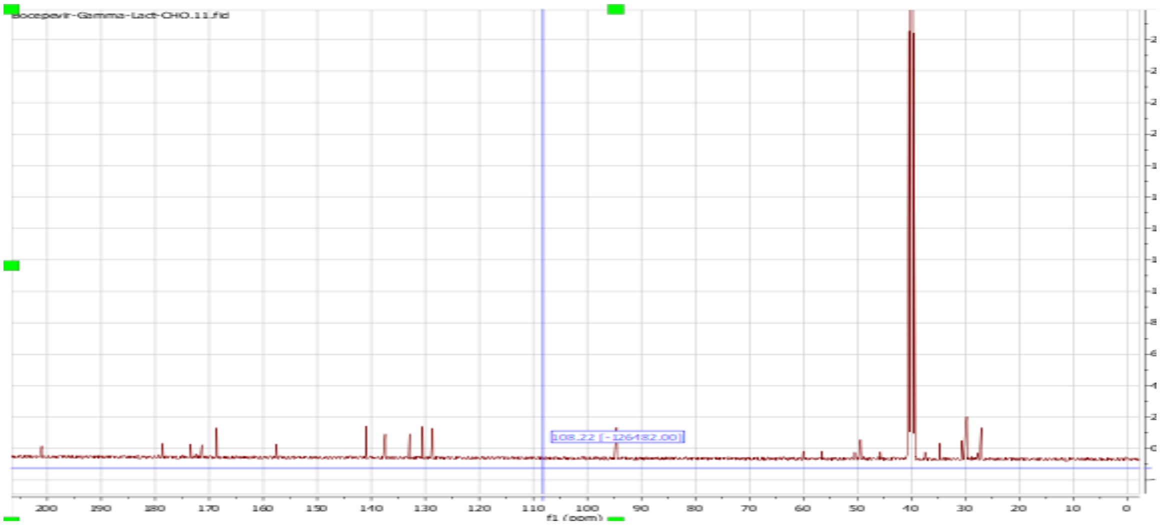
f1 (ppm)

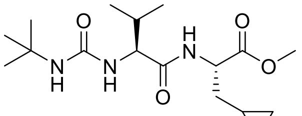

O
N
H
N
H
N
H
O
O
O


**39b**

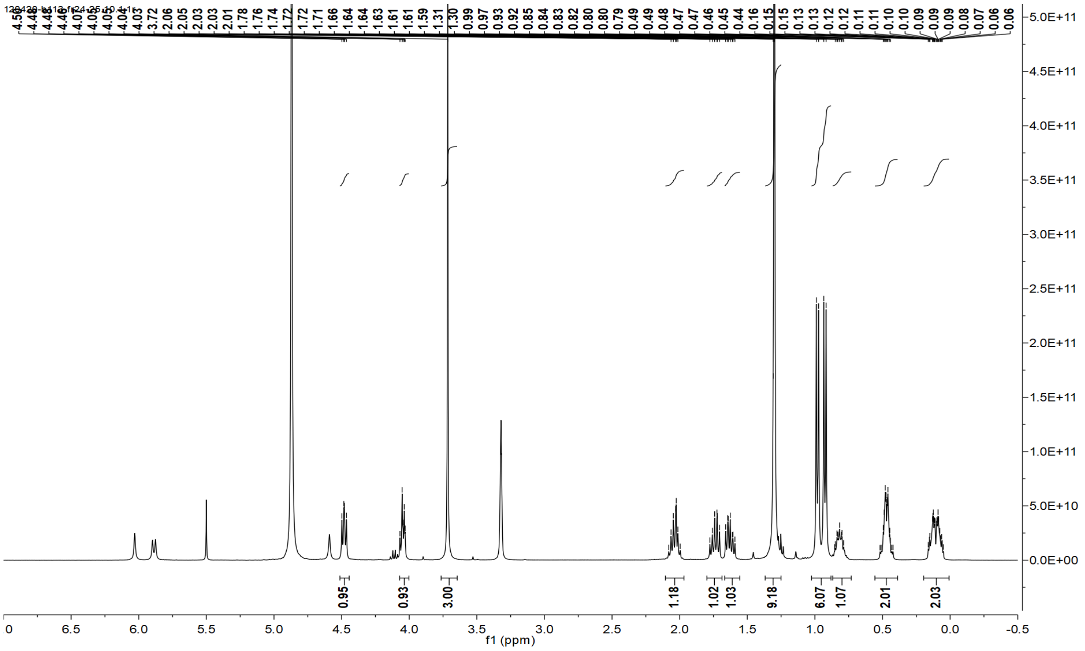

0.95
0.93
3.00
1.18
1.02
1.03
9.18
6.07
1.07
2.01
2.03
f1 (ppm)


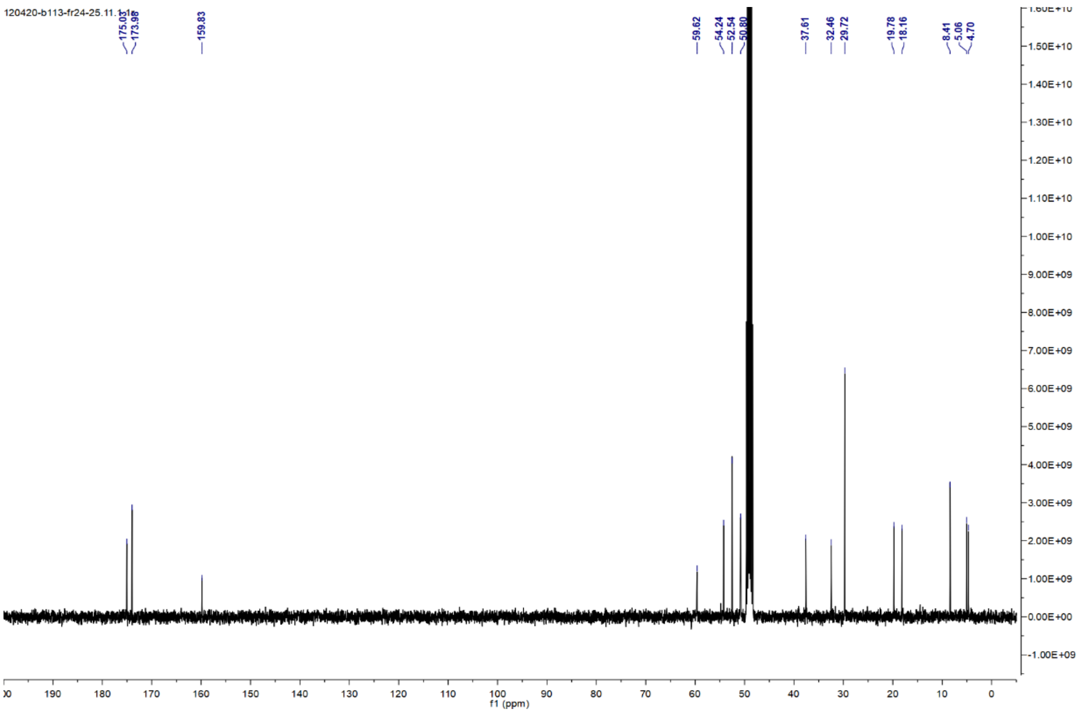

120420-b113-fr24-25.11.1.1r
175.01
159.83
59.62
54.24
52.54
50.80
37.61
32.46
29.72
19.78
18.16
8.41
5.06
4.70
f1 (ppm)

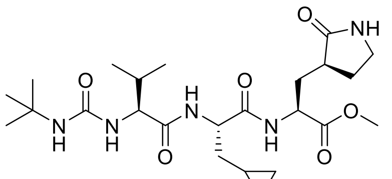
O
NH
N
H
O
N
H
N
H
O
O
O
N
H
O

**39d**

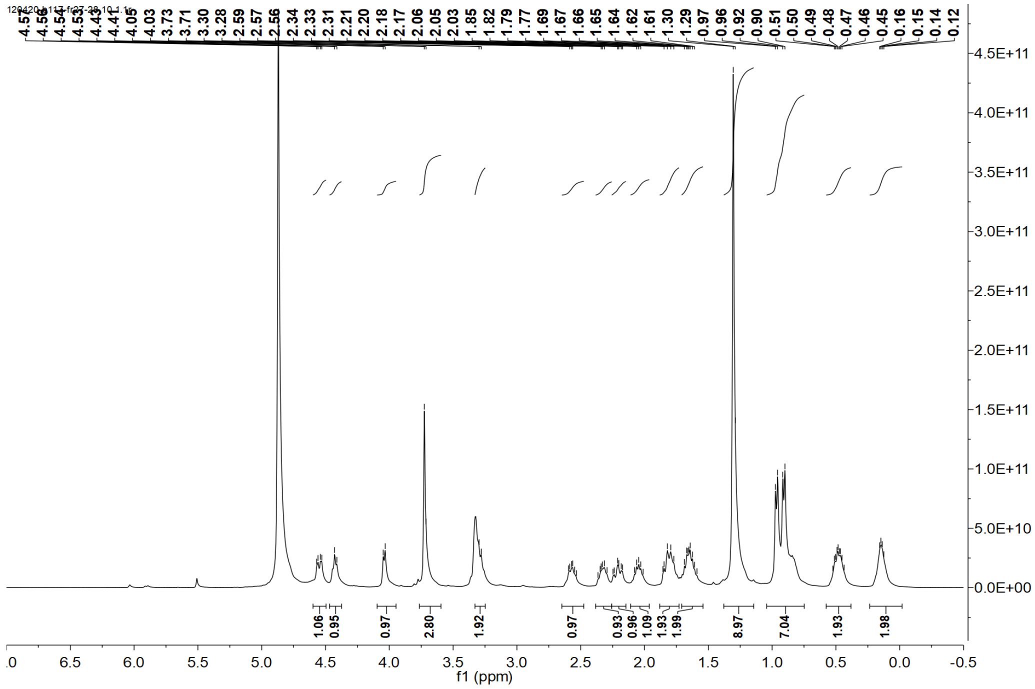
4.56
4.54
4.53
4.43
4.41
4.05
4.03
3.73
3.71
3.30
3.28
2.59
2.57
2.56
2.34
2.33
2.31
2.21
2.20
2.18
2.17
2.06
2.05
2.03
1.85
1.82
1.79
1.77
1.69
1.67
1.66
1.65
1.64
1.62
1.61
1.30
1.29
0.97
0.96
0.92
0.90
0.51
0.50
0.49
0.48
0.47
0.46
0.45
0.16
0.15
0.14
0.12
1.06
0.95
0.97
2.80
1.92
0.97
0.93
0.96
1.09
1.93
1.99
8.97
7.04
1.93
1.98
f1 (ppm)

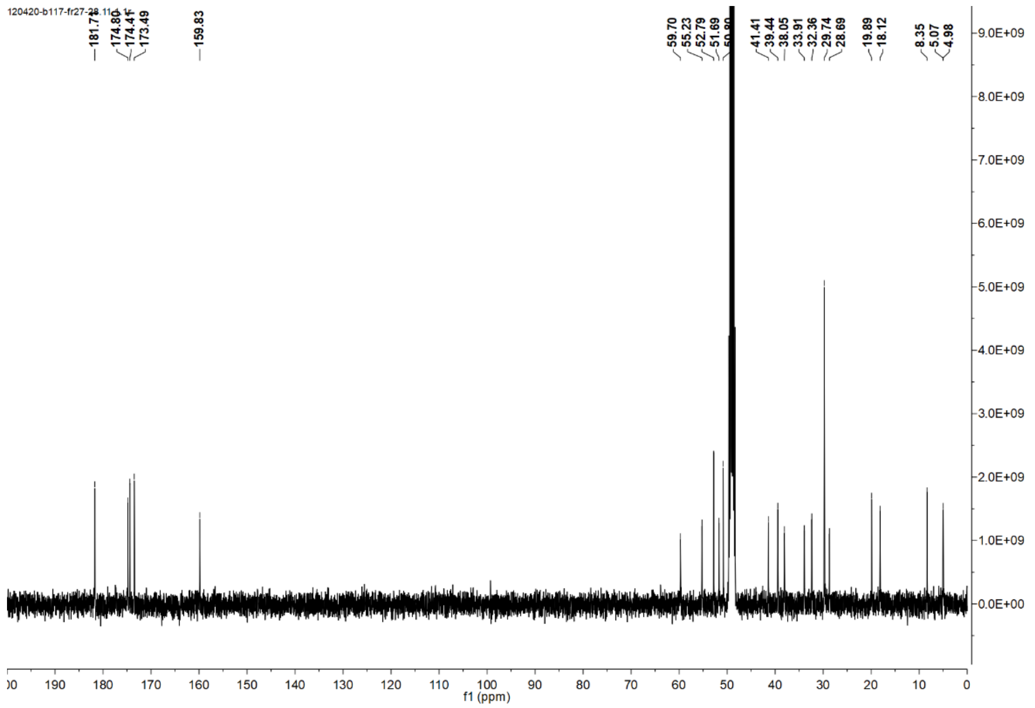
174.86
174.41
173.49
159.83
59.70
55.23
52.79
51.69
41.41
39.44
38.05
33.91
32.36
29.74
28.69
19.89
18.12
8.35
5.07
4.98
f1 (ppm)

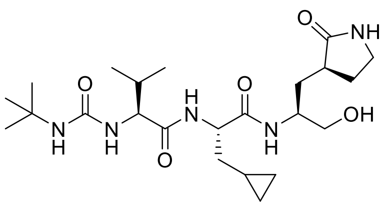
O
NH
N
H
OH

**39e**

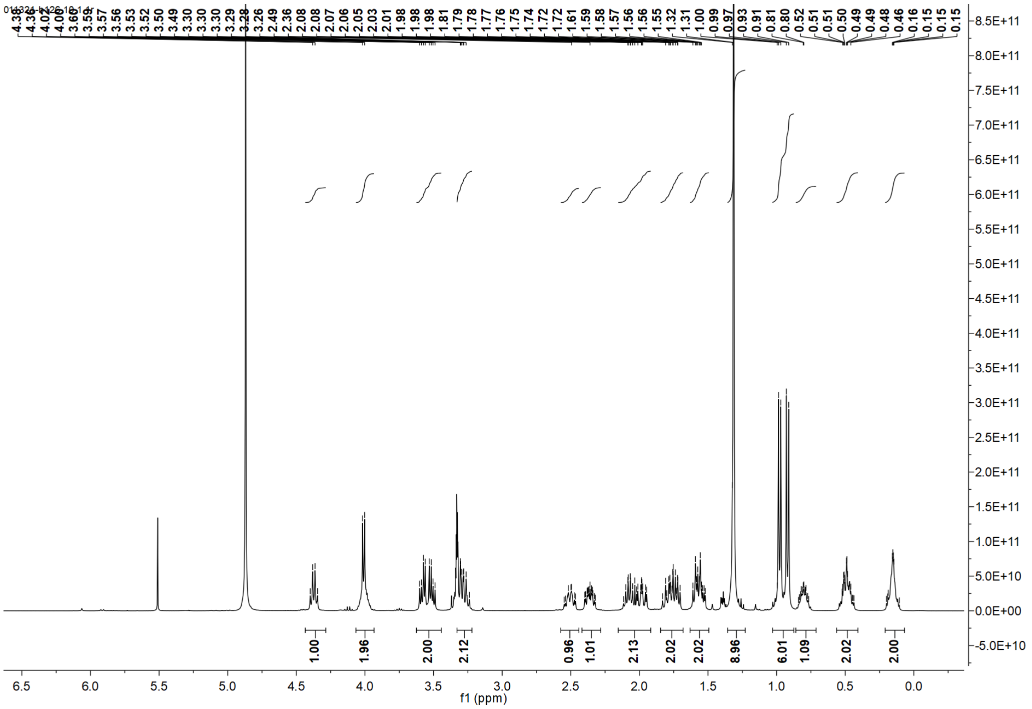
4.38 4.36 4.02 4.00 3.60 3.59 3.57 3.56 3.53 3.52 3.50 3.49 3.30 3.30 3.30 3.29 3.28 3.26 2.49 2.36 2.08 2.08 2.07 2.06 2.05 2.03 2.01 1.98 1.98 1.98 1.81 1.79 1.78 1.77 1.76 1.75 1.74 1.72 1.72 1.61 1.59 1.58 1.57 1.56 1.56 1.55 1.32 1.31 1.00 0.99 0.97 0.93 0.91 0.81 0.80 0.52 0.51 0.51 0.50 0.49 0.49 0.48 0.46 0.16 0.15 0.15
1.00 1.96 2.00 2.12 0.96 1.01 2.13 2.02 2.02 8.96 6.01 1.09 2.02 2.00
f1 (ppm)

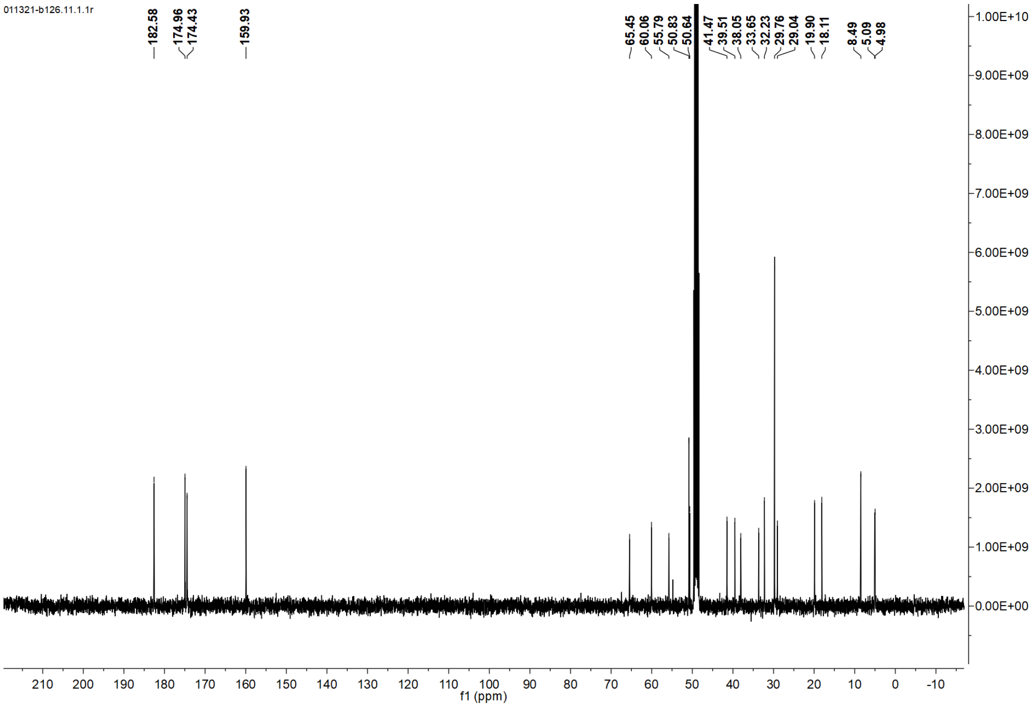
011321-b126.11.1.1r
182.58
174.96
174.43
159.93
65.45
60.06
55.79
50.83
50.64
41.47
39.51
38.05
33.65
32.23
29.76
29.04
19.90
18.11
8.49
5.09
4.98
f1 (ppm)

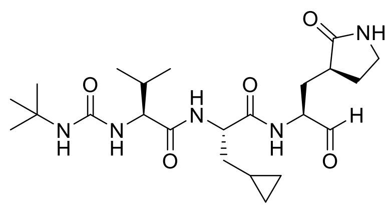

**MPI39**

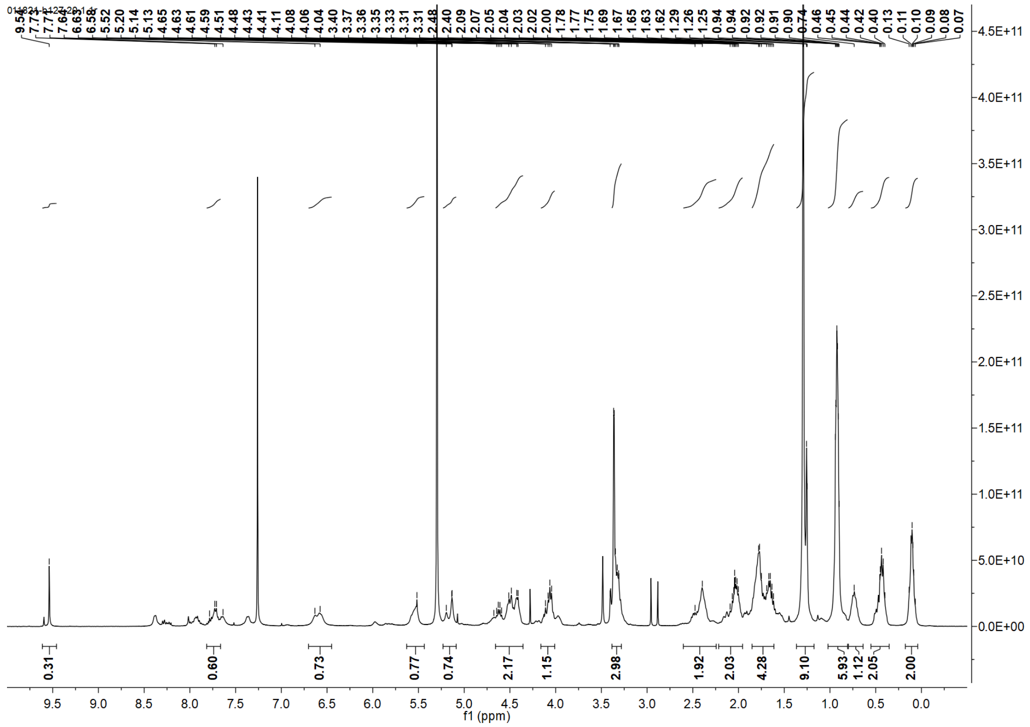

9.54
7.71
7.71
7.64
6.63
6.58
5.52
5.20
5.14
5.13
4.65
4.63
4.61
4.59
4.51
4.48
4.43
4.41
4.11
4.08
4.06
4.04
3.40
3.37
3.36
3.35
3.33
3.31
3.31
2.48
2.40
2.09
2.07
2.05
2.04
2.03
2.02
2.00
1.78
1.77
1.75
1.69
1.67
1.65
1.63
1.62
1.29
1.26
1.25
0.94
0.94
0.92
0.92
0.91
0.90
0.74
0.46
0.45
0.44
0.42
0.40
0.13
0.11
0.10
0.09
0.08
0.07
4.5E+11
4.0E+11
3.5E+11
3.0E+11
2.5E+11
2.0E+11
1.5E+11
1.0E+11
5.0E+10
0.0E+00
0.31
0.60
0.73
0.77
0.74
2.17
1.15
2.98
1.92
2.03
4.28
9.10
5.93
1.12
2.05
2.00
9.5
9.0
8.5
8.0
7.5
7.0
6.5
6.0
5.5
5.0
4.5
4.0
3.5
3.0
2.5
2.0
1.5
1.0
0.5
0.0
f1 (ppm)

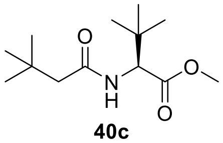

**40c**

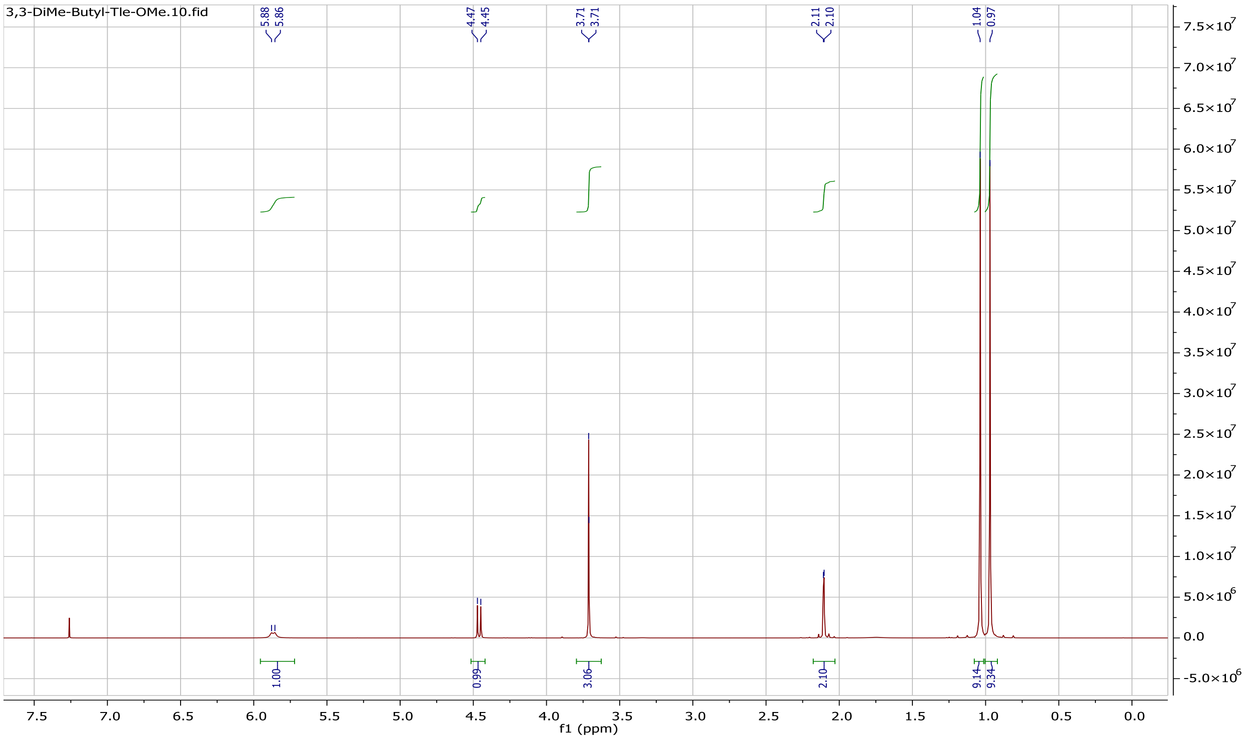
3,3-DiMe-Butyl-Tle-OMe.10.fid
f1 (ppm)

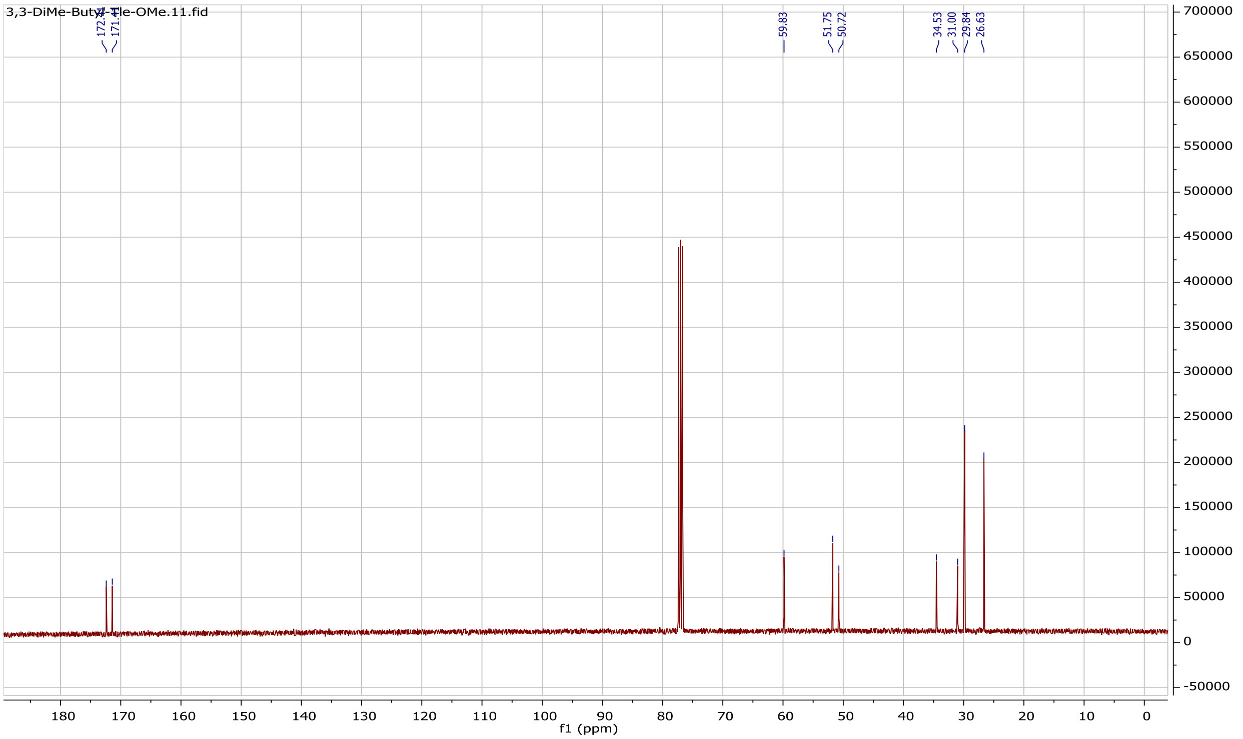
3,3-DiMe-Butyl-Tle-OMe.11.fid
f1 (ppm)

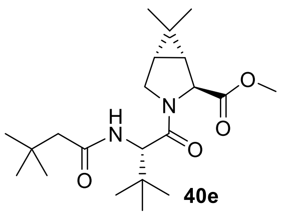

**40e**

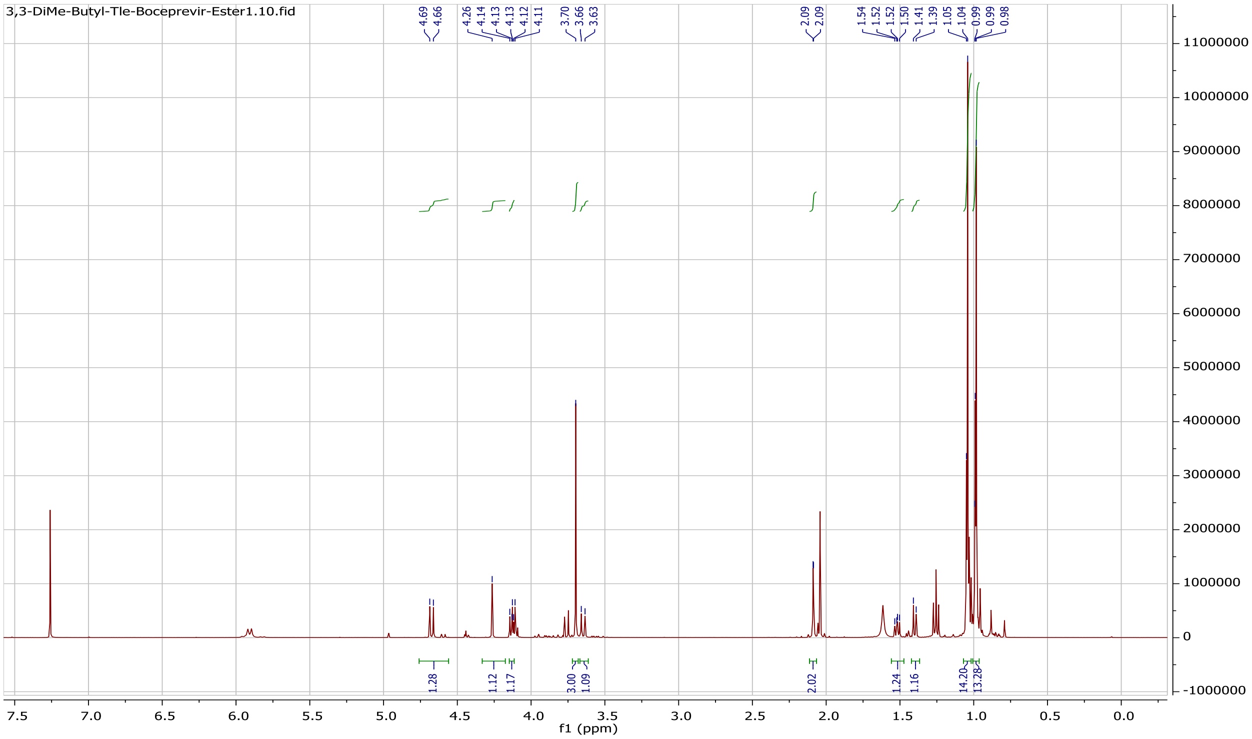

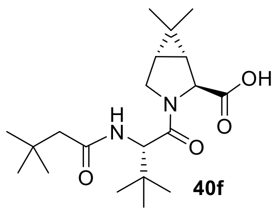


40f

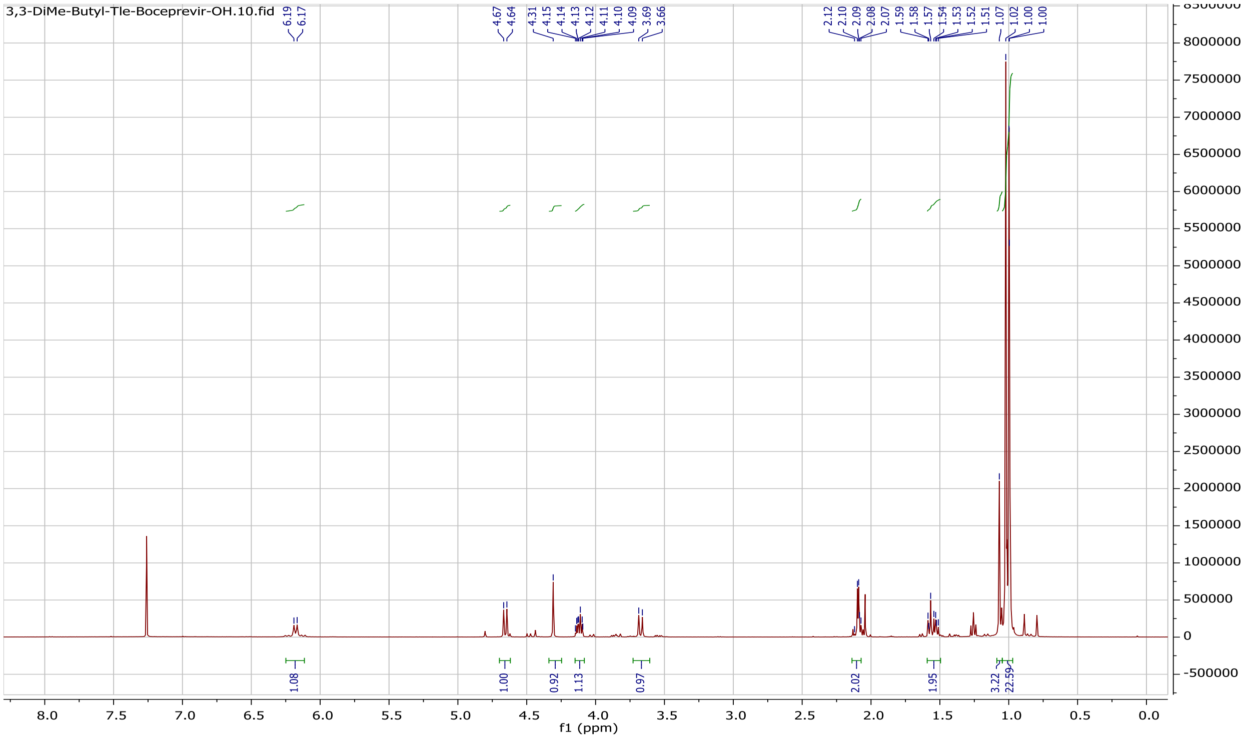

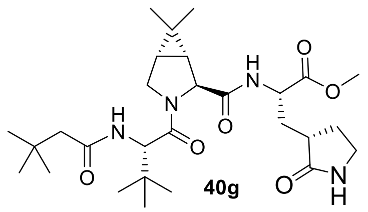

**40g**

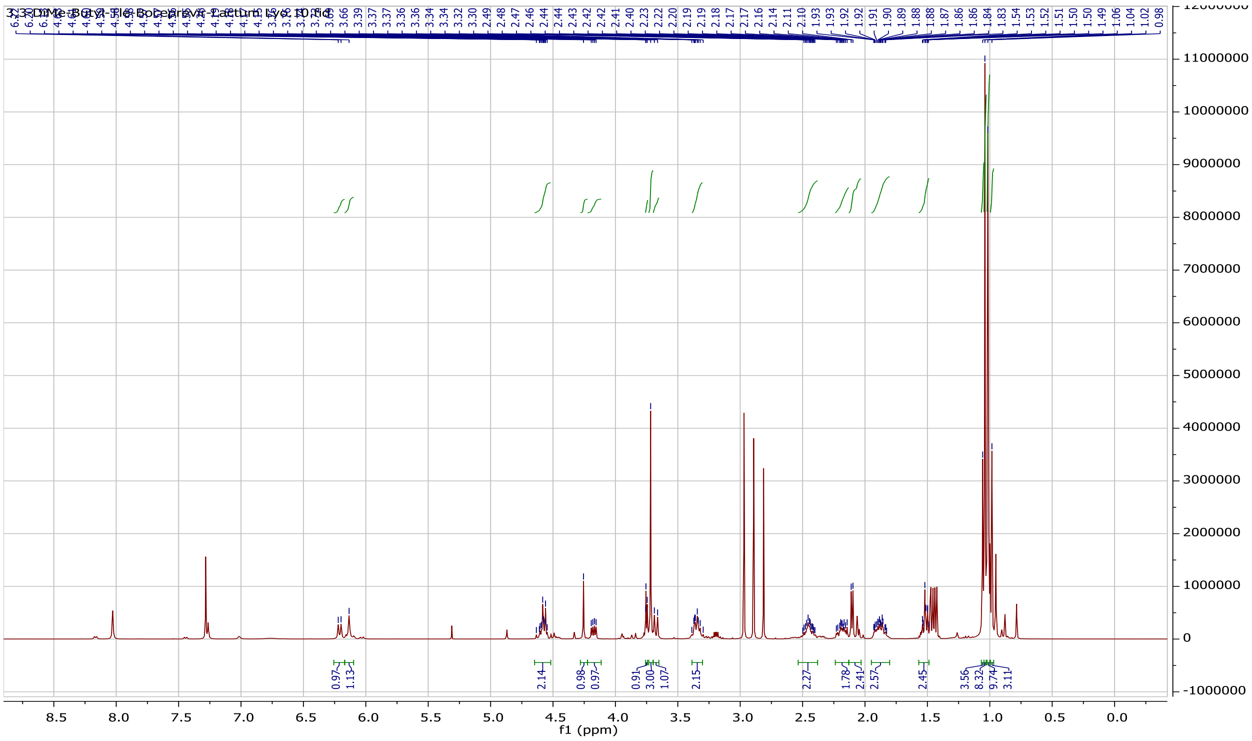

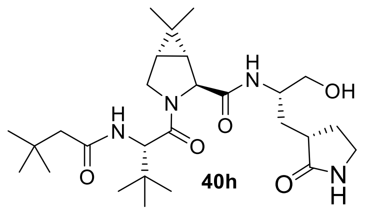

H
N
OH
O
H
N
N
O
O
O
NH
H


**40h**

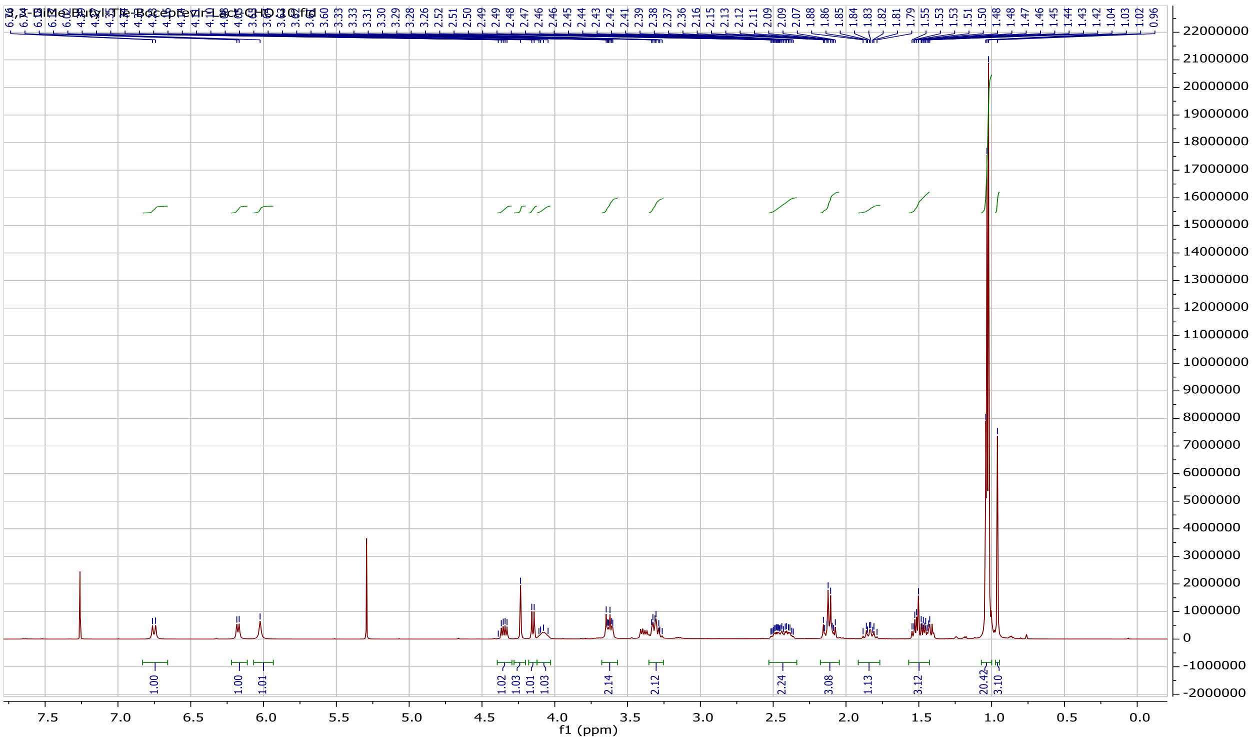

3,3-DiMe-Butyl-Tle-Boceprevir-Lact-CHO.10.fid
1.00
1.00
1.01
1.02
1.03
1.01
1.03
2.14
2.12
2.24
3.08
1.13
3.12
20.42
3.10
f1 (ppm)


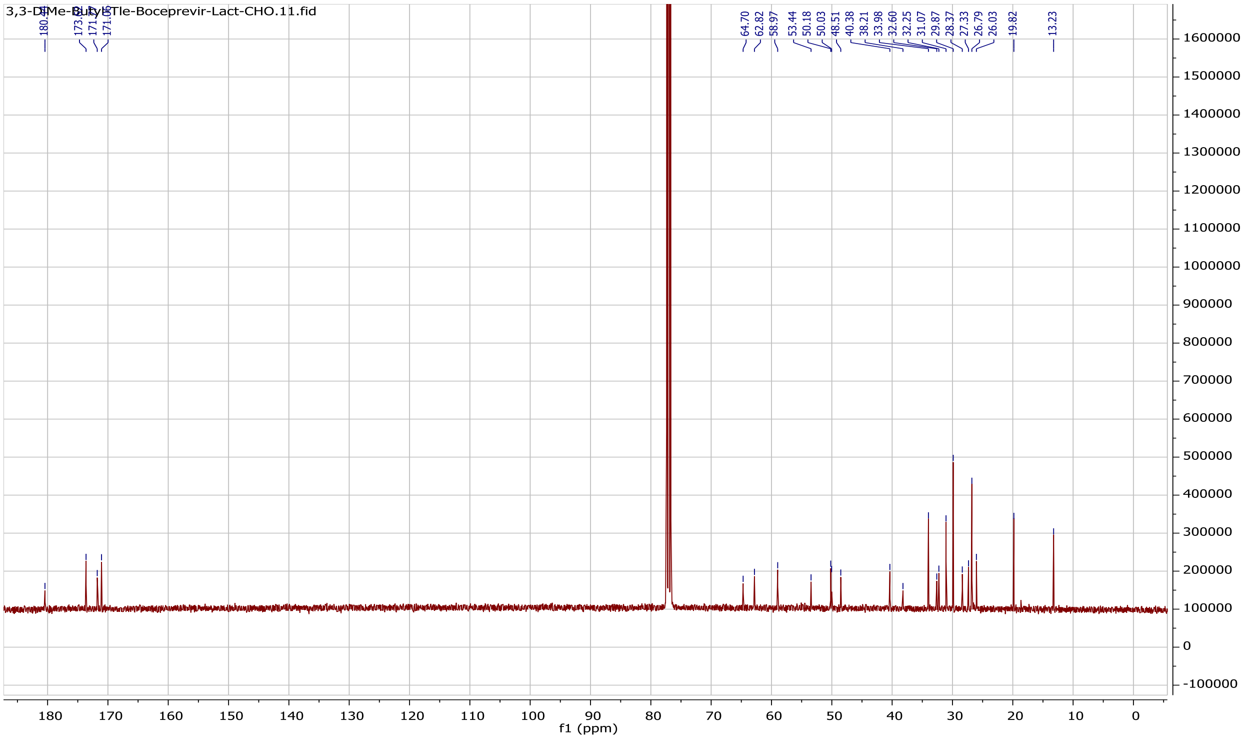

3,3-DiMe-Butyl-Tle-Boceprevir-Lact-CHO.11.fid
64.70
62.82
58.97
53.44
50.18
50.03
48.51
40.38
38.21
33.98
32.60
32.25
31.07
29.87
28.37
27.33
26.79
26.03
19.82
13.23
f1 (ppm)

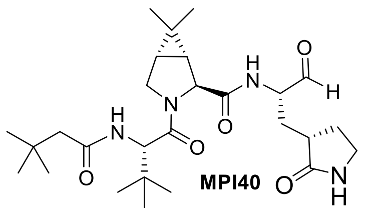

MPI40

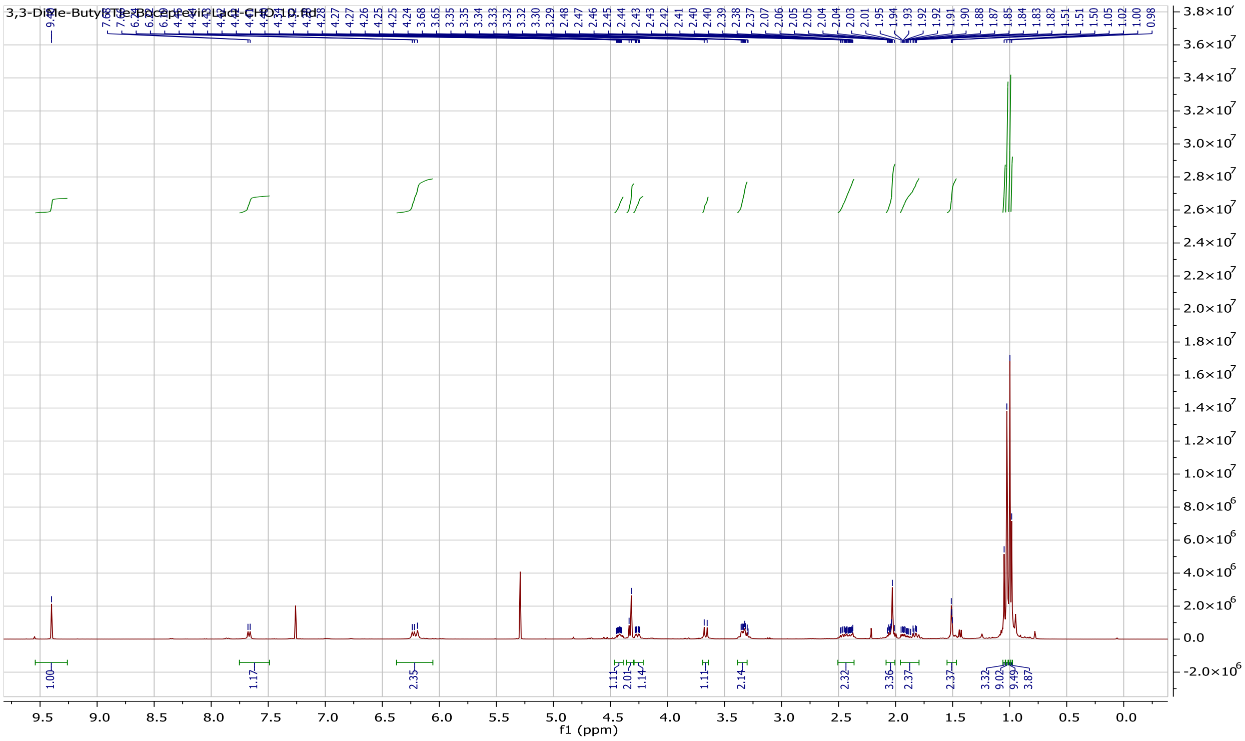
f1 (ppm)

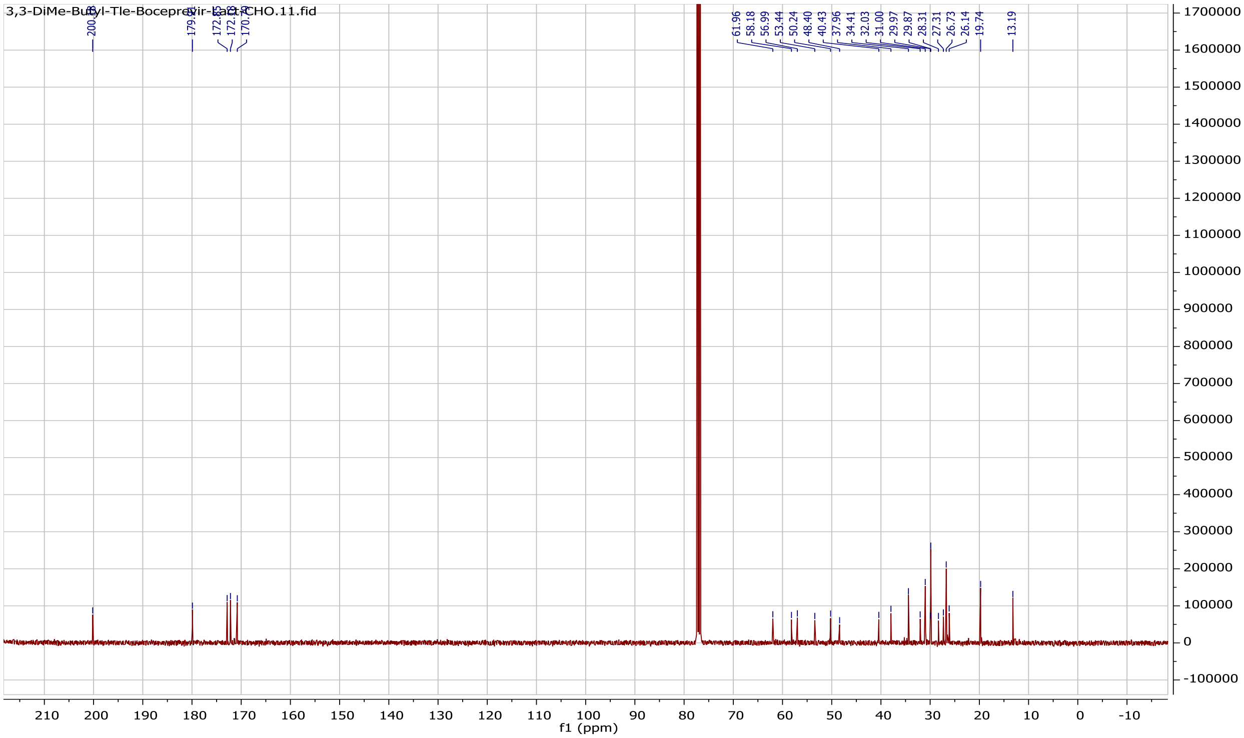
3,3-DiMe-Butyl-Tle-Boceprevir-Lact-CHO.11.fid
61.96
58.18
56.99
53.44
50.24
48.40
40.43
37.96
34.41
32.03
31.00
29.97
29.87
28.31
27.31
26.73
26.14
19.74
13.19
f1 (ppm)

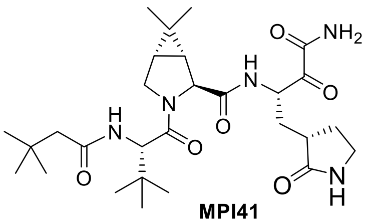

**MPI41**

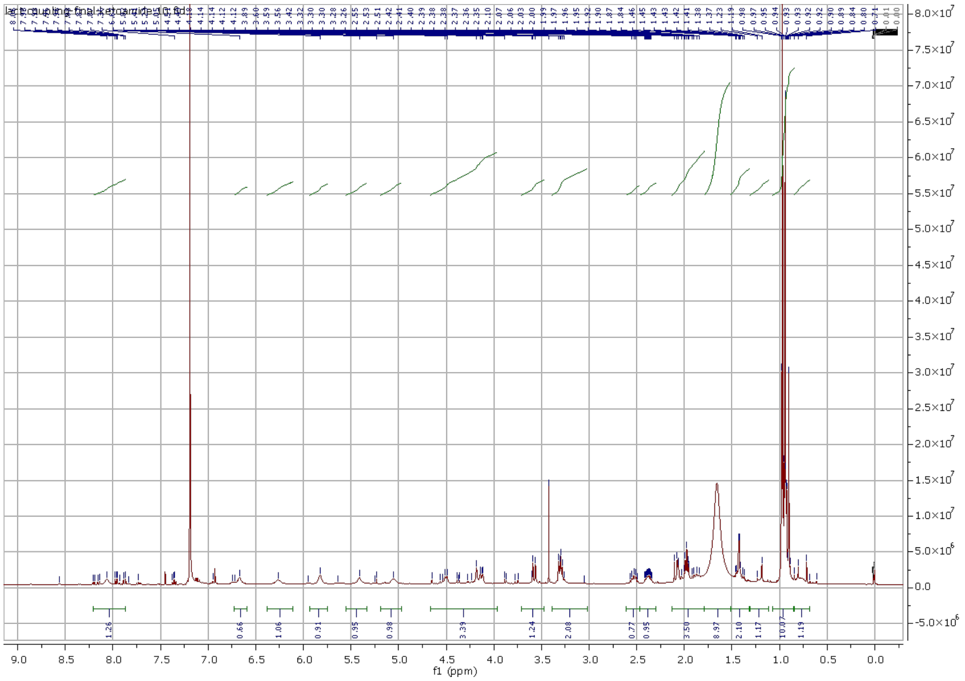

1.26
0.66
1.06
0.91
0.95
0.98
3.39
1.24
2.08
0.77
0.95
3.50
8.97
2.10
1.17
10.07
1.19
9.0 8.5 8.0 7.5 7.0 6.5 6.0 5.5 5.0 4.5 4.0 3.5 3.0 2.5 2.0 1.5 1.0 0.5 0.0
f1 (ppm)

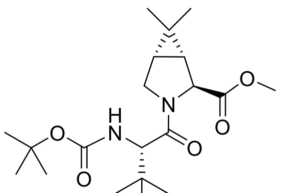
H
N
O
O
O
N
O
O

**42b**

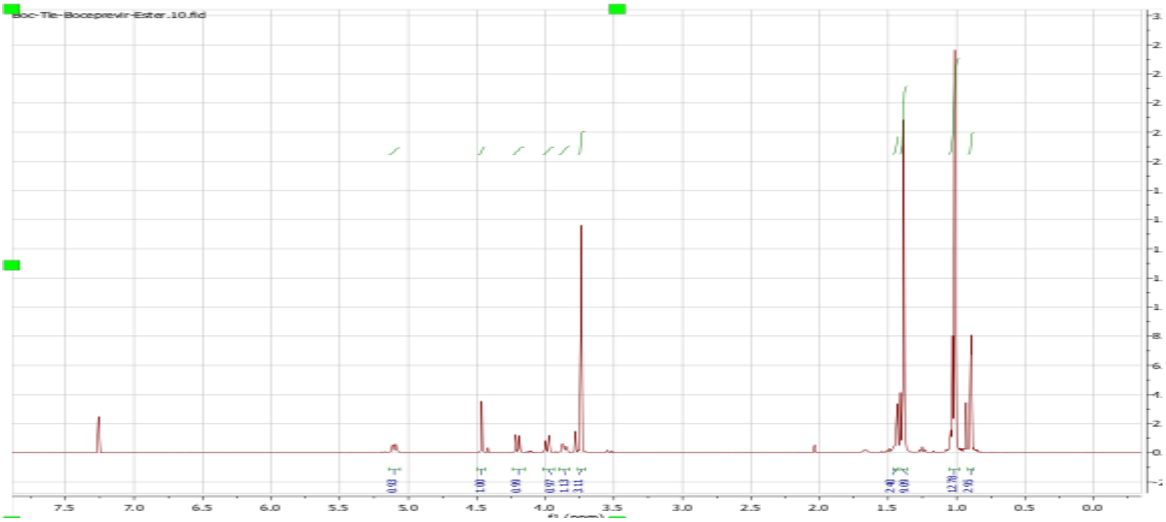
Boc-Tle-Boceprevir-Ester.10.fid
7.5
7.0
6.5
6.0
5.5
5.0
4.5
4.0
3.5
3.0
2.5
2.0
1.5
1.0
0.5
0.0

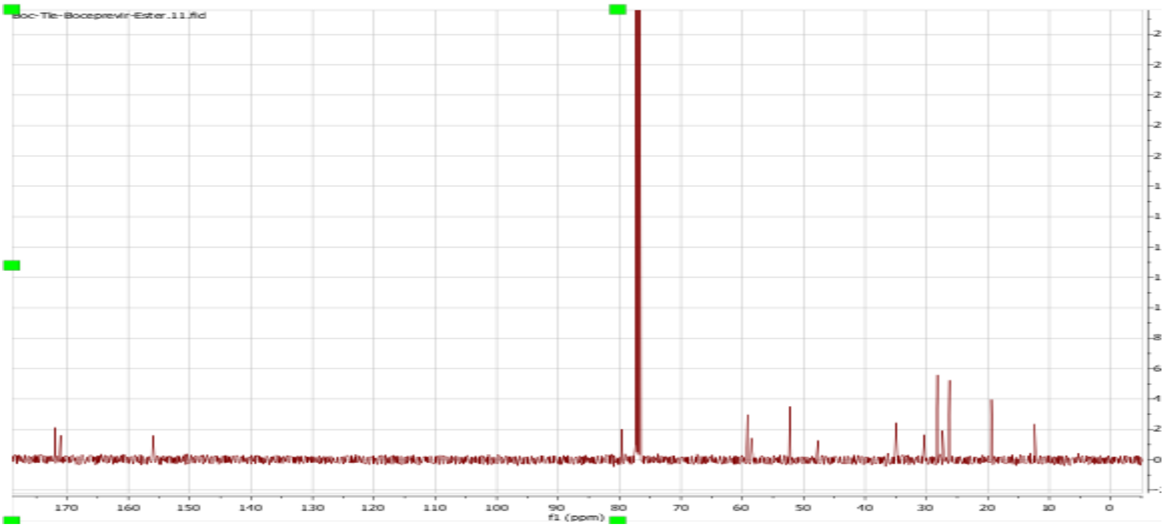
Boc-Tle-Boceprevir-Ester.11.fid
170
160
150
140
130
120
110
100
90
80
70
60
50
40
30
20
10
0
f1 (ppm)

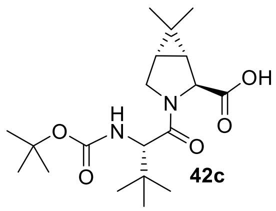

**42c**

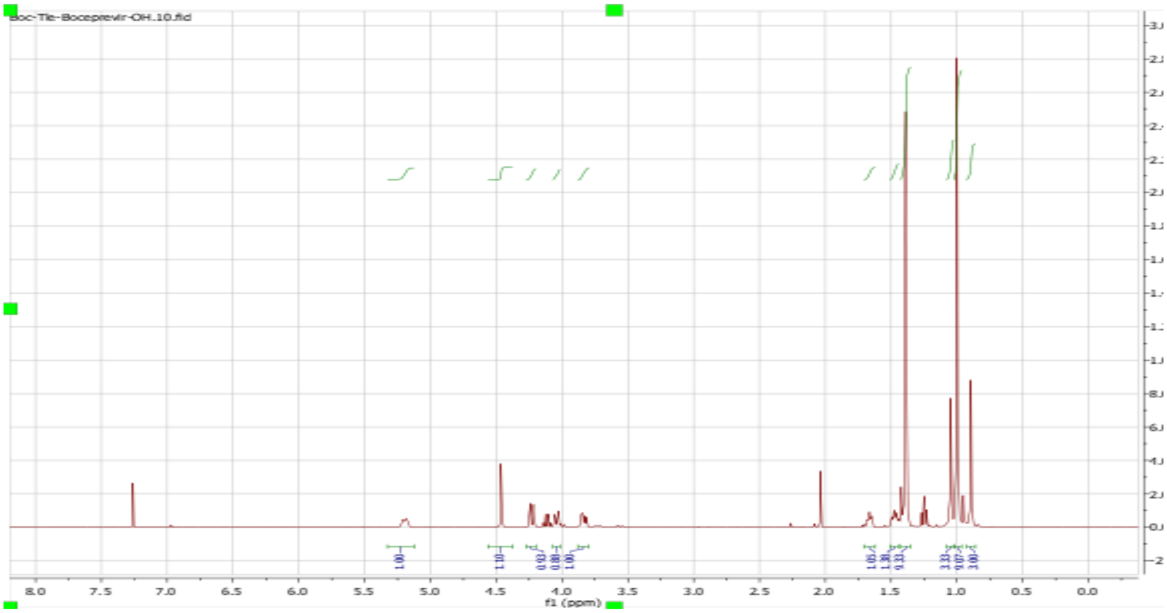

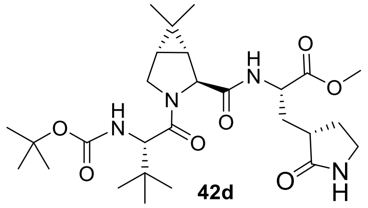

H
N
O
O
O
O
N
H
N
O
O
O
N
H


**42d**

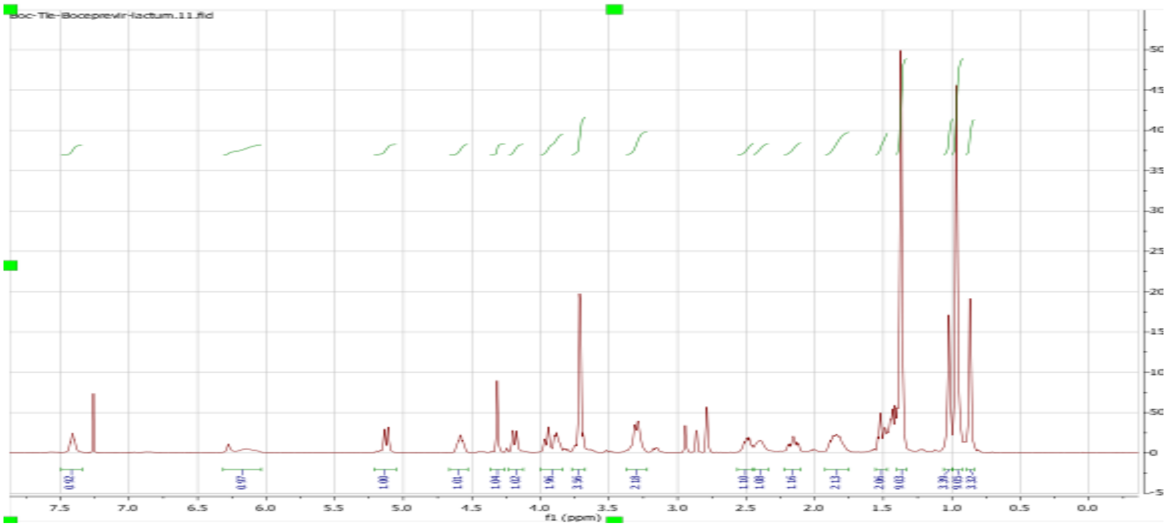

Boc-Tle-Boceprevir-lactum.11.fid
f1 (ppm)


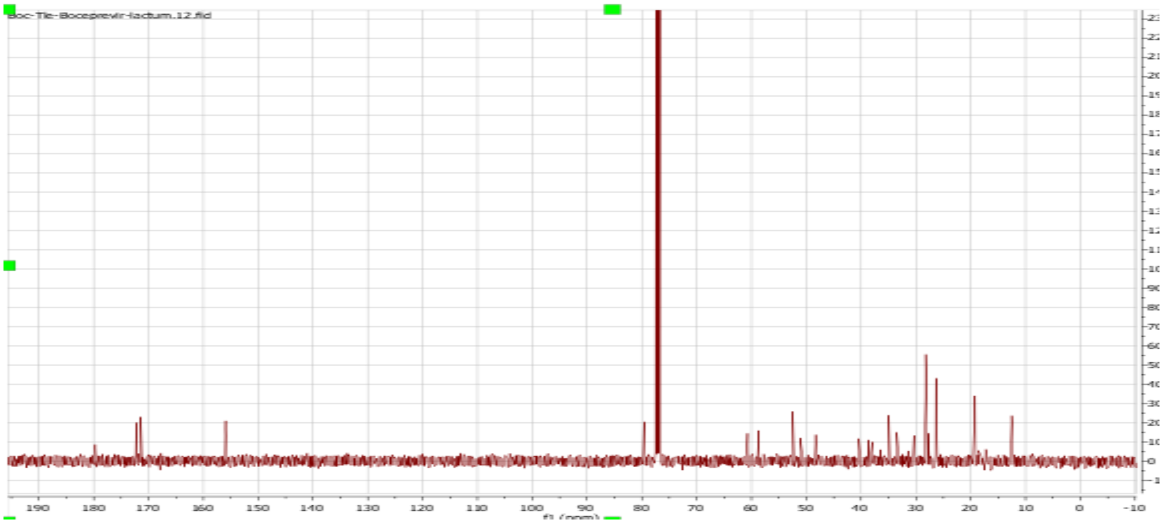

Boc-Tle-Boceprevir-lactum.12.fid
f1 (ppm)

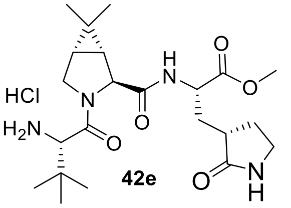
HCl
$H_2N$

**42e**

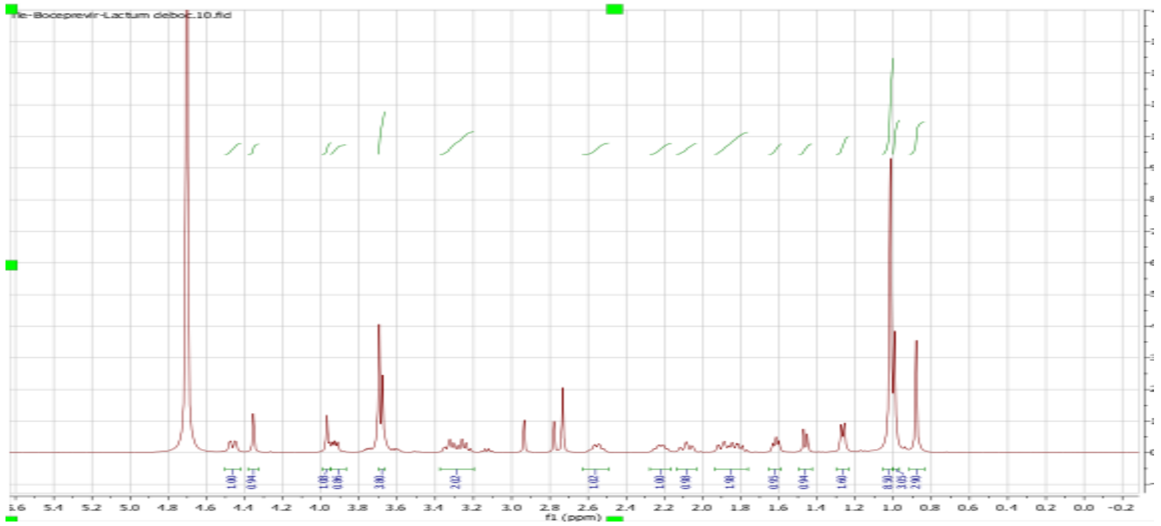
Tle-Boceprevir-Lactum deboc.10.fid
f1 (ppm)

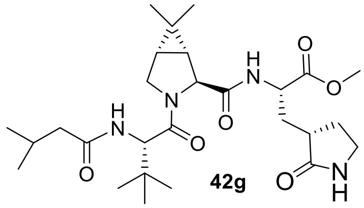

**42g**

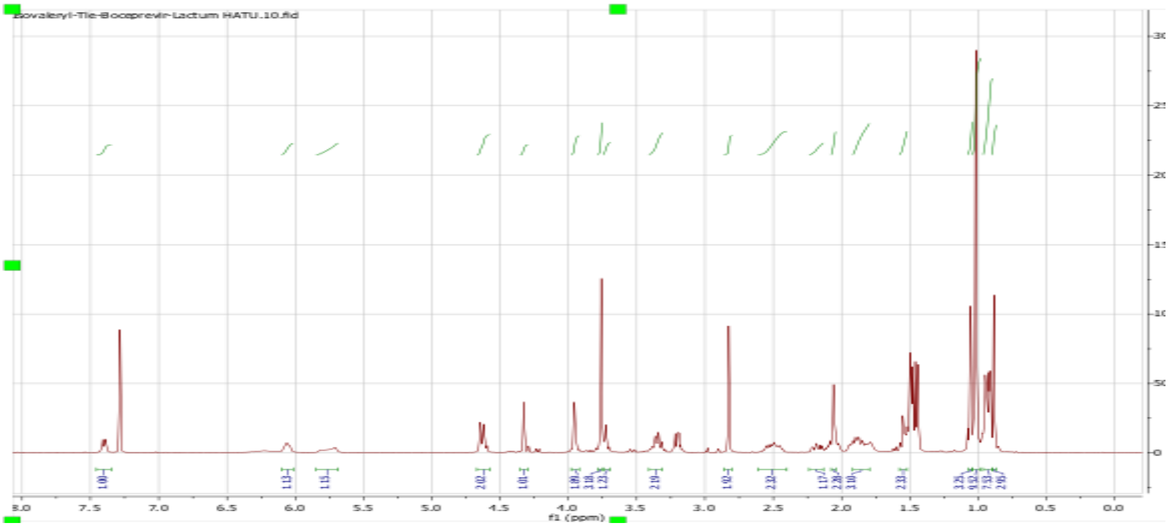
Isovaleryl-Tle-Boceprevir-Lactum HATU.10.fid
f1 (ppm)

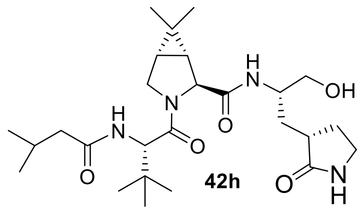

OH
O
O
N
H
O
N
H
N
O
NH
H


**42h**

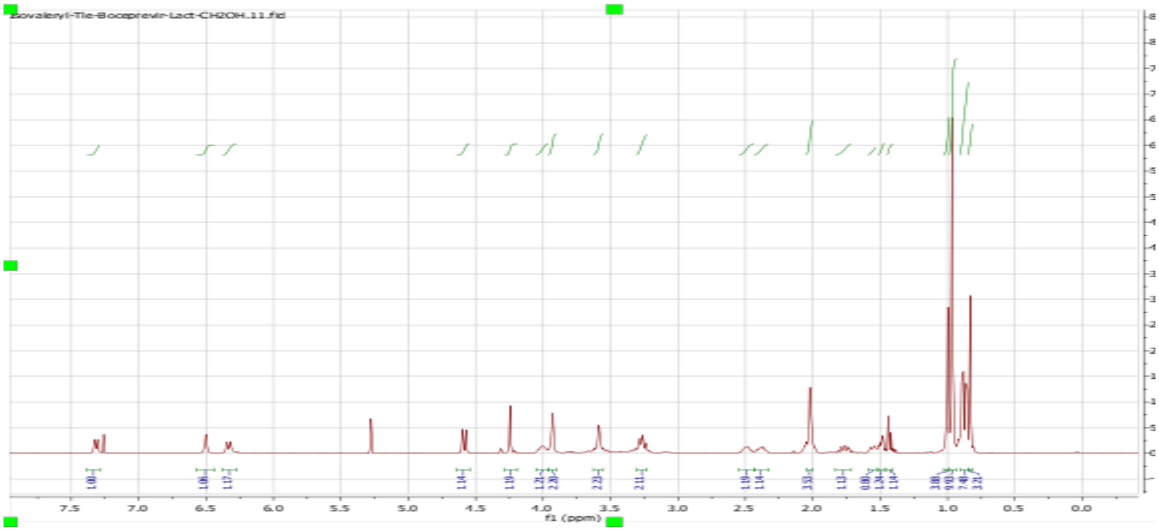

Isovaleryl-Tle-Boceprevir-Lact-CH2OH.11.fid
f1 (ppm)


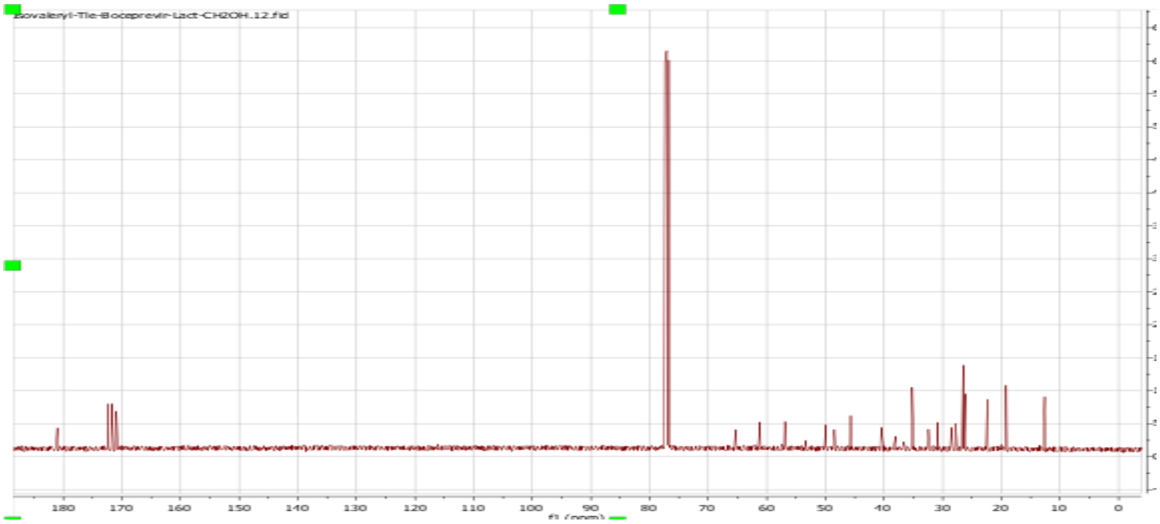

Isovaleryl-Tle-Boceprevir-Lact-CH2OH.12.fid
f1 (ppm)

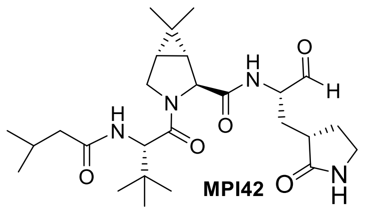

**MPI42**

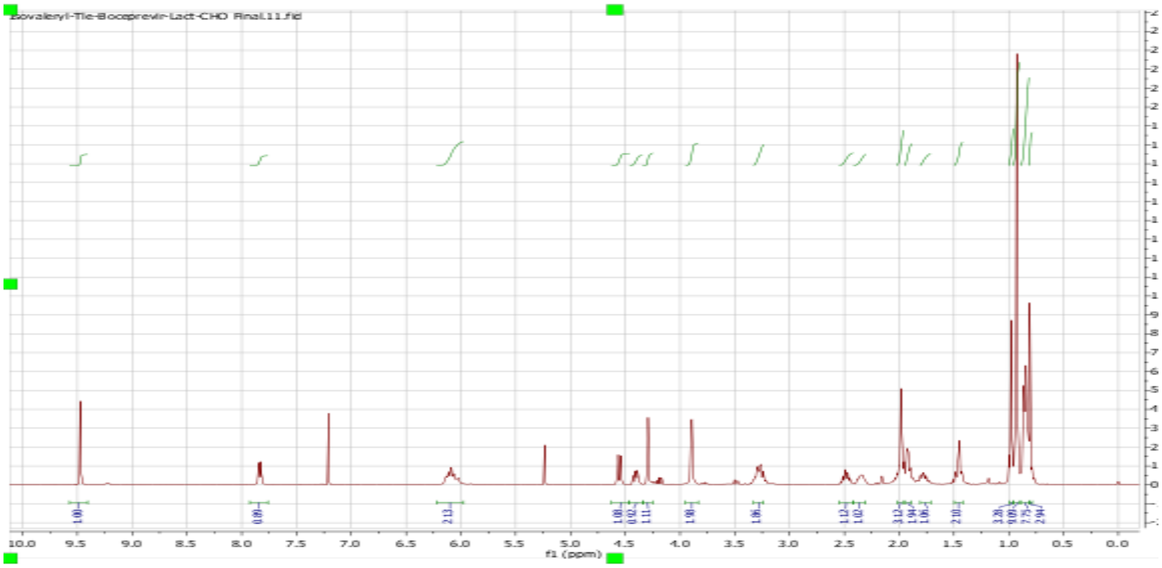

Isovaleryl-Tle-Boceprevir-Lact-CHO Final.11.fid
f1 (ppm)


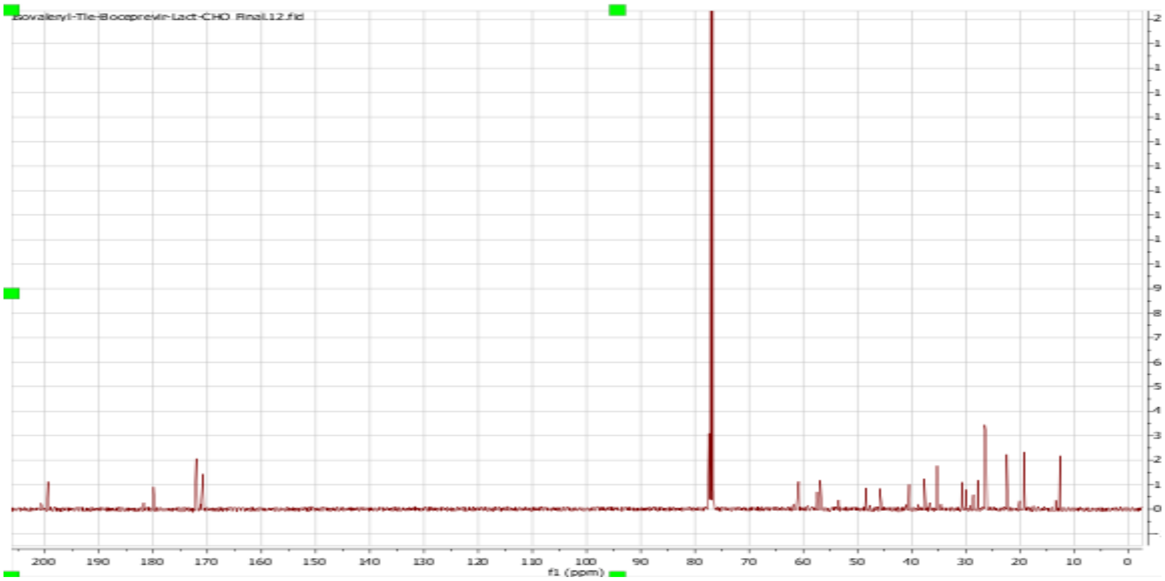

Isovaleryl-Tle-Boceprevir-Lact-CHO Final.12.fid
f1 (ppm)

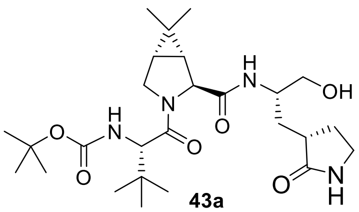
OH
O
N
H
H
N
O
O
H
N
O
O
N
H


**43a**

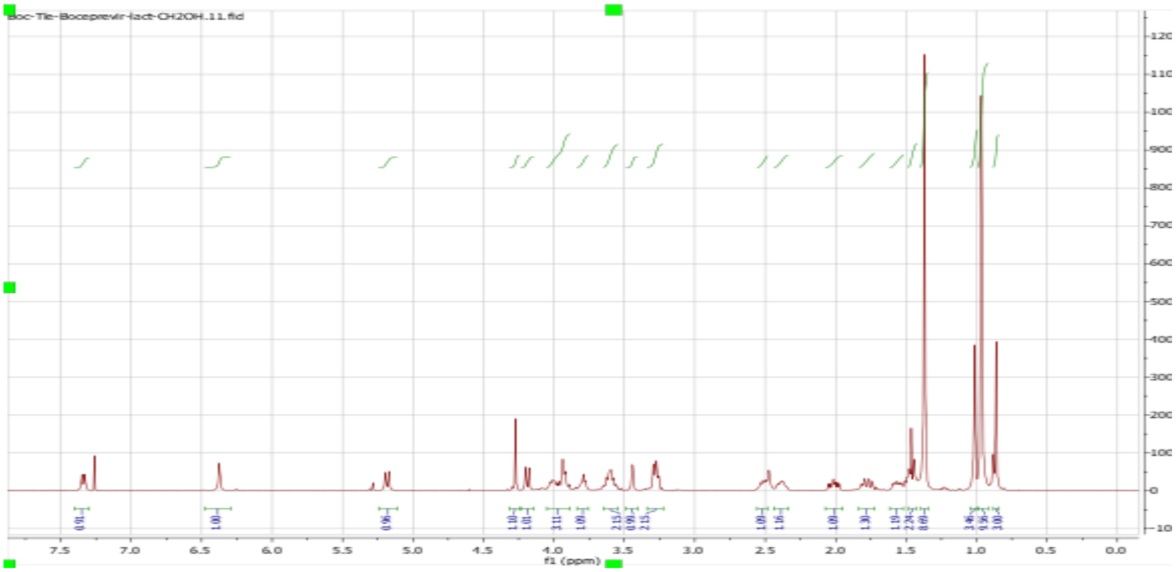
oc-Tle-Boceprevir-lact-CH2OH.11.fid
f1 (ppm)


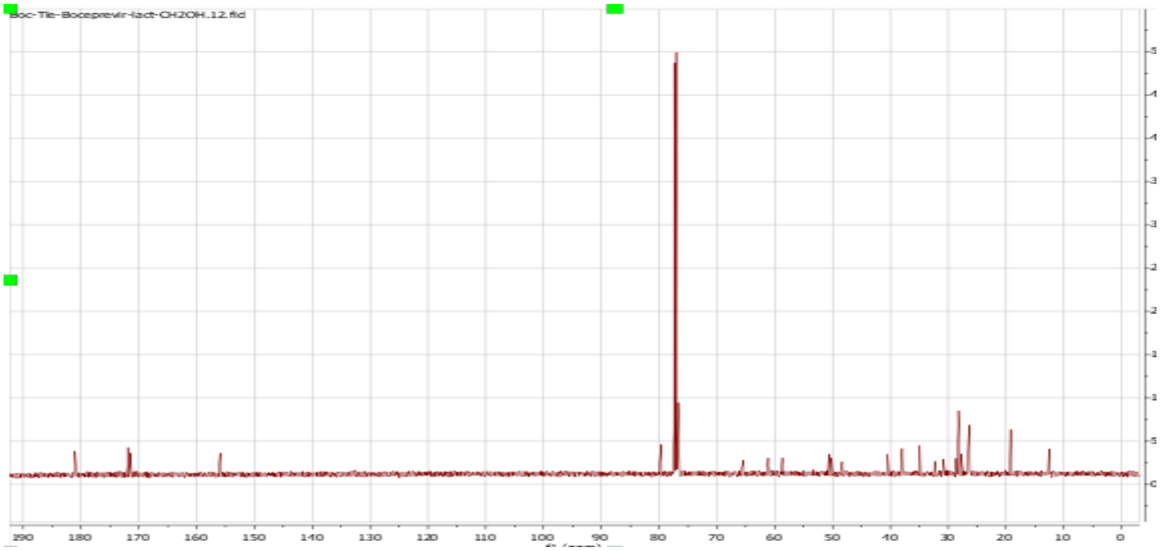
oc-Tle-Boceprevir-lact-CH2OH.12.fid

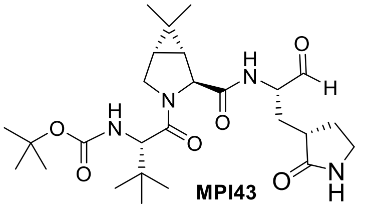

**MPI43**

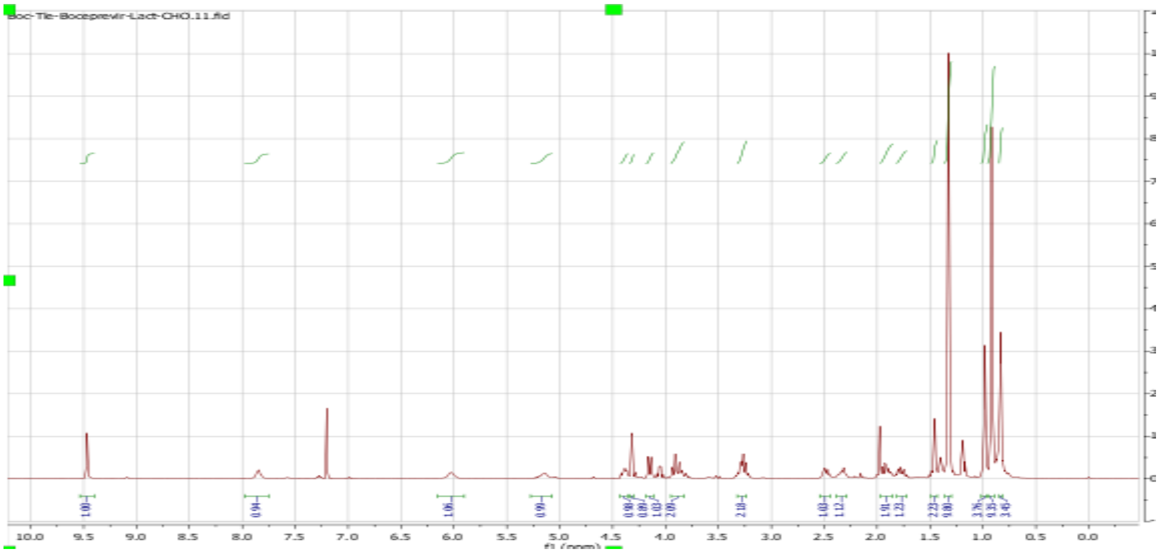
oc-Tle-Boceprevir-Lact-CHO.11.fid
f1 (ppm)

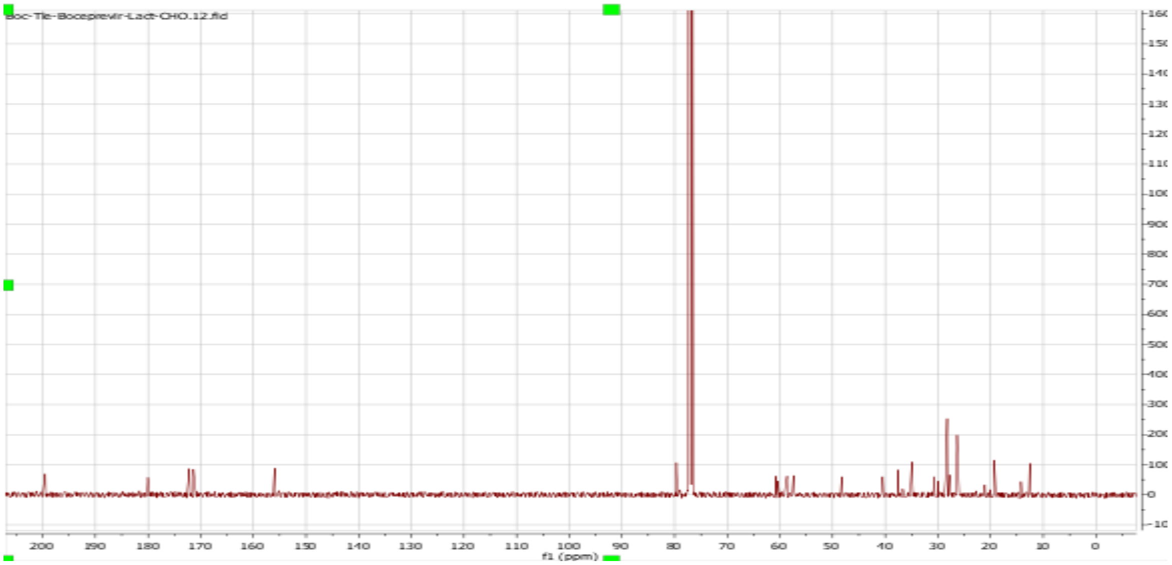
oc-Tle-Boceprevir-Lact-CHO.12.fid
f1 (ppm)

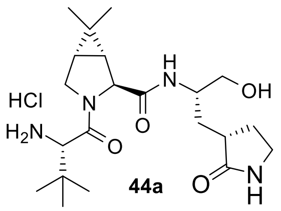
HCl
$H_2N$
OH
O
N
H
NH

**44a**

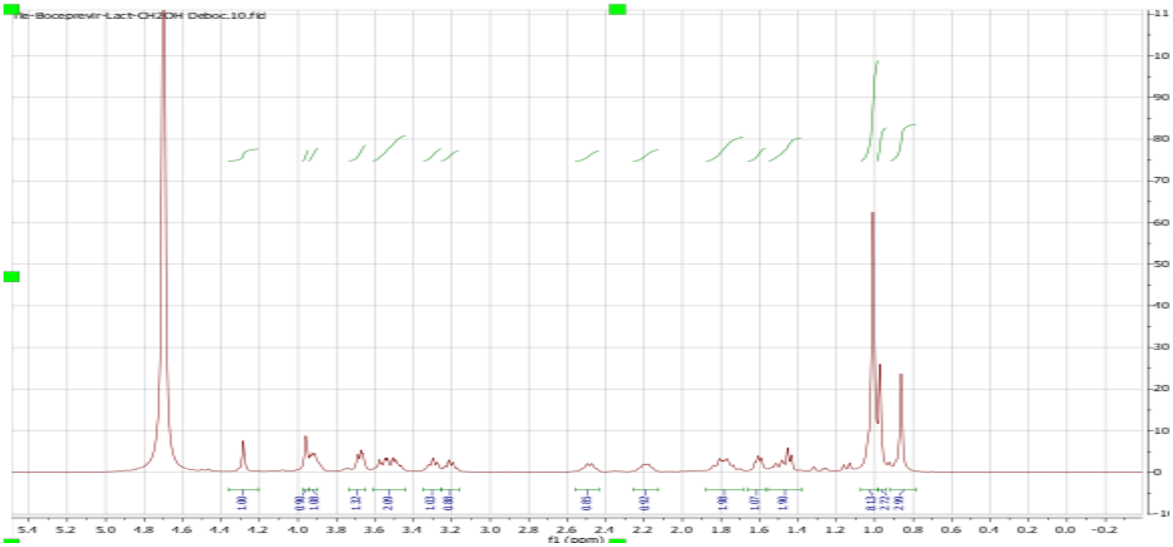
ile-Boceprevir-Lact-CH2OH Deboc.10.fid
f1 (ppm)

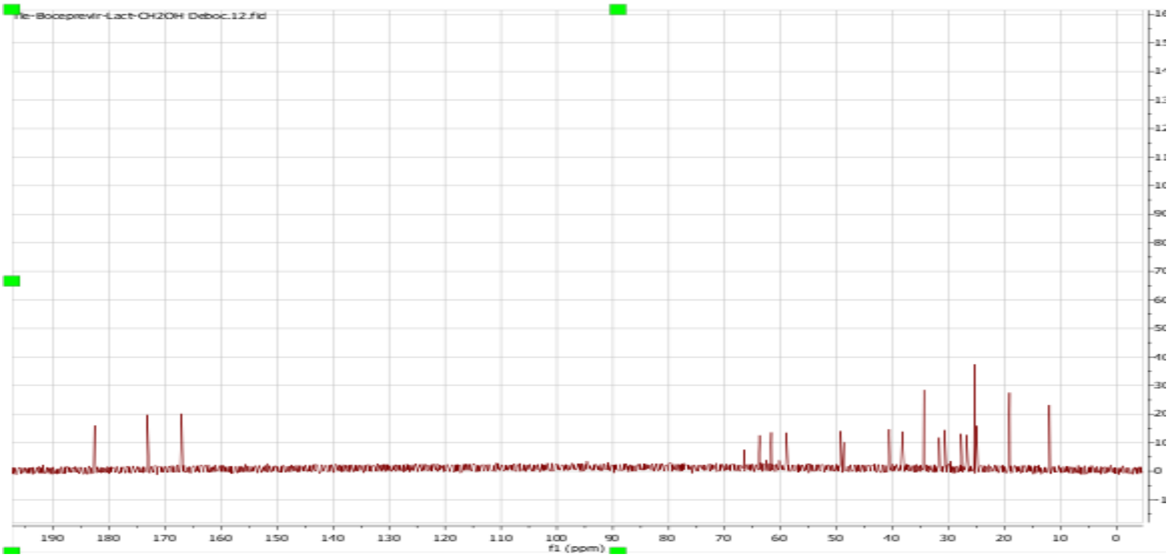
ile-Boceprevir-Lact-CH2OH Deboc.12.fid
f1 (ppm)

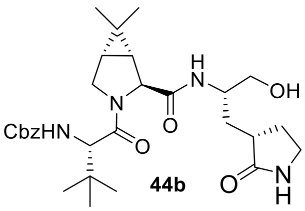
CbzHN
OH
O
N
H

**44b**

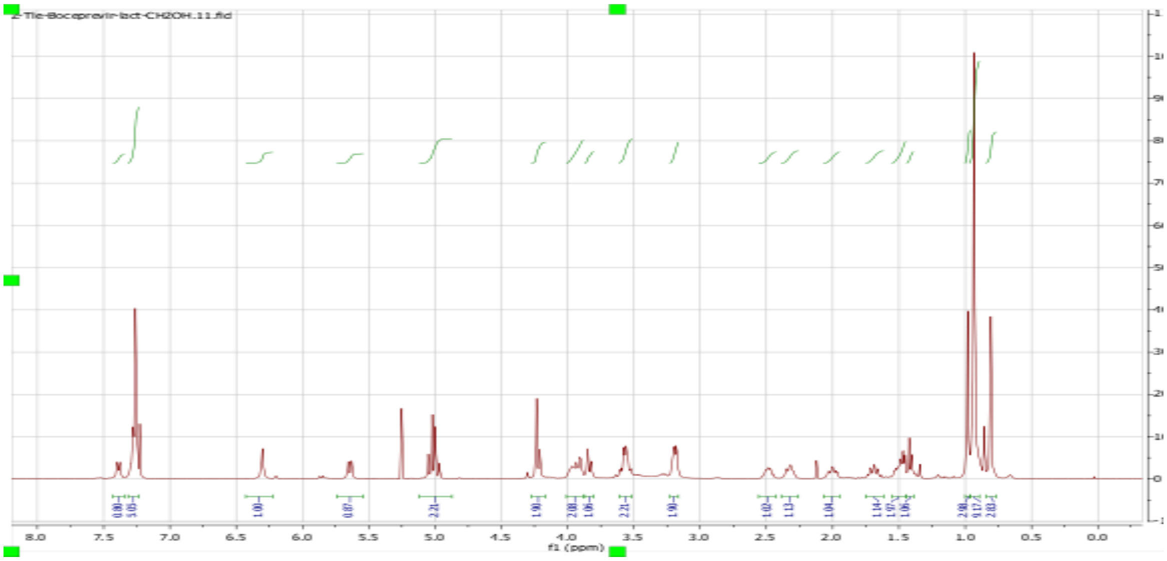
Z-Tle-Boceprevir-lact-CH2OH.11.fid
f1 (ppm)

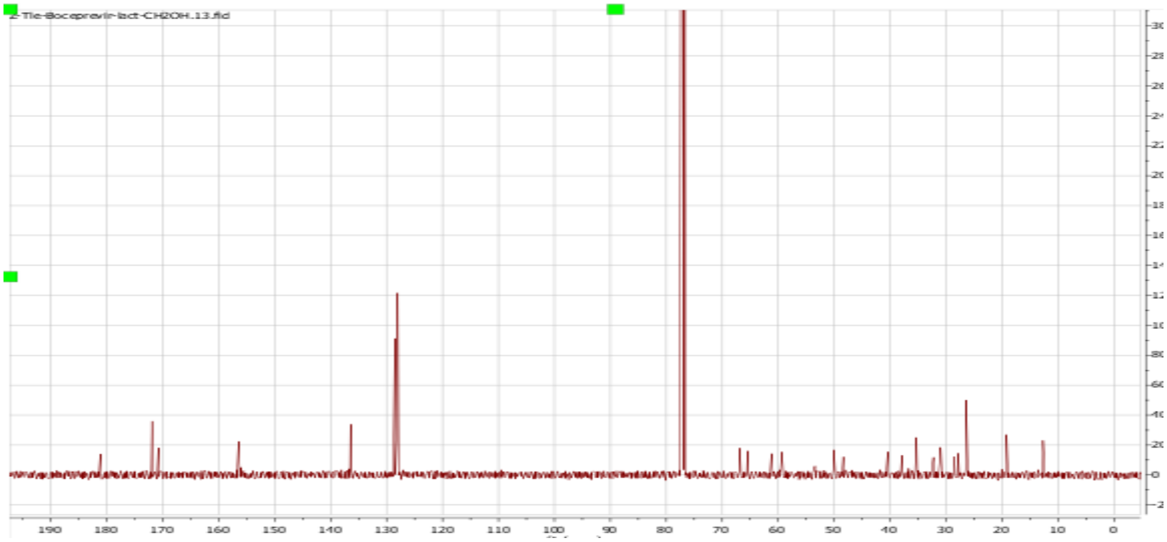
Z-Tle-Boceprevir-lact-CH2OH.13.fid

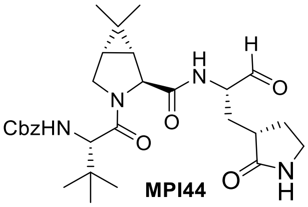
CbzHN
MPI44

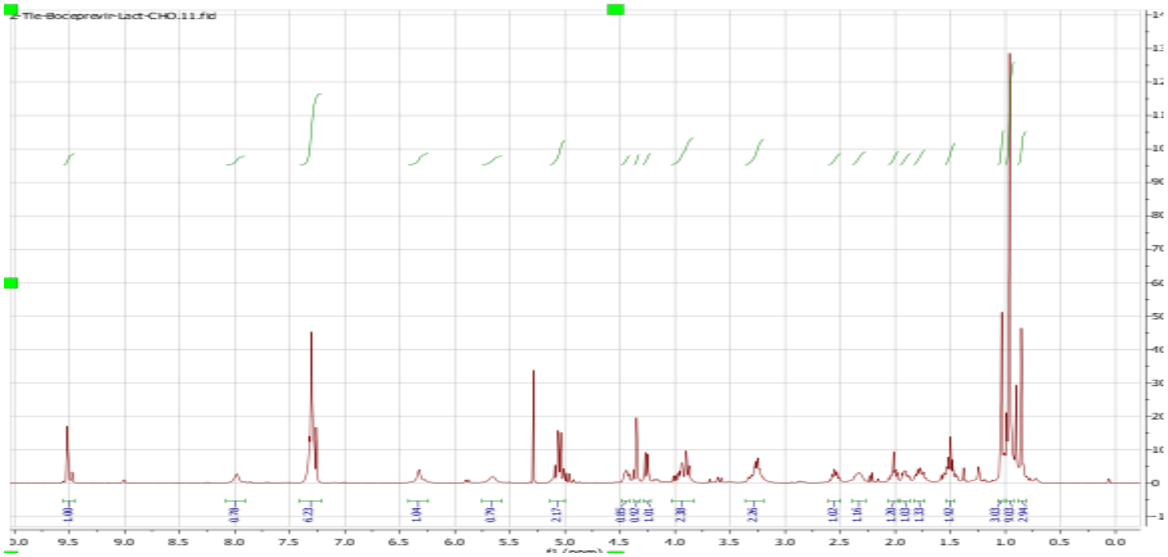
Z-Tle-Boceprevir-Lact-CHO.11.fid

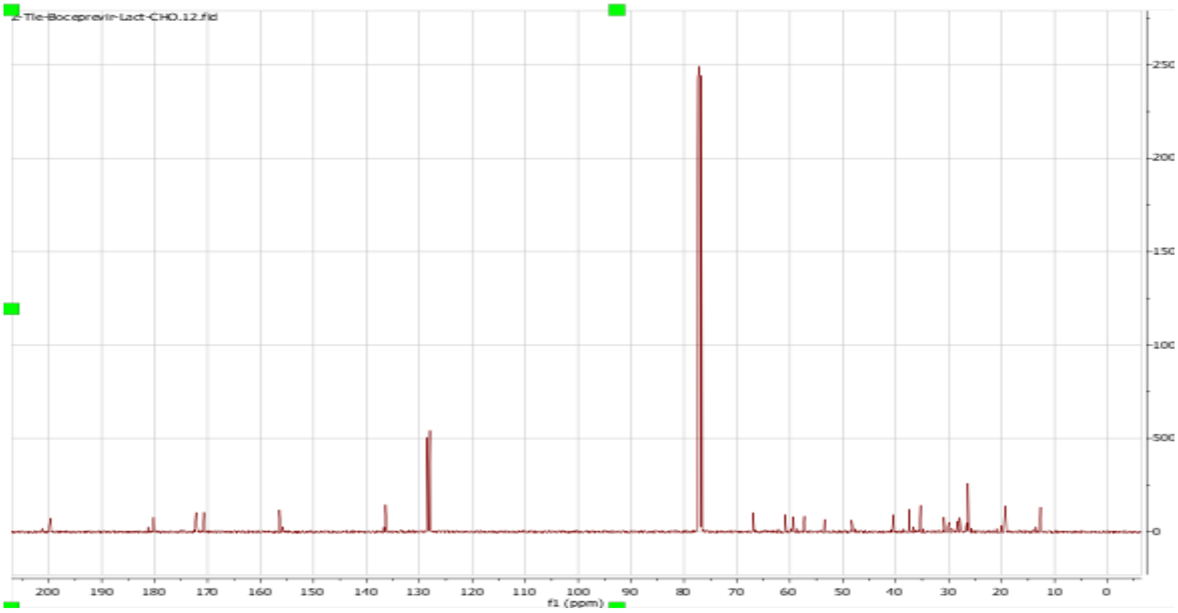
Z-Tle-Boceprevir-Lact-CHO.12.fid
f1 (ppm)

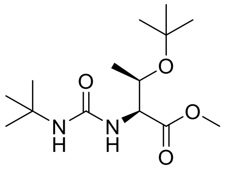

**45d**

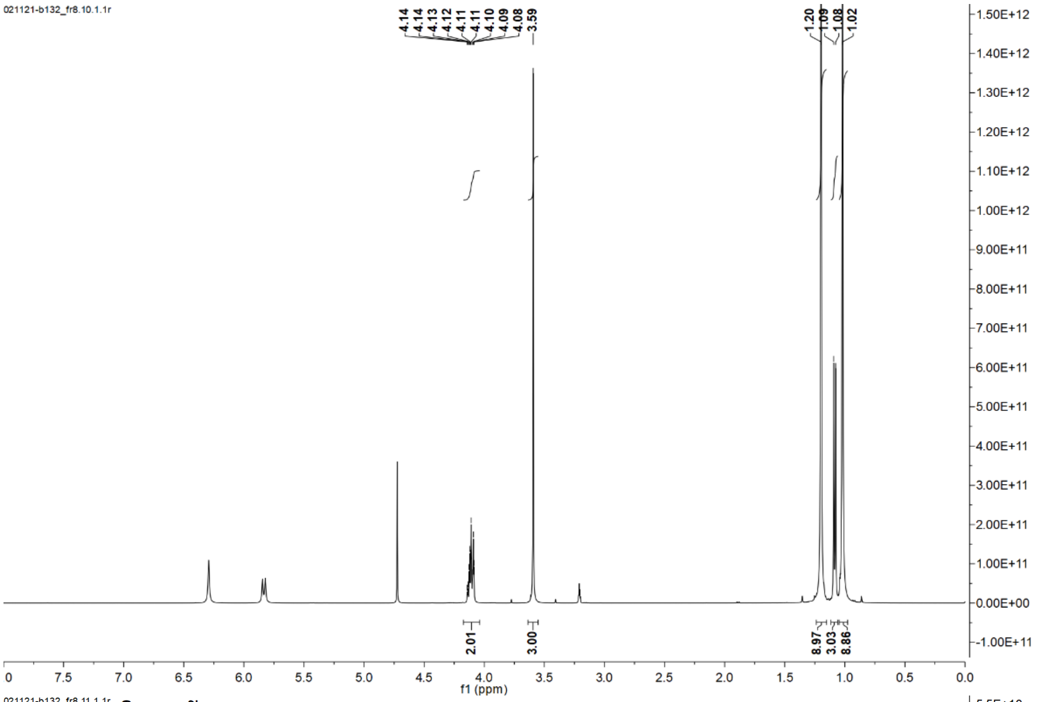
021121-b132_fr8.10.1.1r
4.14 4.14 4.13 4.12 4.11 4.11 4.10 4.09 4.08 3.59
1.20 1.09 1.08 1.02
2.01
3.00
8.97 3.03 8.86
f1 (ppm)

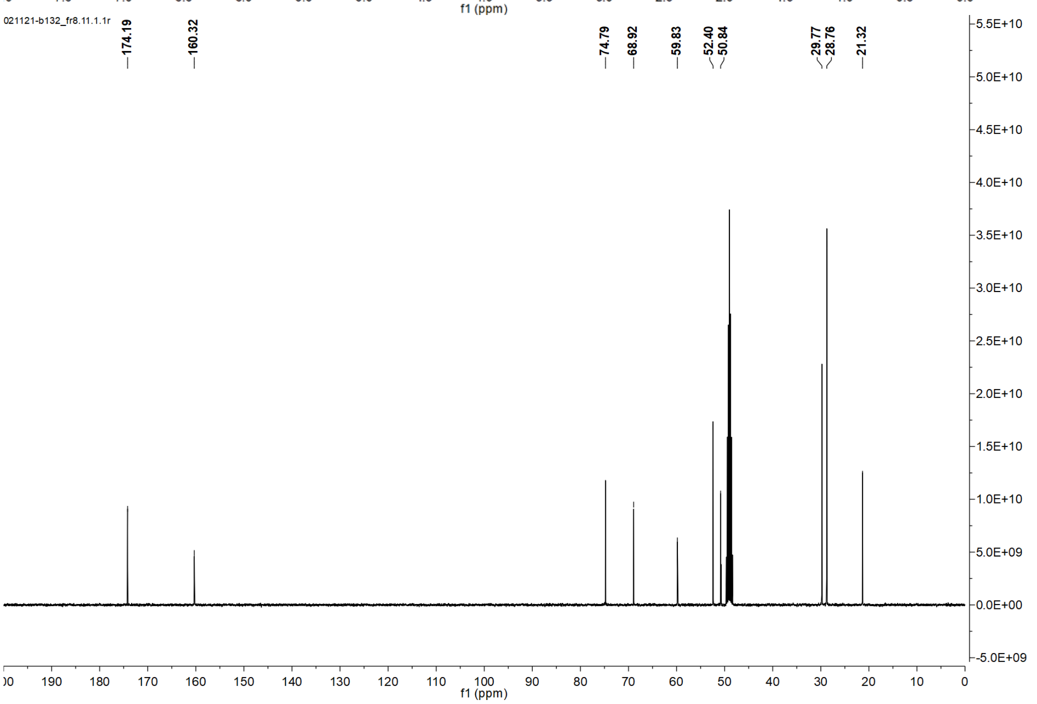
021121-b132_fr8.11.1.1r
174.19
160.32
74.79
68.92
59.83
52.40
50.84
29.77
28.76
21.32
f1 (ppm)

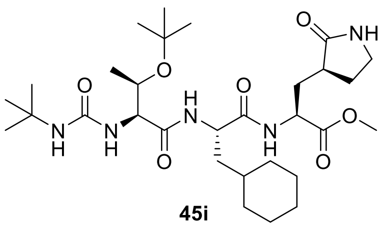

**45i**

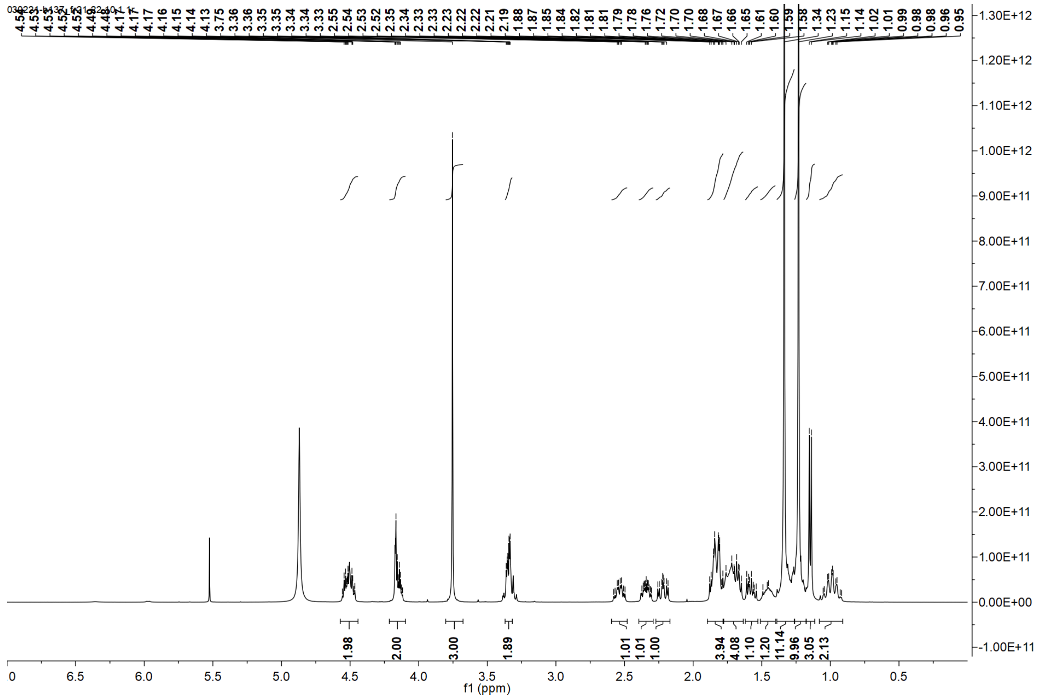

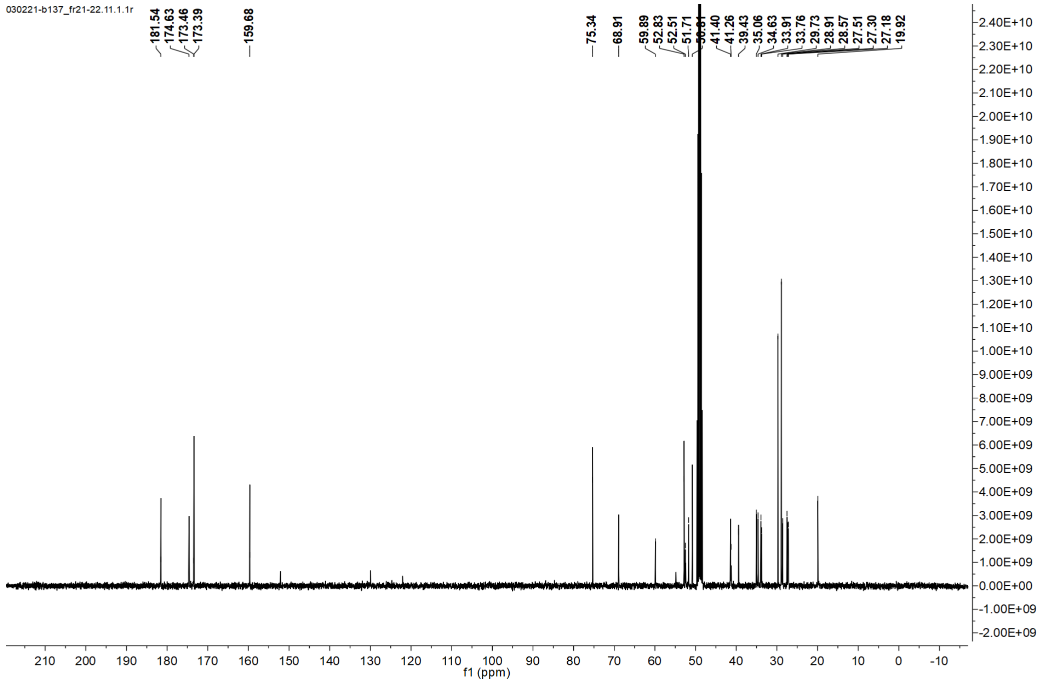

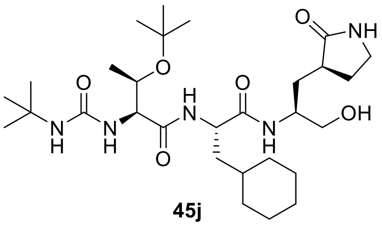
O
NH
O
O
NH
H
N
N
H
H
N
O
N
H
OH

**45j**

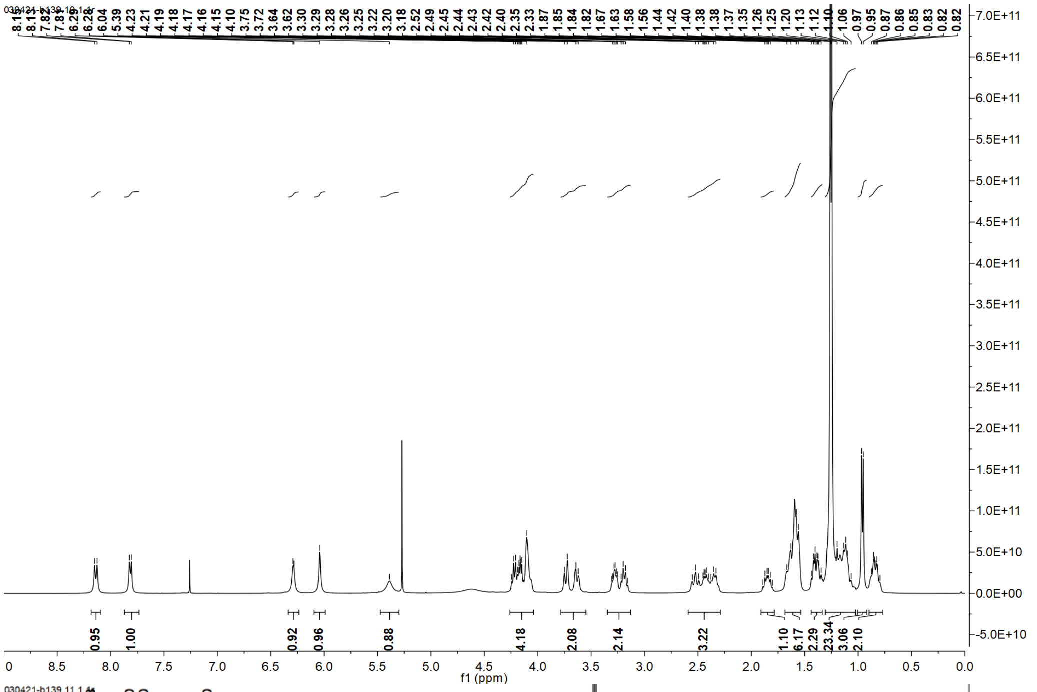
f1 (ppm)

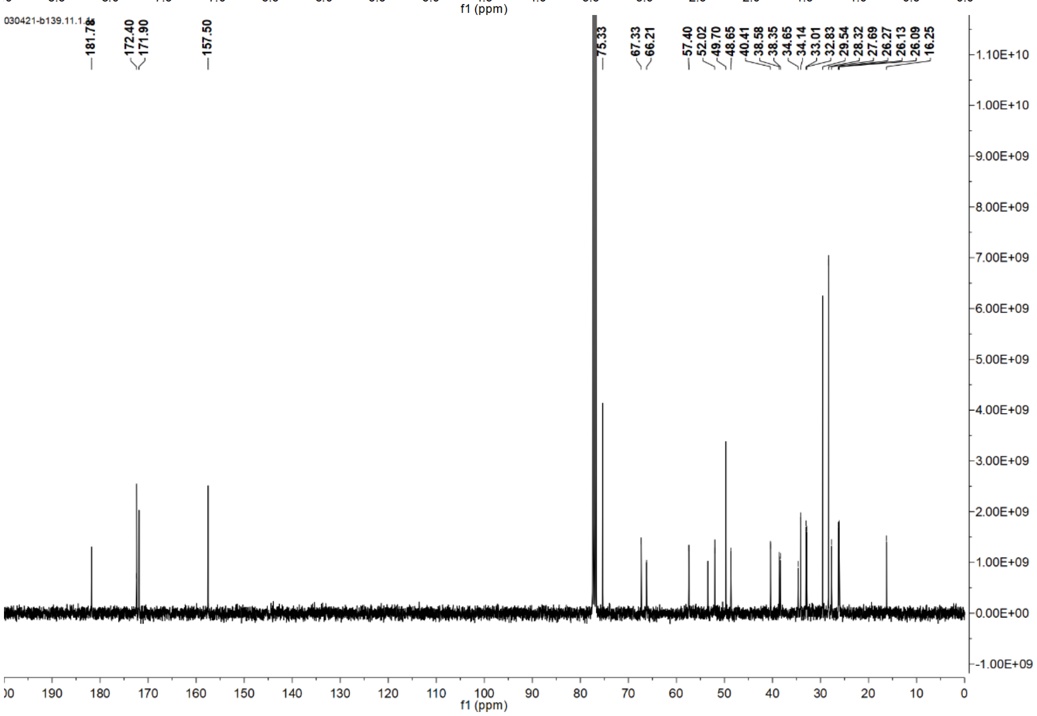
030421-b139.11.1
181.78
172.40
171.90
157.50
75.33
67.33
66.21
57.40
52.02
49.70
48.65
40.41
38.58
38.35
34.65
34.14
33.01
32.83
29.54
28.32
27.69
26.27
26.13
26.09
16.25
f1 (ppm)

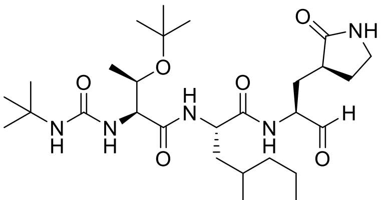
O
NH
O
H
N
H
N
H
O
N
H
O
N
H
H
O

**MPI45**

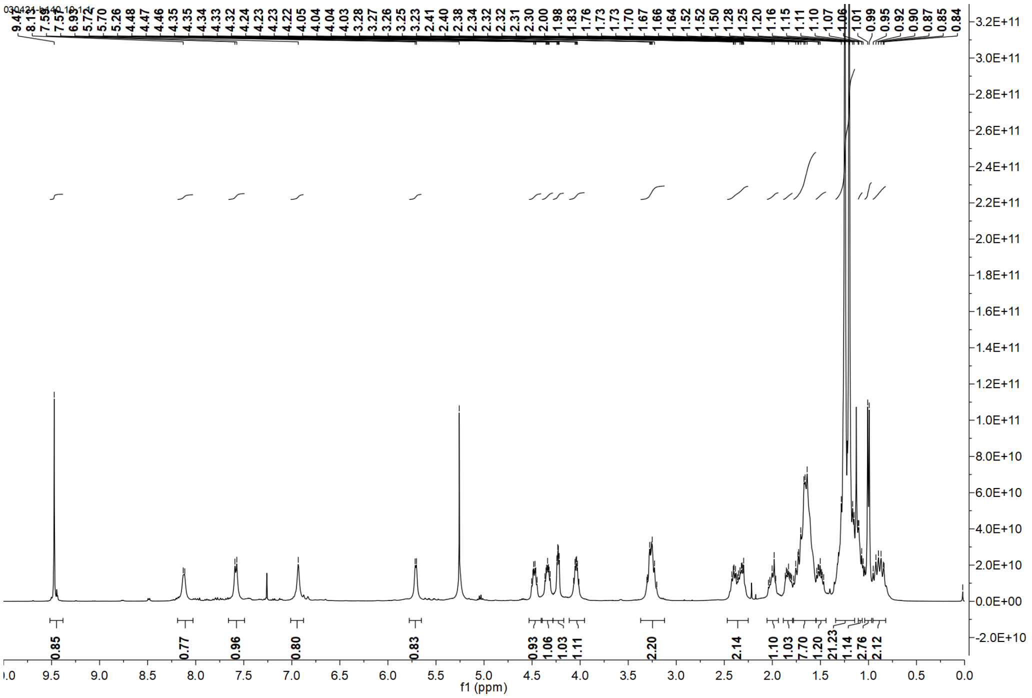
0.85
0.77
0.96
0.80
0.83
0.93
1.06
1.03
1.11
2.20
2.14
1.10
1.03
7.70
1.20
21.23
1.14
2.76
2.12
f1 (ppm)

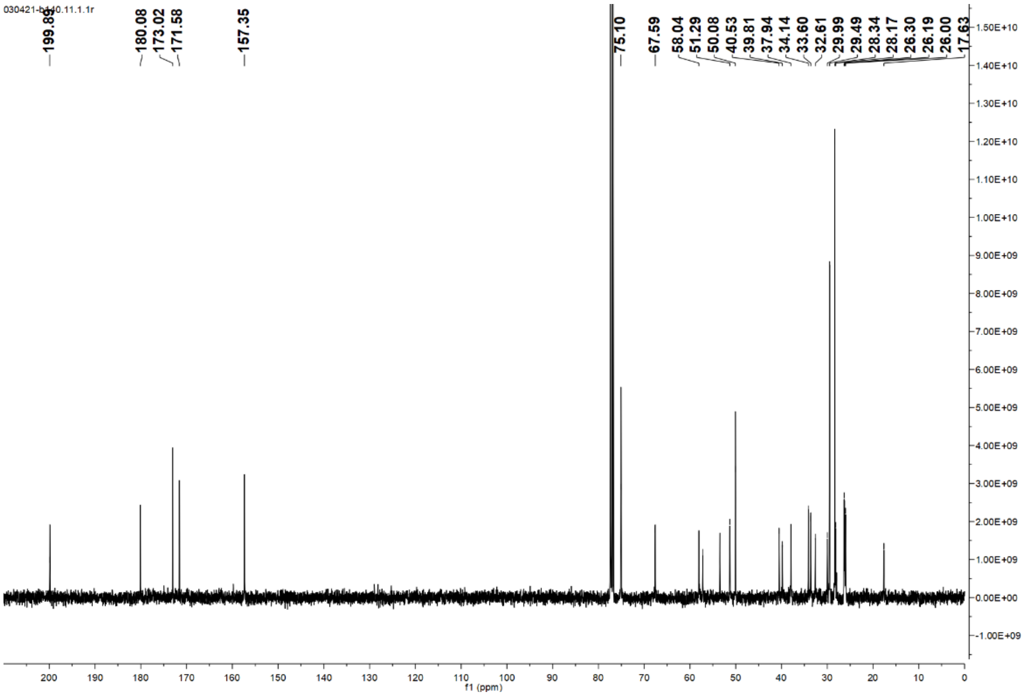
199.89
180.08
173.02
171.58
157.35
75.10
67.59
58.04
51.29
50.08
40.53
39.81
37.94
34.14
33.60
32.61
29.99
29.49
28.34
28.17
26.30
26.19
26.00
17.63
f1 (ppm)

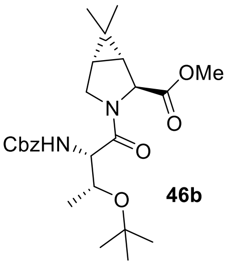
OMe
CbzHN
O
O
N

**46b**

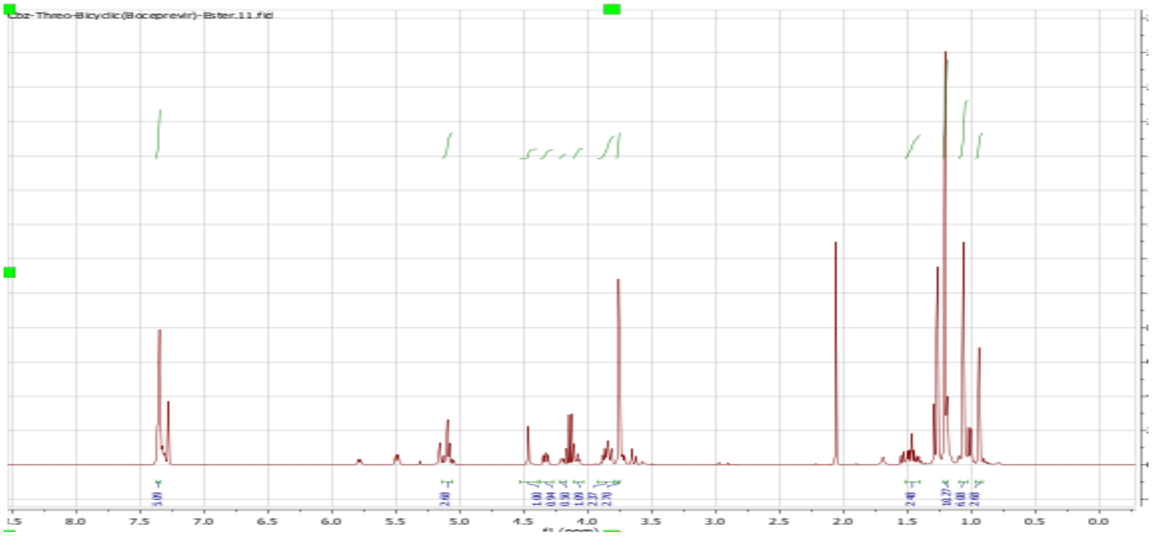
Cbz-Threo-Bicyclic(Boceprevir)-Ester.11.fid

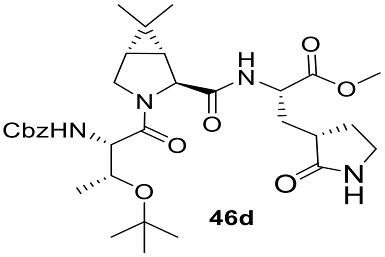
CbzHN
H
N
O
O
NH

**46d**

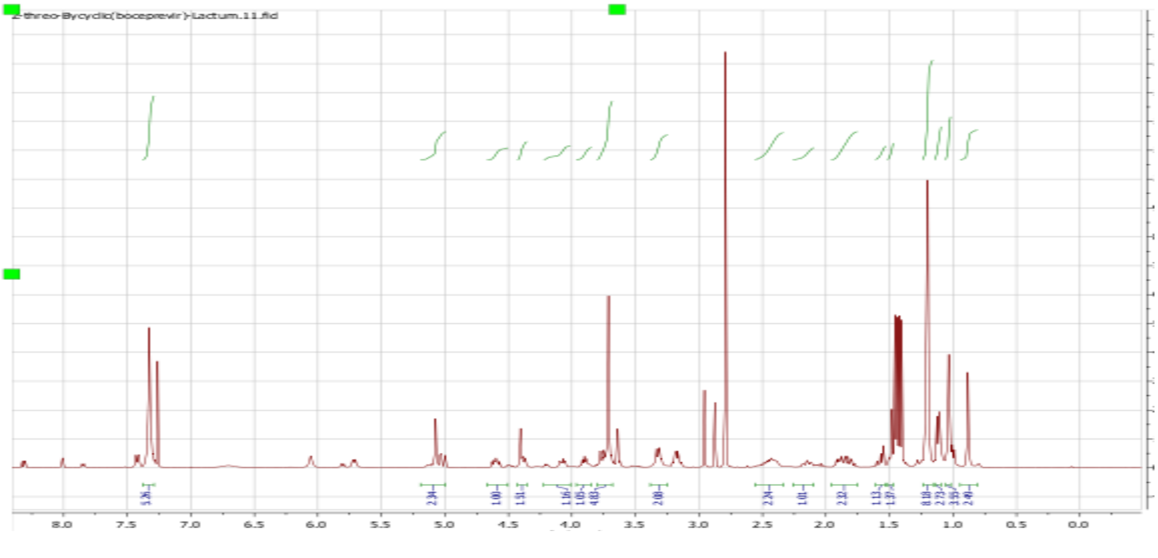
Z-threo-Bycyclic(boceprevir)-Lactum.11.fid

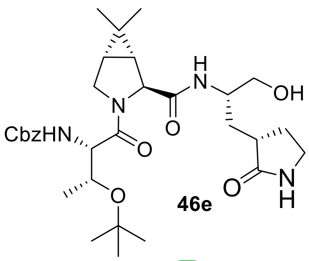
CbzHN
N
O
H
N
OH
O
NH
O
H
O
46e

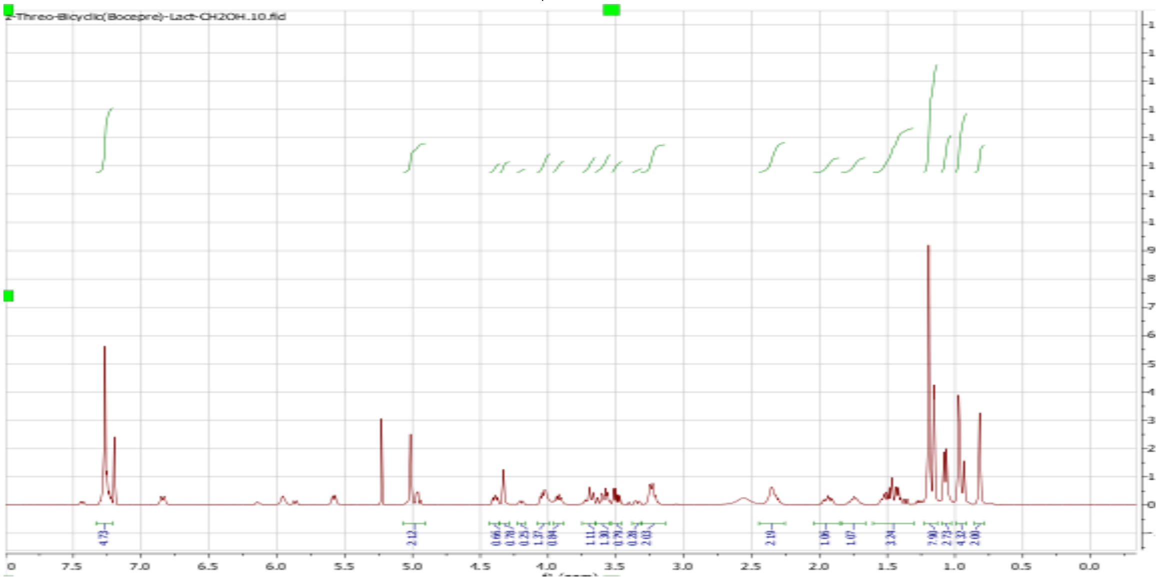
Z-Threo-Bicyclic(Bocepre)-Lact-CH2OH.10.fid
4.73
2.12
2.19
1.06
1.07
3.24
7.90
2.73
4.32
2.00
7.5
7.0
6.5
6.0
5.5
5.0
4.5
4.0
3.5
3.0
2.5
2.0
1.5
1.0
0.5
0.0

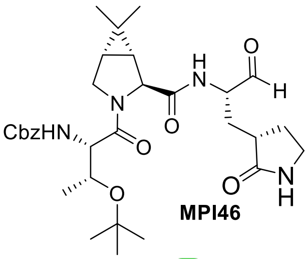


**MPI46**

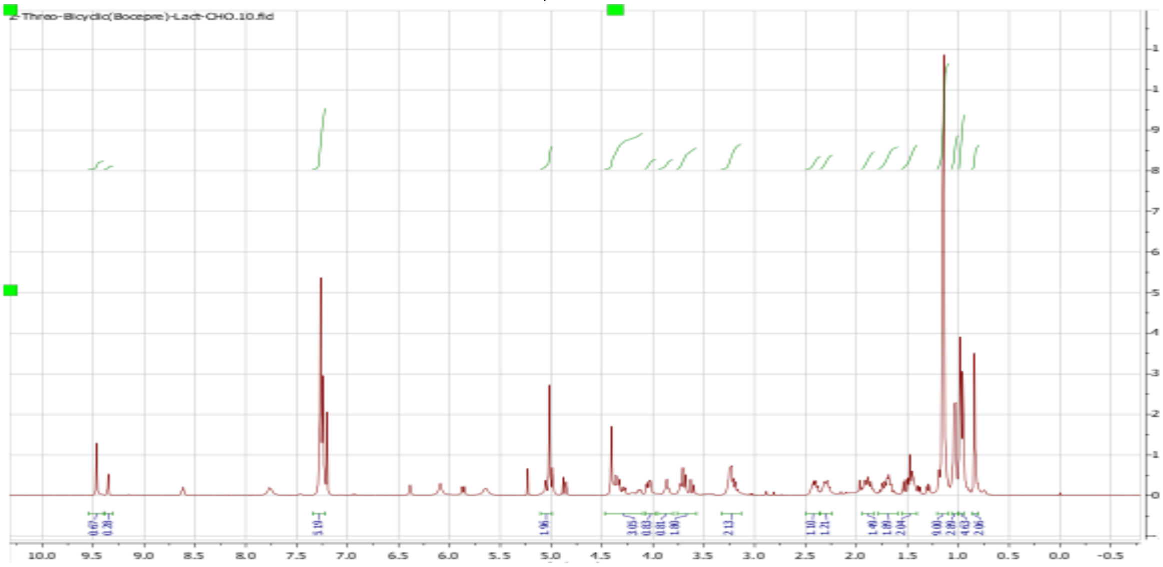


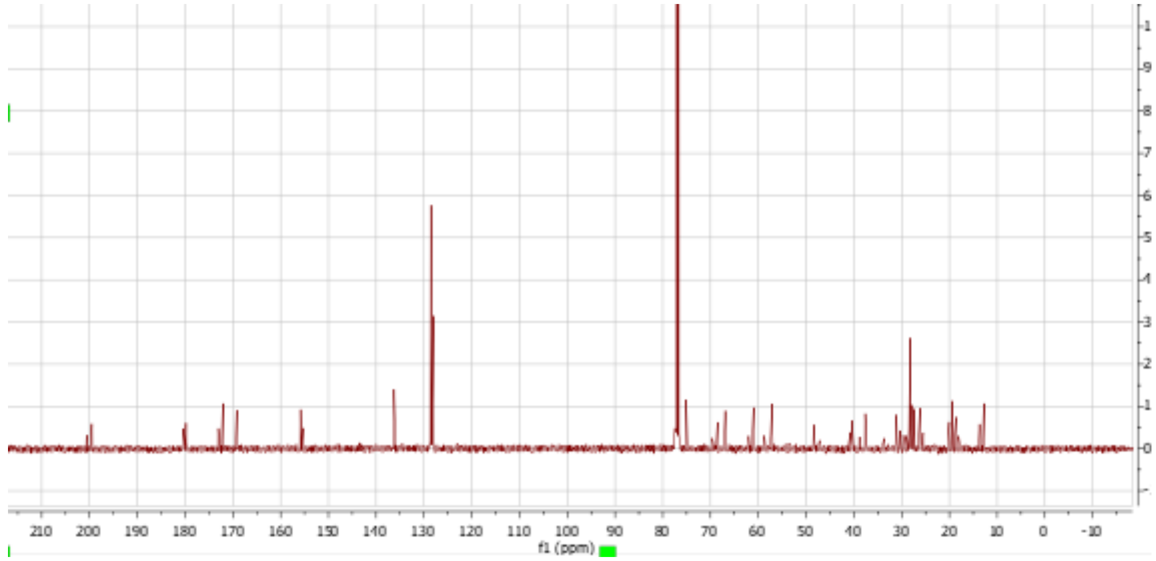

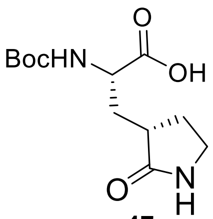
BocHN
O
OH
O
N
H

**47a**

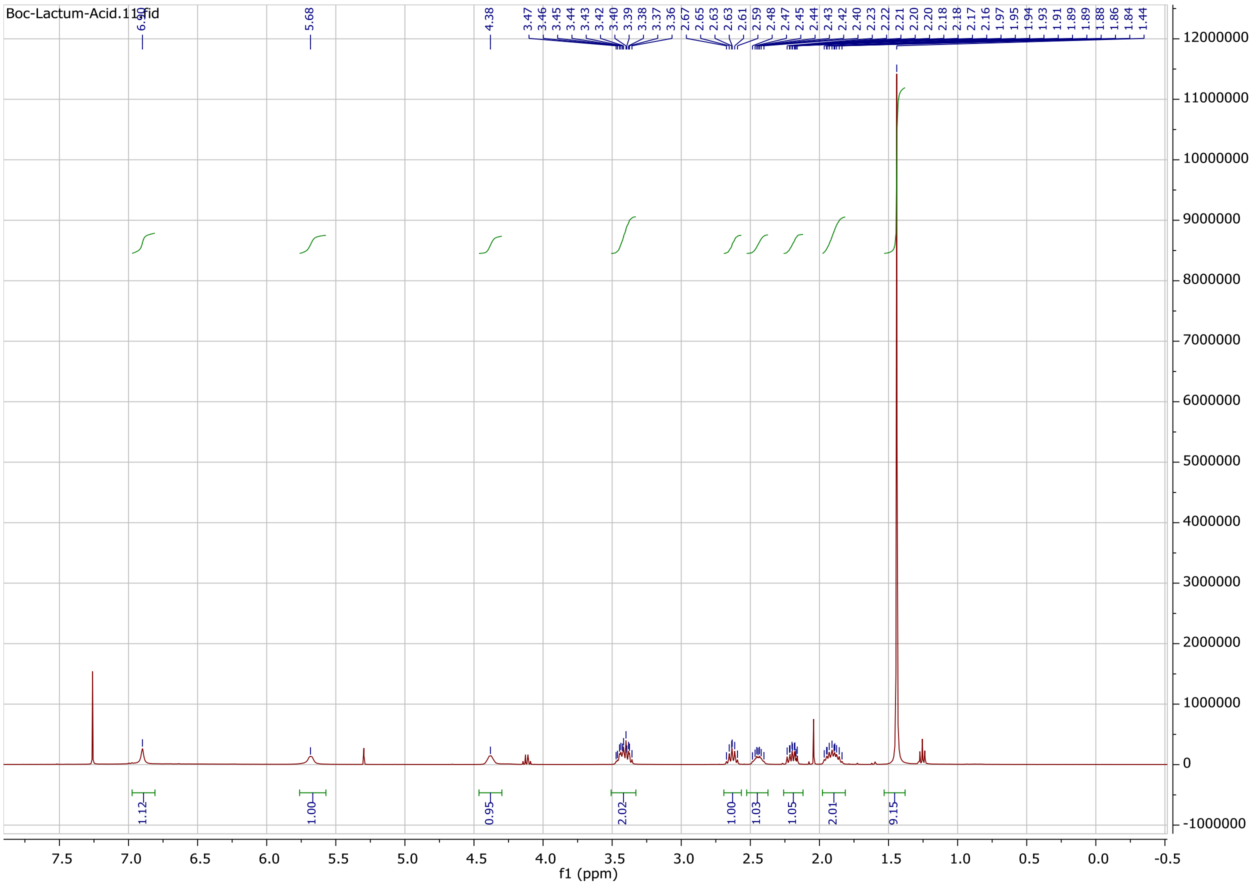
Boc-Lactum-Acid.11.fid
5.68
4.38
3.47 3.46 3.45 3.44 3.43 3.42 3.40 3.39 3.38 3.37 3.36 2.67 2.65 2.63 2.63 2.61 2.59 2.48 2.47 2.45 2.44 2.43 2.42 2.40 2.23 2.22 2.21 2.20 2.20 2.18 2.18 2.17 2.16 1.97 1.95 1.94 1.93 1.91 1.89 1.89 1.88 1.86 1.84 1.44
1.12
1.00
0.95
2.02
1.00
1.03
1.05
2.01
9.15
12000000
11000000
10000000
9000000
8000000
7000000
6000000
5000000
4000000
3000000
2000000
1000000
0
-1000000
7.5 7.0 6.5 6.0 5.5 5.0 4.5 4.0 3.5 3.0 2.5 2.0 1.5 1.0 0.5 0.0 -0.5
f1 (ppm)

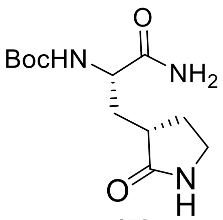


**47b**

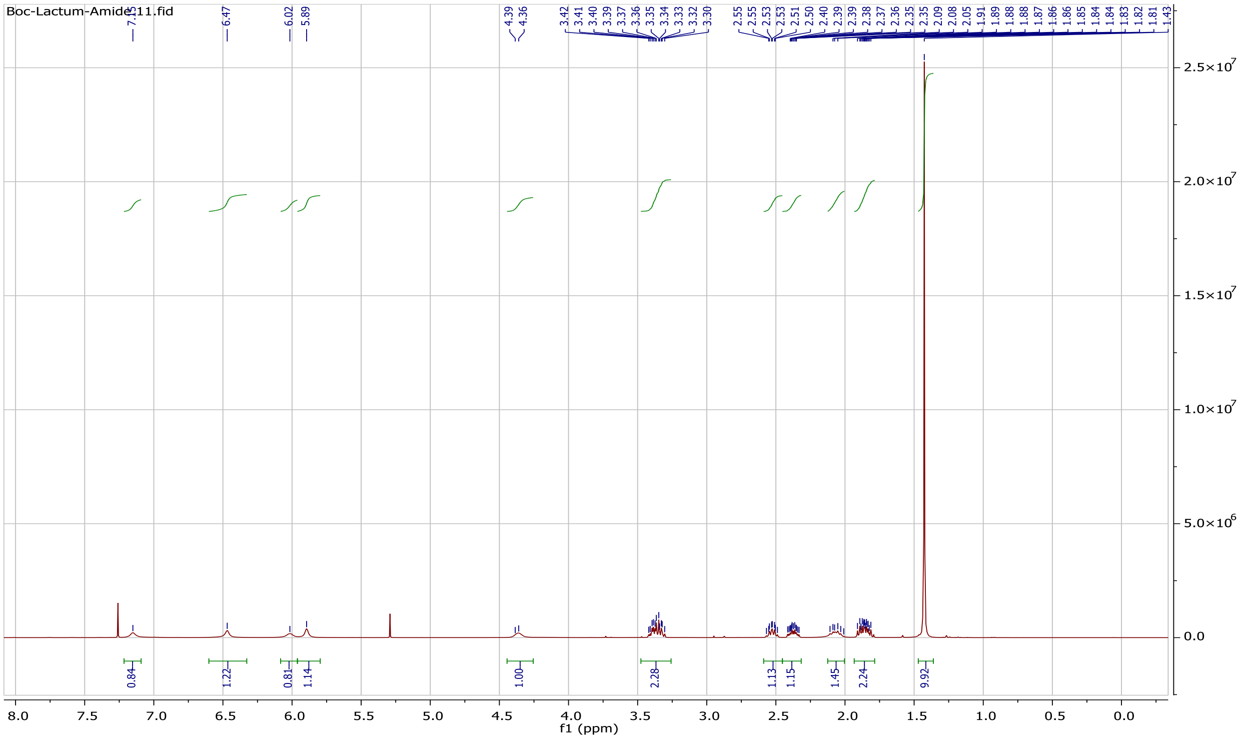

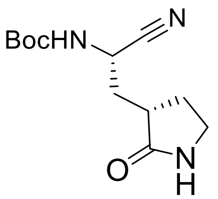

**47c**

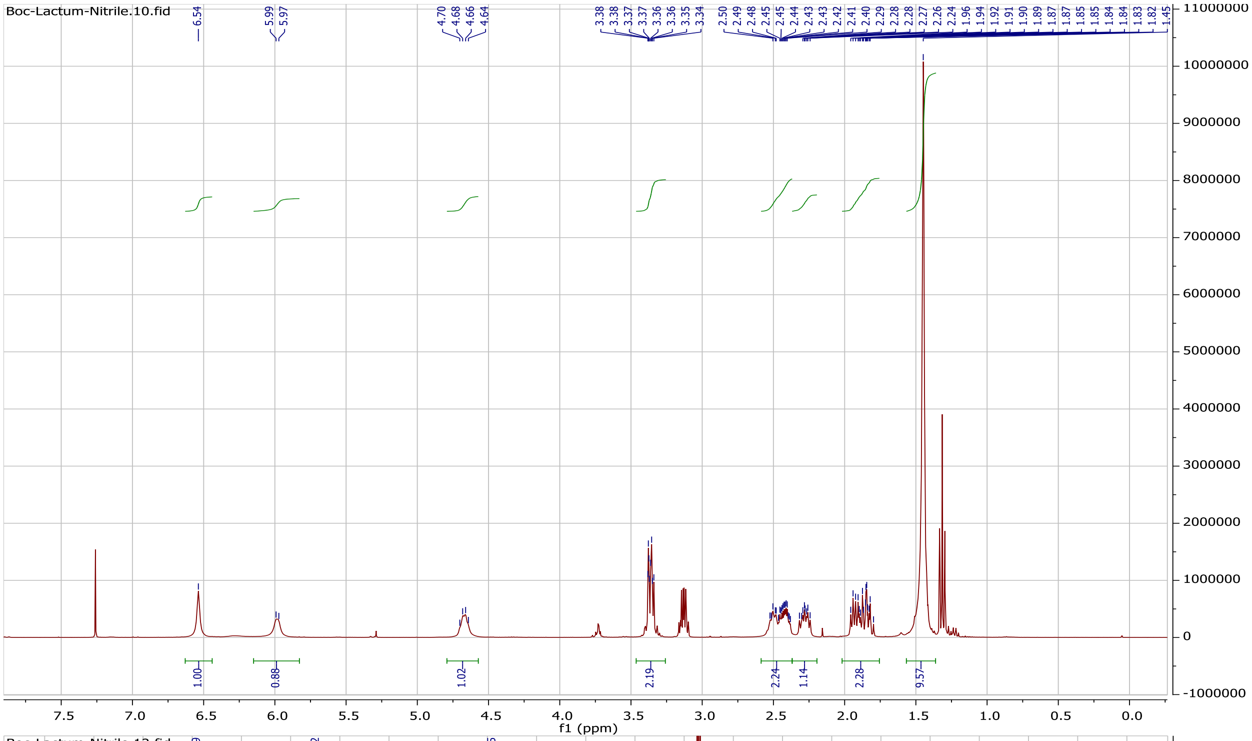
Boc-Lactum-Nitrile.10.fid
f1 (ppm)

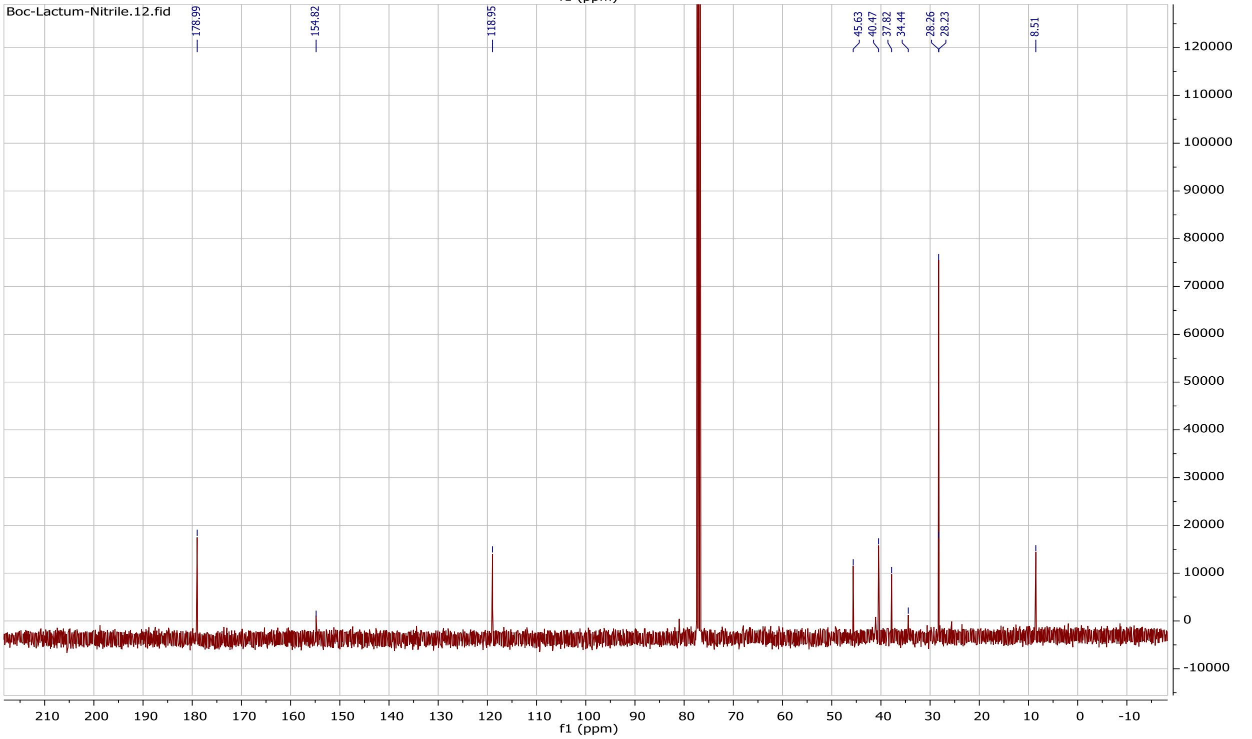
Boc-Lactum-Nitrile.12.fid
178.99
154.82
118.95
45.63
40.47
37.82
34.44
28.26
28.23
8.51
f1 (ppm)

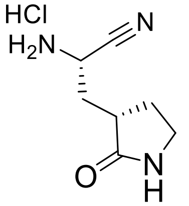


**47d**

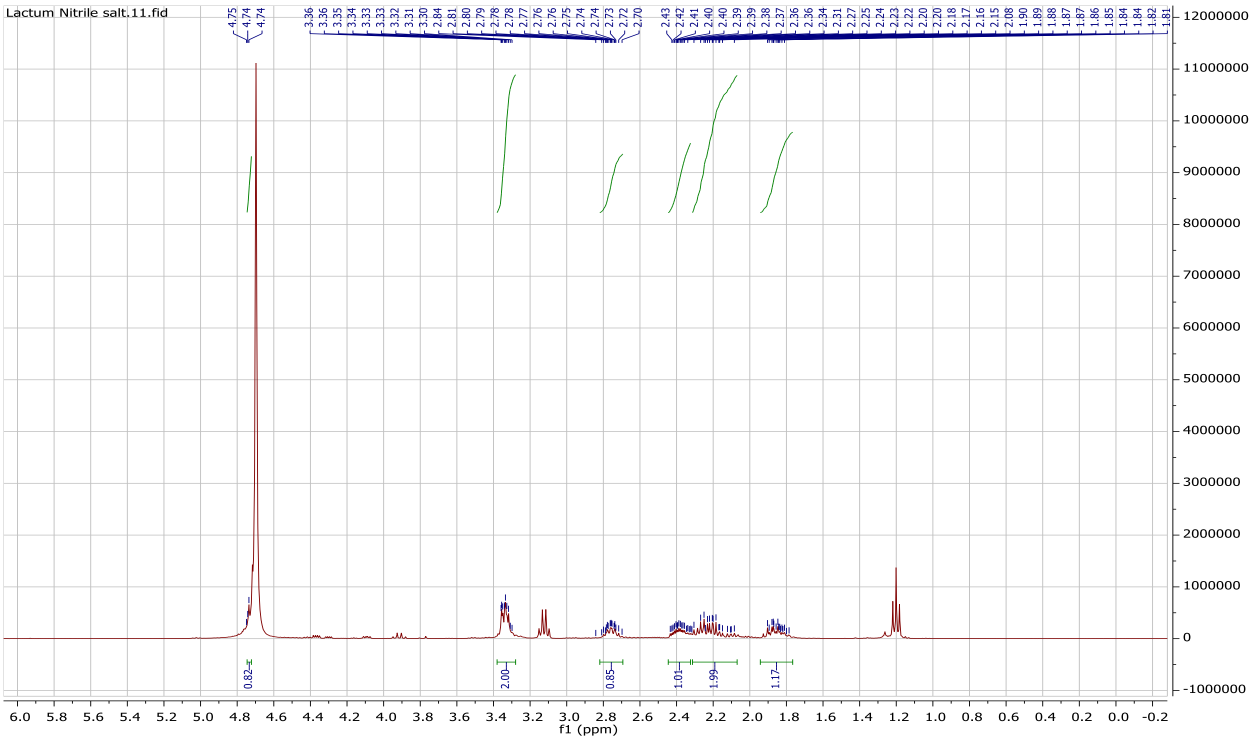

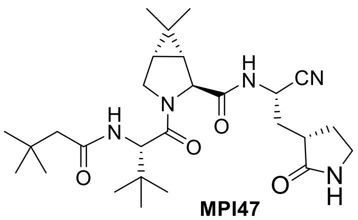

**MPI47**

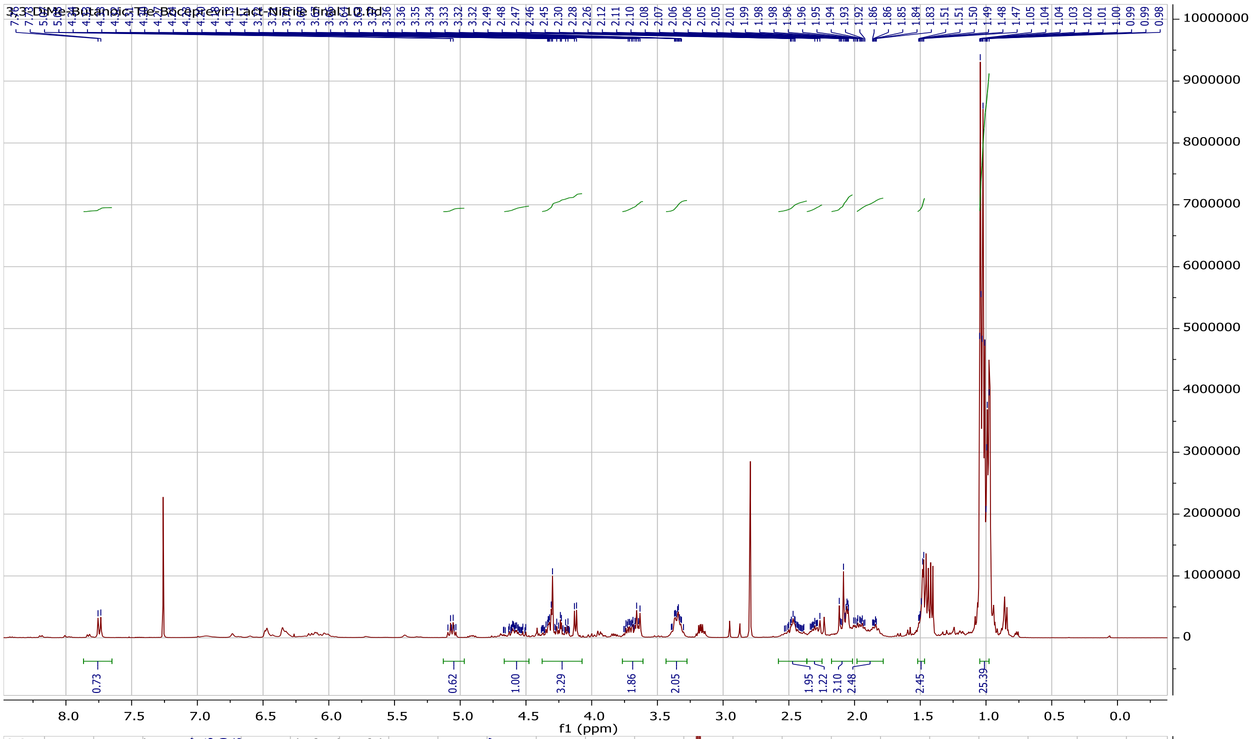


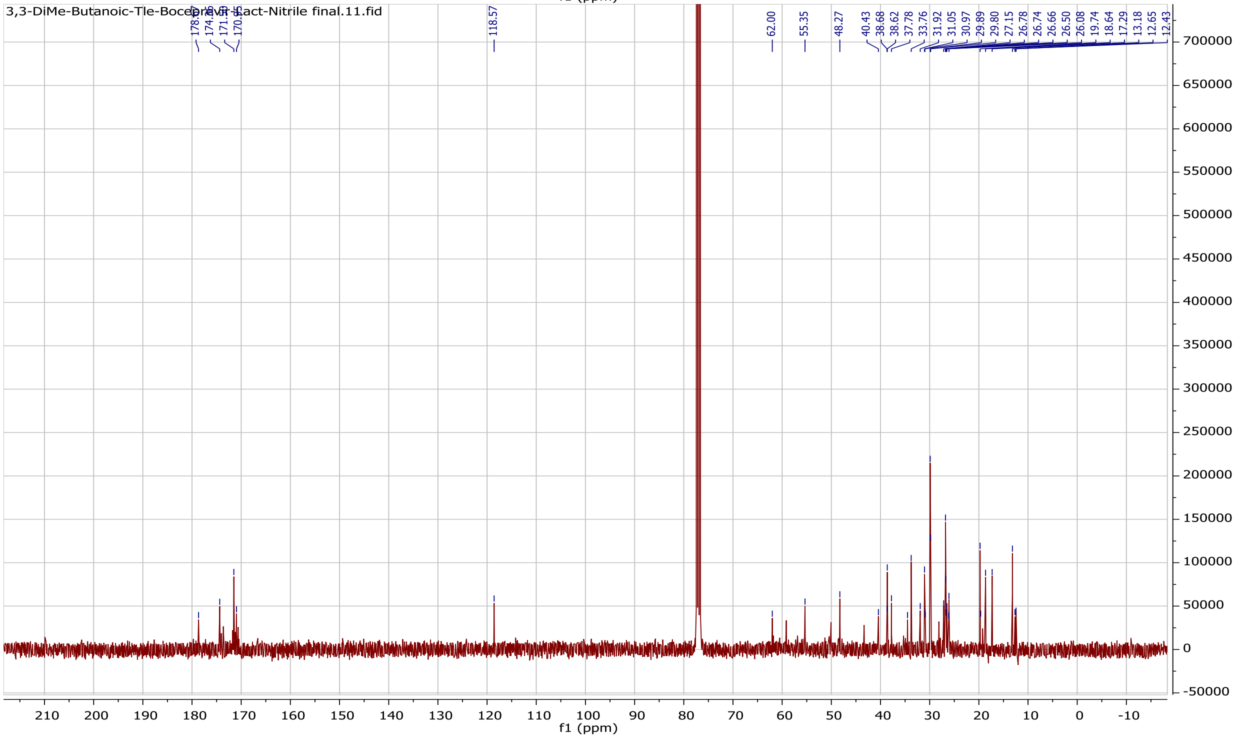

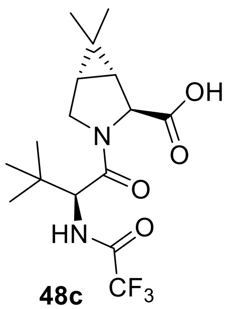

**48c**

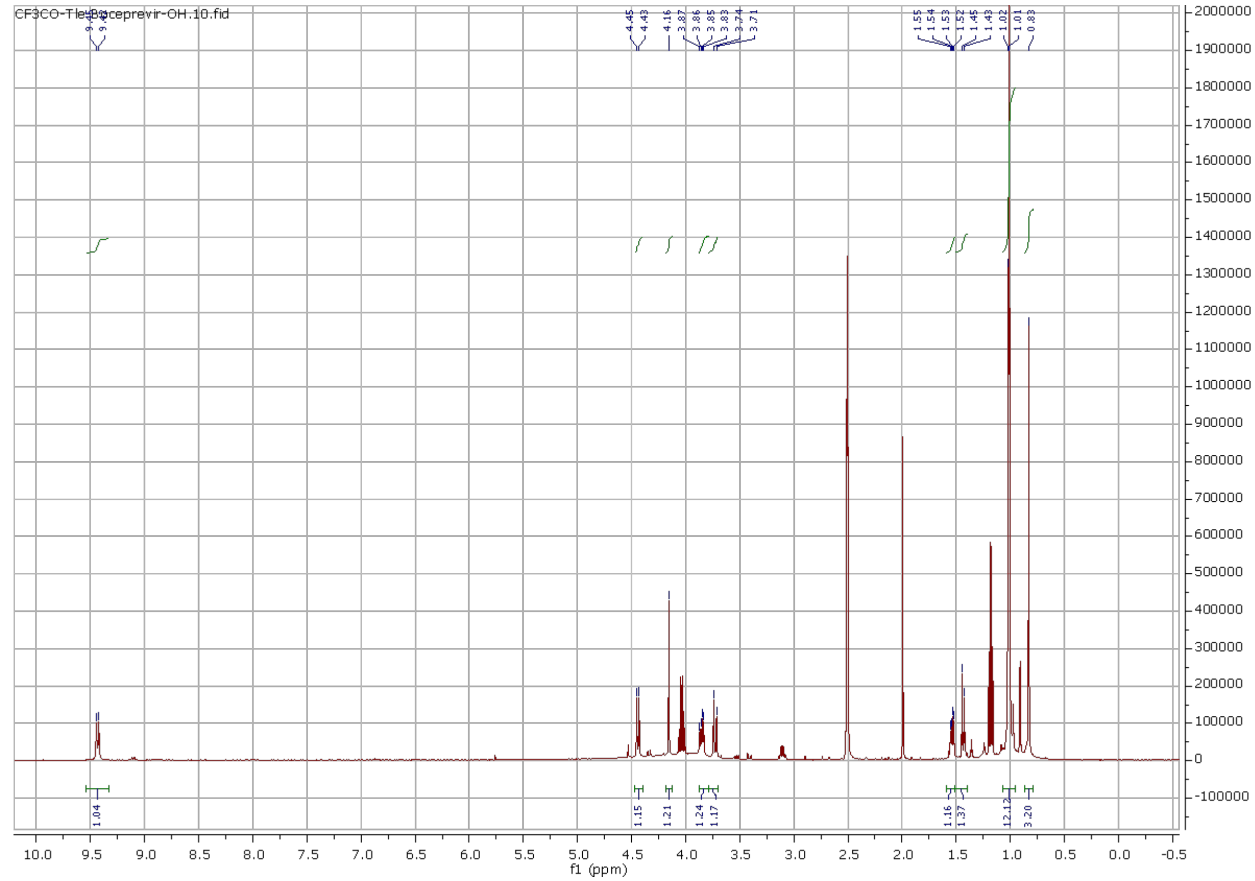
CF3CO-Tle-Boceprevir-OH.10.fid
f1 (ppm)
1.04
1.15
1.21
1.24
1.17
1.16
1.37
12.12
3.20

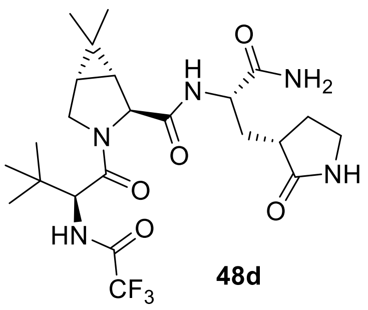

**48d**

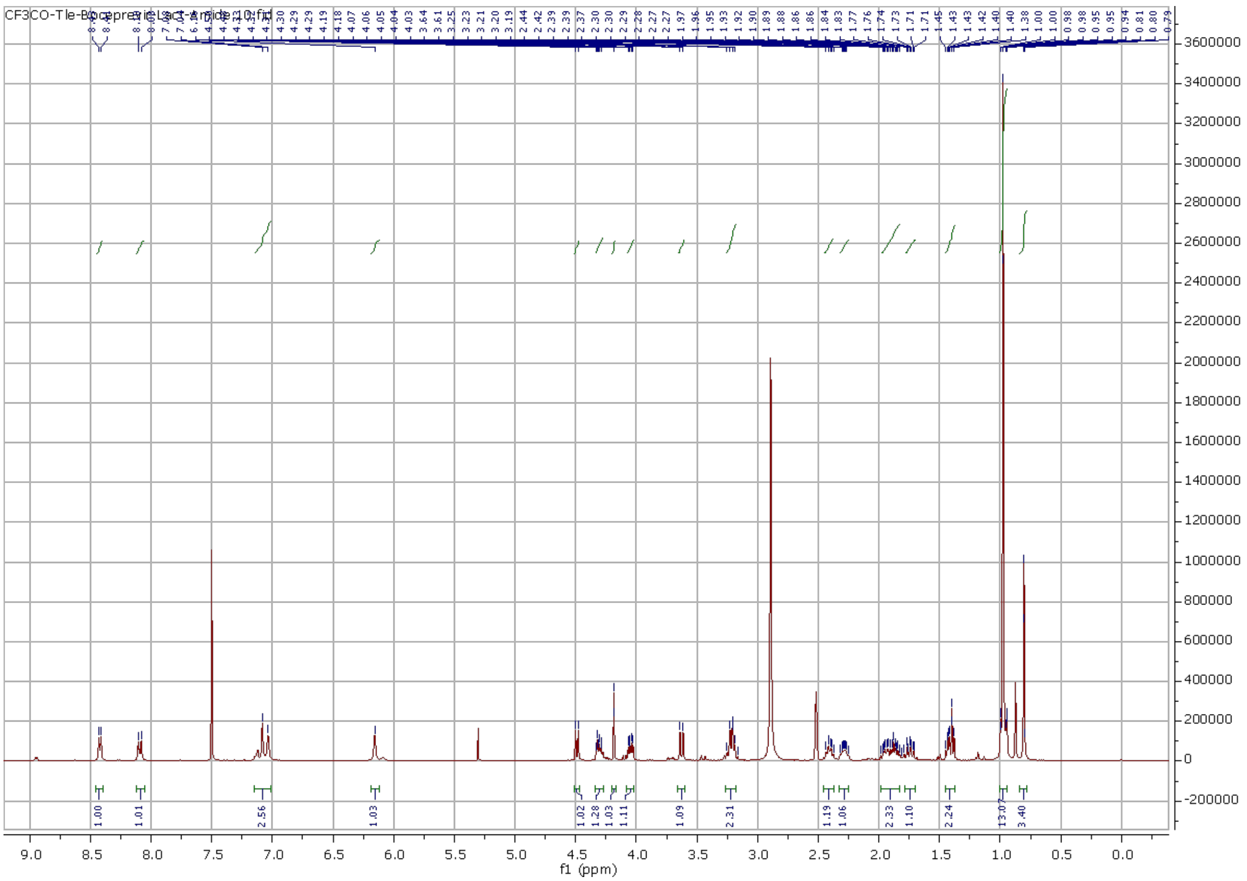

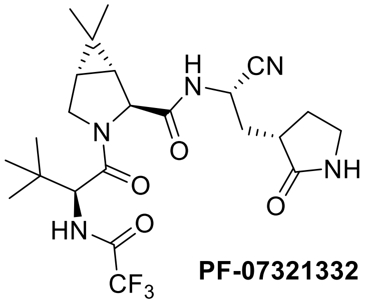

**PF-07321332**

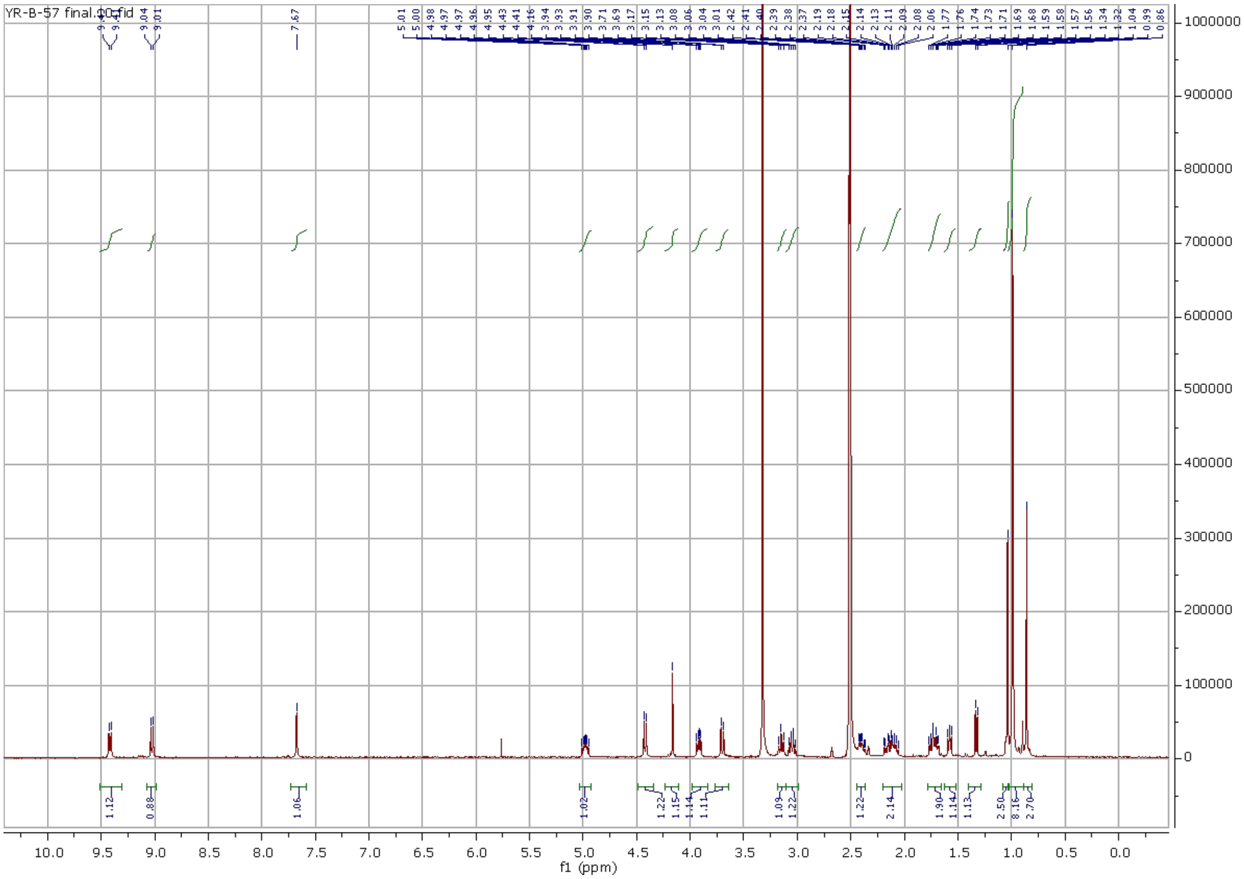

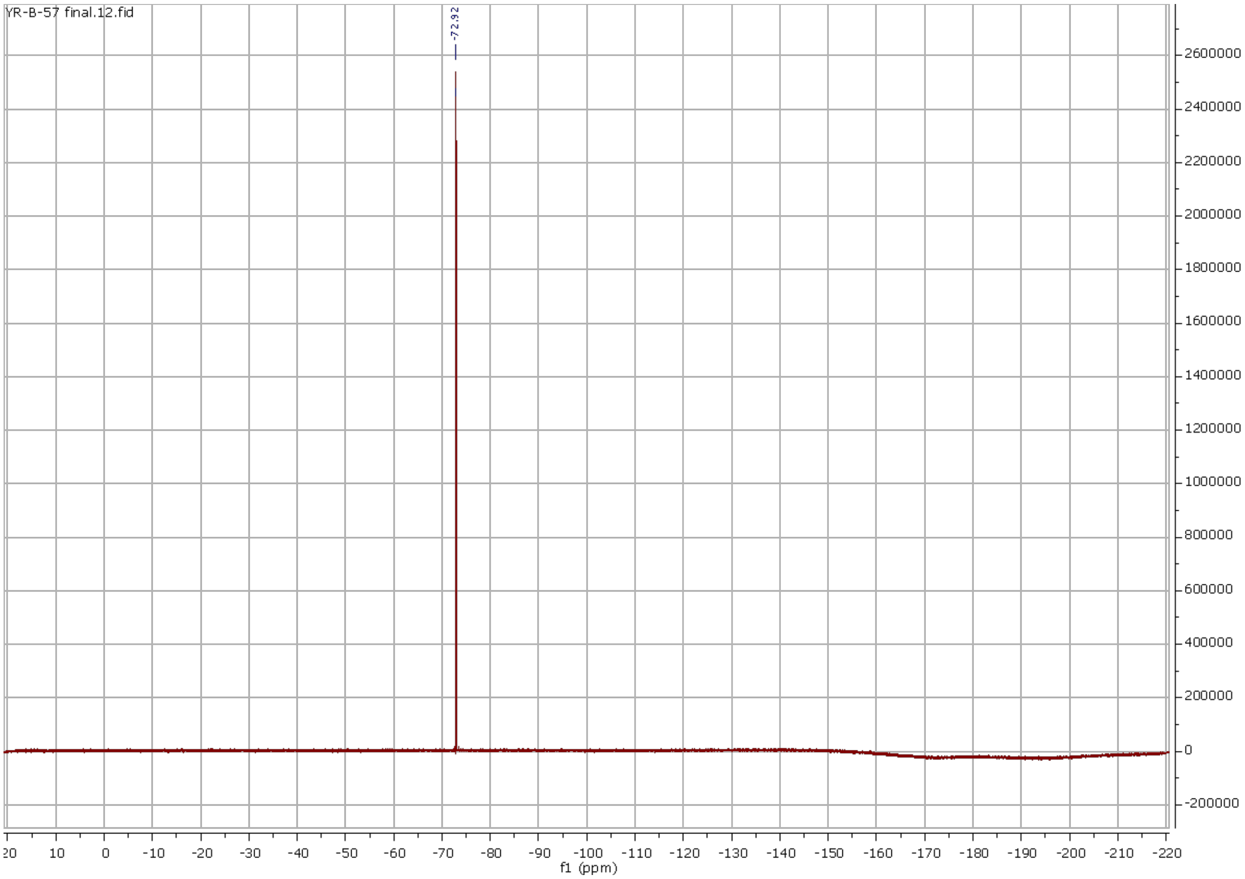
YR-B-57 final.12.fid
-72.92
f1 (ppm)

### HPLC Chromatographies of MPIs

#### MPI 40

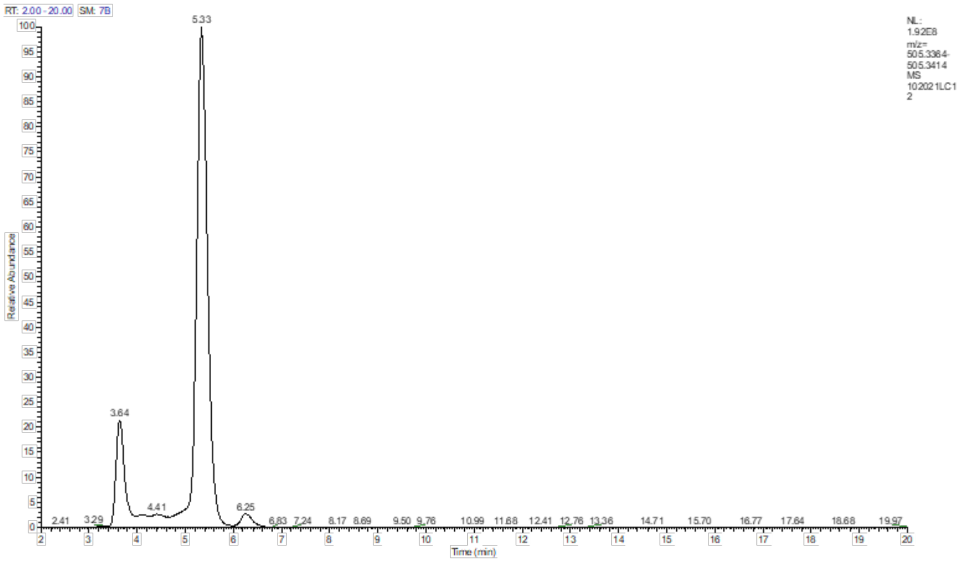


| | | | | | | |
|---|---|---|---|---|---|---|
| 1 | **Sample** | **MPI-40** | | | | |
| 2 | | | | | | |
| 3 | PEAK LIST | | | | | |
| 4 | 102021LC12.raw | | | | | |
| 5 | RT: 0.00 - 20.00 | | | | | |
| 6 | Number of detected peaks: 3 | | | | | |
| 7 | Apex RT | Start RT | End RT | Area | %Area | **Purity** |
| 8 | 3.64 | 3.48 | 4.18 | 459484399 | 13.4 | **13%** |
| 9 | 5.33 | 4.77 | 5.74 | 2883153622 | 84.3 | **84%** |
| 10 | 6.25 | 6.02 | 6.73 | 78381597 | 2.3 | |

**Experimental Condition**

| No | Time | Flow [ml/min] | %B | %C | %D | Curve |
|---|---|---|---|---|---|---|
| **1** | 0.000 | Equilibration | | | | |
| **2** | 0.000 | 0.300 | 0.0 | 70.0 | 0.0 | 5 |
| **3** | *New Row* | | | | | |
| **4** | 0.000 | Run | | | | |
| **5** | 5.000 | 0.300 | 0.0 | 70.0 | 0.0 | 5 |
| **6** | *New Row* | | | | | |
| **7** | 20.000 | Stop Run | | | | |

**Note:** two peaks ($t_R$ = 3.64 min and $t_R$ = 5.33 min) are diastereomers.

### MPI 43

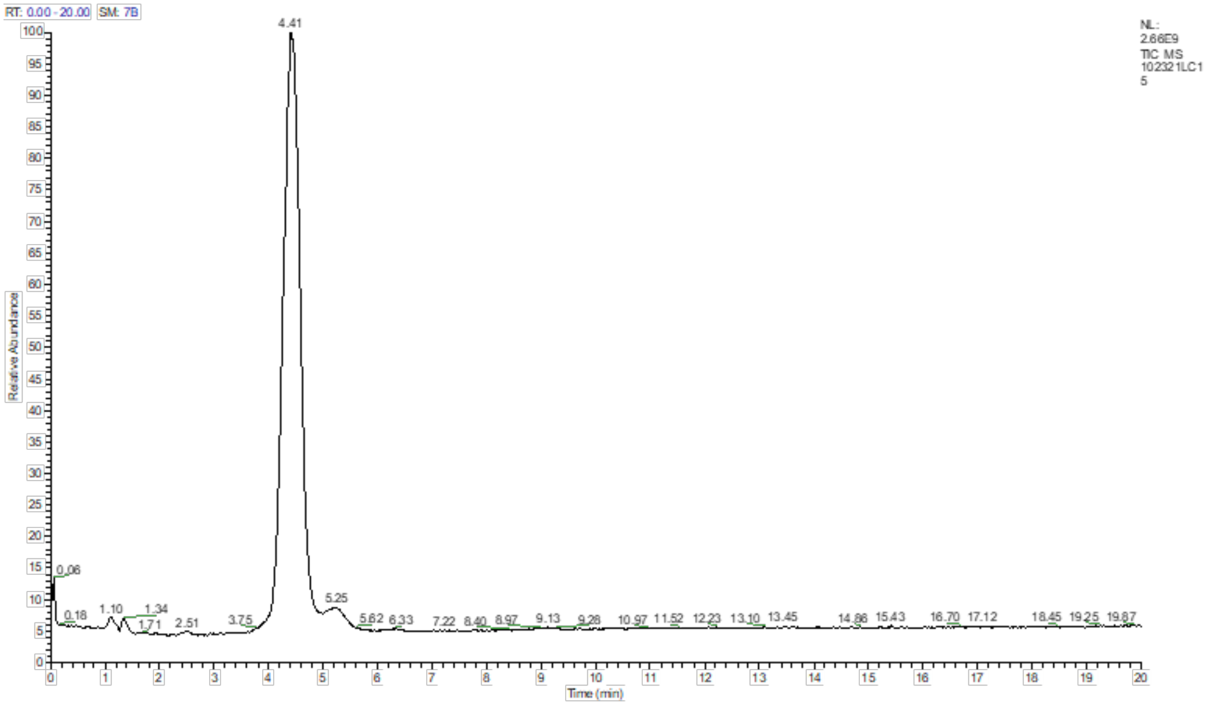


| | | | | | | |
|---|---|---|---|---|---|---|
| 1 | **Sample ID** | **MPI43** | | | | |
| 2 | | | | | | |
| 3 | PEAK LIST | | | | | |
| 4 | 102321LC15.raw | | | | | |
| 5 | RT: 0.00 - 20.00 | | | | | |
| 6 | Number of detected peaks: 3 | | | | | |
| 7 | Apex RT | Start RT | End RT | Area | %Area | **PURITY** |
| 8 | 2.51 | 2.32 | 2.69 | 1.82E+08 | 0.31 | |
| 9 | 4.41 | 3.78 | 5.01 | 5.65E+10 | 94.85 | **95%** |
| 10 | 5.25 | 5.02 | 5.85 | 2.89E+09 | 4.85 | |

#### Experimental Condition

| No | Time | Flow [ml/min] | %B | %C | %D | Curve |
|---|---|---|---|---|---|---|
| 1 | 0.000 | Equilibration | | | | |
| 2 | 0.000 | 0.300 | 0.0 | 70.0 | 0.0 | 5 |
| 3 | *New Row* | | | | | |
| 4 | 0.000 | Run | | | | |
| 5 | 5.000 | 0.300 | 0.0 | 70.0 | 0.0 | 5 |
| 6 | *New Row* | | | | | |
| 7 | 20.000 | Stop Run | | | | |

### MPI 44

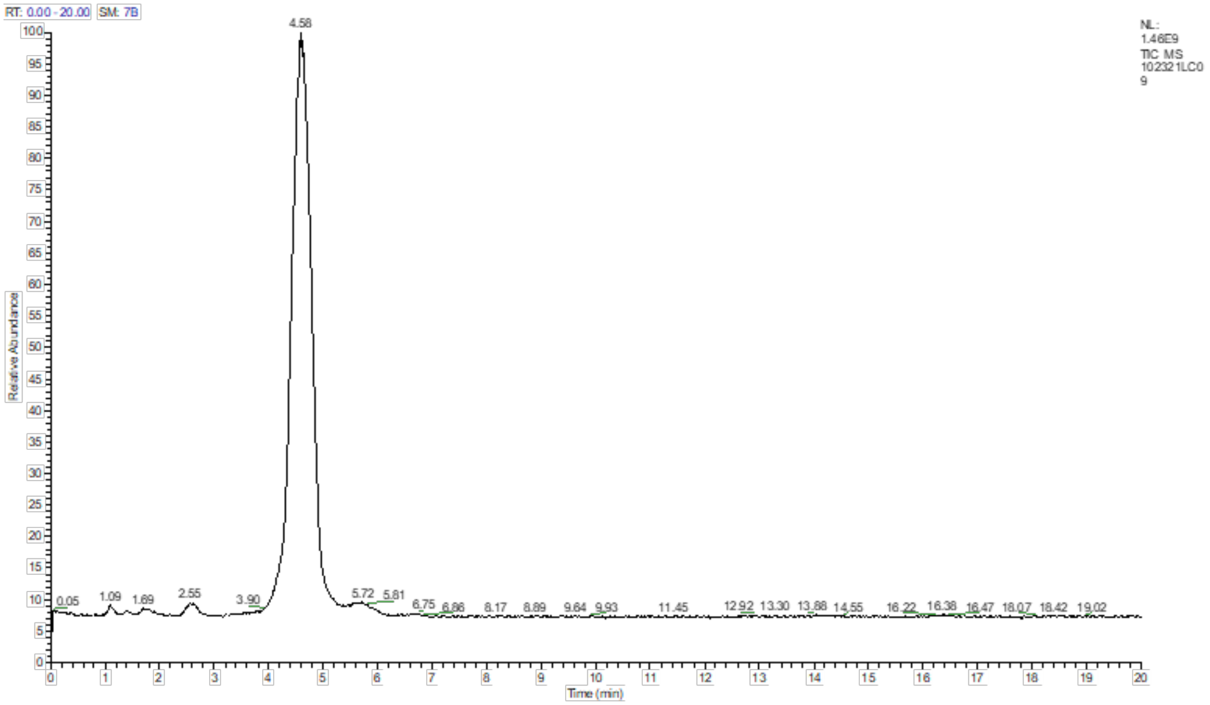


| | | | | | | |
|---|---|---|---|---|---|---|
| 1 | **Sample** | **MPI44** | | | | |
| 2 | | | | | | |
| 3 | PEAK LIST | | | | | |
| 4 | 102321LC09.raw | | | | | |
| 5 | RT: 0.00 - 20.00 | | | | | |
| 6 | Number of detected peaks: 3 | | | | | |
| 7 | Apex RT | Start RT | End RT | Area | %Area | **PURITY** |
| 8 | 2.55 | 2.39 | 2.79 | 3.82E+08 | 1.08 | |
| 9 | 4.58 | 3.93 | 5.34 | 3.42E+10 | 96.04 | **96%** |
| 10 | 5.72 | 5.34 | 6.25 | 1.03E+09 | 2.88 | |

**Experimental Condition**

| No | Time | Flow [ml/min] | %B | %C | %D | Curve |
|---|---|---|---|---|---|---|
| 1 | 0.000 | Equilibration | | | | |
| 2 | 0.000 | 0.300 | 0.0 | 70.0 | 0.0 | 5 |
| 3 | *New Row* | | | | | |
| 4 | 0.000 | Run | | | | |
| 5 | 5.000 | 0.300 | 0.0 | 70.0 | 0.0 | 5 |
| 6 | *New Row* | | | | | |
| 7 | 20.000 | Stop Run | | | | |

### MPI 46

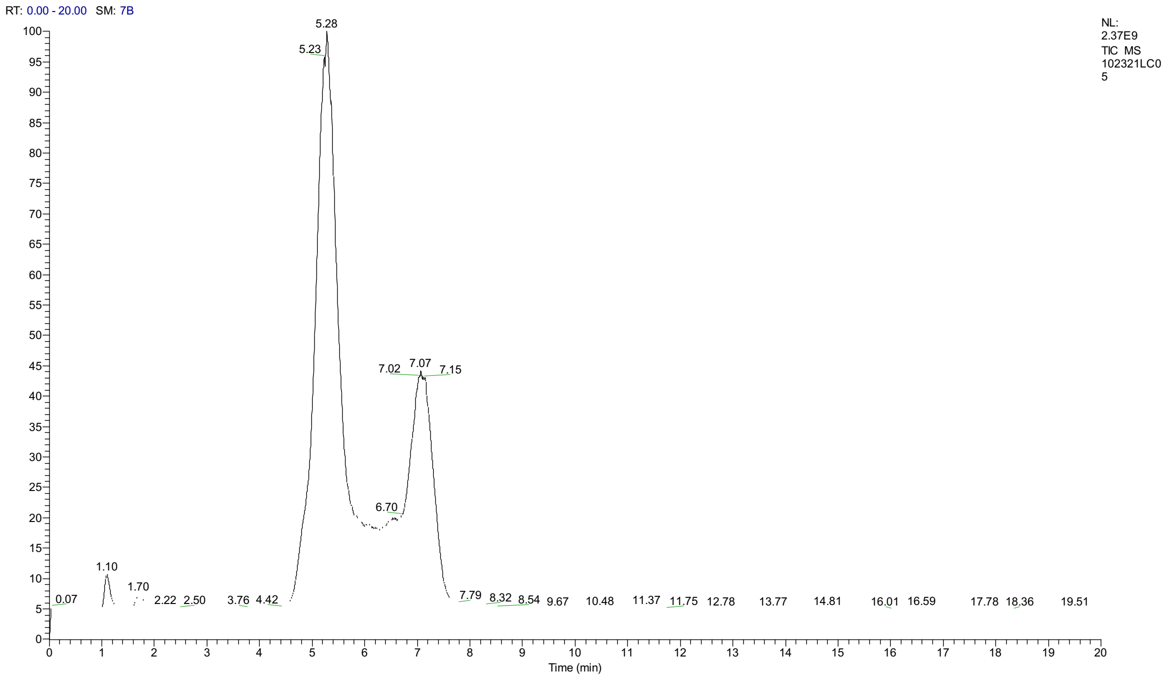


| | | | | | | |
|---|---|---|---|---|---|---|
| 1 | **Sample ID** | **MPI46** | | | | |
| 2 | | | | | | |
| 3 | PEAK LIST | | | | | |
| 4 | 102321LC05.raw | | | | | |
| 5 | RT: 0.00 - 20.00 | | | | | |
| 6 | Number of detected peaks: 2 | | | | | |
| 7 | Apex RT | Start RT | End RT | Area | %Area | **PURITY** |
| 8 | 4.9 | 4.55 | 5.07 | 1928030399 | 3.831643 | |
| 9 | 5.28 | 4.05 | 6.28 | 34425896325 | 68.41581 | **68%** |
| 10 | 6.57 | 6.28 | 6.8 | 704564290.1 | 1.400206 | |
| 11 | 7.07 | 6.41 | 7.67 | 13260138847 | 26.35234 | **26%** |

#### Experimental Condition

| No | Time | Flow [ml/min] | %B | %C | %D | Curve |
|---|---|---|---|---|---|---|
| **1** | 0.000 | Equilibration | | | | |
| **2** | 0.000 | 0.300 | 0.0 | 70.0 | 0.0 | 5 |
| **3** | *New Row* | | | | | |
| **4** | 0.000 | Run | | | | |
| **5** | 5.000 | 0.300 | 0.0 | 70.0 | 0.0 | 5 |
| **6** | *New Row* | | | | | |
| **7** | 20.000 | Stop Run | | | | |

**Note:** peaks ($t_R$ = 5.25 min and $t_R$ = 7.07 min) are diastereomers.

**Flow Cytometry Images for MPIs**
(compound concentration labeled)

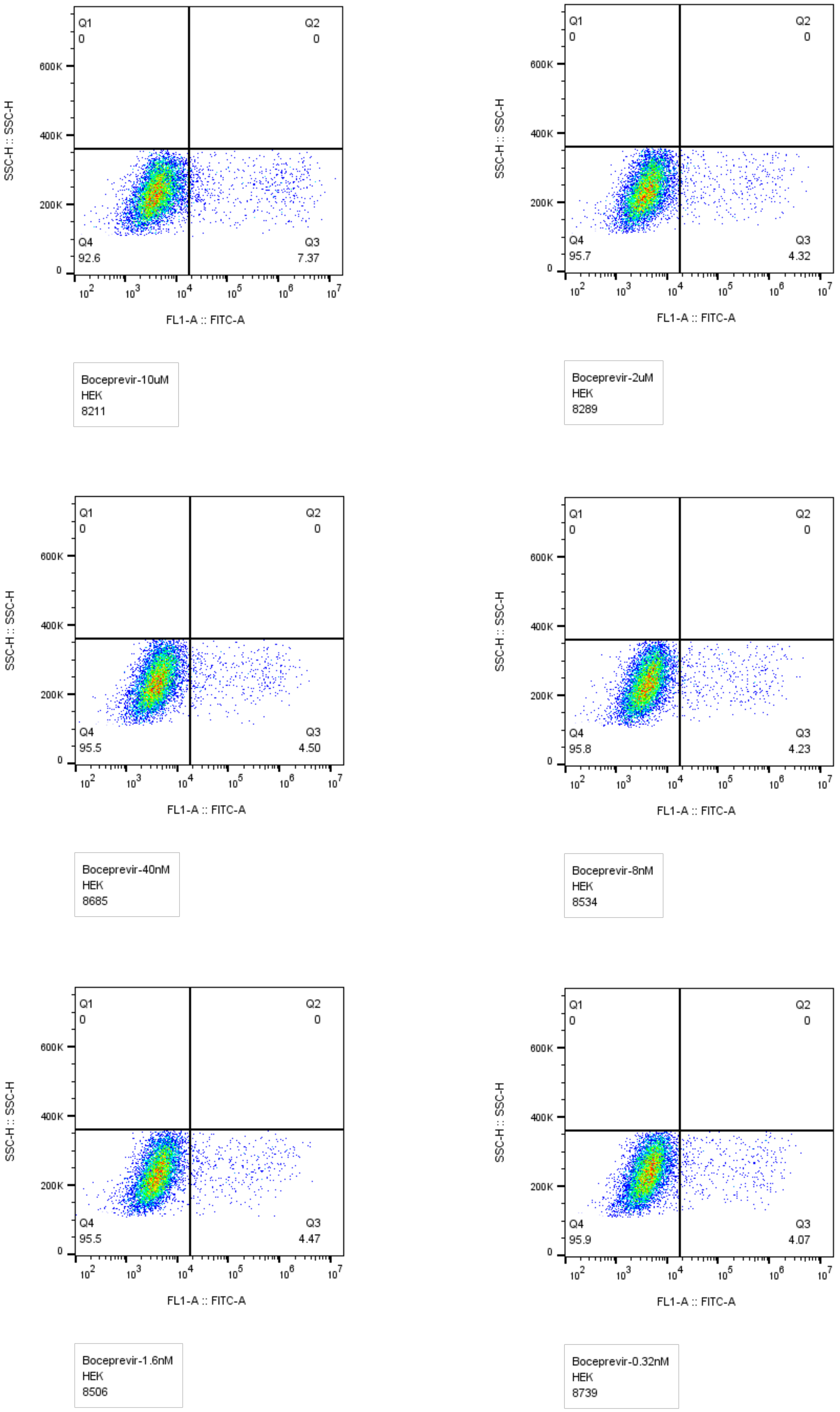


Flow cytometry images for Boceprevir, compound concentration labeled.

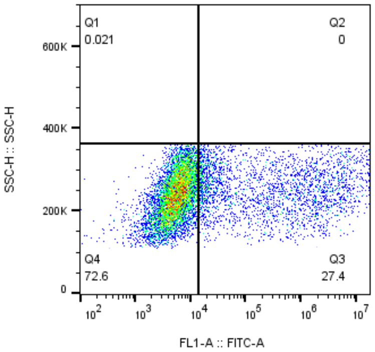


MPI29-10uM
HEK
9706

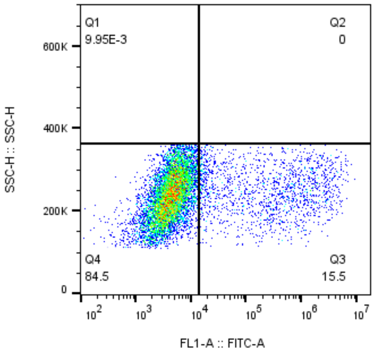


MPI29-2uM
HEK
10050

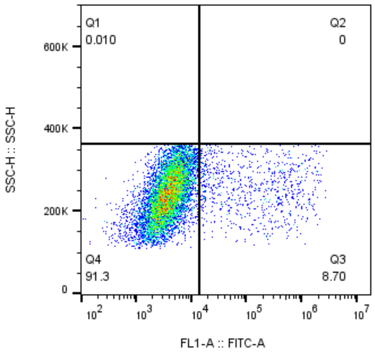


MPI29-40nM
HEK
9751

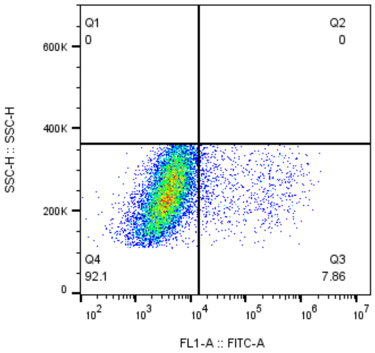


MPI29-8nM
HEK
10042

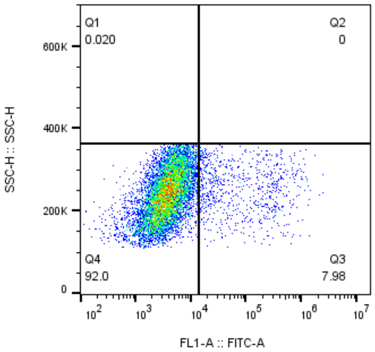


MPI29-1.6nM
HEK
9965

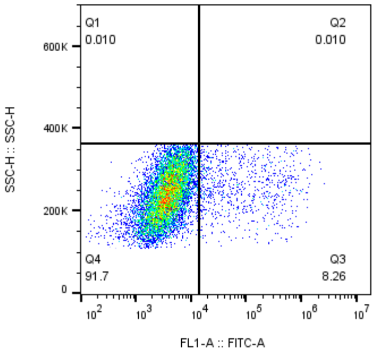


MPI29-0.32nM
HEK
9926

MPI29

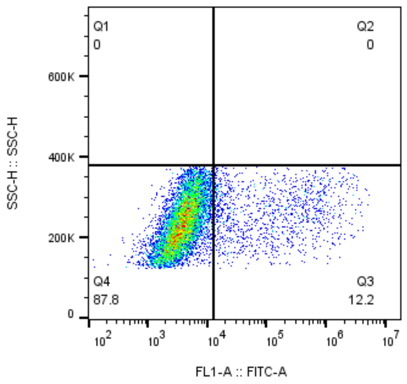


MPI30-10uM
HEK
9760

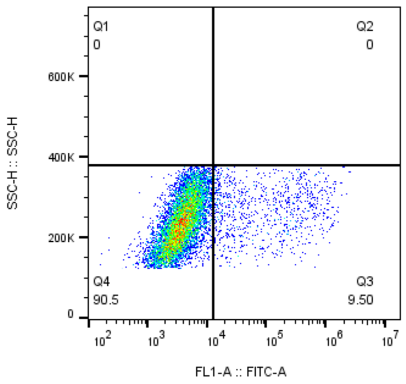


MPI30-2uM
HEK
9831

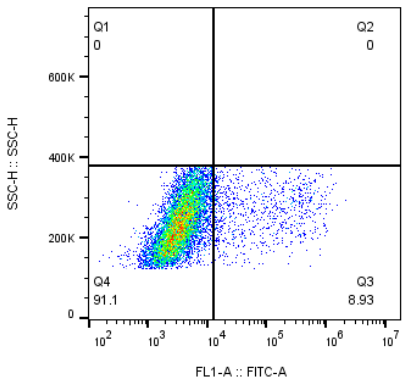


MPI30-40nM
HEK
9687

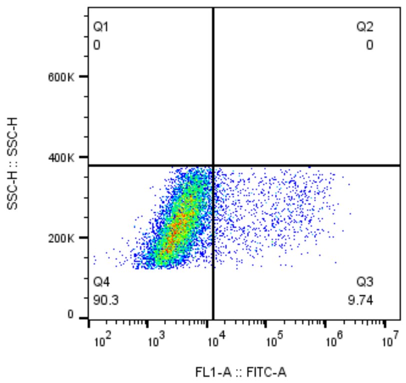


MPI30-8nM
HEK
9912

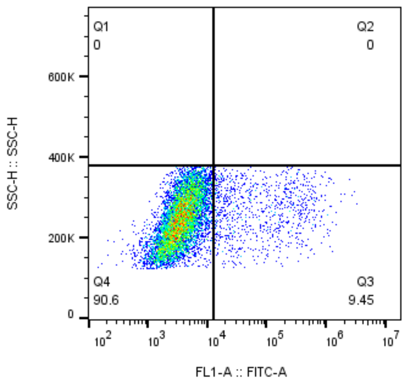


MPI30-1.6nM
HEK
9546

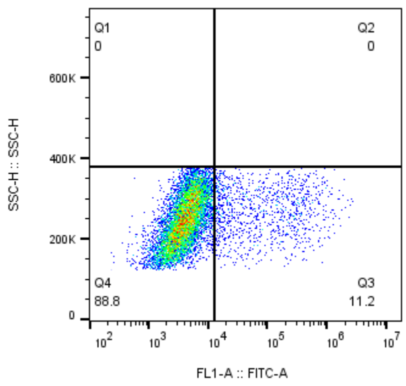


MPI30-0.32nM
HEK
9704

MPI30

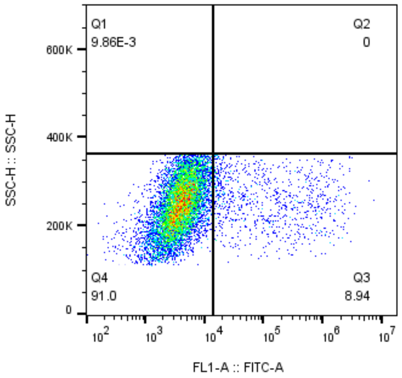


MPI31-10uM
HEK
10144

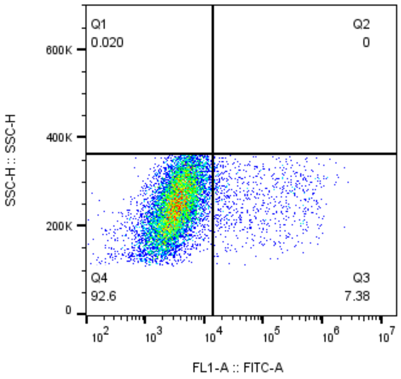


MPI31-2uM
HEK
10126

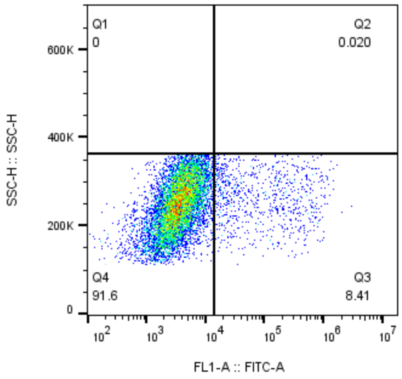


MPI31-40nM
HEK
9795

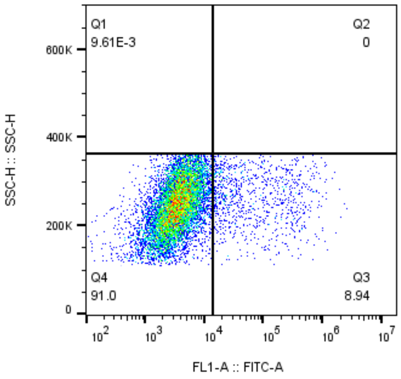


MPI31-8nM
HEK
10401

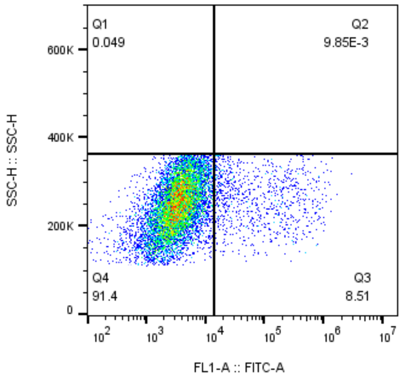


MPI31-1.6nM
HEK
10152

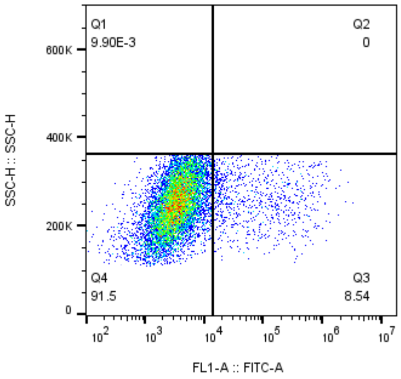


MPI31-0.32nM
HEK
10098

### MPI31

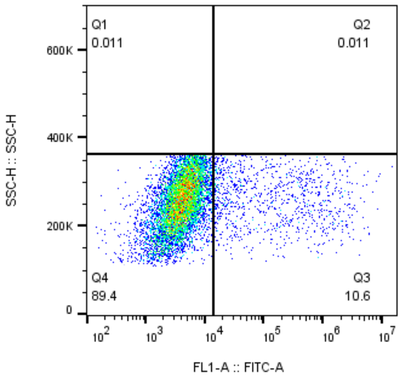


MPI32-10uM
HEK
9482

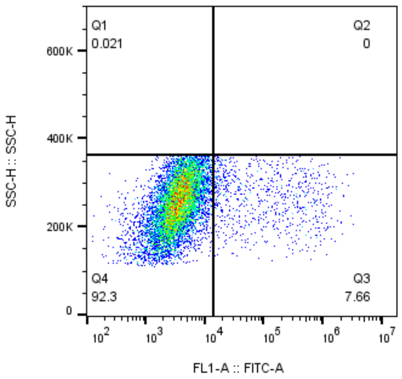


MPI32-2uM
HEK
9732

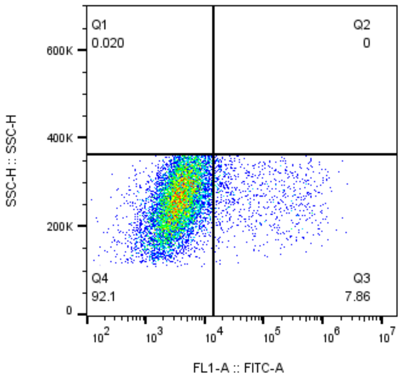


MPI32-40nM
HEK
9856

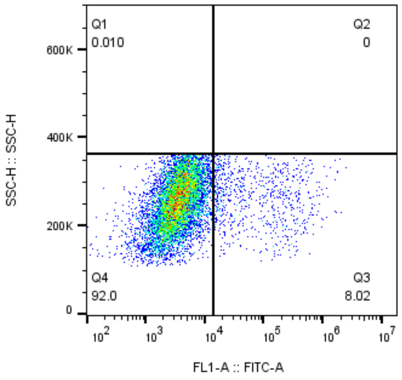


MPI32-8nM
HEK
9641

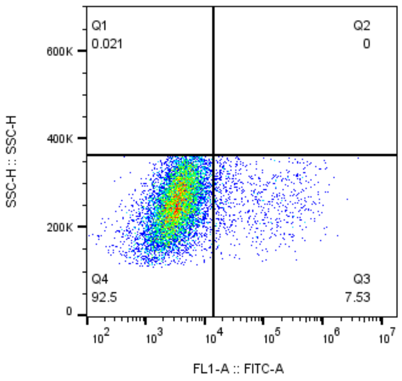


MPI32-1.6nM
HEK
9592

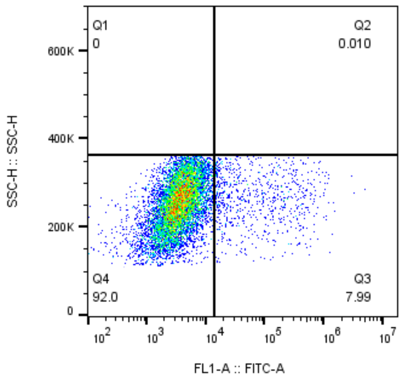


MPI32-0.32nM
HEK
9660

### MPI32

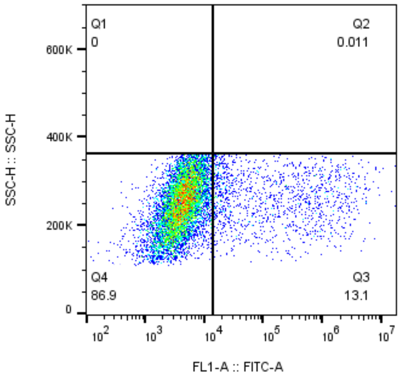


MPI33-10uM
HEK
9514

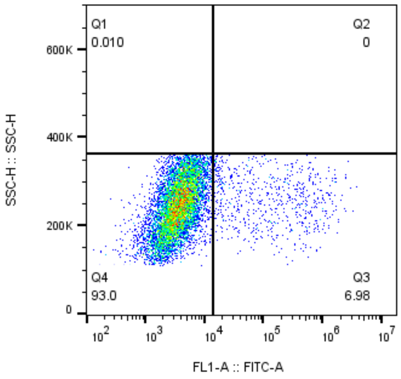


MPI33-2uM
HEK
9534

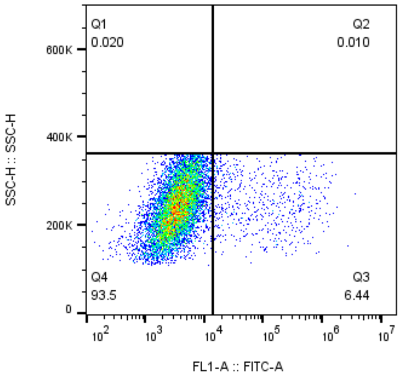


MPI33-40nM
HEK
9776

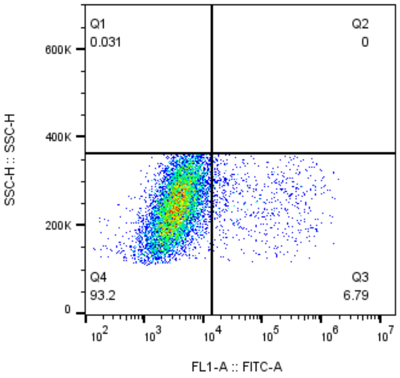


MPI33-8nM
HEK
9752

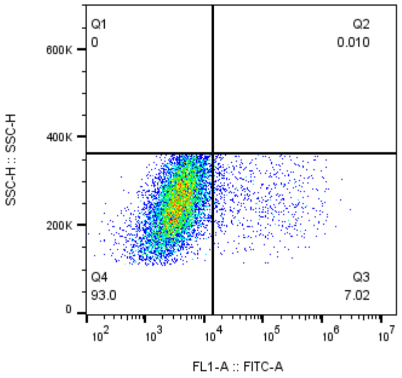


MPI33-1.6nM
HEK
9704

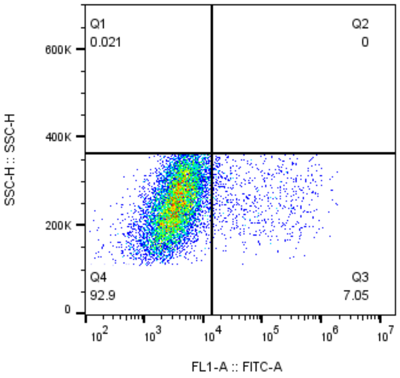


MPI33-0.32nM
HEK
9444

### MPI33

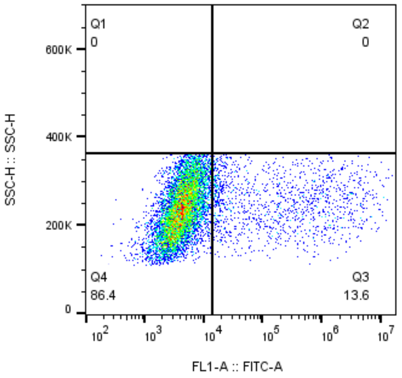


MPI34-10uM
HEK
9581

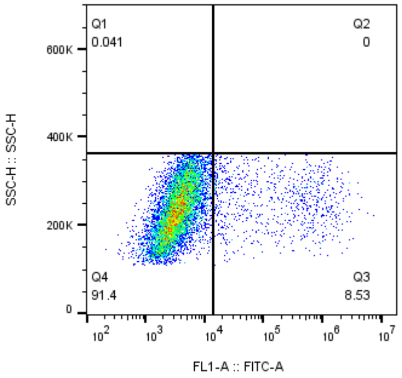


MPI34-2uM
HEK
9837

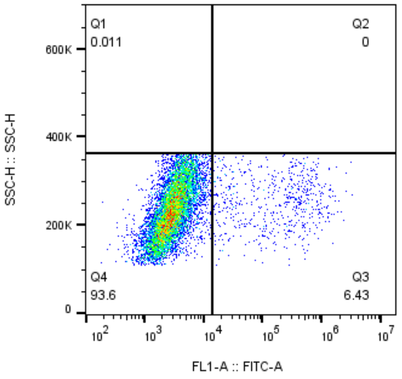


MPI34-40nM
HEK
9212

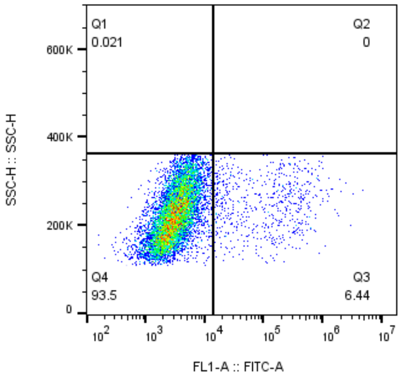


MPI34-8nM
HEK
9520

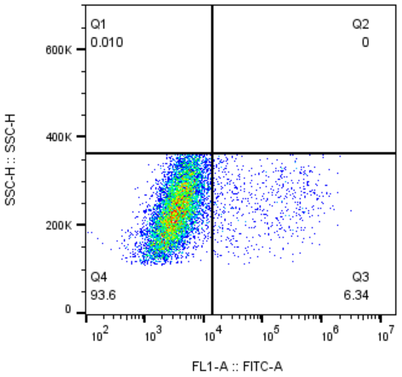


MPI34-1.6nM
HEK
9730

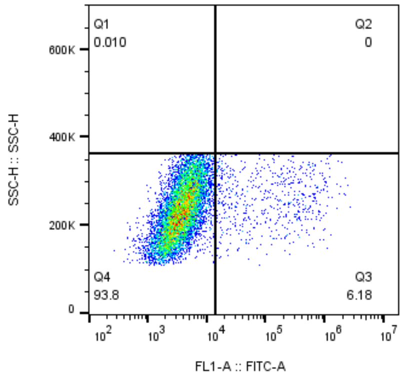


MPI34-0.32nM
HEK
9758

### MPI34

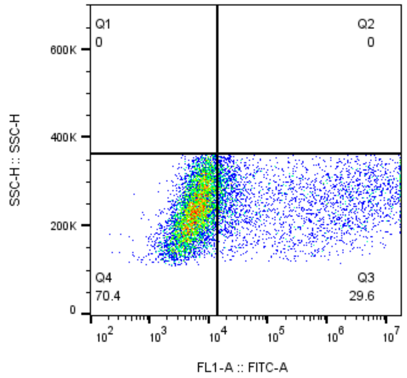


MPI35-10uM
HEK
9322

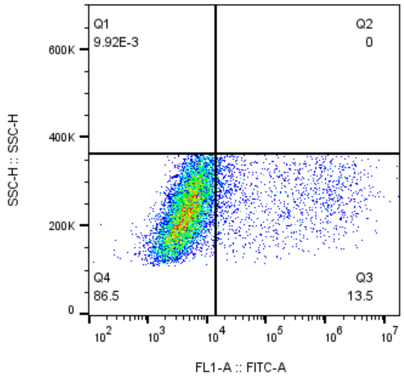


MPI35-2uM
HEK
10080

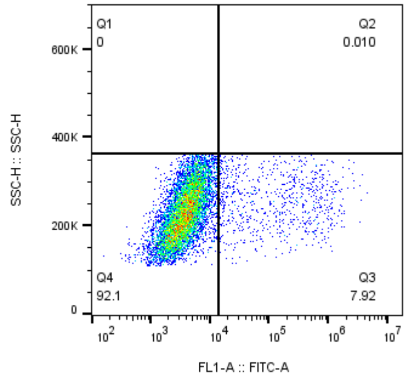


MPI35-40nM
HEK
9561

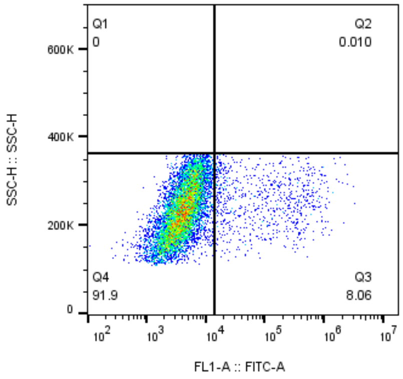


MPI35-8nM
HEK
9723

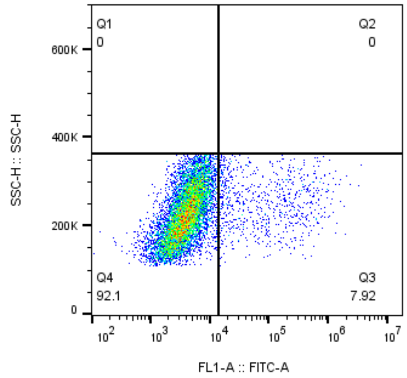


MPI35-1.6nM
HEK
9447

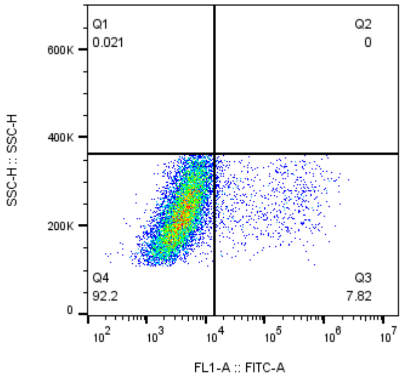


MPI35-0.32nM
HEK
9592

### MPI35

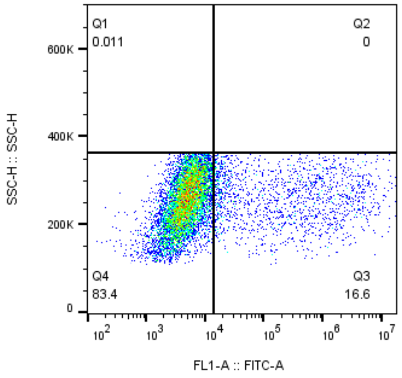


MPI36-10uM
HEK
9346

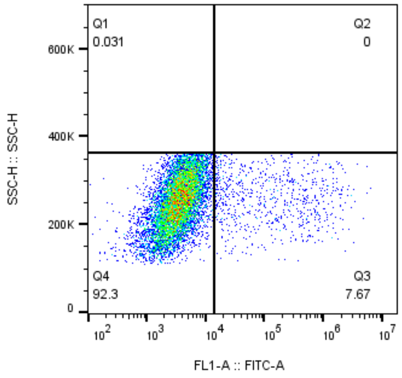


MPI36-2uM
HEK
9580

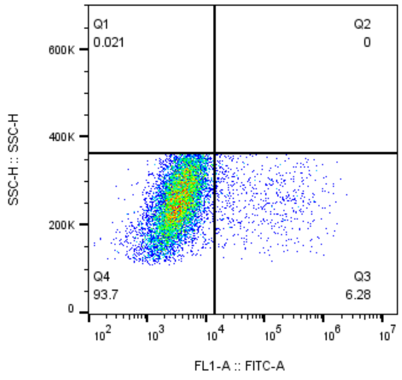


MPI36-40nM
HEK
9452

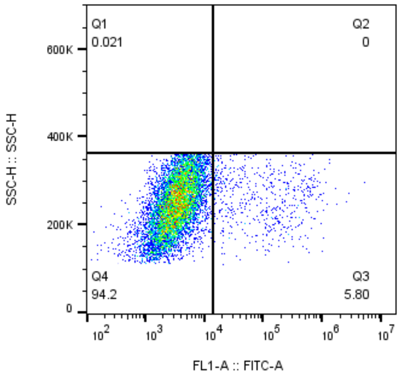


MPI36-8nM
HEK
9314

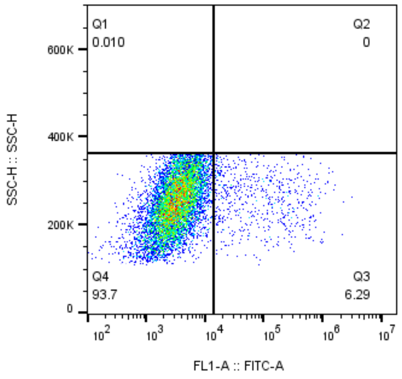


MPI36-1.6nM
HEK
9580

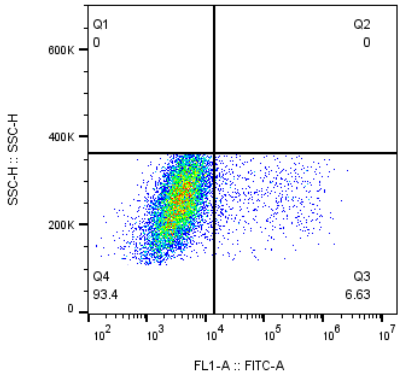


MPI36-0.32nM
HEK
9599

MPI36

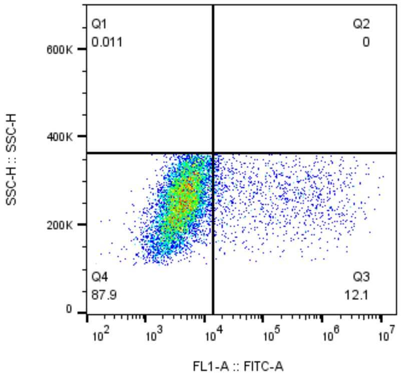


MPI37-10uM
HEK
9024

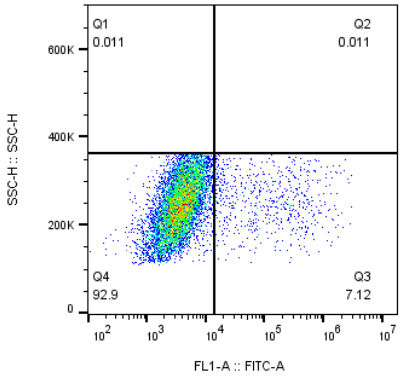


MPI37-2uM
HEK
8838

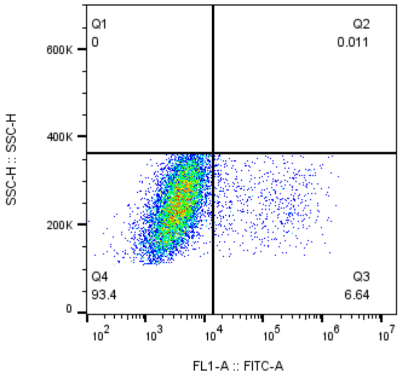


MPI37-40nM
HEK
9026

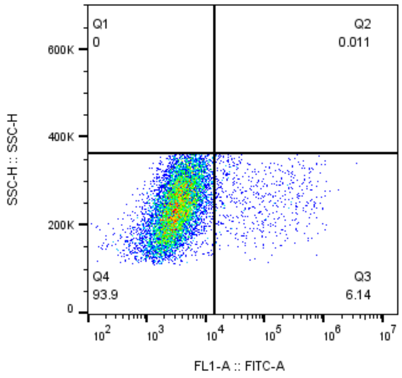


MPI37-8nM
HEK
9302

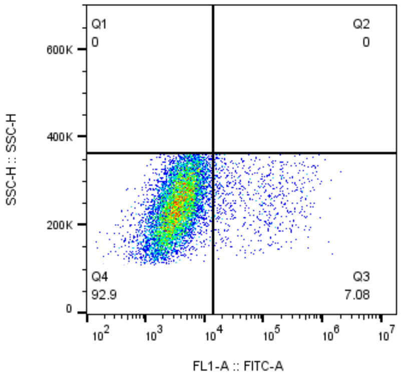


MPI37-1.6nM
HEK
9550

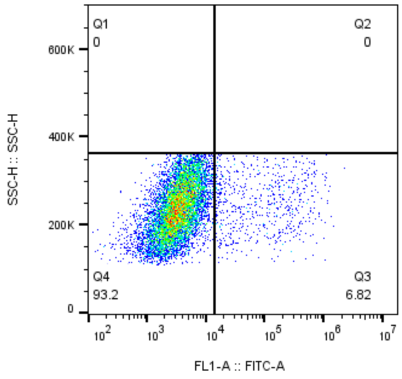


MPI37-0.32nM
HEK
9533

MPI37

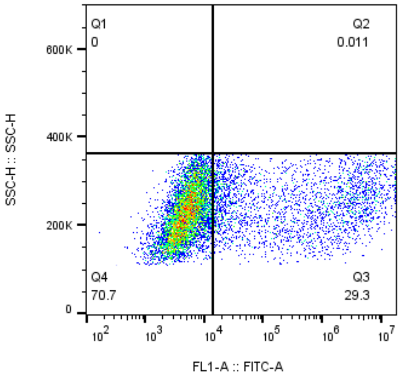


MPI38-10uM
HEK
9383

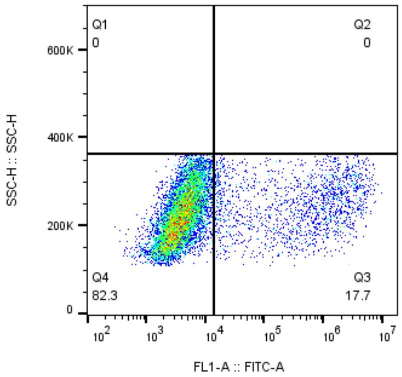


MPI38-2uM
HEK
9829

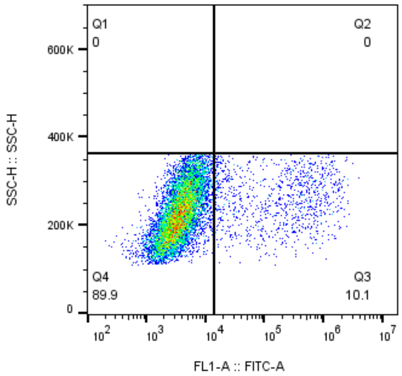


MPI38-40nM
HEK
10007

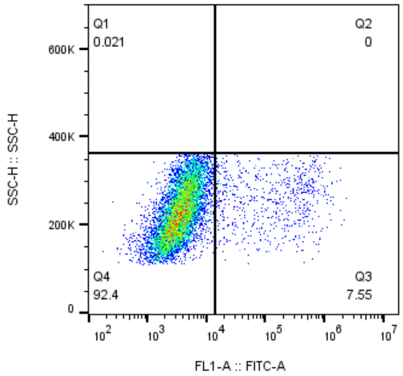


MPI38-8nM
HEK
9365

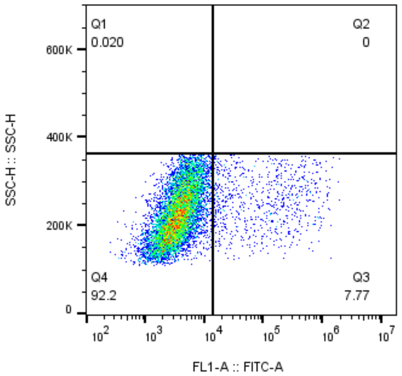


MPI38-1.6nM
HEK
9783

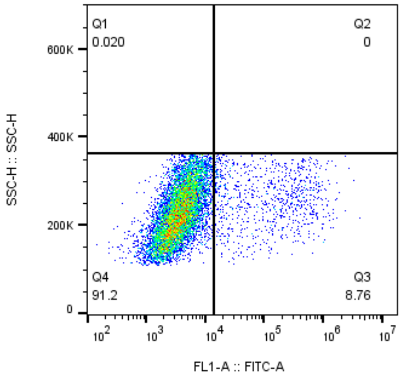


MPI38-0.32nM
HEK
10013

### MPI38

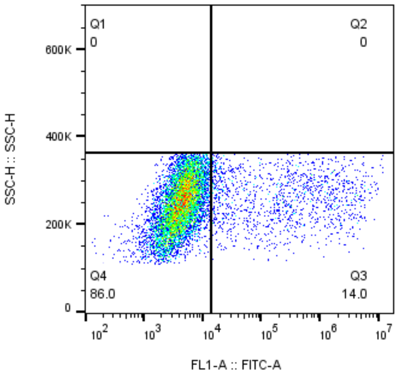


MPI39-10uM
HEK
9458

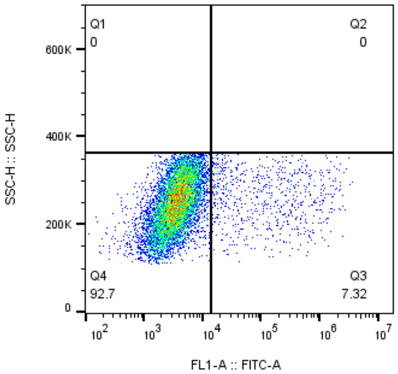


MPI39-2uM
HEK
9850

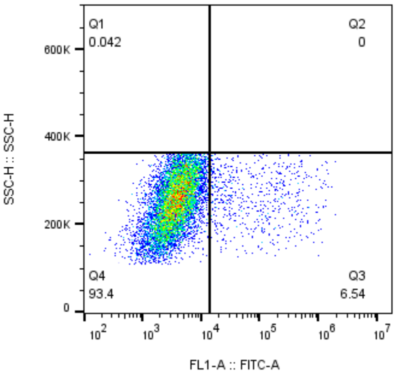


MPI39-40nM
HEK
9456

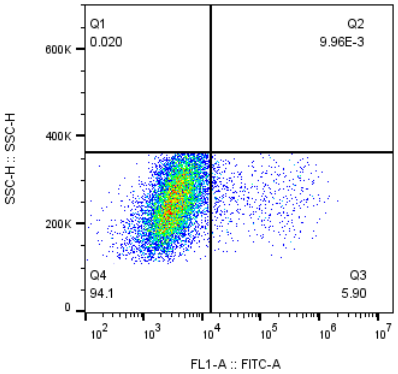


MPI39-8nM
HEK
10043

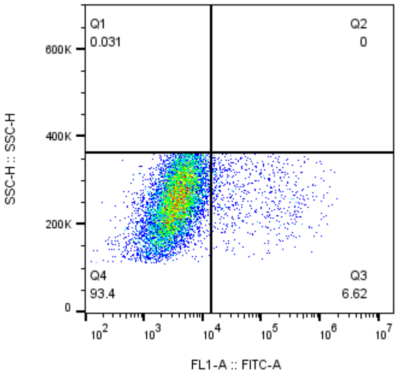


MPI39-1.6nM
HEK
9780

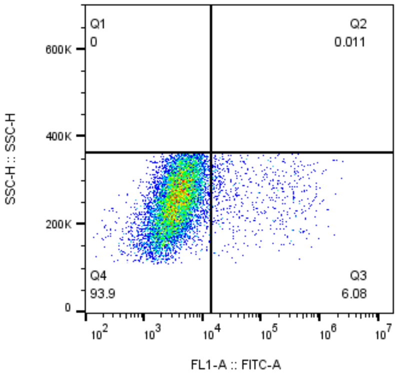


MPI39-0.32nM
HEK
9501

MPI39

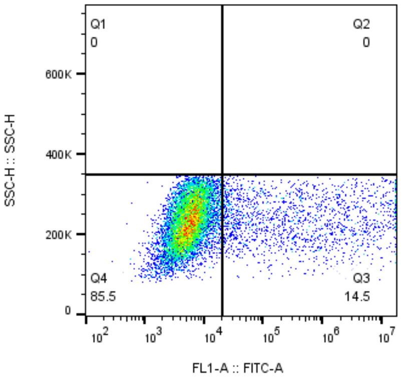


MPI40-10uM
HEK
11632

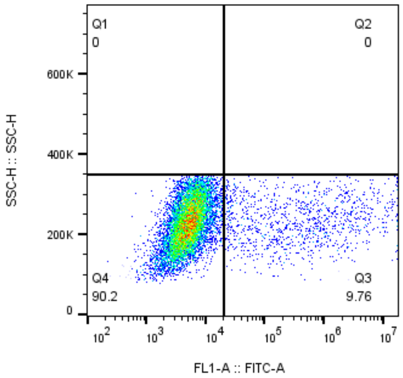


MPI40-2uM
HEK
11442

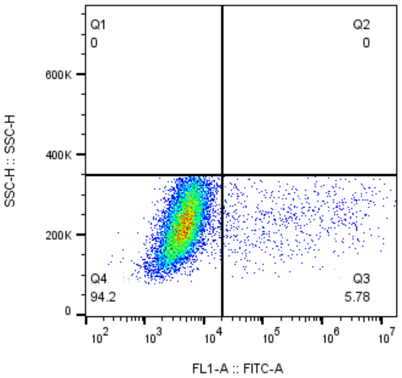


MPI40-40nM
HEK
11342

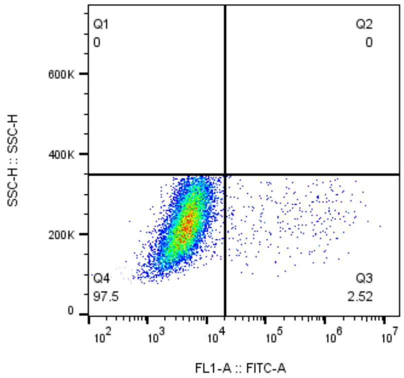


MPI40-8nM
HEK
11317

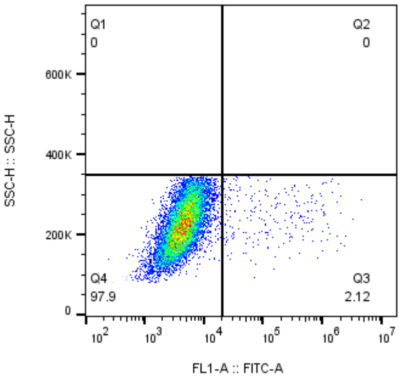


MPI40-1.6nM
HEK
11484

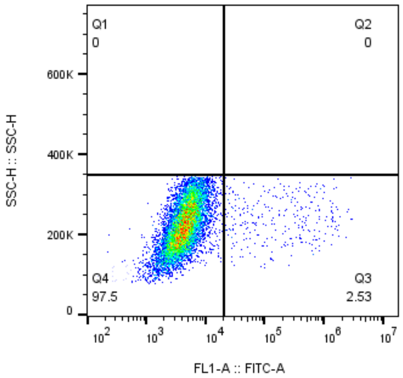


MPI40-0.32nM
HEK
11492

### MPI40

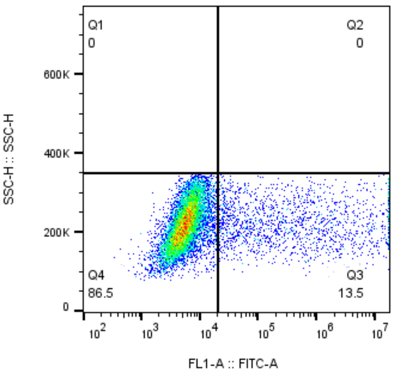


MPI41-10uM
HEK
11813

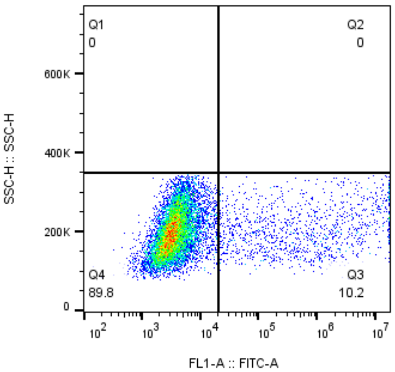


MPI41-2uM
HEK
11591

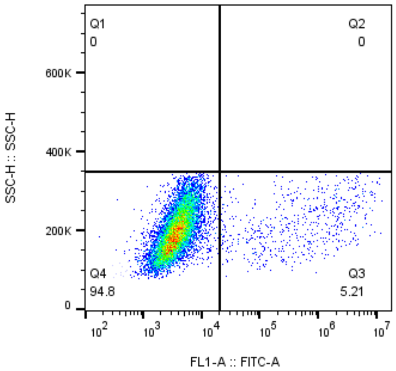


MPI41-40nM
HEK
11585

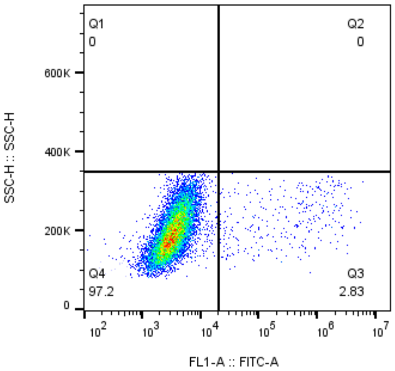


MPI41-8nM
HEK
11476

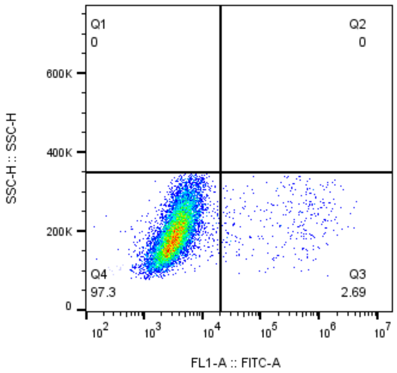


MPI41-1.6nM
HEK
11463

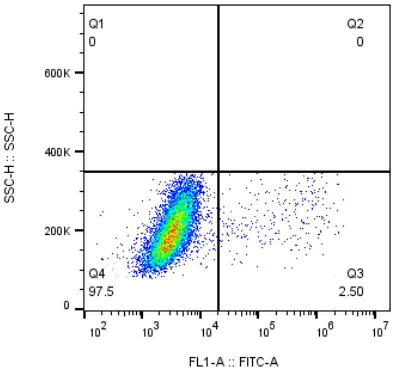


MPI41-0.32nM
HEK
11418

### MPI41

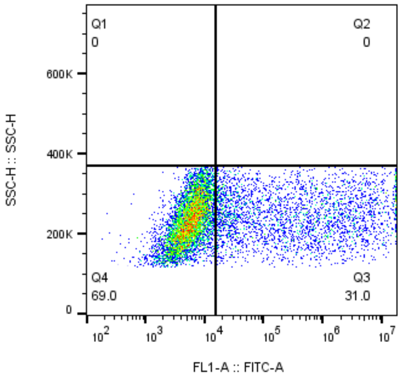


MPI42-10uM
HEK
9620

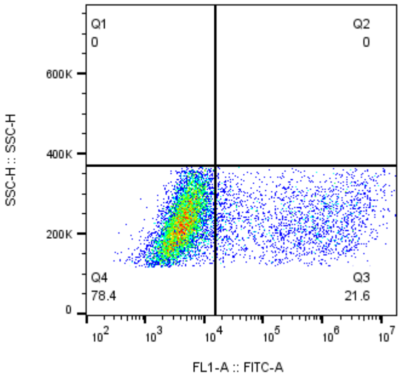


MPI42-2uM
HEK
9371

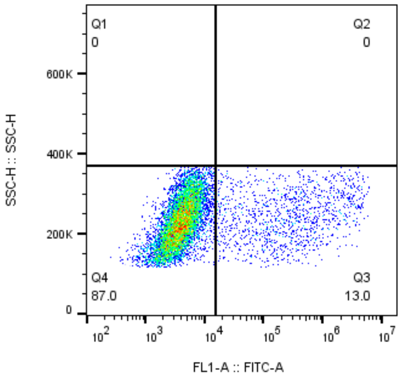


MPI42-40nM
HEK
9573

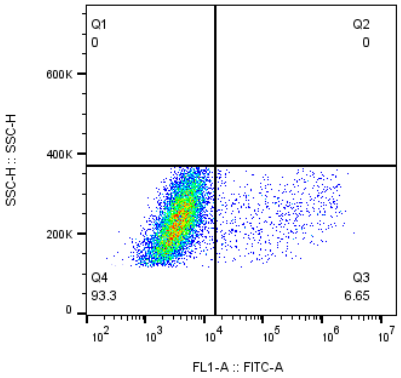


MPI42-8nM
HEK
9457

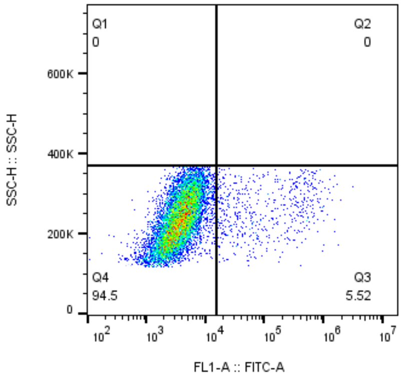


MPI42-1.6nM
HEK
9747

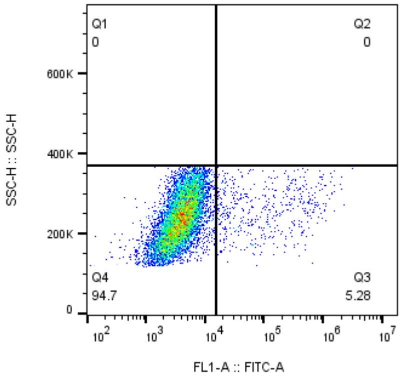


MPI42-0.32nM
HEK
9591

MPI42

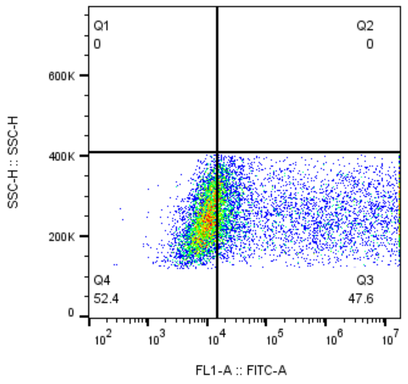


MPI43-10uM
HEK
9520

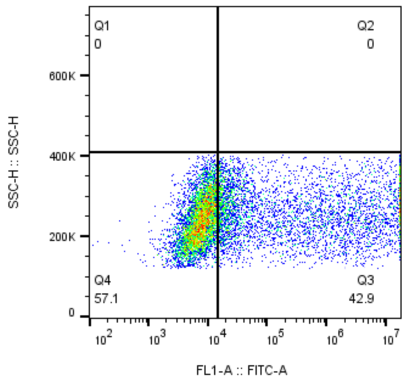


MPI43-2uM
HEK
9906

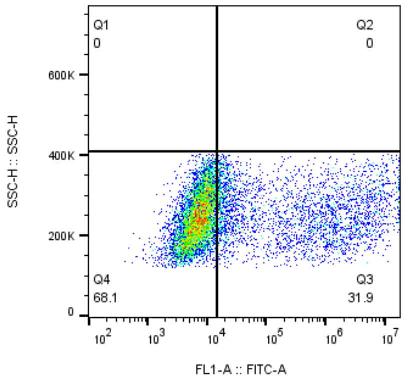


MPI43-40nM
HEK
9707

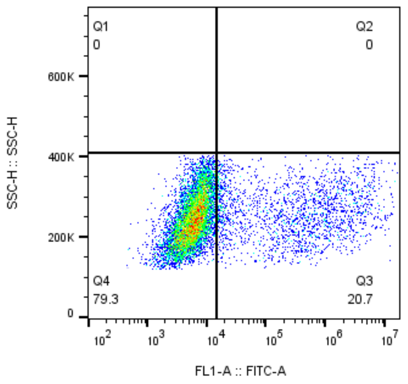


MPI43-8nM
HEK
9950

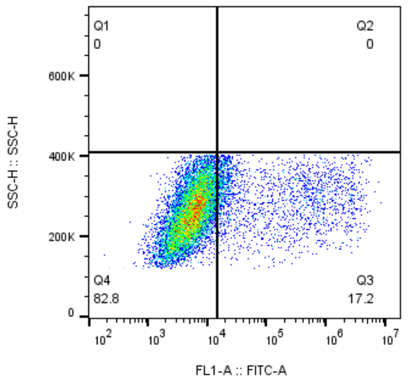


MPI43-1.6nM
HEK
11940

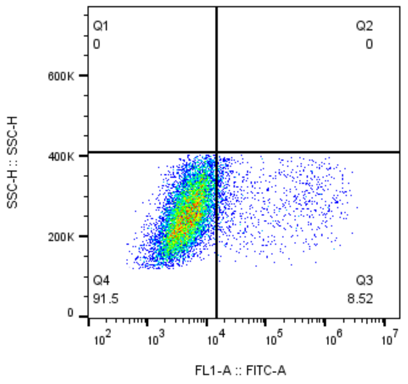


MPI43-0.32nM
HEK
10984

### MPI43

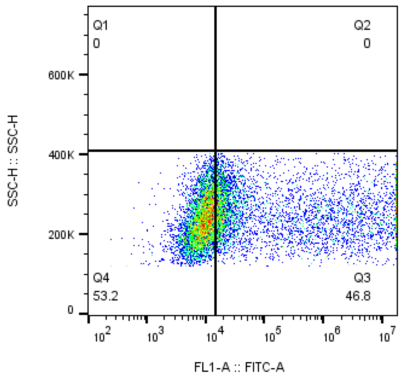


MPI44-10uM
HEK
9688

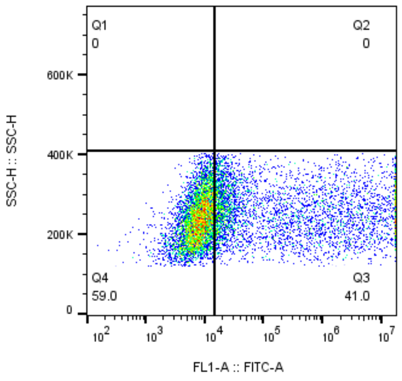


MPI44-2uM
HEK
10140

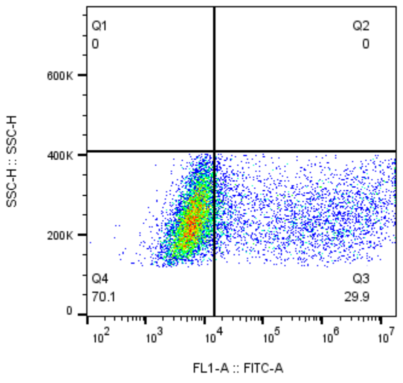


MPI44-40nM
HEK
9920

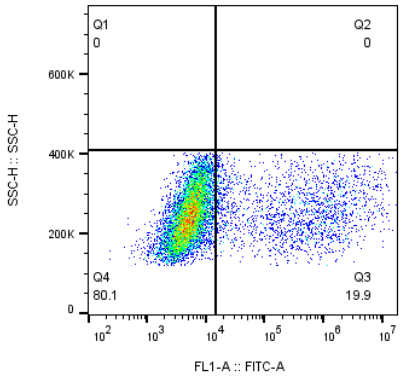


MPI44-8nM
HEK
10035

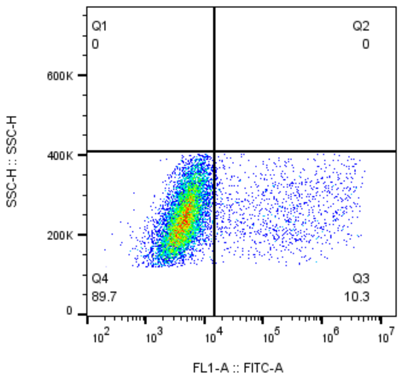


MPI44-1.6nM
HEK
9748

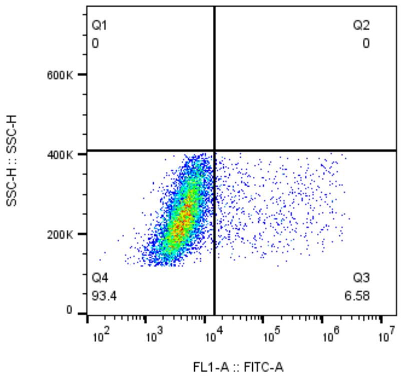


MPI44-0.32nM
HEK
9866

### MPI44

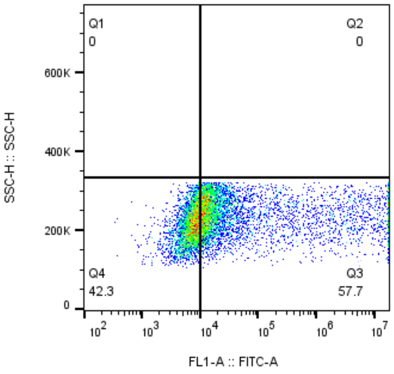


MPI45-10uM
HEK
9060

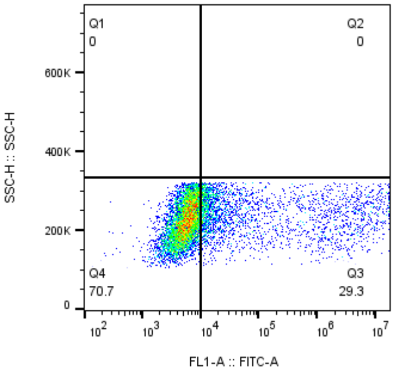


MPI45-2uM
HEK
9108

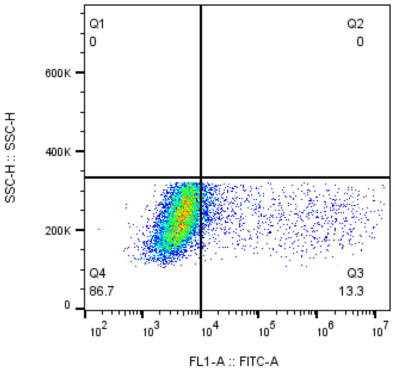


MPI45-40nM
HEK
8912

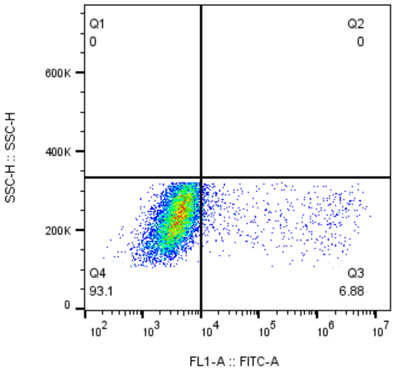


MPI45-8nM
HEK
8842

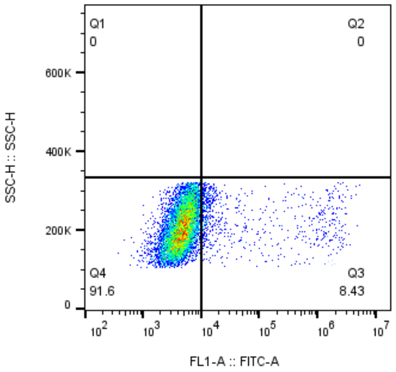


MPI45-1.6nM
HEK
9202

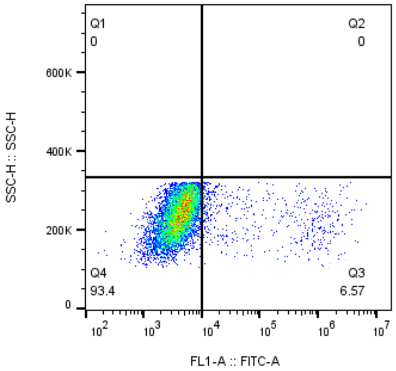


MPI45-0.32nM
HEK
8627

MPI45

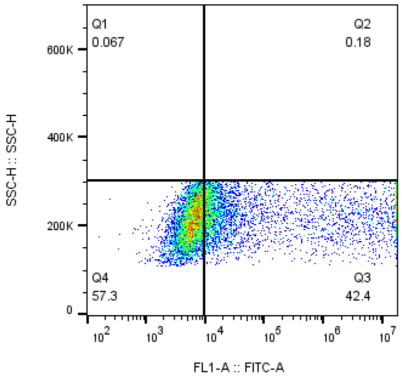


MPI46-10uM
HEK
9009

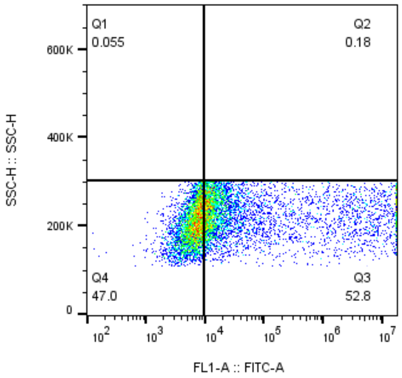


MPI46-2uM
HEK
9117

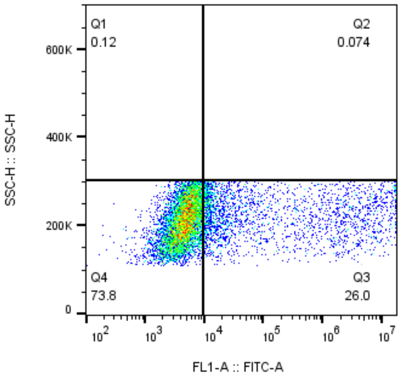


MPI46-40nM
HEK
9445

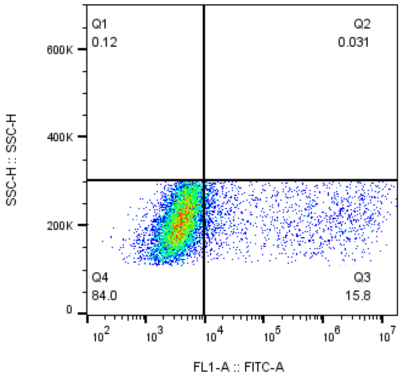


MPI46-8nM
HEK
9708

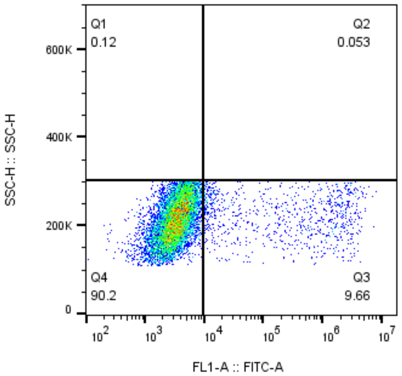


MPI46-1.6nM
HEK
9407

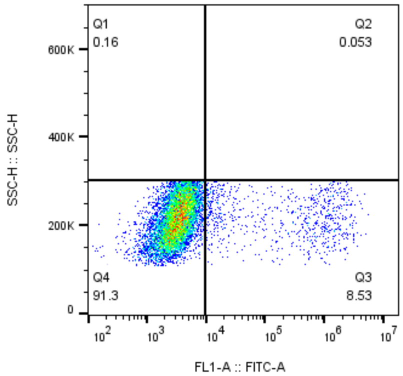


MPI46-0.32nM
HEK
9503

MPI46

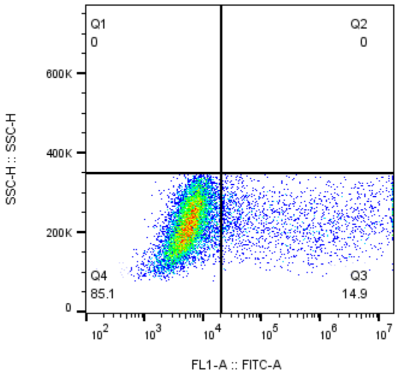


MPI47-10uM
HEK
11639

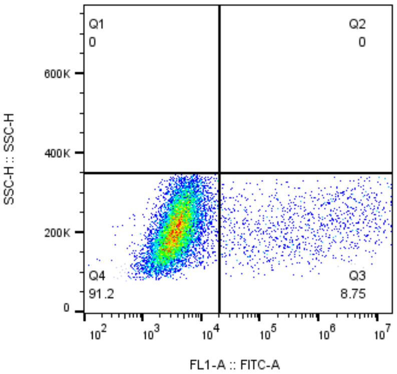


MPI47-2uM
HEK
11548

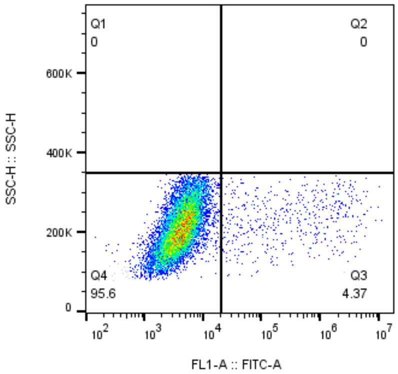


MPI47-40nM
HEK
11664

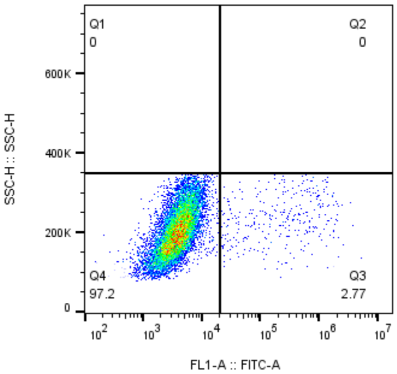


MPI47-8nM
HEK
11700

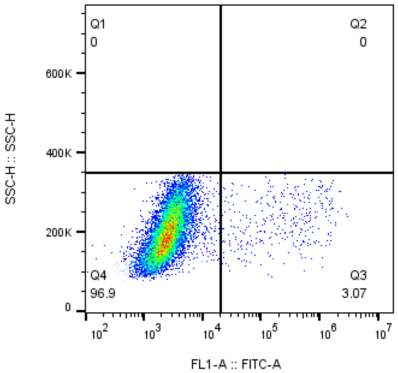


MPI47-1.6nM
HEK
11788

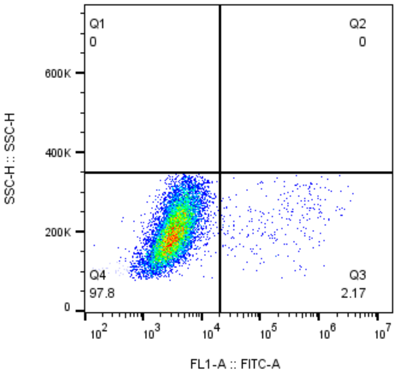


MPI47-0.32nM
HEK
11704

MPI47

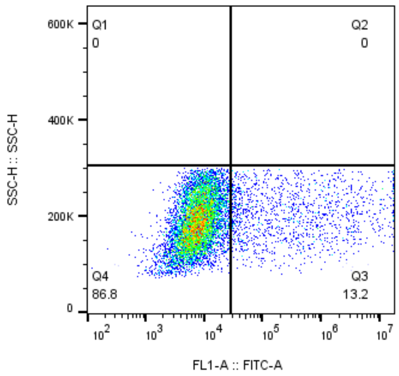


PF-07321332-10uM
HEK
9830

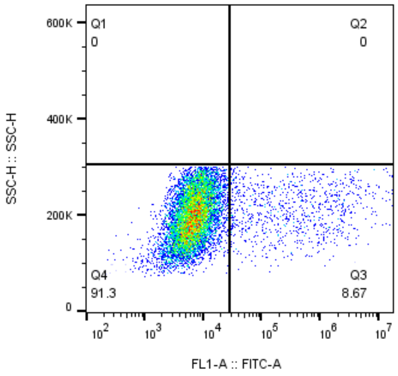


PF-07321332-2uM
HEK
10225

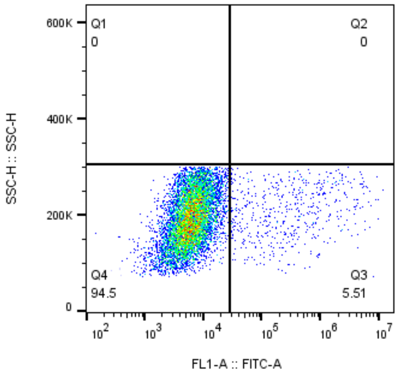


PF07321332-40nM
HEK
9545

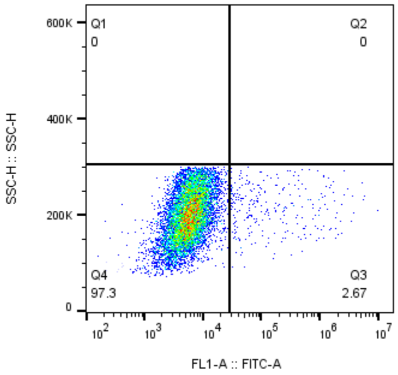


PF07321332-8nM
HEK
10172

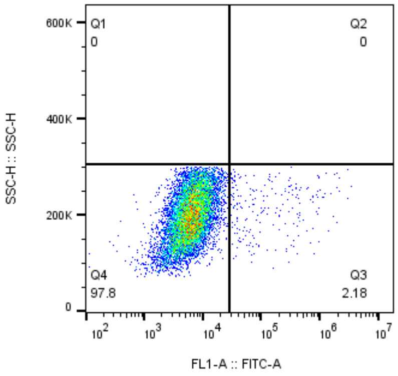


PF07321332-1.6nM
HEK
9658

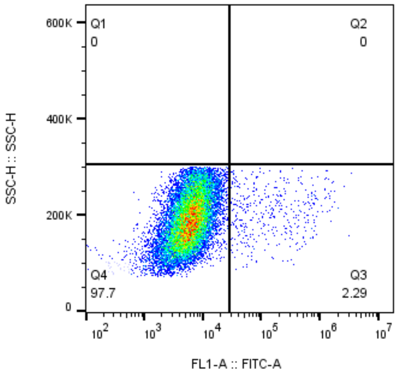


PF07321332-0.32nM
HEK
17273
